# Insights into the genetic architecture of the human face

**DOI:** 10.1101/2020.05.12.090555

**Authors:** Julie D. White, Karlijne Indencleef, Sahin Naqvi, Ryan J. Eller, Jasmien Roosenboom, Myoung Keun Lee, Jiarui Li, Jaaved Mohammed, Stephen Richmond, Ellen E. Quillen, Heather L. Norton, Eleanor Feingold, Tomek Swigut, Mary L. Marazita, Hilde Peeters, Greet Hens, John R. Shaffer, Joanna Wysocka, Susan Walsh, Seth M. Weinberg, Mark D. Shriver, Peter Claes

**Affiliations:** Department of Anthropology, Pennsylvania State University, State College, PA, 16802, USA; Department of Electrical Engineering, ESAT/PSI, KU Leuven, Leuven, 3000, Belgium; Medical Imaging Research Center, UZ Leuven, Leuven, 3000, Belgium; Department of Otorhinolaryngology, KU Leuven, Leuven, 3000, Belgium; Department of Chemical and Systems Biology, Stanford University School of Medicine, Stanford, CA, 94305, USA; Department of Genetics, Stanford University School of Medicine, Stanford, CA, 94305, USA; Department of Biology, Indiana University Purdue University Indianapolis, Indianapolis, IN, 46202, USA; Department of Oral Biology, Center for Craniofacial and Dental Genetics, University of Pittsburgh, Pittsburgh, PA, 15261, USA; Department of Developmental Biology, Stanford University School of Medicine, Stanford, CA, 94305, USA; Applied Clinical Research and Public Health, School of Dentistry, Cardiff University, Cardiff, CF10 3AT, United Kingdom; Department of Internal Medicine, Section of Molecular Medicine, Wake Forest School of Medicine, Winston-Salem, NC, 27101, USA; Center for Precision Medicine, Wake Forest School of Medicine, Winston-Salem, NC, 27101, USA; Department of Anthropology, University of Cincinnati, Cincinnati, OH, 45221, USA; Department of Human Genetics, University of Pittsburgh, Pittsburgh, PA, 15261, USA; Department of Human Genetics, KU Leuven, Leuven, 3000, Belgium; Department of Anthropology, University of Pittsburgh, Pittsburgh, PA, 15261, USA; Murdoch Children’s Research Institute, Melbourne, Victoria, 3052, Australia; Department of Biomedical Engineering, University of Oxford, Oxford, OX1 2JD, United Kingdom

## Abstract

The human face is complex and multipartite, and characterization of its genetic architecture remains intriguingly challenging. Applying GWAS to multivariate shape phenotypes, we identified 203 genomic regions associated with normal-range facial variation, 117 of which are novel. The associated regions are enriched for both genes relevant to craniofacial and limb morphogenesis and enhancer activity in cranial neural crest cells and craniofacial tissues. Genetic variants grouped by their contribution to similar aspects of facial variation show high within-group correlation of enhancer activity, and four SNP pairs display evidence of epistasis, indicating potentially coordinated actions of variants within the same cell types or tissues. In sum, our analyses provide new insights for understanding how complex morphological traits are shaped by both individual and coordinated genetic actions.

## Main Text

> *“One of the major problems confronting modern biology is to understand how complex morphological structures arise during development and how they are altered during evolution”*
>
> *Atchley and Hall, 1991*^*1*^, *p*.*102*

The ‘problem’ described by Atchley and Hall continues to enthrall biologists, geneticists, anthropologists, and clinicians almost three decades later. In their review, the authors describe a “complicated developmental choreography” in which intrinsic genetic factors, epigenetic factors, and interactions between the two make up the progeny genotype, which engages with the environment to ultimately produce a complex morphological trait, defined thus by its composition from a number of separate component parts^1^. We now understand that the intrinsic genetic factors ultimately contributing to complex morphological traits consist not only of single variants altering protein structure and/or function, but also non-coding variants and interactions among variants, each affecting multiple tissues and developmental timepoints. This realization necessitates the development and utilization of methods capable of describing the genetic architecture of complex morphological traits, which includes identifying the individual genetic variants contributing to morphological variation as well as their interactions^2,3^.

The human face is an exemplar complex morphological structure. It is a highly multipartite structure resulting from the intricate coordination of genetic, cellular, and environmental factors^4–6^. Through prior genetic association studies of quantitative traits, 51 loci have been implicated in normal-range craniofacial morphology, and an additional 50 loci have been associated with self-reported nose size or chin dimples in a large cohort study^7^ (Table S1). However, as with all complex morphological traits, our ability to identify and describe the genetic architecture of the face is limited by our ability to accurately characterize its phenotypic variation^4^, identify variants of both large and small effect^8^, and identify interactions between variants. We previously described a novel data-driven approach to facial phenotyping, which facilitated the identification and replication of 15 loci involved in global-to-local variation in facial morphology^9^. Here, we apply this phenotyping approach to two much larger cohorts from the US and UK (*n* = 8,246; Table S2) and use advanced multivariate techniques to uncover new biological insights into the genetic architecture of the human face. We now identify 203 signals, located in 138 cytogenetic bands, associated with normal-range facial morphology (Fig. 1). Many of these loci harbor genes involved in craniofacial syndromes, which we show also affect normal facial variation, and a large number are novel signals, potentially pointing to previously unknown genes and pathways involved in normal and abnormal facial development. Using bioinformatics tools and ChIP-seq databases for epigenomic characterization, we show that variants at our GWAS peaks are involved in regulating enhancer activity in cell types controlling facial morphogenesis across the developmental timeline. Furthermore, we reveal interactions between variants at different loci affecting similar aspects of facial shape variation, exposing some of the hidden assemblages of genes that work in concert to build human faces. With this work, we not only push forward our understanding of the human face, but also illustrate the potential for researchers to confront Atchley and Hall’s problem, by intensively characterizing complex morphological variation and using advanced methods to identify factors involved in the developmental choreography of complex morphological structures.

**Fig. 1.**
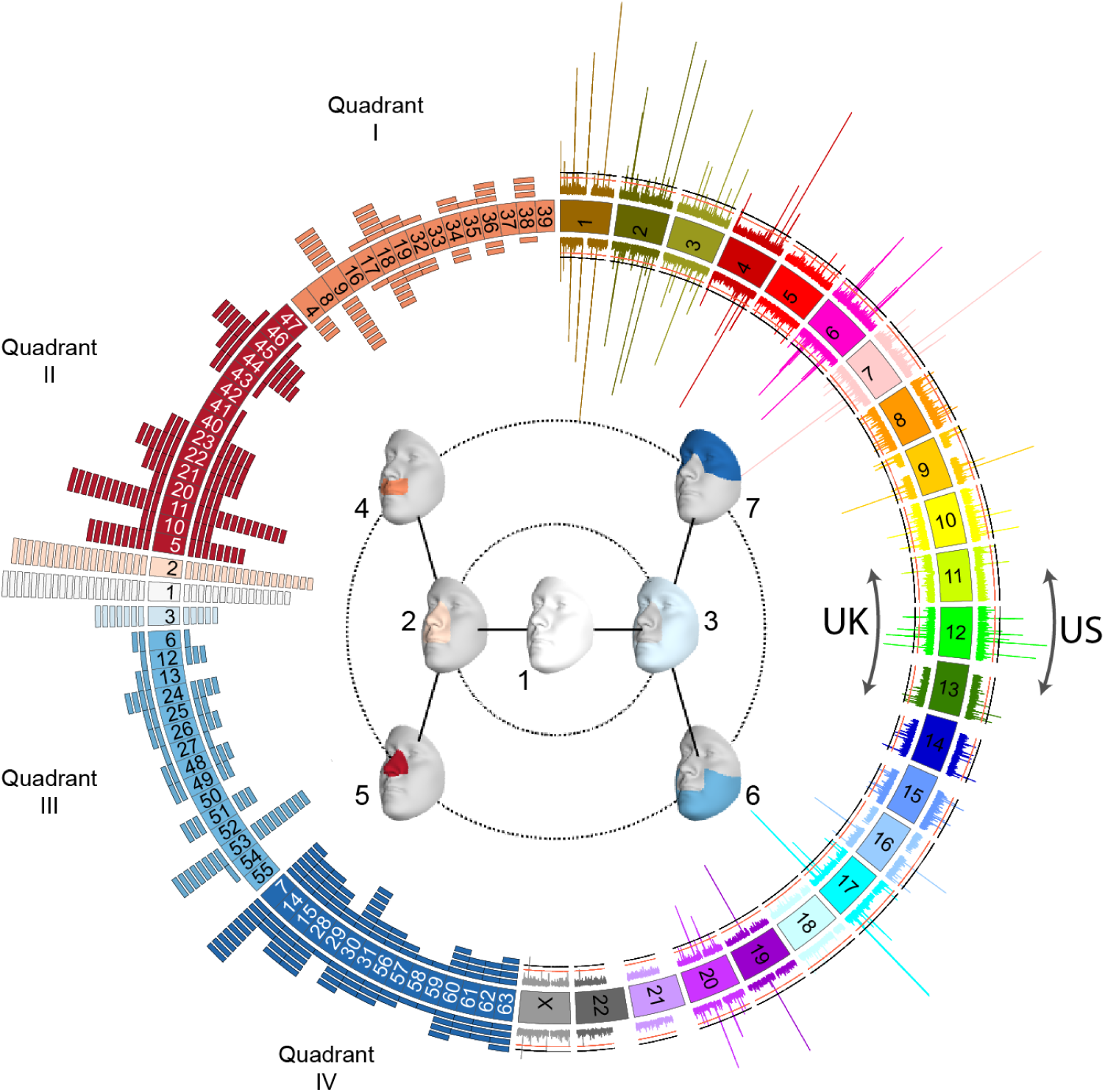
Overall results of US-driven and UK-driven meta-analyses. On the left, blocks representing the facial segments arranged and colored according to quadrant (I = orange; II = red; III = light blue; IV = dark blue), and the full face (white), and segments 2 (light orange) and 3 (ice blue). The histogram arranged on the left side represents the number of lead SNPs reaching their lowest p-value in each segment, colored by quadrant, with the US-driven meta-analysis results on the outside circle and the UK-driven meta-analysis results on the inside circle. In the center, the first three levels of the facial segmentation [segments 1 - 7], also colored to match with the quadrants on the left. On the right, a Miami plot of the US-driven meta-analysis p-values on the outside and the UK-driven meta-analysis p-values on the inside, with chromosomes colored and labeled. P-values are -log10 scaled (range: [0-80]). The red line represents the genome-wide significance threshold (*p* = 5 × 10^−8^) and the black line represents the study-wide threshold (*p* = 6.25 × 10^−10^). Created using Circos v0.69-8^101^.

## Multivariate phenotyping and meta-analysis framework

To study facial variation at both global and local scales, we start with a set of 3D facial surface scans, upon which we map a dense mesh of homologous vertices^10^. We then apply a data-driven facial segmentation approach, defined by grouping vertices that are strongly correlated using hierarchical spectral clustering^9^. The configurations of each of the resulting 63 segments are then independently subjected to a Generalized Procrustes analysis, after which principal components analysis is performed in conjunction with parallel analysis to capture the major phenotypic variation in each facial segment^11^ (Fig. S1). Within each segment, instead of *a priori* selecting the principal components (PCs) of interest, we use canonical correlation analysis (CCA) to first identify the linear combination of components maximally correlated with the SNP being tested, under an additive genetic model, in the identification cohort (we call this combination of PCs the “phenotypic trait”). Subsequently, the verification cohort is projected onto each of these learned phenotypic traits, creating univariate phenotypic variables which are then tested for genotype-phenotype associations. The identification and verification p-values are then meta-analyzed using Stouffer’s method^12,13^. The whole process is then repeated, switching the dataset used for identification and verification, thereby resulting in two sets of meta-analysis p-values from each permutation of identification and verification (p_META-US_ and p_META-UK_; Fig. S2).

We first assessed the degree to which variation in each facial segment shares the same patterns of genetic association across the genome by computing the Spearman correlation among all genetic association results for each pair of facial segments. We then used these pairwise correlations, which we verified were not due to linkage disequilibrium either across the genome or with genome-wide significant SNPs (Fig. S3), to both hierarchically cluster the facial segments and visualize between-segment correlations (Fig. S4). These analyses revealed two key features of the global association patterns. First, the correlations were highest between segments of the same facial quadrant (i.e. lips, nose, lower face, upper face), validating the hierarchical clustering used to initially define the segments. Clustering the facial segments based on the genetic association results resulted in four main clusters, each corresponding to segments from the same quadrant of the polar dendrogram (Fig. S5). Second, despite substantial within-quadrant similarity, there were notable correlations between groups of segments from different quadrants. Some of these specific correlations reflect close physical proximity of the segments in different quadrants (e.g. segment 51-19; Fig. S4), but some correlations seem to reflect the shared embryological origins of groups of segments. Specifically, segments representing the nose (Quadrant II) and upper face (Quadrant IV) cluster together, and segments representing the lips (Quadrant I) and lower face (Quadrant III) cluster together (Fig. S5). Quadrants II and IV together approximate the frontonasal prominence, which appears earlier in development than the mandibular and maxillary prominences, which are approximated by Quadrants I and III^14^. Together, these results indicate that hierarchical spectral clustering of the face based on structural correlations effectively partitions underlying genetic signals into biologically coherent groups.

In total, we identified 17,612 SNPs with p-values (p_META-US_ and/or p_META-UK_) lower than the genome-wide threshold (*p* ≤ 5 × 10^−8^), and 11,319 SNPs with p-values lower than the study-wide Bonferroni threshold (*p* ≤ 6.25 × 10^−10^) (Fig. S6). For each peak, we designated the SNP with the lowest p-value across all facial segments as the “lead SNP,” refining our results to 218 lead SNPs, all below the genome-wide threshold. Of these, 203 showed consistent effects of the phenotypic trait identified in the US- and the UK-driven meta-analyses in the facial segment with the lowest p-value for that SNP (Fig 1; Table S3). Trait similarity was tested using a regression of slopes of each of the phenotypic traits found in the identification stage for each permutation, with results considered sufficiently similar if they were below a false discovery rate of *p* ≤ 3.66 × 10^−2^. The output of our meta-analyses is twofold. First, the meta-analyses p-values facilitate quantification of the statistical evidence of association between the SNPs and these discovered traits. In addition, for each SNP and in each segment, we allow the data to drive the identification of the phenotypic trait most associated with that SNP in the segment. This opens the possibility that SNPs could have associations with many different traits across segments, allowing us to better describe genetic influences on human facial morphology.

## Genes near lead SNPs are enriched for both craniofacial and limb development

Using FUMA^15^ and GREAT^16^ analyses, we established that genes located within 500 kb of the lead SNPs were highly enriched for processes and phenotypes associated with craniofacial development and morphogenesis in humans and mice (Fig. S7A). Notably, the top human phenotype was orofacial clefting, indicating a substantial overlap between the genes involved in normal facial variation and those implicated in the most common craniofacial birth defect in humans. Furthermore, many of the surrounding genes to which the lead SNPs were annotated are known to be involved in pathways relevant for craniofacial development, such as the WNT signaling and TGFB pathways (Fig. S7B). Our GWAS signals were also enriched for processes associated with limb development and related phenotypes, pointing to a shared genetic architecture between faces and limbs (Fig. S7A). A number of genes near our GWAS loci (e.g. *Dlx* homeobox genes, *BMPs*, and *FGFR2*) have well-established roles in limb development^17^. These findings are also supported by the large number of human syndromes that present with both facial and limb malformations^18^.

## Facial GWAS peaks are enriched for enhancers specific to cell types across the timeline of facial development

To assess the likely cell-types and developmental timepoints in which our GWAS regions are active, we used epigenomic mapping datasets generated from human cranial neural crest cells (CNCCs) and other cell types relevant to embryonic development. To gauge activity, we analyzed ChIP-seq signals of acetylation of histone H3 on lysine K27 (H3K27ac), which is a marker of the promoters of transcriptionally active genes and active distal enhancers^19,20^. We compiled H3K27ac ChIP-seq signals from approximately 100 different cell types and tissues, including CNCCs, fetal and adult osteoblasts, mesenchymal stem cell-derived chondrocytes, as well as dissected embryonic craniofacial tissues (Carnegie stages 13-20). Both CNCCs and craniofacial tissues showed the highest H3K27ac signals in the vicinity of the 203 lead SNPs, whereas no H3K27ac signal was observed for 203 random SNPs matched for allele frequency and distance to the nearest gene (Fig. 2A). These observations are consistent with an embryonic origin for human facial variation across the timeline of facial development, as CNCCs represent an early time point in facial development whereas the craniofacial tissues represent progressively later time points.

**Fig. 2.**
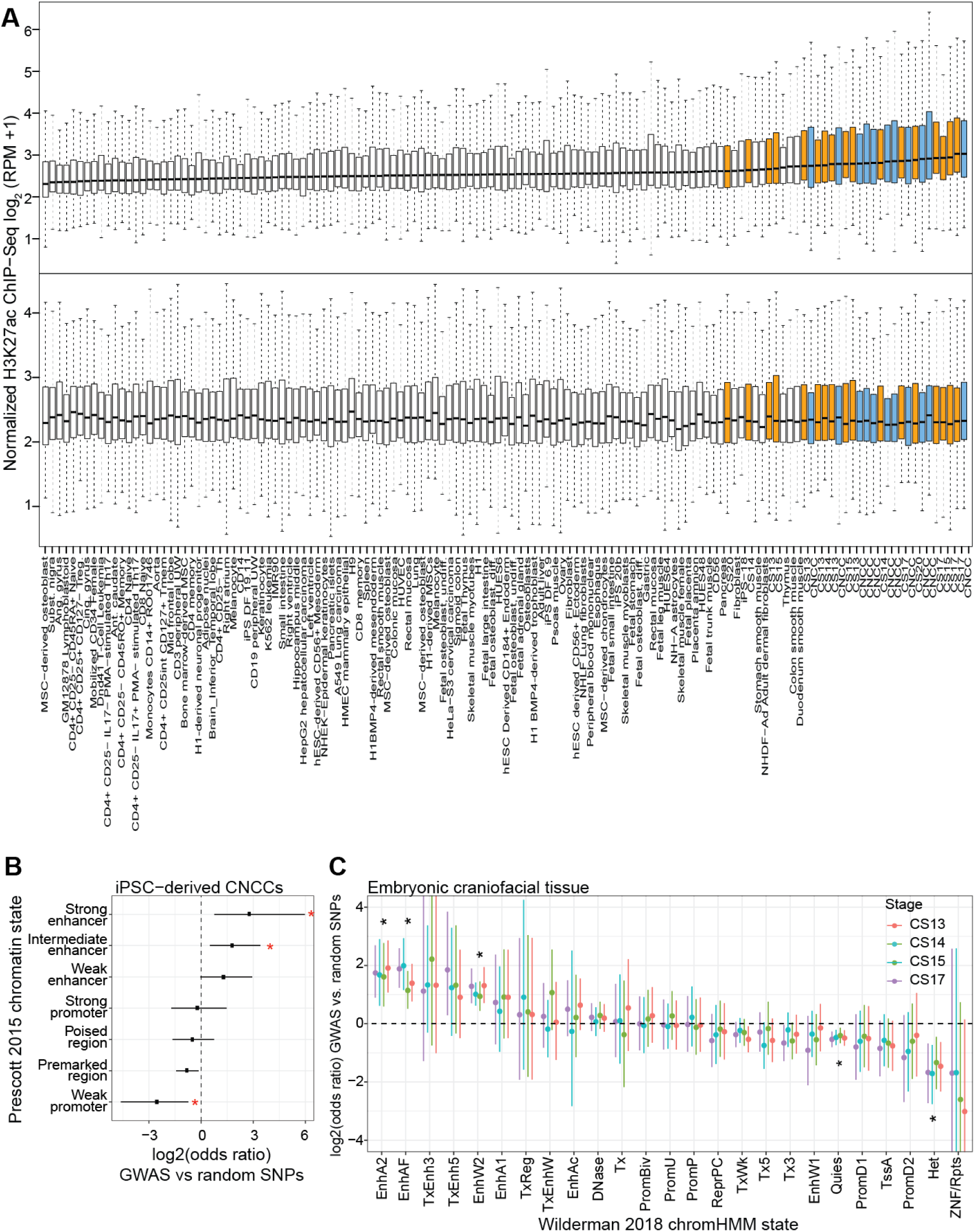
Regions near the 203 lead SNPs are enriched for enhancers preferentially active in cranial neural crest cells and embryonic craniofacial tissue. **(A)** Each boxplot represents the distribution of H3K27ac signal in 20 kb regions around the 203 lead SNPs (*top*) or 203 random SNPs (*bottom*) in one sample, with cranial neural crest cells and embryonic craniofacial tissue highlighted in blue and orange, respectively. For each class of regulatory element in either cranial neural crest cells (**B**) or embryonic craniofacial tissue (**C**), the number of elements within 10 kb of the 203 lead SNPs was compared to the number within 10 kb of 203 random SNPs by Fisher’s exact test. Points represent estimated odds ratio and 95% confidence intervals. Asterisk (*) indicates any adjusted p-value < 0.05. For embryonic craniofacial tissue, enrichments were calculated for each Carnegie stage separately, as Wilderman et al.^22^ performed chromatin state segmentation for each stage separately. Descriptions of mnemonics for significantly enriched or depleted chromHMM states are as follows: EnhA2, active enhancer 2; EnhAF, active enhancer flank; EnhW2, weak enhancer 2; Quies, quiescent/low; Het, heterochromatin. Descriptions of all mnemonics can be found at: https://egg2.wustl.edu/roadmap/web_portal/imputed.html#chr_imp.

H3K27ac marks activity at both coding and noncoding elements; to distinguish enrichment between the two, we examined chromatin signals in CNCCs and embryonic craniofacial tissues in more detail, using ChIP-seq data on additional chromatin marks and transcription factors for either cell-type^21,22^. In both CNCCs and craniofacial tissue at all sampled developmental stages, candidate regulatory regions in the vicinity of the 203 lead SNPs were significantly enriched for predicted enhancers (CNCCs: strong and intermediate enhancers; craniofacial tissue: active, flanking, and weak enhancers), and not promoters (Fig. 2B and C). This is an especially intriguing result, as recent evidence has described the action of multiple enhancers, each showing different tissue or timing specificity, in modulating expression levels to affect craniofacial development^23^. Thus, though promoters can have an equally important role in development, the enrichment of our lead SNPs in predicted enhancer regions signifies the likely importance of enhancers in regulating normal-range facial variation.

While CNCCs and craniofacial tissues showed the strongest enrichments when considering the 203 lead SNPs together, we hypothesized that cell-type-specific activity patterns could be used to further subdivide the 203 lead SNPs. We therefore clustered the 203 lead SNPs into six distinct groups on the basis of H3K27ac signal across all ∼100 cell types. As expected, the greatest fraction of lead SNPs showed specific activity for CNCCs and craniofacial tissue (e.g. cluster 5; Fig. 3); interestingly, however, some SNPs showed preferential activity for either CNCCs or craniofacial tissue (e.g. clusters 1 and 2; Fig. 3). Greater specificity for CNCCs could arise because CNCCs constitute a relatively small proportion of the cells present in craniofacial tissue at Carnegie stages 13-20, while greater specificity for craniofacial tissue could be due to activity in further differentiated cell-types of the face. Complementing our earlier results indicating that some genes near our GWAS peaks are involved in both facial and limb development, a subset of SNPs showed preferential activity in additional in vitro-derived cell types relevant to both the face and the rest of the skeletal system, including osteoblasts, chondrocytes, differentiating skeletal muscle myoblasts, fibroblasts, and keratinocytes (e.g. cluster 3; Fig. 3). As an illustration, lead SNP rs1367228 is located within a transcription factor binding sequence in an intron of the *EFEMP1* gene (2p16.1), and is associated in this study with facial variation around the eye sockets and upper cheekbones and showed preferential activity in fetal osteoblast cell lines and MSC-derived chondrocytes (Fig. S9). Together, these results suggest that genetic variation underlying facial morphology operates by modulating enhancer activity across multiple cell types throughout the timeline of embryonic facial development.

**Fig. 3.**
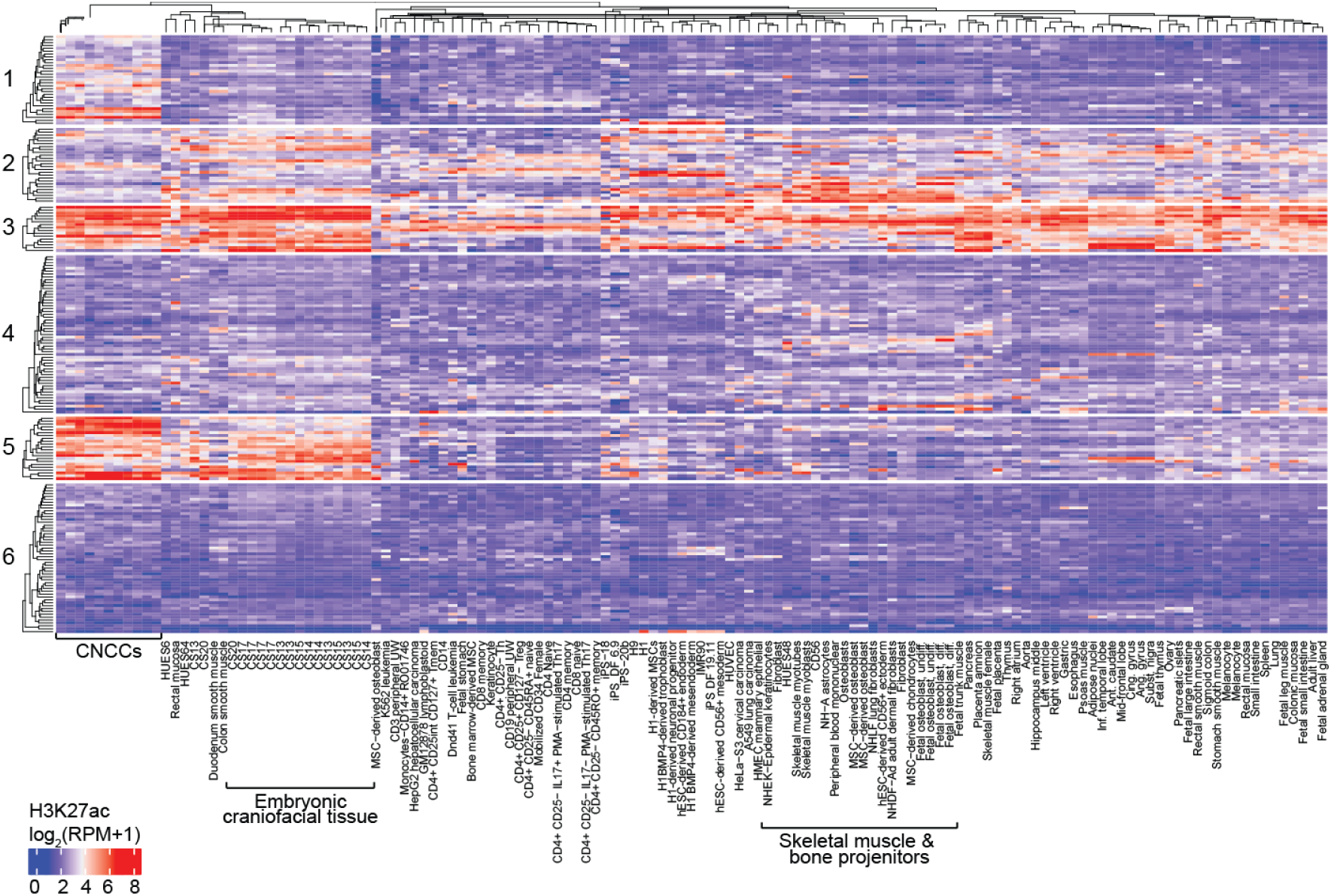
Activity of 203 peak SNPs in all cell-types studied. H3K27ac signal calculation and k-means clustering of SNPs were performed as described in Methods. Average linkage clustering on Euclidean distances was performed both within each of the 6 row clusters and for all columns. SNPs in cluster are active primarily in CNCCs, representing activity in an early time point in development. Cluster 2, 3, and 5 represent SNPs likely active across development, displaying activity in both CNCCs and craniofacial tissue, with SNPs in clusters 2 and 3 additionally active across many cell types and tissues, suggesting broad roles in general development.

## Known and novel loci and SNP effects on multiple facial phenotypes

We identified 86 GWAS peaks that overlap with the results of prior association studies of normal-range facial phenotypes. In several such instances, we observed new details providing a more nuanced understanding of the underlying genetic architecture. For example, variants at the *TBX15/WARS2* locus (1p12) were previously reported to be associated with forehead prominence^9^ and self-reported chin dimples^7^, already indicating that this locus has multiple spatially separated effects on the face. In our current analysis, we see the same influence on forehead morphology (lead SNP rs3936018, located in the promoter region of *WARS2*), as previously reported (Fig. 4)^9^. Interestingly, this lead SNP overlaps in location with rs12027501, but each SNP is associated with a separate linkage disequilibrium cluster and rs12027501 was most significant in segment 48, representing part of the cheeks (Fig. 4). We also see a signal approximately 275kb upstream of *TBX15* (rs7513680) that was most significantly associated with morphology in segment 51, representing the bottom cheek area at the corners of the mouth (Fig. 4). Lastly, another GWAS peak is present approximately 301 kb downstream of *WARS2* (rs17023457) with an effect in the same area, though the two are approximately 725 kb apart (Fig. 4). These results, in which we are able to finely parse out the effect of a SNP even within a complex genomic region, highlight the utility of using hierarchical clustering to segment the face. Of interest, we observed twenty-four such loci with multiple peaks that each affect different facial phenotypes, suggesting that these variants might overlap with or be impacted by regulatory elements that affect the face in highly specific ways (Table S4, Fig. S10).

**Fig. 4.**
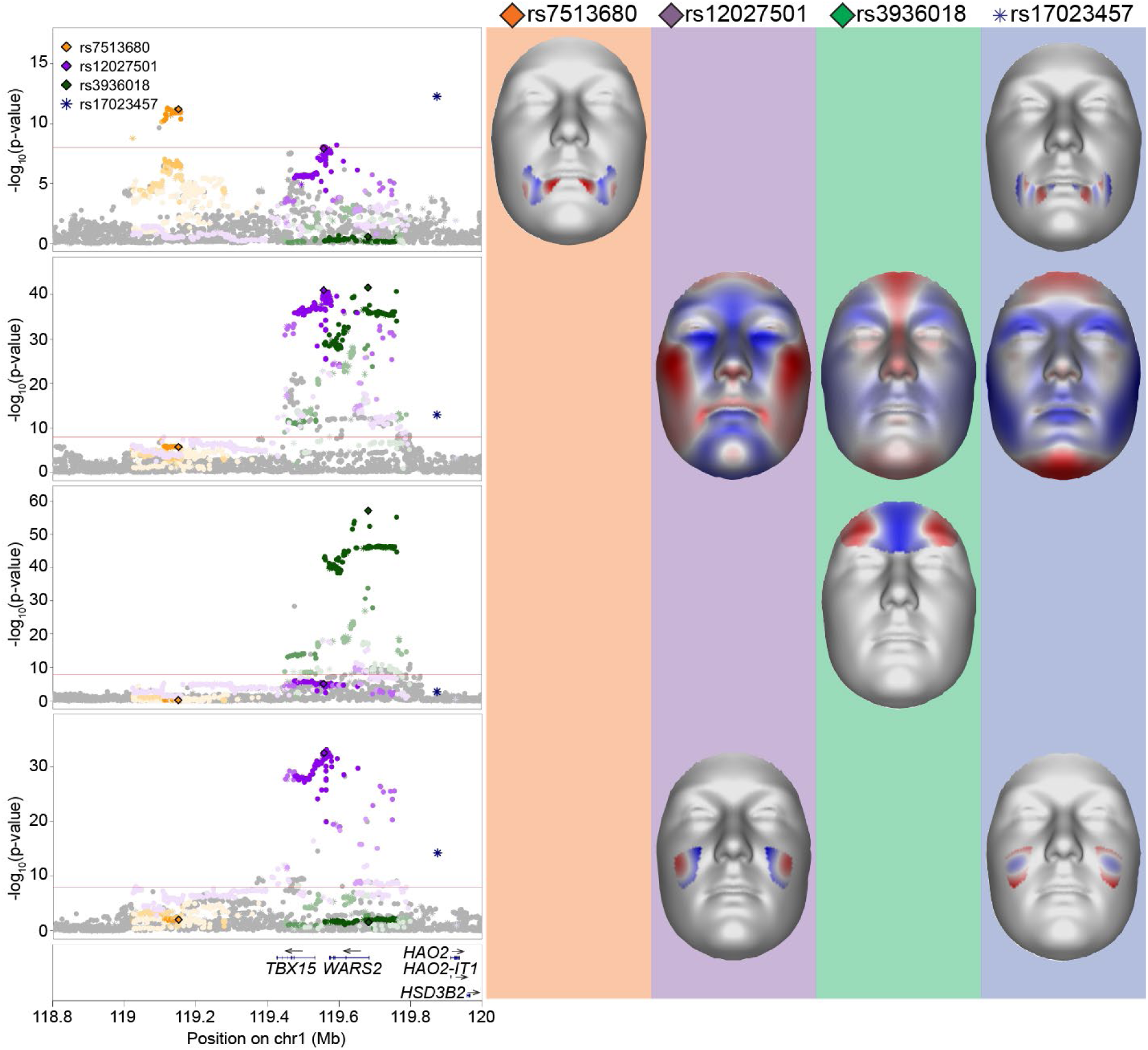
*TBX15-WARS2* multi-peak locus. LocusZoom plots and facial effects for four association signals near the *TBX15-WARS2* locus. Clustering based on linkage disequilibrium was performed to separate independent signals, resulting in the identification of four lead SNPs. Color for each SNPs is based on cluster association, with saturation indicating linkage disequilibrium with the lead SNP. SNPs represented by diamonds are the lead SNPs also present in the 1000G Phase 3 dataset; SNPs represented by circles are adjacent SNPs also present in the 1000G Phase 3 dataset; SNPs represented by asterisks are those not present in the 1000G Phase 3 dataset. For the segment in which each lead SNP had its lowest effect, we plot the facial effects for the lead SNPs reaching significance in that segment as the normal displacement (displacement in the direction normal to the facial surface) in each quasi-landmark going from minor to major allele, with red colored areas shifting outward while blue colored areas shift inwards.

A total of 64 GWAS peaks observed in our analysis are located at loci harboring putative craniofacial genes (implicated from human malformations or animal models), but which had not yet been observed in GWAS for normal-range facial morphology. For instance, *MSX1* has been implicated in orofacial clefting in humans^24,25^ and mice^25,26^, and is also widely expressed in lip and dental tissues during development^27^. We observed two distinct peaks at the *MSX1* locus (4p16.2), one approximately 55 kb upstream of *MSX1* with a pronounced effect on the lateral upper lip (lead SNP rs13117653) and a second peak, about 323 kb upstream of *MSX1* and located in the intron of *STX18*, involving the lateral lower lip and mandible (lead SNP rs3910659; Fig. S11). This result could indicate a potential role of *STX18* in craniofacial development, or provide further evidence that complex phenotypic effects seen in our human sample could be due to the action of multiple regulatory elements within a single locus. In support of this, Attanasio *et al*., demonstrated that the activity of *Msx1* in the second pharyngeal arch and maxillary process of the e11.5 mouse embryo is recapitulated by the combined activity of two separate enhancers, located 1 and 235 kb upstream of the gene promoter^23^.

Our GWAS additionally revealed 53 peaks at loci harboring genes with no previously known role in craniofacial development or disease, though many of the implicated genes at these loci are known to have a general role in developmental processes critical to proper morphogenesis. For example, *DACT1* is an established antagonist of the WNT signaling pathway, which is known to be involved in craniofacial development^28^, though *DACT1* has not previously been associated with facial morphology. *DACT1* is mostly studied for its involvement in gastric cancer, however it has also been shown to inhibit the delamination of neural crest cells, further supporting its involvement in facial development^29^. In the current study, variants at the *DACT1* locus are associated with mandibular morphology, particularly in the chin region (Fig. S11). *DACT1*, and the other 52 signals not previously associated with craniofacial morphology, are promising new candidates of potential roles in facial morphogenesis.

Given the proximity of our GWAS peaks to enhancers, and the understanding that some genomic regions are known to influence multiple phenotypes, we hypothesized that some lead SNPs could have additional associations with facial phenotypic traits besides that with which they were originally associated during the CCA. To test this, we performed pairwise linear regressions between each of the facial phenotypic traits identified during the CCA step and the genotypes at all other lead SNPs (see cross-peak association in Methods; Fig. S12), and identified 13 SNPs with associations with multiple phenotypic traits across multiple quadrants (Table S5). For example, rs1370926, a transcription factor binding site located 34 kb upstream of *PAX3* (based on Ensembl GRCh37 annotation^30^), showed associations with phenotypic traits in segments relating to morphology of the nose (Quadrant II; segments 5, 10, 11, 20, 22, 23, 40, 44, 45, and 47), upper lip (quadrant I; segments 9, 36), forehead (Quadrant IV; segment 28), and eyes (Quadrant IV; segment 60). This indicates an association between *PAX3* and many facial regions, which is consistent with previous GWAS findings implicating *PAX3* in general facial morphology, independent of phenotyping method or sample^7,31–34^. These 13 SNPs might show associations in multiple segments because of proximity to enhancer sequences, expansive effects across multiple cell populations, or early actions in facial development, perhaps in multiple facial prominences, that are then carried along to facial regions with little phenotypic correlation in our sample (i.e. different facial segments). *PAX3*, for example, plays a key role early in development by influencing the induction of the neural crest and the proliferation of craniofacial neural crest cells^35^. Seven of these SNPs can be grouped based on H3K27ac signal into clusters which had the highest activity in CNCCs and craniofacial tissues (clusters 1, 2, and 3; Fig. 3). Higher activity in craniofacial tissue is consistent with effects across multiple parts of the face, since higher activity within a tissue with as many different cell types as the embryonic face likely means that these SNPs are active in many cell types relevant to facial variation.

## Genetic interactions impacting facial variation

Given the complexity of the human face and its component traits, it is likely that the genetic architecture contributing to facial variation includes not only the genomic regions impacting multiple parts of the face, as described above, but also groups of genomic regions that contribute to the same facial trait, perhaps through actions in similar cell types or explicit interactions among variants. To better analyze and rank the effects of multiple genotypes on a facial trait, we used structural equation modeling (SEM) to refine our understanding of which groups of variants best explain the variance observed in each facial segment. SEM is a multivariate statistical analysis technique that has the ability to analyze structural relationships between measured variables (e.g. genetic variants and covariates) and latent constructs (univariate phenotypes derived from the PCs of the analyzed facial segment). This was done in an iterative manner, eventually resulting in 50 SEM models (corresponding to 50 facial segments) that converged successfully and were considered well-fitting based on recommended model fit parameters (Table S6). For each of these 50 models, the output included a univariate latent variable and a list of variants ranked by their estimated contribution to that variable (between 1 and 60, depending on the model), highlighting the polygenic nature of facial variation captured by the latent variable. Importantly, SNPs that significantly explained variance in the same segment showed higher correlations of cross-sample H3K27ac activity than when compared to non-significant SNPs for that segment, indicating that the SEM-refined lists of SNPs for each segment are likely those that are similar in either their spatial or temporal cellular activity (Fig. S13). We further analyzed these refined SNP lists for explicit genotype interactions by assessing, for each of the 50 models, whether interactions between the genotypes increased or decreased the median distribution of the latent variable. Four SNP combinations showed significant pairwise epistatic interactions (Table 1; Fig. 5; Fig. S14). For example, rs76244841 (*PRDM16* associated) and rs62443772 (*GLI3* associated) were found to have a significant interaction in facial segment 9, which covers the premaxillary soft tissue from the base of the columella to the oral commissure. *PRDM16* encodes a zinc finger transcription factor^36,37^ and has been shown to affect palatal shelf elevation through repression of TGFβ signaling^38,39^. *GLI3* encodes a transcriptional activator and a repressor of the sonic hedgehog (Shh) pathway, which has been shown to play a role in limb development^40–42^. In addition, there is evidence that mouse null *Gli3* mutants have a broad nose phenotype^43^ and genome-wide scans have previously implicated *GLI3* in affecting nose morphology^33^. The connection between *PRDM16* and *GLI3* can be seen by their interaction through the *SUFU* intermediary. Multiple studies conducted on *Drosophila melanogaster* have identified evidence for a tetrameric Hedgehog signaling complex comprising Fu, Ci (an ortholog of PRDM16), Cos2, and Su(fu) (an ortholog of SUFU), including evidence that Su(fu) binds directly to Ci^44–46^. SUFU has also been shown to mediate the phosphorylation of GLI3 via GSK3^47^ and has also been shown to interact with the GLI1-3 zinc-finger DNA-binding proteins^48,49^. Overall, these results indicate that the statistical evidence of SNP groups influencing polygenic facial variation, identified through SEM, and explicit variant interactions, suggested by the epistasis analysis, are potentially representative of true biological relationships, but must be confirmed with further study.

**Table 1.**
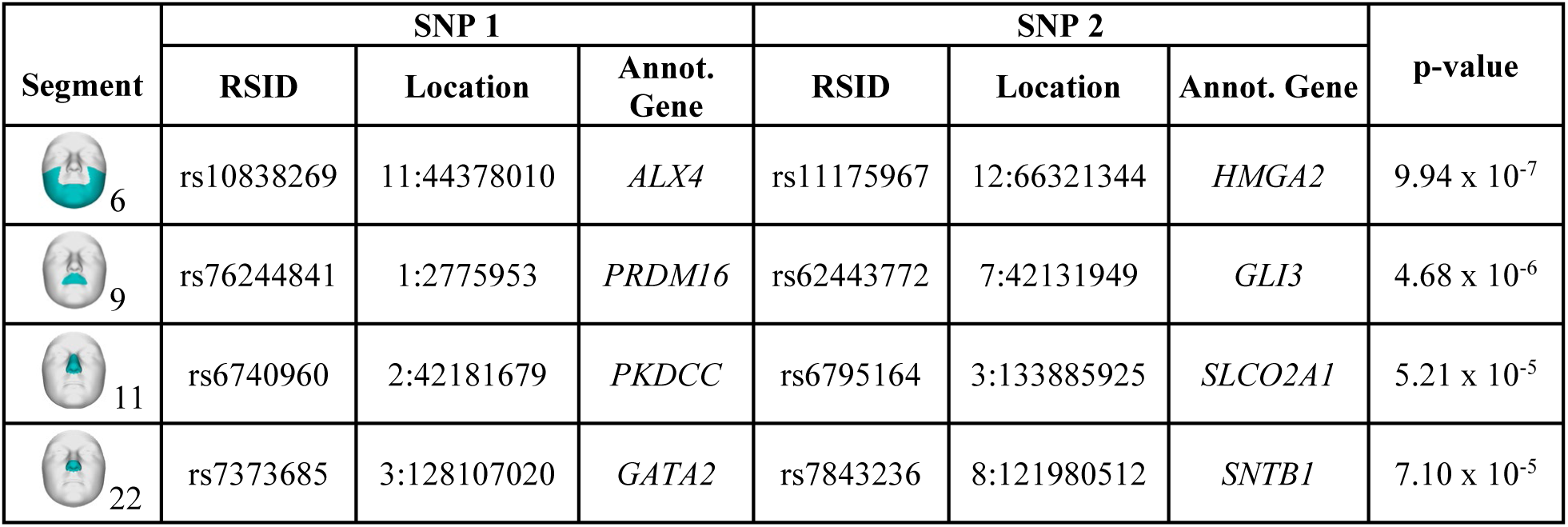
Four SNPs with evidence of epistatic interactions. For each of the 50 segments with a refined SEM model, we used the latent variables and SNP lists to test for evidence of epistasis. For the four SNP pairs with significant evidence of epistatic interactions, this table lists the epistasis p-value, rsID, GRCH37 location, and gene annotation. The phenotypic and marginal distributions for the pairs are depicted as boxplots in Fig. 5 and Fig. S14.

| Segment | SNP 1 |  |  | SNP 2 |  |  | p-value |
| --- | --- | --- | --- | --- | --- | --- | --- |
|  | RSID | Location | Annot. Gene | RSID | Location | Annot. Gene |  |
| 6 | rs10838269 | 11:44378010 | *ALX4* | rs11175967 | 12:66321344 | *HMGA2* | $9.94 \times 10^{-7}$ |
| 9 | rs76244841 | 1:2775953 | *PRDM16* | rs62443772 | 7:42131949 | *GLI3* | $4.68 \times 10^{-6}$ |
| 11 | rs6740960 | 2:42181679 | *PKDCC* | rs6795164 | 3:133885925 | *SLCO2A1* | $5.21 \times 10^{-5}$ |
| 22 | rs7373685 | 3:128107020 | *GATA2* | rs7843236 | 8:121980512 | *SNTB1* | $7.10 \times 10^{-5}$ |

**Fig. 5.**
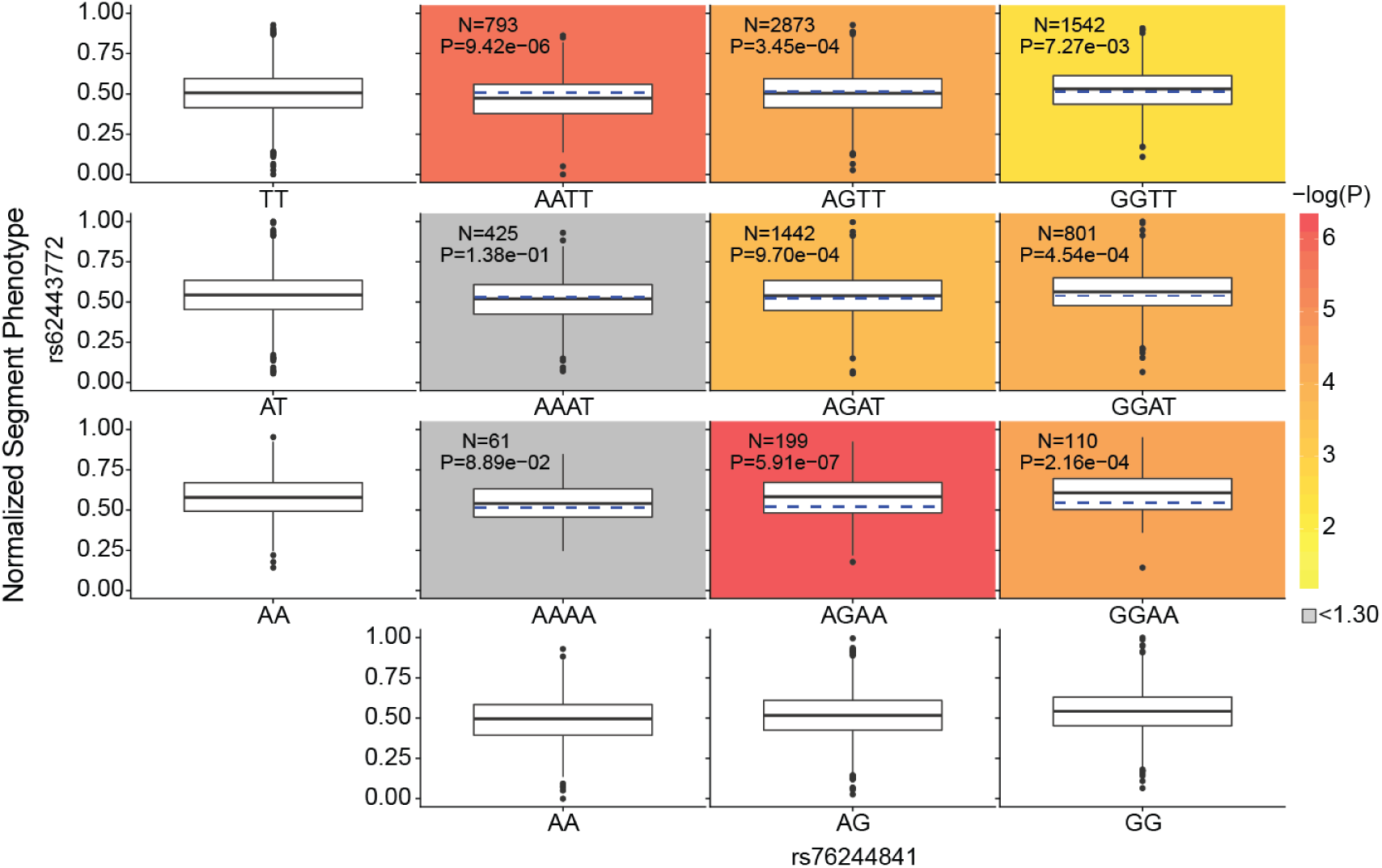
Phenotypic and marginal distributions for rs62443772 - rs76244841 epistatic pair. Plotted in the first column and last row are the marginal phenotypic distributions of the genotypes, which shows the phenotypic distribution that would occur if the two genotypes were acting independently. The median phenotype based on the two genotype distributions (blue dashed lines on the colored plots) was also calculated for each diplotype. The observed diplotype median (black line on the colored plots) was compared to the expected diplotype median (blue dashed lines on the colored blots) via Mood’s Median test^95^ with one degree of freedom. The resulting log transformed p-value was used to color the boxplots to illustrate significance, unless the difference was non-significant (i.e *p* > 0.05; log(*p*) < 1.30), in which the color was automatically set to grey. Within each colored boxplot is the untransformed Mood’s median p-value as well as the number of individuals used for significance testing. The phenotypic distribution facial image in the lower left corner was constructed by comparing the principal components.

## Conclusions

In sum, these results illustrate an avenue for investigating the coordinated processes underlying complex morphological structures, like the human face, at a deeper level than single associations between genotype and univariate phenotype. Having identified the largest number of signals associated with facial morphology to date, including a mix of known and novel loci, we additionally sought to better describe the processes and interactions through which the implicated genomic regions might influence the face. The genomic regions identified in this study are likely active across the timeline of facial development, evidenced by preferential H3K27ac activity in cells representing early timepoints (CNCCs) and progressively later timepoints (craniofacial tissues), and likely influence facial morphology by modulating the activity of nearby enhancers. Our results additionally illustrate that we are now able to pick up on the early shared developmental processes linking multiple complex structures, here both face and limb development. This is supported by genes near our lead SNPs being enriched for patterns and processes relevant to both anatomical regions and by the clustering of some lead SNPs through preferential H3K27ac activity in cell types relevant to both the face and skeleton. Lastly, though we know complex phenotypes like the face are polygenic, for the first time we now explicitly describe refined groups of SNPs influencing the same aspects of facial variation in each facial segment. These groups show correlated H3K27ac activity patterns and four pairs of SNPs exhibit evidence of epistatic interactions influencing facial variation. These results illustrate the potential of bioinformatic techniques to highlight spatial and temporal groupings of SNPs and connections between SNPs, representing a major step forward in our ability to characterize the polygenic genetic architecture of complex morphological structures like the face. We anticipate our results will be useful for researchers seeking to use model organisms and cell expression approaches in future functional investigations to refine the locations, timings, and connections that we suggest in this report, in an effort to better understand the biological forces that shape human and animal morphology.

## Online Methods

### Sample and recruitment details

The samples used for analysis included a combination of three independently collected datasets from the United States (US) and one dataset from the United Kingdom (UK), for a total sample size of *n* = 8,246. The US samples originated from the 3D Facial Norms cohort (3DFN) and studies at the Pennsylvania State University (PSU) and Indiana University-Purdue University Indianapolis (IUPUI). The UK dataset included samples from the Avon Longitudinal Study of Parents and their Children (ALSPAC). Institutional review board (IRB) approval was obtained at each recruitment site, and all participants gave their written informed consent prior to participation. For children, written consent was obtained from a parent or legal guardian. Some individuals from the 3DFN and PSU samples were previously tested for associations with facial morphology in our prior work^9^. A breakdown of the samples used for analysis is shown in Table S2. In all datasets, participants with missing information in sex, age, height, weight, or with insufficient image quality were removed.

For the 3DFN sample, 3D images and genotype data were obtained from the 3D Facial Norms repository^50^. The repository includes 3D facial surface images and self-reported demographic descriptors as well as basic anthropometric measurements from individuals recruited at four US sites: Pittsburgh, PA (PITT IRB PRO09060553 and RB0405013); Seattle, WA (Seattle Children’s IRB 12107); Houston, TX (UT Health Committee for the Protection of Human Subjects HSC-DB-09-0508); and Iowa City, IA (University of Iowa Human Subjects Office IRB (200912764 and 200710721). Recruitment was limited to individuals aged 3 to 40 years old and of self-reported European ancestry. Individuals were excluded if they reported a personal or family history of any birth defect or syndrome affecting the head or face, a personal history of any significant facial trauma or facial surgery, or any medical condition that might alter the structure of the face. The intersection of unrelated participants with quality-controlled images, covariates, and genotype data from individuals of European descent resulted in 1,906 individuals for analysis.

The PSU sample included 3D images and genotypes of participants recruited through several studies at the Pennsylvania State University and sampled at the following locations: Urbana-Champaign, IL (PSU IRB 13103); New York, NY (PSU IRB 45727); Cincinnati, OH (UC IRB 2015-3073); Twinsburg, OH (PSU IRB 2503); State College, PA (PSU IRB 44929 and 4320); Austin, TX (PSU IRB 44929); and San Antonio, TX (PSU IRB 1278). Participants self-reported information on age, ethnicity, ancestry, and body characteristics, and data were gathered on height and weight. Individuals were excluded from the analysis if they were below 18 years of age and if they reported a personal history of significant trauma or facial surgery, or any medical condition that might alter the structure of the face. No restriction on ancestry or ethnicity was imposed during recruitment, but only individuals of European descent were used in this study. The intersection of unrelated European participants with quality-controlled images, covariates, and genotype data resulted in 1,990 individuals for analysis.

The IUPUI sample includes 3D images and genotypic data from individuals recruited in Indianapolis, IN and Twinsburg, OH (IUPUI IRB 1409306349). Participants self-reported information on age, height, weight, and ancestry at the time of the collection. Individuals who were below 18 years of age were included if they had a parent or legal guardian’s signature. Similar to the PSU sample cohort, no restrictions were placed on the recruitment of participants, but only individuals of European descent and those meeting all quality control criteria were used in this study (*n* = 784).

The UK sample was derived from the ALSPAC dataset, a longitudinal birth cohort in which pregnant women residing in Avon with an expected delivery date between 1 April 1991 and 31 December 1992 were recruited^51,52^. At the time, 14,541 pregnant women were recruited and DNA samples were collected for 11,343 children. Genome-wide data was available for 8,952 subjects of the B2261 study, titled “Exploring distinctive facial features and their association with known candidate variants.” In addition to this, 4,731 3D images were available along with information on sex, age, weight, height, ancestry, and other body characteristics. The ALSPAC study website contains details of all the data that is available through a fully searchable data dictionary (http://www.bris.ac.uk/alspac/researchers/our-data/). The intersection of participants of European ancestry with quality-controlled images, covariates, and genotype data included 3,566 individuals. Ethical approval for the study was obtained from the ALSPAC Ethics and Law Committee and the Local Research Ethics Committees. Informed consent for the use of data collected via questionnaires and clinics was obtained from participants following the recommendations of the ALSPAC Ethics and Law Committee at the time. Consent for biological samples has been collected in accordance with the Human Tissue Act (2004).

### Genotyping platform

Genotyping of the 3DFN sample was performed at the Center for Inherited Disease Research at Johns Hopkins University. Participants, including 70 duplicate samples and 72 HapMap control samples, were genotyped on the Illumina OmniExpress + Exome v1.2 array, plus 4,322 investigator-chosen SNPs included to capture variation in specific regions of interest involved in the genetics of facial variation. PSU participants were genotyped by 23andMe on the v3 and v4 arrays (Mountain View, CA). Participants sampled at IUPUI were genotyped using Illumina’s Infinium Multi-Ethnic Global-8 v1 array consisting of 1.78M genome-wide markers. Genotyping was performed by the University of Chicago’s DNA Sequencing & Genotyping Facility (Chicago, IL). For the ALSPAC sample, participants were genotyped using the Illumina Human Hap550 quad genome-wide SNP genotyping platform by Sample Logistics and Genotyping Facilities at the Wellcome Trust Sanger Institute (Cambridge, UK) and the Laboratory Corporation of America (Burlington, NC), supported by 23andMe.

### Genetic imputation

Due to the several genotyping platforms used to genotype the US cohort, we chose to impute the samples from each platform separately, then combine the imputed results, following Verma, et al. 2014^53^. For each dataset, standard data cleaning and quality assurance practices were performed based on the GRCh37 (hg19) genome assembly. Specifically, samples were evaluated for concordance of genetic and reported sex (--check-sex), evidence of chromosomal aberrations, genotype call rate (--mind 0.1), and batch effects using PLINK 1.9^54^. SNPs were evaluated for call rate (--geno 0.1), Mendelian errors (--set-me-missing), deviation from Hardy-Weinberg genotype proportions (--hwe 0.01), and sex differences in allele frequency and heterozygosity, also using PLINK 1.9. The genotypes were “harmonized” with 1000 Genomes Project (1000G) Phase 3^55^ using Genotype Harmonizer^56^ with a window size of 200 SNPs, a minimum of 10 variants, and alignment based on minor allele frequency (--mafAlign 0.1). This program was also used to filter out ambiguous SNPs, update the SNP id, and update the reference allele as needed, all in reference to the 1000G Phase 3 genotypes. After genotype harmonization, the study datasets were merged (*n* = 44,383 SNPs in common) and explored using principal components analysis to assure that there were no batch effects by genotyping platform. Relatedness across the entire US sample was also assessed using this intersection and the KING software^57^. Relatives were noted in the per-platform subsets, and the imputation process proceeded for the full number of quality-controlled SNPs from each platform.

Prior to phasing, special quality control steps were performed on each platform. First, the allele frequency of each SNP was compared to the allele frequency of that SNP in the 1000G Phase 3 dataset. SNPs were removed if the allele frequency in the study dataset was not within |0.2| of any one of the 1000G super populations. We also removed SNPs with duplicate positions, any remaining insertions/deletions, copy number variants, and haploid genotypes. Individuals were removed if they had heterozygosity values ±3 standard deviations from the mean. Haplotypes were estimated using SHAPEIT2^58^. The samples were then imputed to the 1000G Phase 3 reference panel using the Sanger Imputation Server^59^ with the Positional Burrows-Wheeler Transform (PBWT) pipeline^60^, resulting in nearly 40 million variants for each dataset. SNP-level (INFO score >0.8) and genotype-per-participant-level (genotype probability >0.9) filters were used to omit poorly imputed variants. The datasets were then merged and filtered by SNP missingness (--geno 0.5), minor allele frequency (--maf 0.01), and Hardy-Weinberg equilibrium (*p* < 1 × 10^−6^) to produce a single merged dataset of all US participants with 7,417,619 SNPs for analysis.

The raw genotype data from ALSPAC was not available and restrictions are in place against merging the ALSPAC genotypes with any other genotypes. For this reason, imputed ALSPAC genotypes were obtained directly from the ALSPAC database and held separately during the analysis. Prior to phasing and imputation, these genotypes were subjected to standard quality control methods. Individuals were excluded on the basis of genetic sex and reported gender mismatches, minimal or excessive heterozygosity, disproportionate levels of individual missingness (>3%), and insufficient sample replication (IBD <0.8). Only individuals of European descent, compared to the Hapmap II dataset by way of multidimensional scaling analysis, were kept for imputation. SNPs were removed if they had a minor allele frequency of <1%, a call rate of <95%, or evidence for violations of Hardy-Weinberg equilibrium (*p* < 5×10^−7^). Haplotypes were estimated using SHAPEIT2^58^ and imputed to the 1000G Phase 1 reference panel (Version 3)^61^ using IMPUTE2^62^. After post-imputation quality control, the ALSPAC dataset contained 8,629,873 SNPs for analysis.

### Ancestry axes and selection of European participants

From the post-imputation merged dataset of the US participants, we identified the European participants by projecting them into a principal component (PC) space constructed using the 1000G Phase 3 dataset. To do this, we first excluded all indels, multi-allelic SNPs, and SNPs with MAF ≤ 0.1 in both the 1000G dataset and the US dataset and identified the SNPs common to both datasets. On this list (*n =* 1,940,221 SNPs), we iteratively performed linkage disequilibrium (LD) pruning (50 bp window, 5 bp step size, 0.2 correlation threshold) on the 1000G dataset until no variants were excluded. We then used this LD-pruned list (*n* = 461,372 SNPs) in a principal component analysis (PCA) to construct a population structure space based upon the 1000G project and projected our dataset onto that PCA space to obtain the ancestry axes of our dataset.

Once in a combined PC space, we calculated the Euclidean distance between all US participants and the 1000G samples. Using a k-th nearest neighbor algorithm, we identified the five nearest 1000G sample neighbors for each US participant in our dataset. The most common 1000G population label (e.g. CEU, GIH, YRI) from these five nearest neighbors was then assigned to the US participant in our dataset. Participants with the 1000G European population labels of CEU, TSI, FIN, GBR, and IBS were then selected for analysis. Prior to association, the genotypes in the US and UK dataset were separately corrected for sex, age, age-squared, height, weight, facial size, the first four genomic ancestry axes, and the camera system, using PLSR (plsregress from Matlab 2017b).

### 3D image acquisition

For all datasets, 3D images were captured using either one of two digital facial stereophotogrammetry systems or one laser scanning system. All participants were asked to have closed mouths and to maintain a neutral facial expression during image capture^63^. For the 3DFN sample, facial surfaces were acquired using the 3dMDface (3dMD, Atlanta, GA) camera system. PSU sample images were obtained with either the 3dMDface or Vectra H1 (Canfield Scientific, Parsippany, NJ) systems. The IUPUI sample was also imaged using the Vectra H1 system. The ALSPAC sample was imaged using a Konica Minolta Vivid 900 laser scanner (Konica Minolta Sensing Europe, Milton Keynes, UK). For this system, two high-resolution facial scans were taken and then processed, merged, and registered using a macro algorithm in Rapidform® software (INUS Technology Inc., Seoul, South Korea).

### 3D image registration and quality control

3D surface images were imported in wavefront.obj format into Matlab 2017b to perform the spatially dense registration process using a series of in-house functions packaged together in the MeshMonk registration framework^10^. Briefly, each image is cleaned to remove hair, ears, clothing, and other imaging artifacts. Five positioning landmarks are roughly indicated to establish image orientation. MeshMonk is then used to map a symmetric (relative to the sagittal plane) anthropometric mask of 7,160 landmarks onto the images and their reflections, which were constructed by changing the sign of the *x* coordinate^64^. This process results in a homologous configuration of spatially dense quasi-landmarks, allowing the image data from different individuals and camera systems to be standardized^10,65^.

Although variation in asymmetric facial features is of interest, in this work we sought to only study variation in symmetric facial shape. Therefore, when discussing facial shape, we always refer to the symmetric quasi-landmark configuration. To obtain the symmetric configuration, the registered original and reflected images were aligned using Generalized Procrustes analysis (GPA) to eliminate differences in position, orientation, and centroid size of both quasi-landmark configurations^66^. The average of the original and reflected quasi-landmark configuration constitutes the symmetric component, while the difference between the two constitutes the asymmetric component.

Outlier images, likely caused by image mapping errors, were identified using two approaches. First, as described in prior work^9^, outlier faces were identified by calculating Z-scores from the Mahalanobis distance between the average face and each individual face. Faces with Z-scores higher than two were manually investigated. Second, a score was calculated that reflects the missing data present in the image due to holes, spikes, and other mesh artifacts, which can be caused by facial hair or errors during the preprocessing steps. Images with high scores, indicating large gaps in the mesh, were also manually investigated. During the manual check, the images were either classified as images of poor quality and removed or were preprocessed and mapped again.

### Segmentation of facial shape

To study global and local effects on facial variation, we performed a data-driven facial segmentation on the UK and US datasets combined, as described previously^9^. Before segmentation, images in the two datasets were separately adjusted for sex, age, age-squared, height, weight, facial size, the first four genomic ancestry axes, and the camera system, using a partial least-squares regression (PLSR, function plsregress from Matlab 2017b). After adjustment, facial segments were defined by grouping vertices that are strongly correlated using hierarchical spectral clustering. The strength of covariation between quasi-landmarks was defined using Escoufier’s RV coefficient^67,68^. The RV coefficient was then used to build a structural similarity matrix that defined the hierarchical construction of 63 facial segments, broken into five levels (Fig. S1A). The configurations of each segment were then independently subjected to a GPA, after which a PCA was performed in combination with parallel analysis to capture the major variance in the facial segments with fewer variables^11,69^.

### Genome-wide association meta-analyses

The meta-analysis framework used consists of three steps: identification, verification, and meta-analysis (Fig. S2). For all analyses, the genotypes were coded additively based on the presence of the major allele. In the identification step, within each of the 63 facial segments, each SNP was associated with phenotypic variation using canonical correlation analysis (CCA, canoncorr in Matlab 2017b). CCA is a multivariate analysis which extracts the linear combination of PCs that are maximally correlated with the SNP, which represent the direction of phenotypic effect in shape space (i.e. a phenotypic trait). The correlation is tested for significance based on Rao’s F-test approximation^70^ (right tail, one-sided test). In sum, the identification step identifies a phenotypic trait most correlated with each SNP and a p-value representing the strength of the correlation. In the verification step, the shape variables (PCs) of the non-identification dataset (i.e. the verification dataset) were projected onto the trait found in the identification stage, which returns a univariate variable (UniVar in Fig. S2). These univariate variables were then tested for genotype-phenotype associations in a standard linear regression with the SNP genotypes of the verification dataset as independent variables and the univariate trait projection score as the dependent variable (regstats in Matlab 2017b). This function employs a t-statistic and a one-sided p-value was obtained with the Student’s T cumulative distribution function^71^ (function tcdf in Matlab 2017b). Next, the identification p-value (from the CCA) and the verification p-value (from the univariate regression) were combined in a meta-analysis using Stouffer’s method^12,13^. This process was repeated, resulting in two meta-analysis p-values accompanied by two identified traits, per segment and per SNP: first using US in the identification stage and UK as verification (META_US_), then using UK in the identification stage and US as verification (META_UK_).

### Sharing of genome-wide signal between facial segments

To assess the extent to which genome-wide signals of association with facial variation were shared between a pair of facial segments, we computed Spearman correlation rho between the two vectors of approximately 6 million SNP association p-values (Fig. S3). This was done separately for the META_US_ and META_UK_ p-values, but the pairwise rho values were very similar between the two datasets, so we used the average rho from META_US_ and META_UK_. This 63 × 63 matrix of correlations was visualized on top of the facial segmentation hierarchy to assess correlation both within and between facial quadrants (Fig. S4) and used to perform average-linkage hierarchical clustering (Fig. S5). To exclude the possibility that the correlations observed were driven primarily by linkage disequilibrium (LD) with the 203 lead SNPs reported, we downloaded the locations of 1,725 approximately independent European LD blocks in the human genome^72^ and re-computed the between-segment correlations when excluding all SNPs in the same LD block as any of the 203 lead SNPs. We also computed the mean SNP association p-value for each LD block and computed between-segment comparisons when using the average p-value for each LD block.

### GWAS peak selection

The analysis strategy yielded 126 p-values and 126 traits for every SNP, representing the 63 segments by two permutations of identification and verification. Per SNP, the lowest p-value was selected, and we noted in which version of the meta-analysis (META_US_ or META_UK_; “Best Permutation”) and segment (“Best Segment”) this p-value occurred. The study-wide Bonferroni threshold (*p* ≤ 6.25 × 10^−10^) was calculated as 5 × 10^−8^ / (40 * 2), where 40 is the number of independent segments and two is the number of datasets used. The FigShare repository for this work provides information on all SNPs reaching suggestive significance (*p* = 5 × 10^−7^). For the initial peak selection, we chose to group SNPs below genome-wide threshold by genomic position and the SNP with the lowest p-value per genomic region was selected as the lead SNP. Within a 1 Mb window (± 500 kb) of the resulting lead SNPs, we further refined the selection by performing a regression of slopes on the traits defined in the identification stage (in Best Permutation and Best Segment) to determine if adjacent SNPs showed consistent effects with the lead SNP. Adjacent SNPs were considered suggestively similar and belonging to the same signal as the lead SNP if the slope of the adjacent SNP trait and the lead SNP trait had a regression p-value lower than 0.2. On the other hand, a p-value higher than 0.2 was considered to indicate a different phenotypic effect by the two SNPs and led to a definition of a new lead SNP within the 1 Mb window. Peak selection on both genomic position and phenotypic effect resulted in 218 lead SNPs. Of these 218 lead SNPs, 203 showed consistent phenotypic effects in the US and UK in the Best Segment (Table S3). The consistency of effect across datasets was also determined with a regression of the CCA traits found in the identification stage for each dataset, with results considered sufficiently similar if they were below a false discovery rate of *p* ≤ 3.66 × 10^−2^. The 203 lead SNPs were mapped to 138 cytogenetic bands (i.e. loci) using the Ensembl GRCh37 locations^30^. This method of peak selection is statistical in nature and is thus not perfect. For example, our inspection of the LocusZoom plots for the *TBX15-WARS2* locus led to the identification of two clusters of SNPs, based on linkage disequilibrium, sharing the same genomic positions and affecting different facial segments, but separating these two clusters was not possible in our initial peak selection and they were considered a single signal until manual investigation. To more comprehensively identify all of the SNPs within a locus contributing to facial morphology, and the specific facial segments affected, fine mapping and other detailed investigations are needed.

### Gene annotation

Genes 500 kb up- and downstream of the lead SNPs were identified using the Table Browser of the UCSC Genome Browser^73^. The most likely candidate gene per lead SNP was identified based on a three-step system. First, we investigated whether any gene in the window was previously associated with craniofacial development or morphology through normal-range facial association studies, genetic disorders with facial dysmorphology as a symptom, or animal models. If this was not the case, we checked whether the gene was a known contributor to facial development based on the paper of Hooper and colleagues, who used transcriptome data from critical periods of mouse face formation to assess gene activity across facial development^74^. If both methods did not deliver a suitable candidate gene, the most likely candidate gene was selected based on the FUMA gene prioritization algorithm^15^.

To investigate the potential roles of the identified lead SNPs, enrichment analyses using FUMA and GREAT^16^ were performed, using preset parameters (Fig. S7). We additionally investigated whether SNPs within ±250 kb of the lead SNPs were previously identified in a GWAS using the NCBI-EBI GWAS catalog^75^. All links with facial morphology from the literature, based on genomic region or gene, are provided in Table S3.

### Cell-type-specific enhancer enrichment

Raw read (fastq) files of H3K27ac ChIP-seq from Prescott et al.^21^ (GSE70751; CNCCs), Najafova et al.^76^ (GSE82295; fetal osteoblast cell line, undifferentiated and differentiated), and Baumgart et al.^77^ (GSE89179; mesenchymal stem cell-derived osteoblasts) were downloaded and aligned to the human genome (hg19) using bowtie2 with default parameters. Aligned read (tagAlign) files of H3K27ac ChIP-seq from Wilderman et al.^22^ (GSE97752; embryonic craniofacial tissue) and the Roadmap Epigenomics Project^78^ (https://egg2.wustl.edu/roadmap/data/byFileType/alignments/consolidated/; various fetal and adult tissues and cell-types) were downloaded. To compare H3K27ac signal in the vicinity of lead SNPs between cell-types in an unbiased manner, we divided the genome into 20 kb windows, and calculated H3K27ac reads per million from each aligned read (bam or tagAlign) file in each window using bedtools coverage. We then performed quantile normalization (using the normalize.quantiles function from the preprocessCore package) on the matrix of 154,613 windows x 133 ChIP-seq datasets. We then selected the windows containing each of the 203 lead SNPs, 203 random SNPs matched for minor allele frequency and distance to the lead SNPs using SNPsnap^79^, or 619 Crohn’s disease-associated SNPs from the NCBI-EBI GWAS catalog^75^. Regions in the vicinity of SNPs associated with Crohn’s disease showed the highest H3K27ac signal in various immune cell types, serving as a positive control for both our approach and dataset (Fig. S8). K-means clustering was performed on the lead SNP H3K27ac signal with k = 6, as we found that this value maximized the number of clusters without significantly impacting cluster quality, as measured by silhouette width (Fig. 3).

### Chromatin state association in CNCCs and embryonic craniofacial tissue

Lists of human CNCC regulatory elements were annotated on the basis of multiple chromatin marks by Prescott et al.^21^ and embryonic craniofacial chromHMM states were computed in combined data from each Carnegie stage by Wilderman et al.^22^. For each set of regulatory regions, all regions within 20 kb of either lead SNPs or the above-described 203 random SNPs were considered. Enrichment/depletion of each class of regulatory region for lead SNPs versus random SNPs was computed using Fisher’s exact test (Fig. 2B, C).

### Cross peak association

For each lead SNP, one or more traits (i.e. linear combination of PCs) were discovered during the identification step in both META_US_ and META_UK_ permutations, depending on the number of segments in which the SNP showed a significant effect. Taking into account the polygenic and complex nature of facial shape, it is possible that a single SNP can be associated with more than one trait. To test this possibility, we executed a cross peak association (CPA), in which we performed a linear regression between all the lead SNPs and the facial phenotypic traits of each of the other lead SNPs, acting as a pool of possible secondary traits for association.

For each lead SNP we took the union of segments in which the SNP had a significant result in META_US_ and META_UK_. We then averaged the traits from the META_US_ and META_UK_ identification steps of each SNP in each selected segment. Accordingly, *n* = 1,405 traits were used as input in the CPA. Next, we projected all of our participants in both datasets onto each of these *n =* 1,405 traits using the same technique as used in the verification step of the GWAS, obtaining univariate variables that encapsulate the location of a participant relative to the other participants along the trait. These univariate variables, together with the genotypes at the lead SNPs, then served as input for linear regressions (regstats in Matlab 2017b) in which we tested for associations between each of the univariate variables and the genotypes at all lead SNPs excluding the lead SNP with which the trait was originally identified. We defined a CPA result as significant when the p-value was lower than a Bonferroni threshold of *p* ≤ 1.7617 × 10^−7^ (overall tests performed = 283,810, resulting from 1,405 traits each associated with 202 SNPs; Figure S12). Across all facial segments, we identified 75 lead SNPs with significant associations with an additional trait besides the trait identified for that SNP in the CCA step. Of these, 36 associations were present with phenotypes in multiple segments. For example, rs1572037 was associated with three different facial traits in segments 23, 45, and 47, all of which are in quadrant II. Because of the hierarchical nature of the facial segmentation, it is possible that associations within the same quadrant are driven by the inheritance of similar phenotypes from an upper level of the hierarchy. Thus, we further refined our results to 13 SNPs with associations to multiple phenotypes across multiple quadrants (Table S5).

### Structural Equation Modeling

To better define the cause-effect relationships between the significant genotypes and their collective phenotypic effects, both the US and UK participants were used as input for structural equation modeling (SEM) using the Lavaan package in R^80^. Mathematically, SEM analyses are a combination of a measurement model, which is constructed via confirmatory factor analysis, and a structural model, which is constructed using path analysis. In general, Lavaan outputs a best fit model that summarizes all genotype, phenotype, and covariate interactions, as well as a latent variable (aka “mask”), which is produced by a built-in dimension reduction that condenses the multidimensional facial phenotype from many principal components down to a single univariate phenotype. Parameters, which represent the interactions between the input variables, are generated by comparing the real covariance matrix between input variables and the estimated matrix created by numerical maximization, in our case carried out via maximum likelihood estimation. To maximize statistical power, Schreiber et al. recommend having at least 10 participants per parameter^81^. For our analyses, separate SEM models were constructed for each segment using each of the 203 lead SNPs and the shape PCs for all 8,246 participants. Missing genotypes were substituted with the most common genotype based on frequency. Covariates of age, sex, height, weight and face size (i.e. centroid size) were also included as model input. Prior to analysis, the distributions of these covariates were plotted and transformed, if necessary, to display near normal distributions. As genotypes are trichotomous, normality was not assessed.

Since analyzing all variants and all principal components for each segment via a single SEM would require the modeling of thousands of interactions and require extensive computational resources, separate SEM models were initially constructed. First, for each segment, we separated the 203 variants into three groups and ran three SEM models on each of these groups, plus all covariates. If any of the three subset SEMs did not converge, we then re-grouped the SNPs into four or more groupings and re-ran the subset SEM models on these groupings. This process was repeated until all subset SEMs converged and we had parameter estimates for all 203 SNPs. Next, for each segment, SNPs with p-values lower than 0.2 in the initial subset SEMs were collected and a unified SEM model for each segment was created and subsequently refined. If the unified SEM model did not converge, then this segment was discarded and no further analysis was performed. If all of the SNPs included in the unified model had p-values lower than 0.2, a cutoff selected to maintain model stability, no further changes were made, and we reported the model fit indices and parameter estimates. For segments where the unified SEM model produced SNP p-values greater than 0.2, the SNPs included in the SEM model were pruned by selecting SNPs with *p* < 0.05 and the model was re-run with this reduced set of SNPs. This process was repeated until all SNPs had p-values lower than 0.2. In the case of segments 7, 16, and 25, this iterative pruning process led to a rapidly declining model, so we elevated the SNP pruning p-value from 0.05 to 0.1 to account for instability in these models. Once the model refinement was complete (i.e. all SNPs had *p* < 0.2), we designated the SNPs with *p* < 0.05 as significantly contributing to variance within the segment.

In general, the number of model parameters generated by the final refined SEM model for each segment ranged between 92 and 217, depending on the number of shape PCs and SNPs included in each model. As 8,246 participants were used, this led to a range of 38-90 participants per parameter, which is well above recommendations^81^. Additional statistical power was lent to our models by having a large number of samples and input variables per latent factor^82^. Of the 63 segments, the SEM models for 13 segments were discarded because they did not converge on a solution, which normally occurs when variants are non-informative for that particular segment or the variance of the segment is low. For each of the 50 SEM models where the refinement process was successful, we evaluated the fit of each model by instituting cutoffs on the following indices: Chi-square (p-value < 0.05), comparative fit index (CFI > 0.90), root mean square error of approximation (RMSEA < 0.08), and standardized root mean square residual (SRMR < 0.08)^83,84^, which generally indicate the strength of how well the SEM models the data. 18 models passed all recommended model fit parameters and 32 models passed all but one of the fit indices, leading to the conclusion that the refined SEM models fit our data well. Final model fit indices and model parameter estimates are provided in Table S6. Reassuringly, for segments that are closely related in the segmentation hierarchy (i.e. segments 5, 11, 23, and 47) there is an average overlap of 46% of the variants meeting the *p* < 0.05 cutoff for significance, compared to 13.6% average overlap for non-hierarchically related segments (i.e. segments 5 and 6). The H3K27ac activity across all cell types was compared for significant variants both within and between segments using Spearman’s rho (Fig. S13).

### Epistasis Analysis

We additionally used the univariate latent variable and the variants passing the *p* < 0.05 significance cutoff from the final 50 refined SEM models (*p* < 0.1 for segments 7, 16, and 25) to assess whether interactions between genotypes increase or decrease the distribution of the latent variable. For each segment, the effect on the latent variable of all diplotype combinations of variants were assessed via a chi-square analysis in Plink 1.9^54^. After correcting for multiple testing, four SNP pairs were significant at *p* < 0.05 (Table 1). For these four pairs, the nine diplotype combinations and their normalized phenotypic and marginal distributions were plotted (Fig. 5; Fig. S14) to assess the genotypic contribution to epistatic masking (i.e. the combination of two variants reduce the output phenotype) and boosting (i.e. the combination of two variants elevate the output phenotype). This was performed using the R packages Agricolae, Cowplot, ggplot2, ggpubr, gridExtra, gtable, grid, Hmisc, psych, and data.table^85–94^. For each diplotype combination, the marginal phenotypic medians of the singular genotypes were averaged in order to visualize the predicted phenotypic distribution that would occur if the two genotypes were acting independently (dashed blue lines). This average was compared to the medians of the combined diplotypes (solid black lines). Significance testing was performed using Mood’s Median test^95^ with one degree of freedom. Follow up data mining on the four epistatic SNP pairs was performed using VarElect^96^, StringDB^97^, and Encode^98,99^.

## Acknowledgments

We are extremely grateful to all the families who took part in this study, the midwives for their help in recruiting them, and the whole ALSPAC team, which includes interviewers, computer and laboratory technicians, clerical workers, research scientists, volunteers, managers, receptionists and nurses. We are also very grateful to all of the US participants for generously donating their time to our research, and to present and former lab members who worked tirelessly to make these analyses possible.

## Author contributions

Conceptualization (Ideas; formulation or evolution of overarching research goals and aims): PC, MDS, SMW, JRS, JW, SW

Data curation (Management activities to annotate (produce metadata), scrub data and maintain research data for initial use and later re-use): JDW, KI, JL, PC, MKL, RJE, SW

Formal analysis (Application of statistical, mathematical, computational, or other formal techniques to analyze or synthesize study data): JDW, KI, SN, RJE, JR, PC

Funding acquisition (Acquisition of the financial support for the project leading to this publication): SR, TS, MLM, JRS, JW, SW, SMW, MDS, PC

Investigation (Conducting a research and investigation process, specifically performing the experiments, or data/evidence collection): JDW, KI, SN, RJE, JR, MKL, JL, JM, PC

Resources (Provision of study materials, computing resources, or other analysis tools): JDW, SN, RJE, JM, SR, EEQ, HLN, TS, MLM, JW, SW, SMW, MDS

Supervision (Oversight and leadership responsibility for the research activity planning and execution, including mentorship external to the core team): PC, SMW, MDS, SW, JW, JRS, MLM, TS, HP, GH

Visualization (Preparation, creation and/or presentation of the published work, specifically visualization/data presentation): JDW, KI, SN, RJE, JR, MKL, PC

Writing (original draft): JDW, KI, SN, RJE, JR

Writing (review and editing): JDW, KI, SN, RJE, JR, MKL, EF, TS, MLM, JRS, JW, SW, SMW, MDS, PC

## Data and materials availability

All of the genotypic markers for the 3DFN dataset are available to the research community through the dbGaP controlled-access repository (http://www.ncbi.nlm.nih.gov/gap) at accession #phs000929.v1.p1. The raw source data for the phenotypes - the 3D facial surface models in .obj format - are available through the FaceBase Consortium (https://www.facebase.org) at accession #FB00000491.01. Access to these 3D facial surface models requires proper institutional ethics approval and approval from the FaceBase data access committee. Additional details can be requested from SMW.

The participants making up the PSU and IUPUI datasets were not collected with broad data sharing consent. Given the highly identifiable nature of both facial and genomic information and unresolved issues regarding risk to participants, we opted for a more conservative approach to participant recruitment. Broad data sharing of the raw data from these collections would thus be in legal and ethical violation of the informed consent obtained from the participants. This restriction is not because of any personal or commercial interests. Additional details can be requested from MDS and SW for the PSU and IUPUI datasets, respectively.

The ALSPAC (UK) data will be made available to bona fide researchers on application to the ALSPAC Executive Committee (http://www.bris.ac.uk/alspac/researchers/data-access). Ethical approval for the study was obtained from the ALSPAC Ethics and Law Committee and the Local Research Ethics Committees.

KU Leuven provides the MeshMonk spatially dense facial mapping software, free to use for academic purposes (https://github.com/TheWebMonks/meshmonk). Matlab implementations of the hierarchical spectral clustering to obtain facial segmentations are available from a previous publication^100^ (https://doi.org/10.6084/m9.figshare.7649024.v1). The statistical analyses in this work were based on functions of the statistical toolbox in Matlab 2017b as mentioned throughout the Methods.

All relevant data to run future replications and meta-analysis efforts are provided in Matlab format online (https://doi.org/10.6084/m9.figshare.c.4667261.v1). This includes the anthropometric mask used, facial segmentation cluster labels, PCA shape spaces for all 63 facial segments, CCA loadings and all association statistics for the lead SNPs.

