## Supplemental Information for "Insights into the genetic architecture of the human face"

**This PDF file includes:**

Supplementary Note
Figs. S1 to S9, S11 to S14
Captions for Figure S10
Captions for Tables S1, S3 to S6
Table S2

**Other Supplementary Materials for this manuscript include the following:**

Fig. S10 (PDF)
Tables S1, S3, S4, S5, and S6 (Excel)

**Supplementary Note**

Literature evidence of epistatic interactions

The strongest SNP x SNP epistatic interaction was found between rs10838269 and rs11175967 ($p = 9.94 \times 10^{-7}$) within segment 6, which covers the area of the face from the zygoma to the mandible. Rs11175967 is an intronic variant mapped to the *HMGA2* transcription factor (12q14.3), which has been associated with Silver-Russell syndrome, symptoms of which include a triangular face shape and broad foreheads[1,2]. Rs10838269, its epistatic partner, is an intergenic variant whose nearest protein coding gene is the transcription factor *ALX4* (11p11.2), which is expressed in the mesenchyme of developing bones and has been shown to play a vital role in craniofacial development[3]. Previous morphology studies identifying both *HMGA2* and *ALX4* show that the genes respectively contribute to ear morphology and stature in sheep[4]. In addition, genomic analyses on finches have shown that alterations of *ALX1*, the protein product of which is functionally redundant to Alx4 in mice[5], and *HMGA2* have been associated with beak shape[6] and size[7].

Rs6740960 (candidate gene *PKDCC*) and rs6795164 (candidate gene *SLCO2A1*) ($p = 5.21 \times 10^{-5}$), and rs7373685 (candidate gene *GATA2*) and rs7843236 (candidate gene *SNTB1*) ($p = 7.10 \times 10^{-5}$) were significant pairs in facial segments 11 and 22, respectively, which are hierarchical segments that include areas surrounding the base of the nose. Due to the overlapping nature of the segments, these variants were analyzed as a collective group. The nature of the relationship between these four variants is less clear, however some trends are evident. The first is that there appears to be a connection between GATA2 and SLCO2A1 through AKT1. AKT1 is one of 3 related serine/threonine-protein kinases first characterized in mouse models[8], which regulate multiple processes, such as metabolism, cell survival, growth, proliferation, and angiogenesis. It has also been indicated in Proteus Syndrome, whose symptoms include bone development abnormalities[9,10]. SLCO2A1 is a solute carrier involved in the release and transport of prostaglandin[11,12] and has also been shown to be involved in hypertrophic osteoarthropathy[13–15]. SLCO2A1 regulates AKT1 and the Akt pathway through prostaglandin[16,17]. Furthermore, the PI3K/Akt signal pathway has been shown to negatively regulate the transcriptional activator GATA2[18]. There were not any connections found between PKDCC and SNTB1, however, there was an interesting connection between SNTB1 and GATA2 via Dystrophin (DMD). DMD serves as a key component of the dystrophin-associated glycoprotein complex, which helps stabilize the sarcolemma[19]. SNTB1 is an adapter protein that has been suggested to link receptors to the dystrophin glycoprotein complex[20,21]. GATA2 has also been shown to be a transcriptional factor of DMD[22]. Finally, there is evidence in mouse models that supports a connection between the Akt signalling pathway and DMD[23], which serves as another underlying link between three of the four epistatic hits. While there were no evident links between PKDCC and the other epistatic hits, it may be worth noting that this tyrosine-protein kinase has been previously shown to be involved in bone growth[24–26].

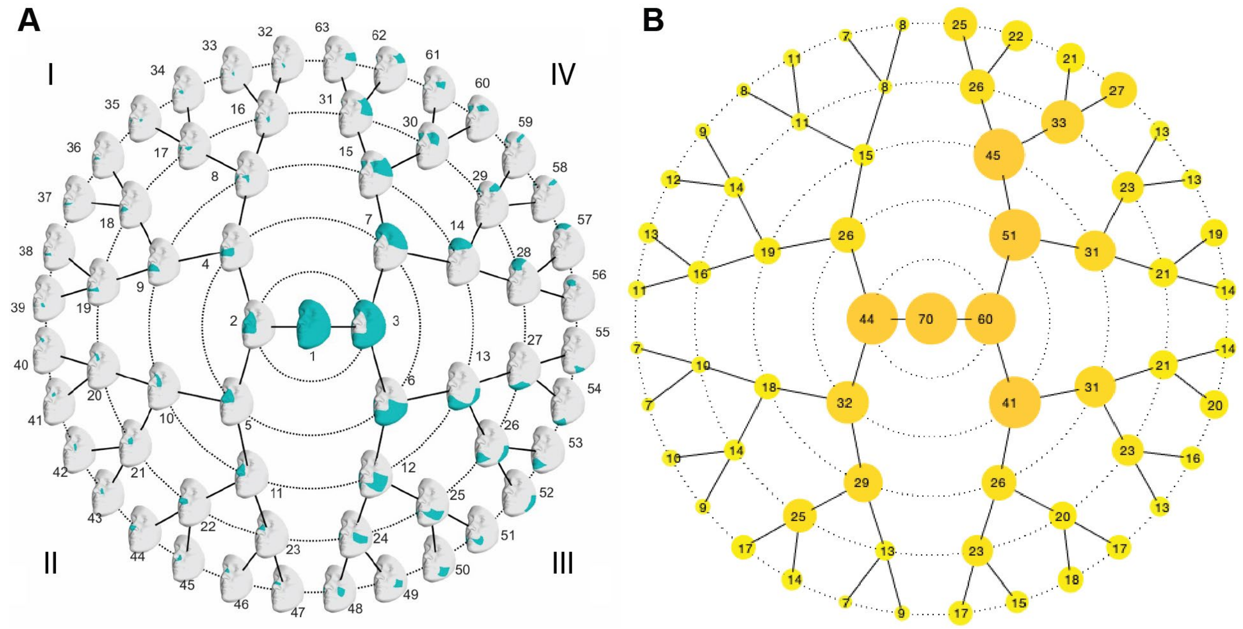


**Fig. S1. Hierarchical spectral clustering of facial shape.**
(**A**) Global to local facial segmentation of all 3D images included (*n* = 8,246) obtained using hierarchical spectral clustering. Segments are colored in teal. Roman numerals represent "quadrants" of facial segments. (**B**) The number of principal components retained after parallel analysis for each facial segment.

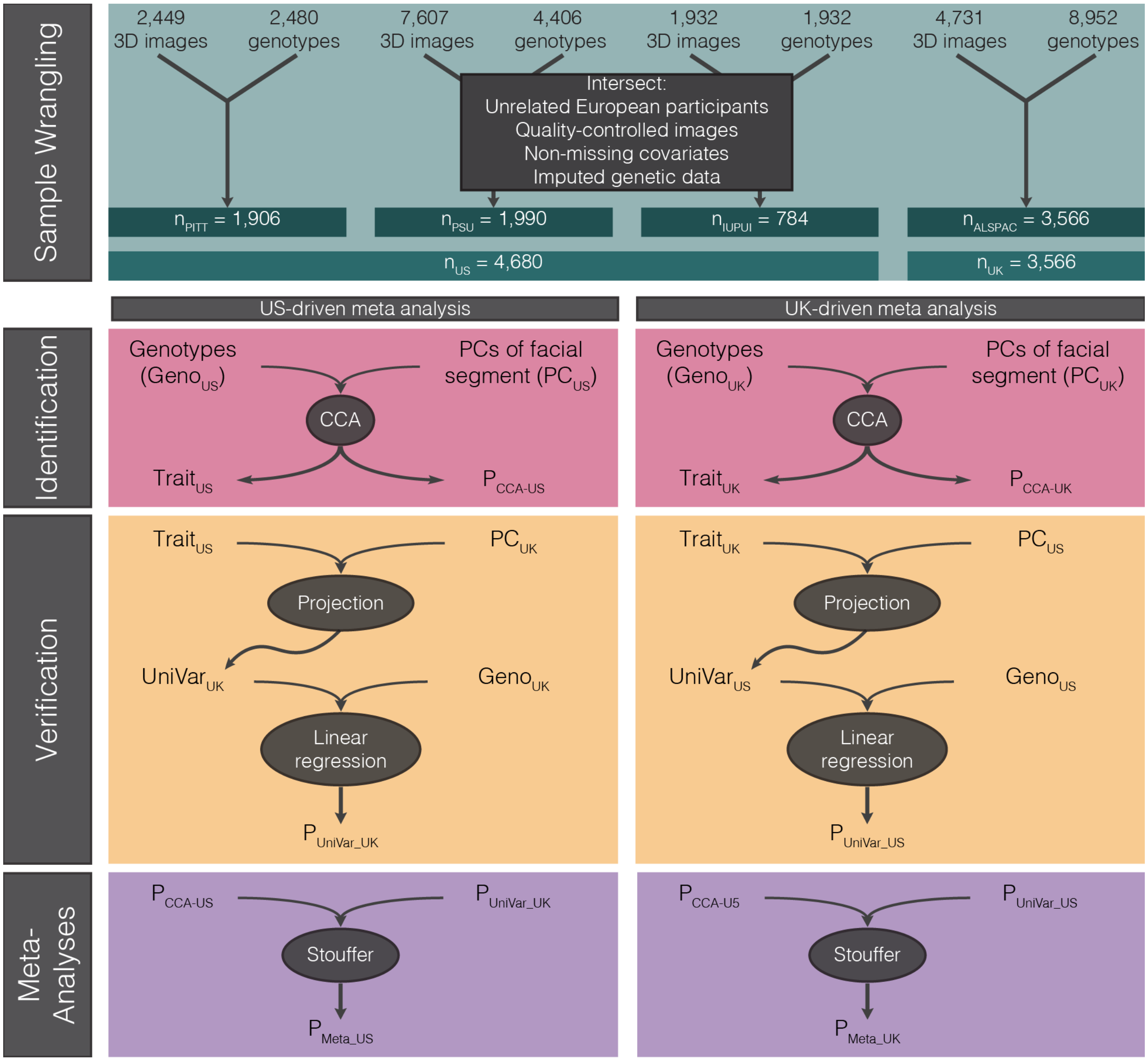


**Fig. S2. Study design.**

*Sample Wrangling*: Images and genotypes from each study were intersected and unrelated participants of European ancestry, with quality controlled images, covariates, and imputed genetic data were selected to obtain the analyzed data. *Identification*: Within each facial segment, canonical correlation analysis (CCA) was used to identify the facial principal components most correlated with the genotypes, which led to a p-value ($P_{CCA\text{-}US}$ or $P_{CCA\text{-}UK}$) and facial trait most correlated with each SNP ($Trait_{US}$ and $Trait_{UK}$). *Verification*: The principal components of the other dataset were then projected onto this trait to obtain a univariate variable representing the distribution of participants from the verification dataset for the trait identified in the identification dataset ($UniVar_{UK}$ and $UniVar_{US}$). The genotypes of the verification dataset are then tested against this variable via linear regression, resulting in an additional p-value ($P_{UniVar\text{-}UK}$ and $P_{UniVar\text{-}US}$). *Meta-Analysis*: The p-values from identification and verification are meta-analyzed using Stouffer's method, resulting in the final set of p-values from each meta-analysis permutation ($P_{META\text{-}US}$ and $P_{META\text{-}UK}$).

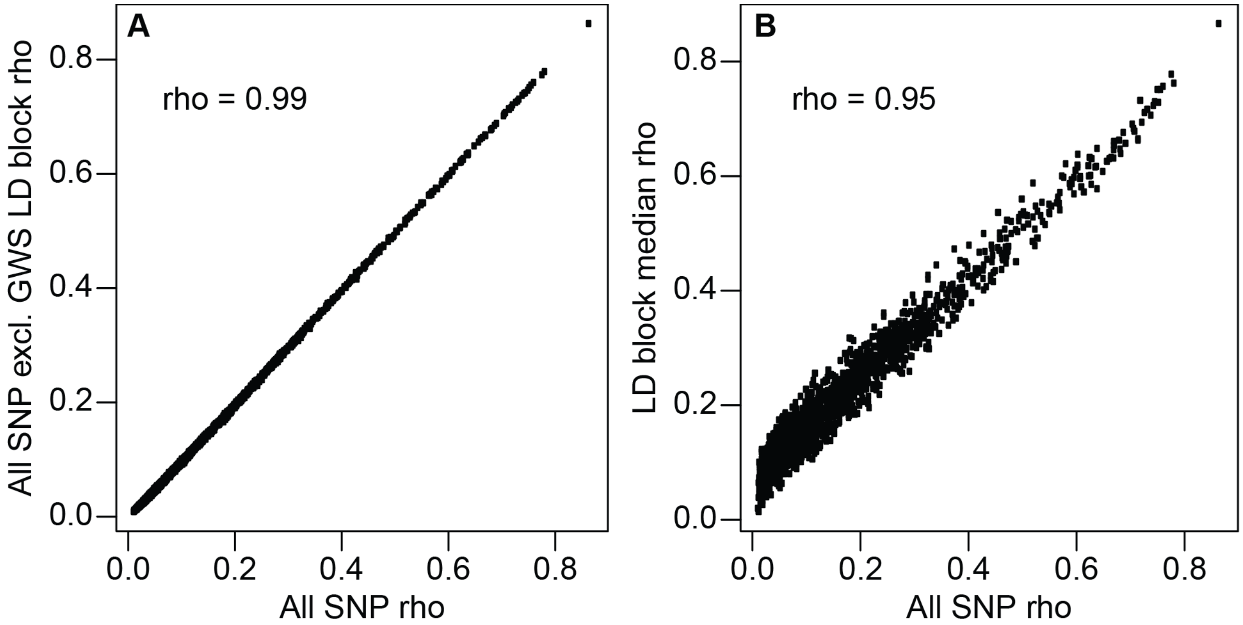


**Fig. S3. Correlations between facial segments are not driven by linkage to genome-wide significant variants or linkage disequilibrium in general.**
Correlation between rho values for segment-segment pairs calculated on the basis of p-values for all SNPs (x-axis) and either all SNPs excluding those in the same linkage disequilibrium block as the 203 genome-wide significant SNPs (y-axis, **A**), or SNP p-values averaged within each linkage disequilibrium block (y-axis, **B**). Linkage disequilibrium blocks in Europeans from Berisa and Pickrell (2016) were used[27].

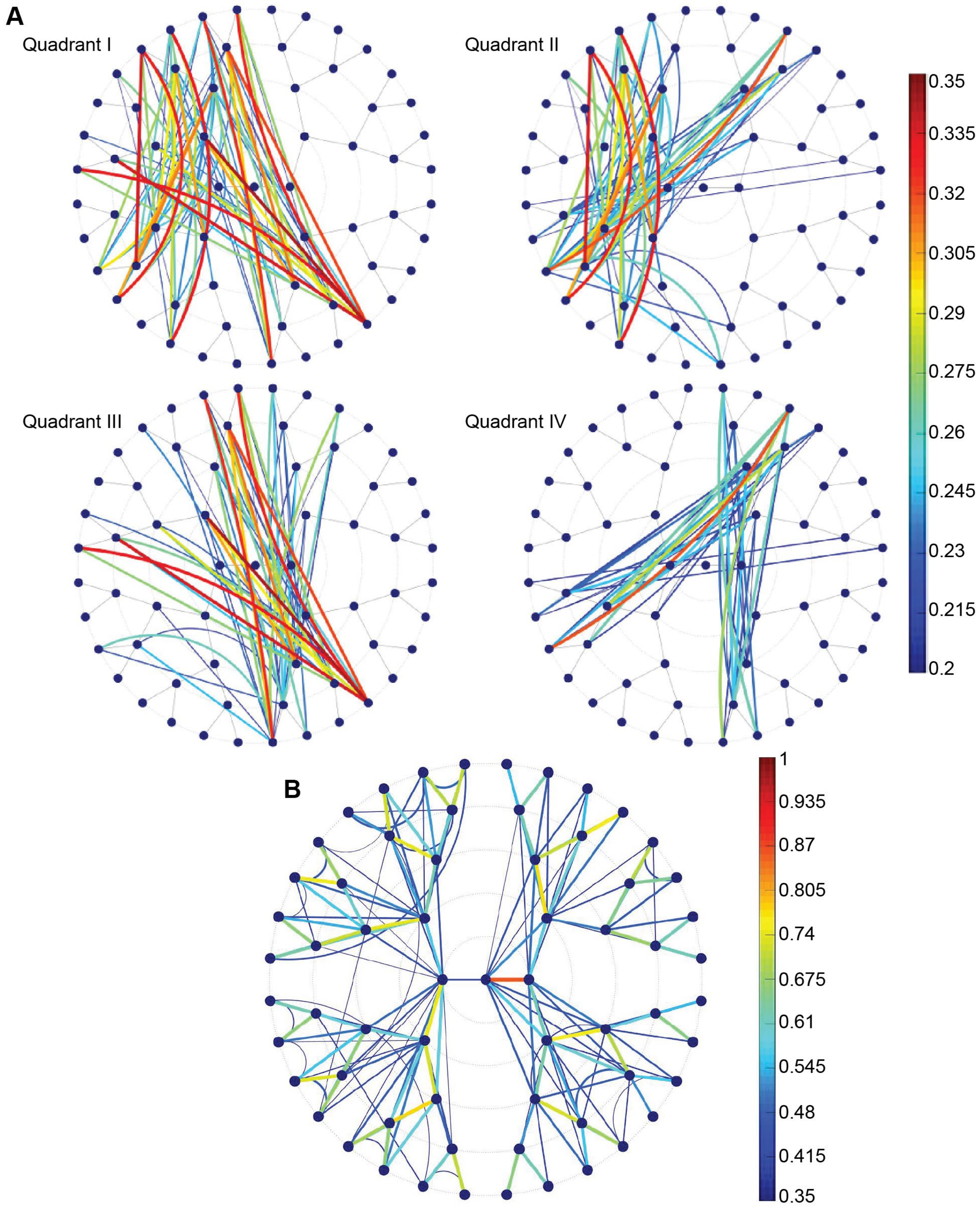


**Fig. S4. Genomic signal correlations.**
Spearman correlations, rho (calculated using genome-wide association p-values) between segments. (**A**) Correlations ranging from 0.2-0.35, which are all between segments from different quadrants. (**B**) Correlations ranging from 0.35-1.

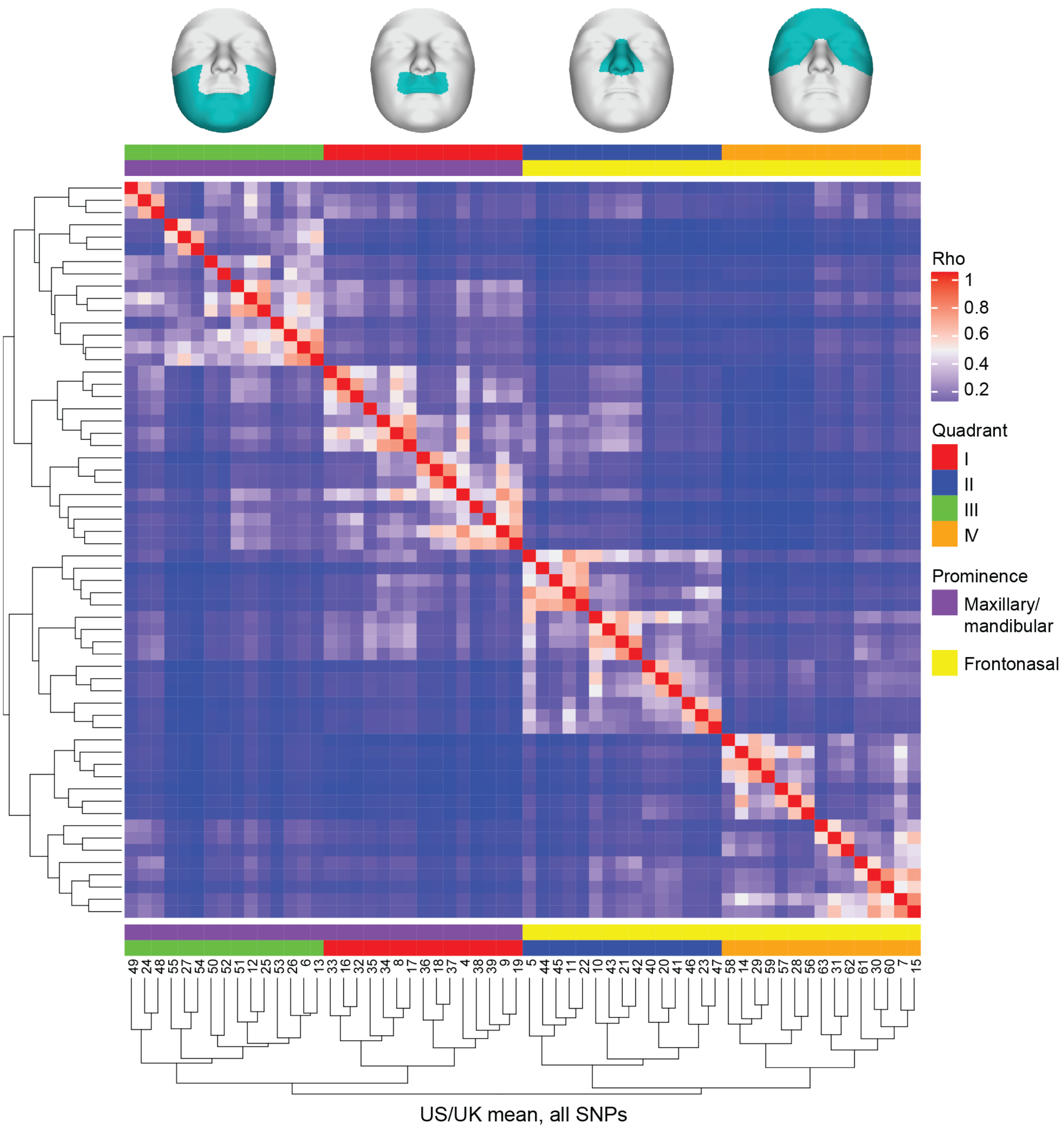


**Fig. S5. Clustering of facial segments on the basis of shared genetic signals.** Correlations between facial segments on the basis of SNP p-values were calculated as described in Methods, and average linkage clustering was performed using the matrix of correlation values. Quadrant colors in legend refer to the quadrant of the polar dendrogram in which the facial segment lies in, also represented by the facial images at the top, and embryonic facial prominences are assigned to each facial segment.

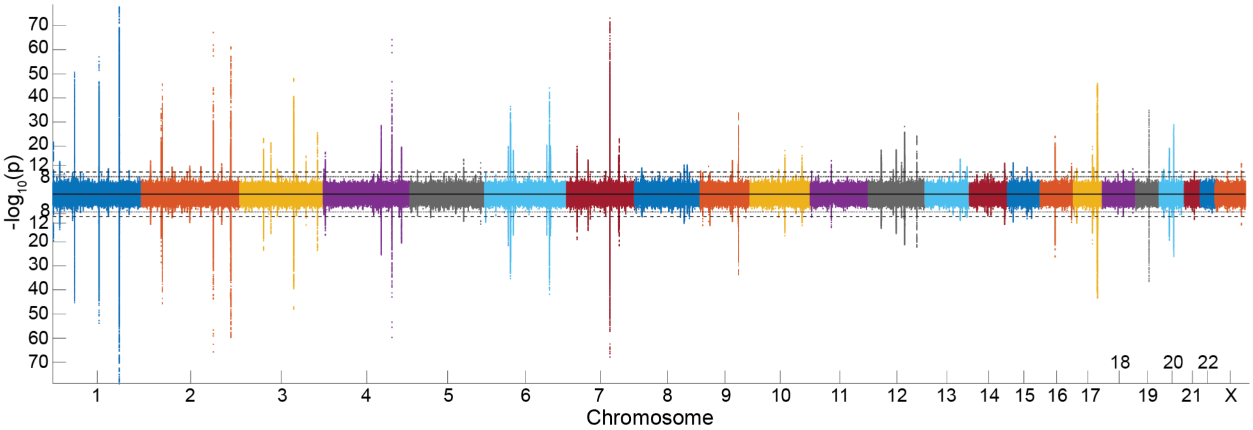


**Fig. S6. Miami plot of all results.**
Miami plot of all 63 facial segments combined, illustrating the chromosomal position of the associated loci from the meta-analysis permutation with the US dataset as identification (*top*) and UK as identification (*bottom*). The dotted horizontal line represents the genome-wide significance threshold ($p = 5 \times 10^{-8}$) and the dashed horizontal line represents the study-wide threshold ($p = 6.25 \times 10^{-10}$).

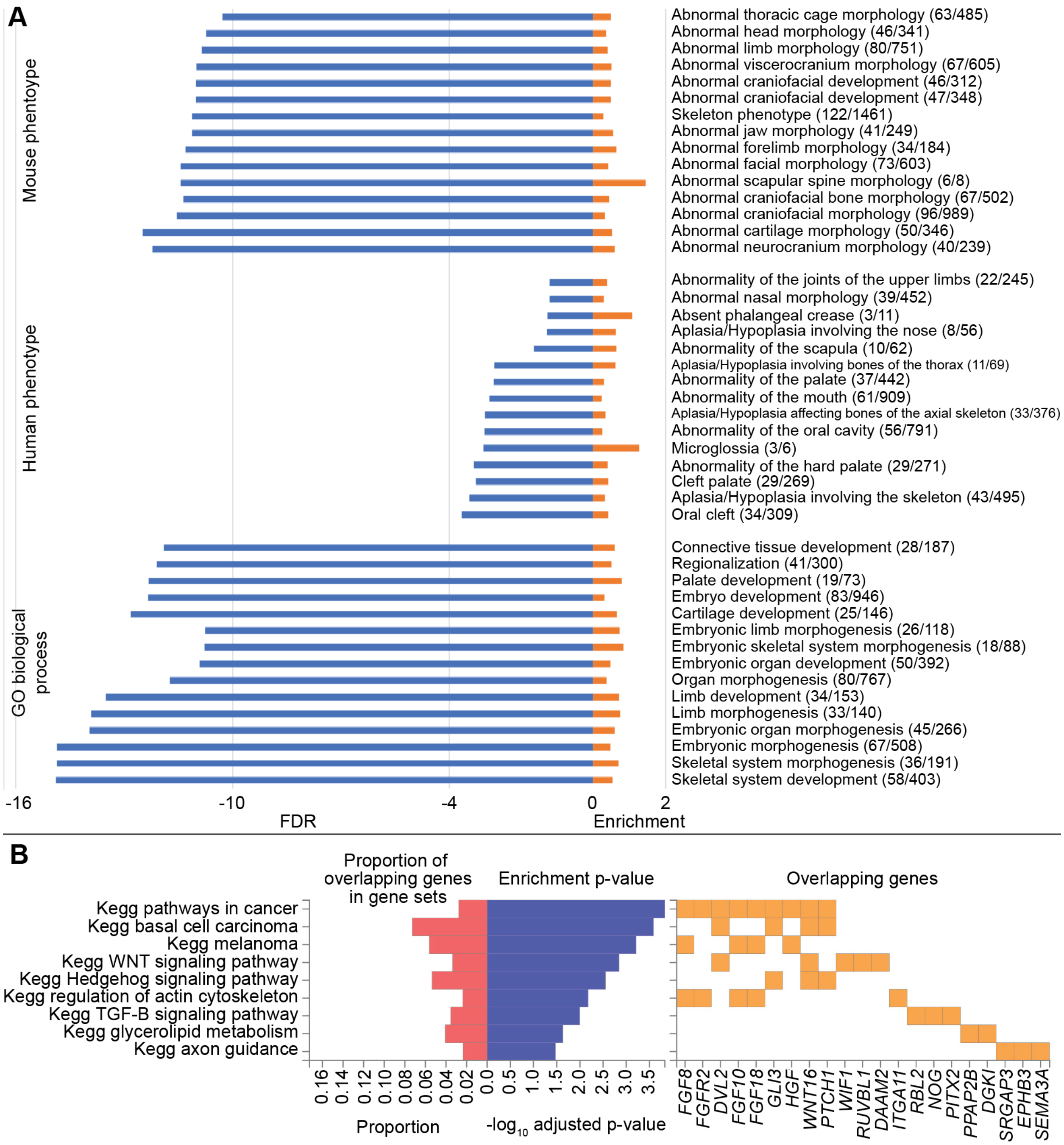


**Fig. S7. GREAT and FUMA analyses showing enrichment for craniofacial and limb development.**

(**A**) GREAT analysis, Plotted is the binomial test FDR (blue) and binomial enrichment (orange). We indicate the actual number of genomic regions in the test set with the annotation compared to the observed region hits (expected number of genomic regions in the test set with the annotation) behind every term. (**B**) FUMA analysis, indicating the KEGG pathways that were enriched in our results. Multiple pathways are relevant for craniofacial development. The right panel shows the genes that are involved in the pathways.

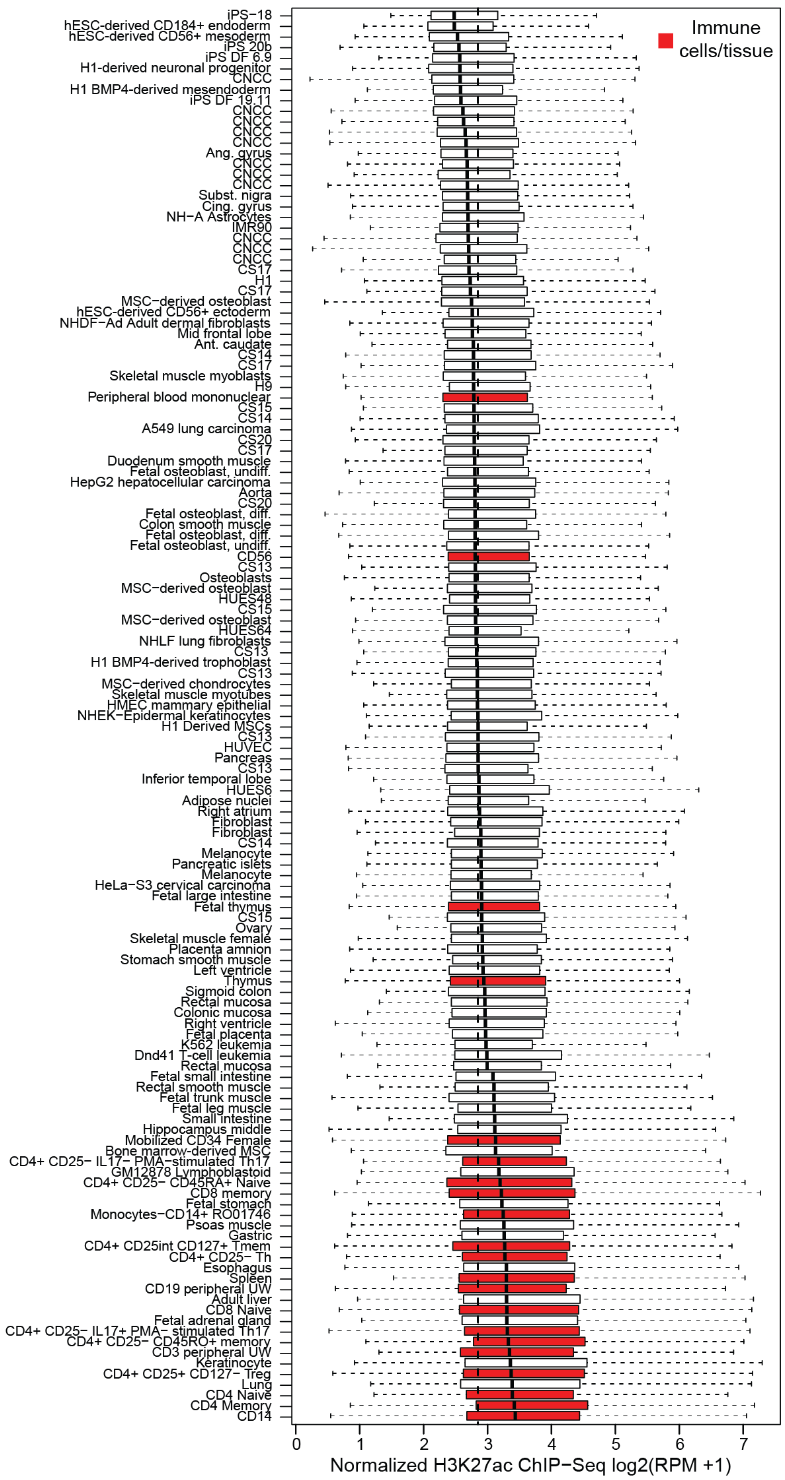


**Fig. S8. Regions nearby previously published SNPs associated with risk for Crohn's disease are preferentially active in immune cells and tissues.**

Each boxplot represents the distribution of H3K27ac signal in 20 kb regions around 619 Crohn's disease-associated SNPs from the NCBI-EBI GWAS catalog in one sample. See Methods for details on calculation of H3K27ac signal. Samples corresponding to immune cells and tissues are highlighted in red.

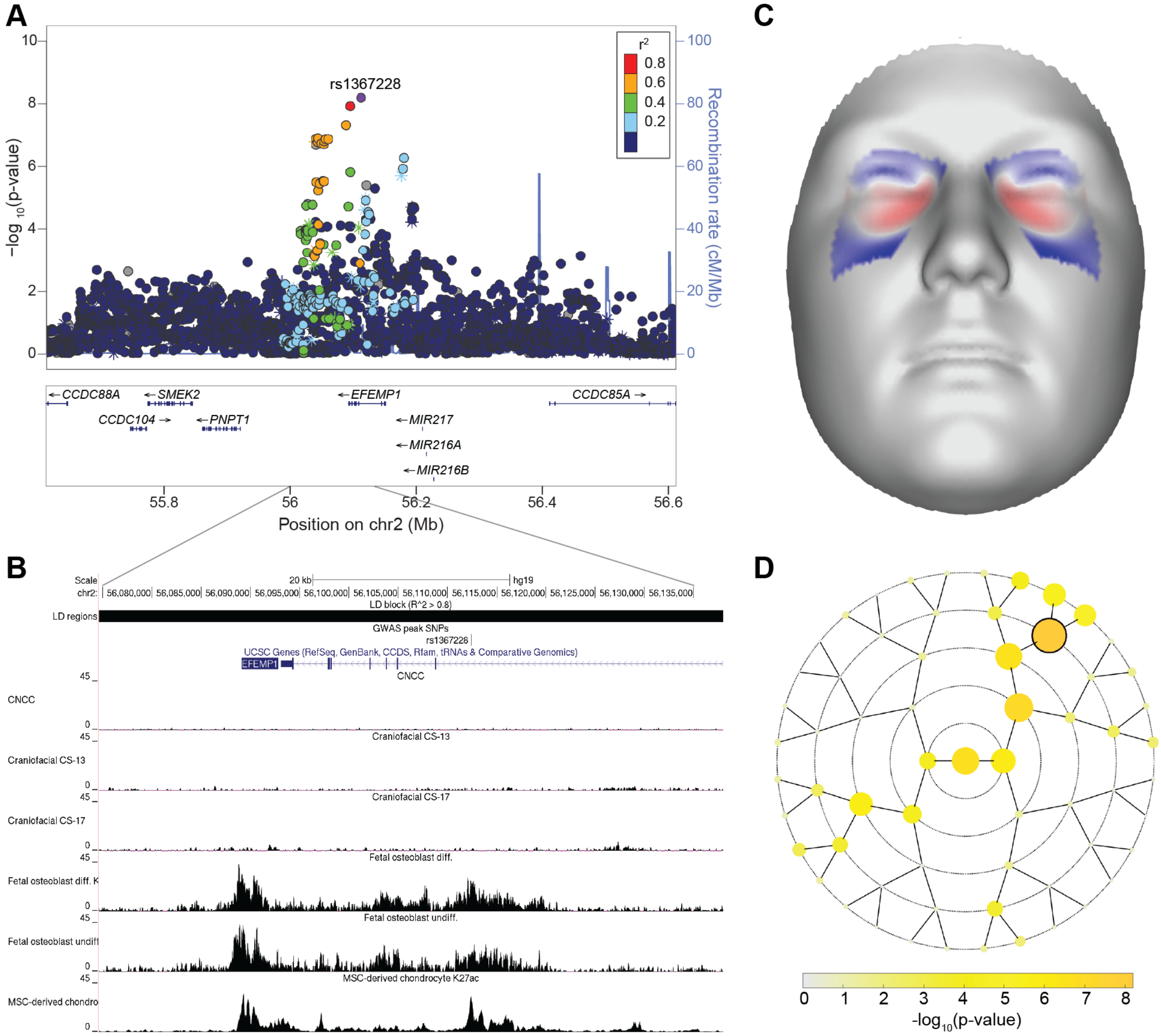


**Fig. S9. A locus containing the extracellular matrix protein *EFEMP1* and affecting shape around the eye sockets is specifically active in osteoblasts and chondrocytes.**
(**A**) Locus zoom plot of region surrounding rs1367228. Points are colored based on linkage disequilibrium with the labeled SNP. Asterisks indicate genotyped SNPs and circles indicate imputed SNPs. (**B**) UCSC browser plot for the region surrounding *EFEMP1*, H3K27ac signal tracks are displayed after input subtraction and reads per million normalization in 50 bp windows. (**C**) Effect of rs1367228 in segment 30 in the UK-derived meta-analysis, where the META$_{UK}$ p-value was lowest for this SNP. Plotted is the normal displacement (displacement in the direction normal to the facial surface) in each quasi-landmark going from minor to major allele, red colored areas shift outward while blue colored areas shift inwards. (**D**) Rosette plot showing the size and location of UK-derived meta-analysis p-values, with the genome wide significant result outlined in black. Segment locations correspond to Fig. S1A.

**Additional Figure S10 (Separate File). Overview of 24 multi-peak loci**
For each of the twenty-four multi-peak loci (listed in Table S4): (**A**) $-\log_{10}$(p-value) of the meta-analysis p-value per facial segment in $META_{US}$ and $META_{UK}$ permutations. Black-encircled facial segments have reached genome-wide significance ($p = 5x10^{-8}$). (**B**) The normal displacement (displacement in the direction locally normal to the facial surface) in each quasi landmark of the facial segment reaching the lowest p-value in $META_{US}$ and $META_{UK}$, going from the minor to the major allele. Blue indicates inward depression; red indicates outward protrusion. (**C**) LocusZoom plots in $META_{US}$ (top) and $META_{UK}$ (bottom), for the segment in which the SNP had its lowest p-value. Points are colored based on linkage disequilibrium ($r^2$) in the 1000 Genomes Phase 3 EUR population. Asterisks represent genotyped SNPs and circles represent imputed SNPs.

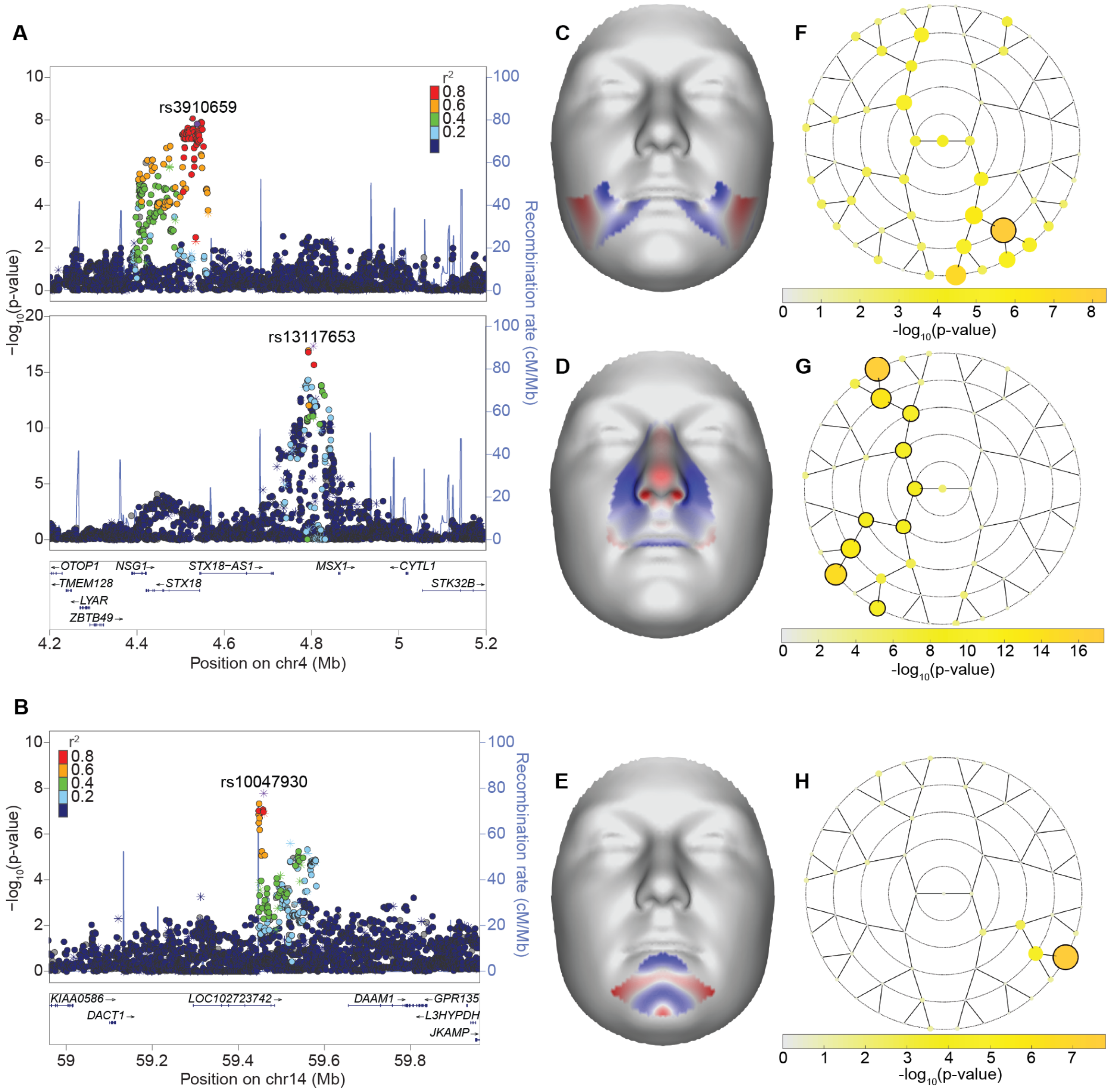


**Fig. S11.** ***MSX1*** **and** ***DACT1*** **loci.**

LocusZoom plots for the two association signals nearby *MSX1* (**A**), which has previously been implicated in orofacial clefting in humans and mice, and *DACT1* (**B**), which is a novel result. Points are colored based on linkage disequilibrium with the labeled SNP. Asterisks indicate genotyped SNPs and circles indicate imputed SNPs. Facial effects for the two association signals nearby *MSX1*: rs3910659 (**C**) and rs13117653 (**D**) and the signal nearby *DACT1*: rs10047930 (**E**). Effects are the normal displacement (displacement in the direction locally normal to the facial surface) in each quasi landmark of the lowest facial segment reaching genome-wide significance in $META_{UK}$, going from the minor to the major allele. Blue indicates inward depression; red indicates outward protrusion. Yellow rosette plots depict the $-log_{10}$(p-value) of the meta-analysis p-value per facial segment in $META_{UK}$ permutation. Black-encircled facial segments have reached genome-wide significance ($p = 5x10^{-8}$). ($-log_{10}$(p-value) of the meta-analysis p-value per facial segment in $META_{US}$ and $META_{UK}$ permutations. Black-encircled facial segments have reached genome-wide significance ($p = 5x10^{-8}$). (**F**) rs3910659; (**G**) rs13117653; (**H**) rs10047930.

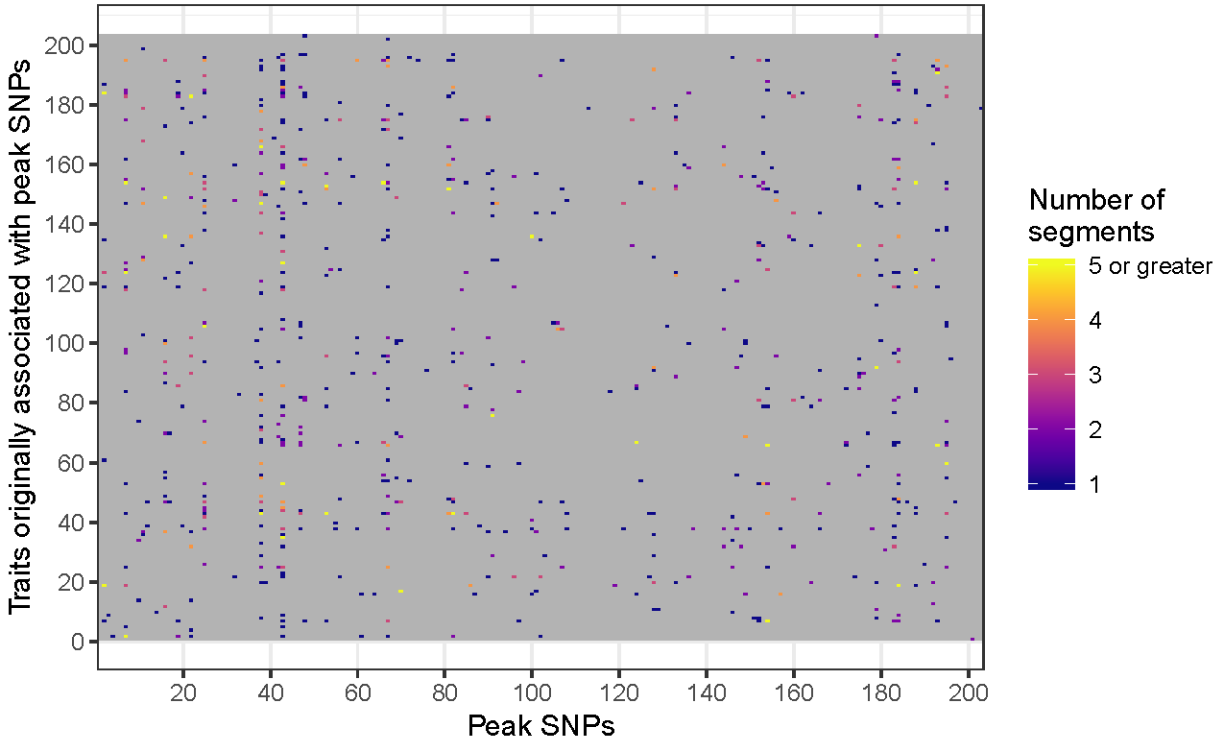


**Fig. S12. Significant cross peak association results and designation of multi-effect SNPs.** The number of segments in which the association between a trait and peak SNP was lower than the Bonferroni-corrected significance threshold ($p \leq 1.76 \times 10^{-7}$). Grey values are those in which the association did not reach significance in any segment. Viewing column-wise, we see certain SNPs are associated with many different traits. SNPs associated with traits in segments spanning multiple quadrants were considered "multi-effect" SNPs and are listed in Table S5.

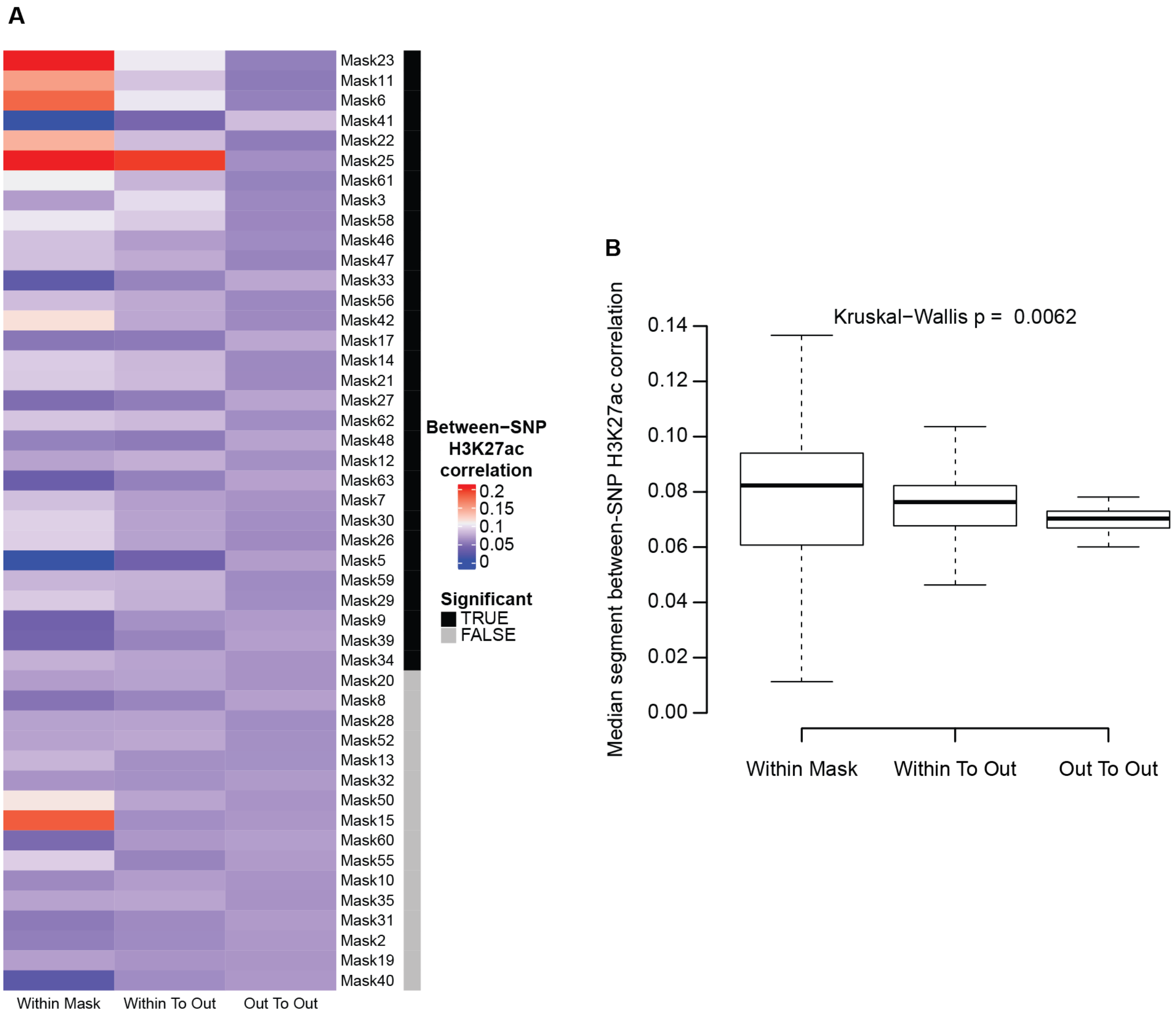


**Fig. S13. Correlation of H3K27ac activity among SEM models.**
(**A**) For all segments (aka "masks"), we compared the H3K27ac activity for significant SNPs from the refined SEM model for variation at that facial segment. Plotted is the Spearman's rho correlation between pairs of SNPs significant in the same SEM model ("Within Mask"); pairs of SNPs where one is from the SEM model and the other is not ("Within To Out"); and where both SNPs in the pair are from a different SEM model ("Out To Out"). Segments where the distribution of correlation across all cell types was significantly different ($p < 0.05$) are indicated in black. (**B**) For all cell types, the median correlation across all segments is plotted for each of the three SNP groupings. Significance between the means was determined using Kruskal-Wallis test.

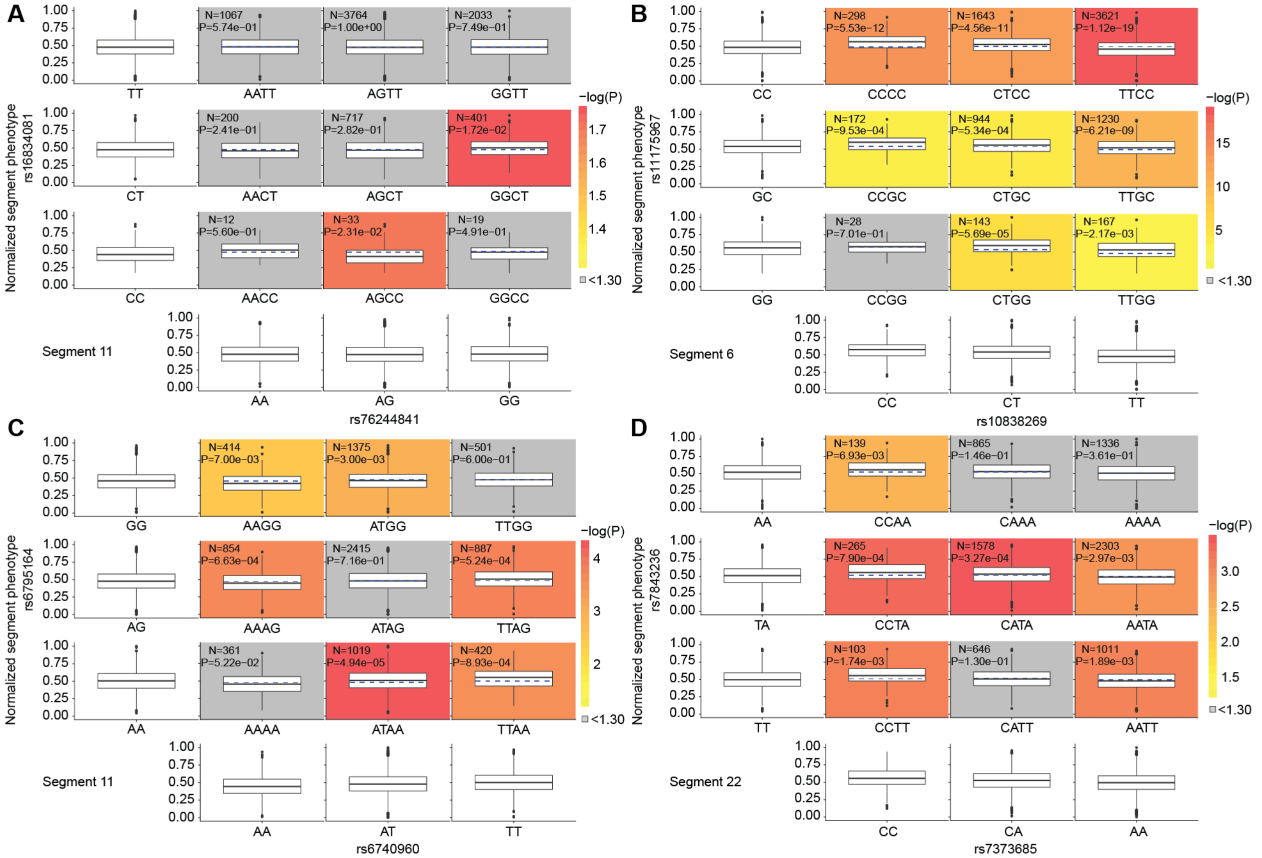


**Fig. S14. Phenotypic and marginal distributions for diplotype combinations.** For a random SNP pairing (**A**) and each significant epistasis pair (**B-D**), boxplots are plotted to visualize the epistatic effect on the phenotype. The marginal phenotypic medians of the singular genotypes (first column and last row of boxplots) were used to calculate and visualize the predicted diplotype phenotypic distribution that would occur if the two genotypes were acting independently (dashed blue lines in colored boxplots). This median was compared to the observed medians of the diplotypes (solid black lines; colored boxplots) via Mood's Median test[28] with one degree of freedom. Log transformed p-values were used to color boxplots if there was a significant ($p < 0.05$; $\log(p) > 1.30$) difference between the expected phenotype of the combined genotype and observed diplotype.

**Additional Table S1. (Separate file). SNPs associated with facial morphology through GWAS.** Expanded from Roosenboom et al., 2016[29] with permission. Where possible, we have listed the locus, SNP, gene annotation, and a description of the facial traits identified in the original publication. Bolded entries indicate that this locus, SNP, or gene was also identified or annotated in the current study. Studies with an asterisk (*) indicate an overlap in the study participants used in the referenced study and the current study. The 'Citations' tab contains the full citations for the referenced studies.

| | | **PITT** | **PSU** | **IUPUI** | **ALSPAC** |
|---|---|---|---|---|---|
| **Sex** | **Male** | 734 | 610 | 245 | 1682 |
| | **Female** | 1172 | 1380 | 539 | 1884 |
| **Age** | **Range** | 3 - 40 | 18 - 88 | 7.15 - 78.87 | 14 - 17 |
| | **Mean** | 22.45 | 33.39 | 23.67 | 15.33 |
| | **Std. Dev.** | 9.06 | 16.79 | 10.70 | 0.49 |
| **Height (cm)** | **Range** | 92.4 - 210.82 | 130.81 - 208.28 | 99.06 - 203.30 | 145 - 200.1 |
| | **Mean** | 163.43 | 168.75 | 169.24 | 169.38 |
| | **Std. Dev.** | 20.57 | 9.23 | 11.30 | 8.42 |
| **Weight (kg)** | **Range** | 12.7 - 193.5 | 35.2 - 162.7 | 22.68 - 181.44 | 32.3 - 124.6 |
| | **Mean** | 64.33 | 73.88 | 71.88 | 61.52 |
| | **Std. Dev.** | 22.38 | 17.05 | 18.65 | 11.75 |
| **SNPs passing QC (pre imputation)** | | 581,874 | 23andMe v3: 744,091<br><br>23andMe v4: 426,463 | 480,430 | 465,740 |

**Table S2. Sample breakdown for US and UK participants.**
For each dataset contributing to the US and UK sample, information is given for the sex, age, height, and weight distribution, as well as the number of SNPs used for imputation of unknown variants.

**Additional Table S3 (Separate File). 203 Lead SNPs**

SNPs reaching at least genome-wide significance ($p$ = 5x$10^{-8}$) in either $META_{US}$ or $META_{UK}$ permutations and showing similar phenotypic effects in the trait identified during each permutation. For each SNP, this table contains chromosome, position (GRCh37/hg19), cytogenetic band, allele information, the frequency of the A1 allele in the US sample ($A1_{freq}$ US), the frequency of the of the A1 allele in the UK ($A1_{freq}$ UK), the lowest meta-analysis p-value (Lowest P), the segment where the lowest p-value was found for either meta-analysis permutation (Best Segment), the permutation in which the best segment was found (Best Permutation), whether significance was reached in the Best Segment for both meta-analysis p-values (GWAS Sym Best), the p-value of the regression of slopes for comparing the effect of each SNP in the $META_{US}$ and $META_{UK}$ permutations (P Reg Slopes). We have also included information on whether this SNP was considered to be part of a multi-peak locus (Multipeak), the genomic context gathered from Ensembl[30], the candidate gene to which this signal was annotated, references to previous literature associating either the annotated gene or genomic region ± 250kb of the SNP to facial morphology, and whether we categorized this GWAS hit as being novel or previously implicated in craniofacial morphology by GWAS of normal-range facial variation, human malformations, or animal models. The "Citation" tab contains the DOI for literature referenced.

**Additional Table S4 (Separate File). 24 multi-effect loci**

Listed are SNP pairs within 250 kb of each other that are associated with different facial effects in the segment in which they reached lowest meta-analysis p-values. For each SNP, this table contains the cytogenetic band, chromosome, position (GRCh37/hg19), allele information, the frequency of the A1 allele in the US sample ($A1_{freq}$ US), the frequency of the A1 allele in the UK ($A1_{freq}$ UK), the lowest meta-analysis p-value (Lowest P), the genomic context gathered from Ensembl[30], the candidate gene to which this signal was annotated, and the $r^2$ and D' values for the SNP pairs, based on the 1000 Genomes Phase 3 EUR sample and calculated using LDPair[31]. All pairs have $r^2$ lower than 0.05. To accompany this file, we have generated LocusZoom and facial effect plots for each SNP (Fig. S10).

**Additional Table S5 (Separate File). 13 multi-effect SNPs**

SNPs identified in the cross-peak association that had associations with additional traits spanning multiple facial quadrants. For each SNP, this table contains chromosome, position (GRCh37/hg19), gene annotation, the segment where the additional association occurred, the trait number(s) with which the SNP was additionally associated, the H3K27 cluster membership (refer to Fig. 3), and the genomic context, regulatory features at the SNP position, and regulatory features within ± 10kb of the SNP position, all gathered from Ensembl[30]. The "Trait Key" tab contains the conversion of trait number to the SNP with which that trait was additionally identified during the CCA.

**Additional Table S6 (Separate File). Structural Equation Modeling results**
For each of the 50 segments with well-fitting SEM models, in this table we provide the number of principal components included to represent shape variation in that segment, the number of SNPs that survived the model refinement process (see Methods), the p-value cutoff used to perform the model refinement and determine the SNPs to be used for epistasis, the number of SNPs used in the epistasis analysis for this segment, and values for the $\chi^2$, CFI, RMSE, SRMR model fit indices, which were used to evaluate the models for our analysis. We also include the TLI and GFI model fit indices for completeness. This table also contains internal links to separate tabs where, for each surviving model, we have listed the parameters used and the estimate, standard error, z-score, p-value, and 95% confidence intervals. SNPs which were selected for epistasis are highlighted in green.
