## Supplementary material for "Insights into the genetic architecture of the human face": Figure S10

### Supplemental figure 10

#### Multi-peak loci overview

J. White, K. Indencleef et al.

### Multi-peak locus 1: rs76244841

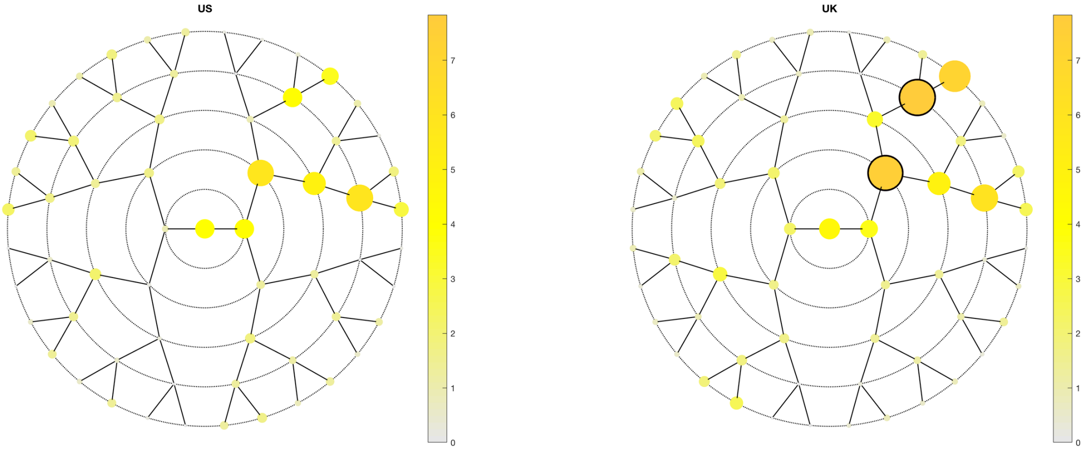

**A**

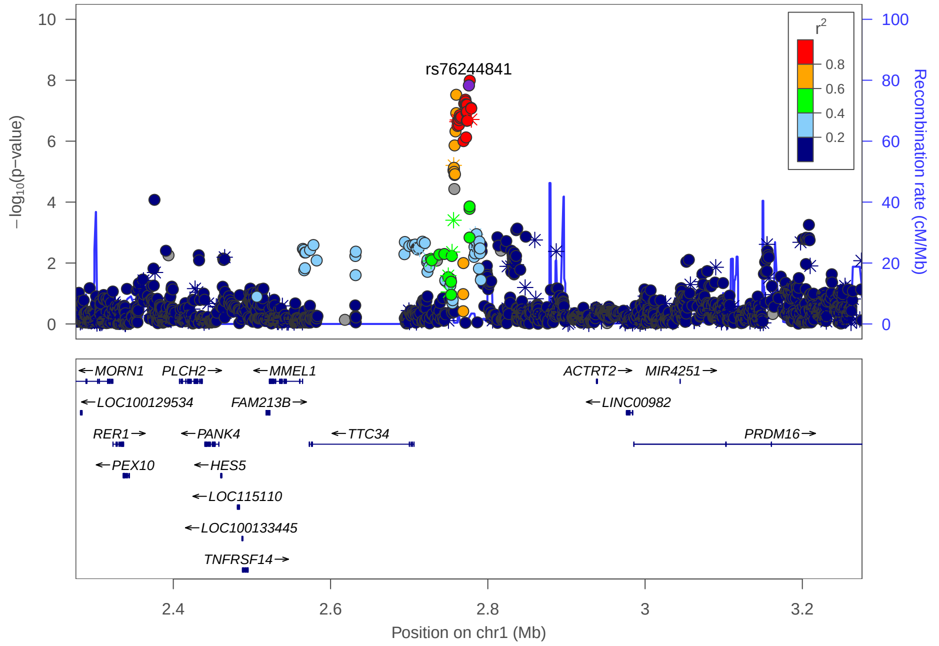

**C**

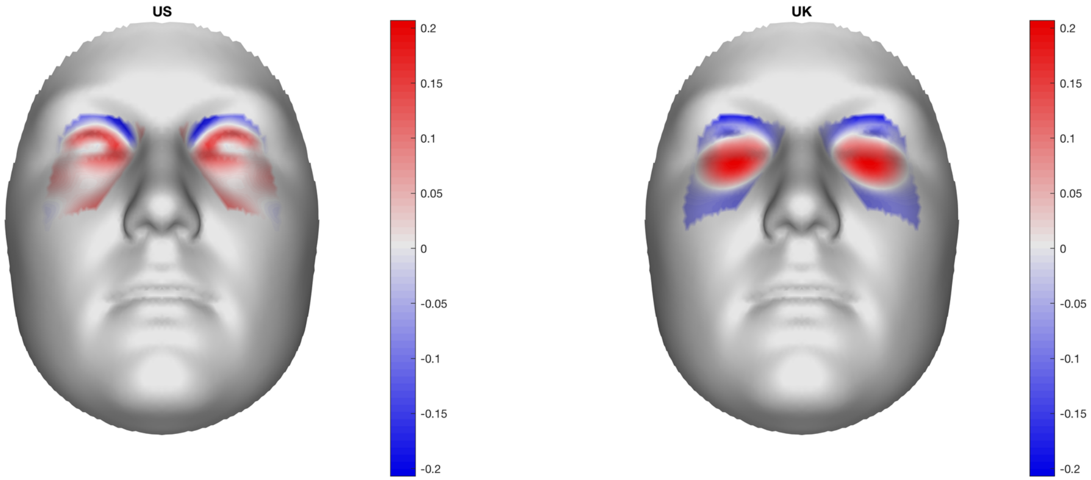

**B**

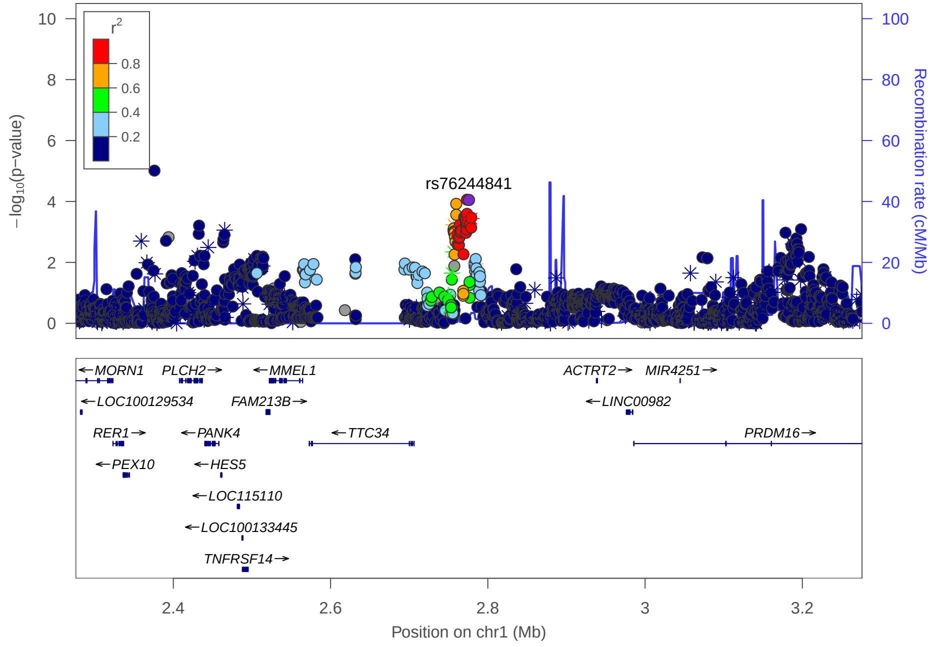

#### Multi-peak locus 1: rs1572037

**CHR1** - See final page for captions

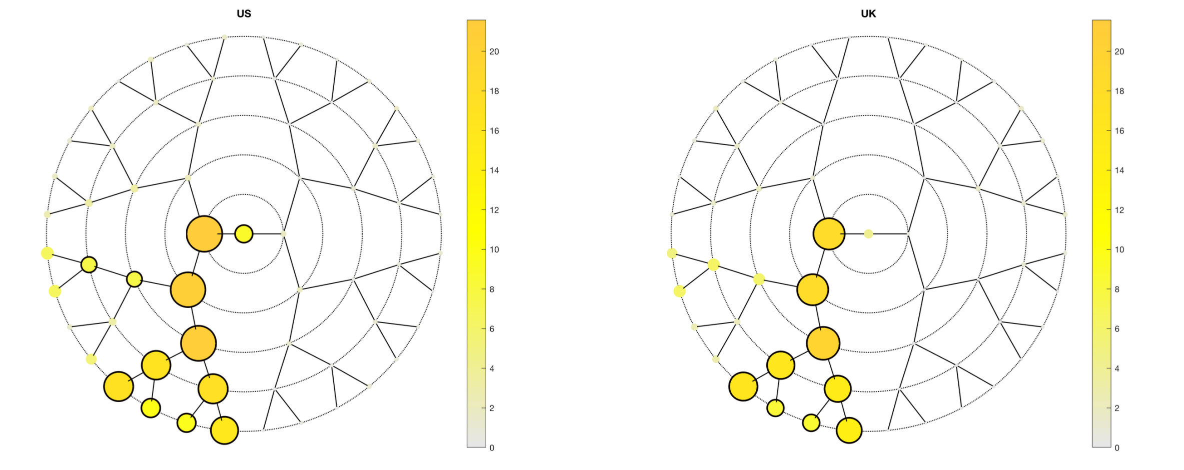

**A**

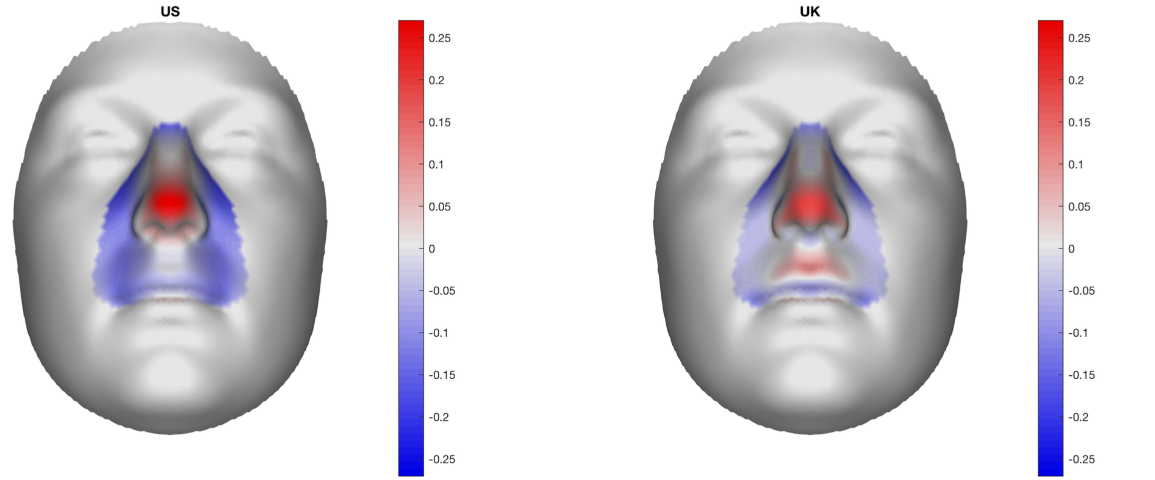

**B**

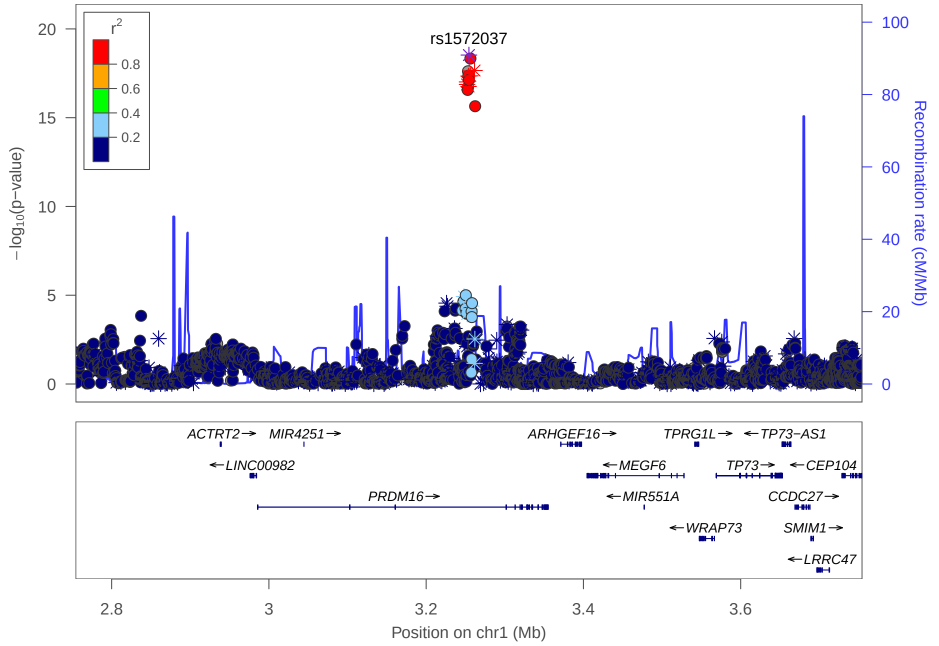

**C**

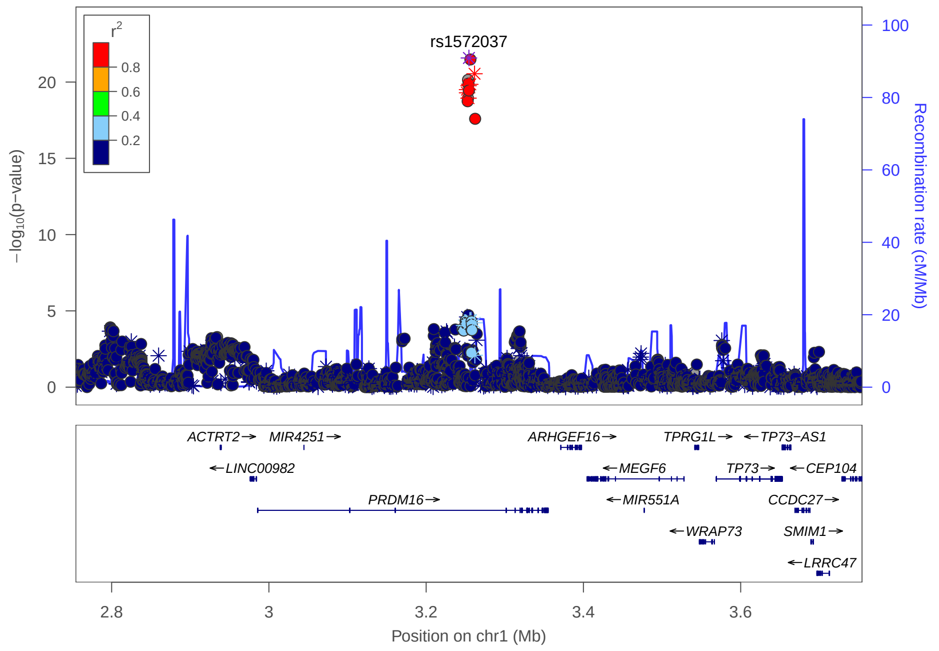

### Multi-peak locus 2: rs3936018

**CHR1** - See final page for captions

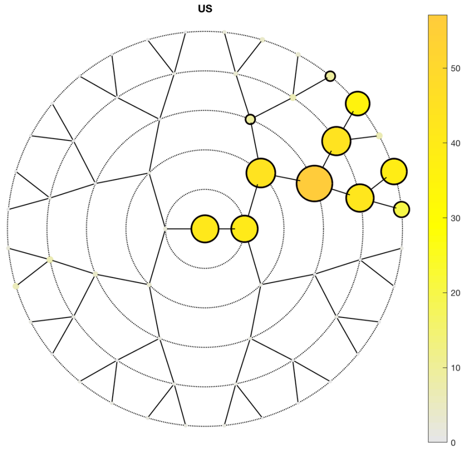

**A**

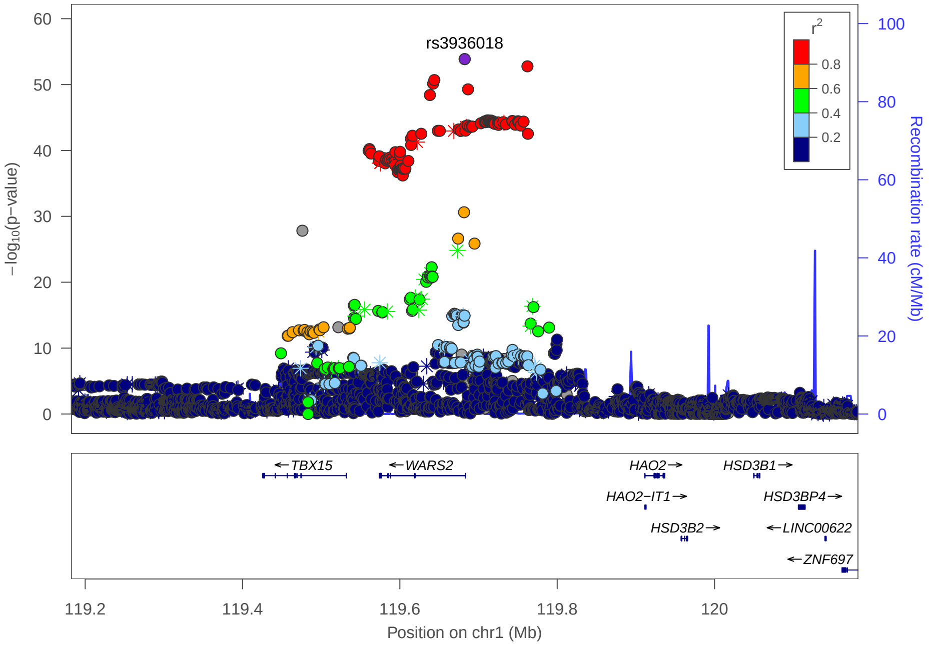

**C**

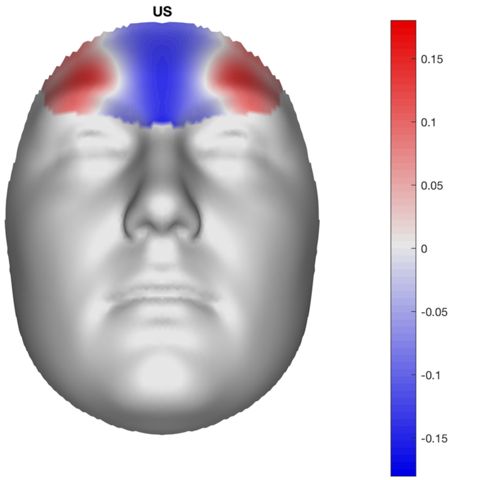

**B**

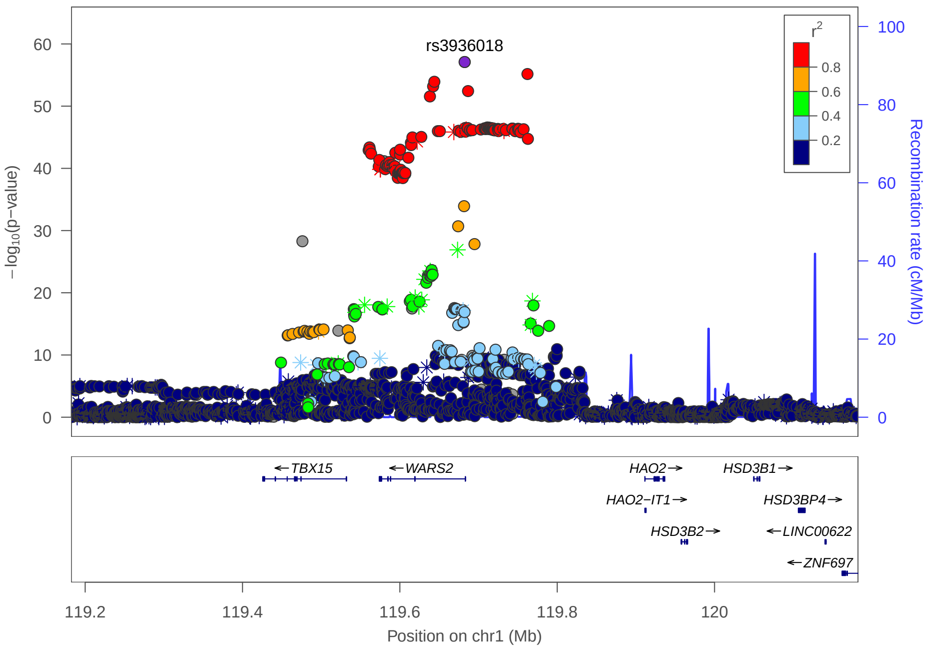

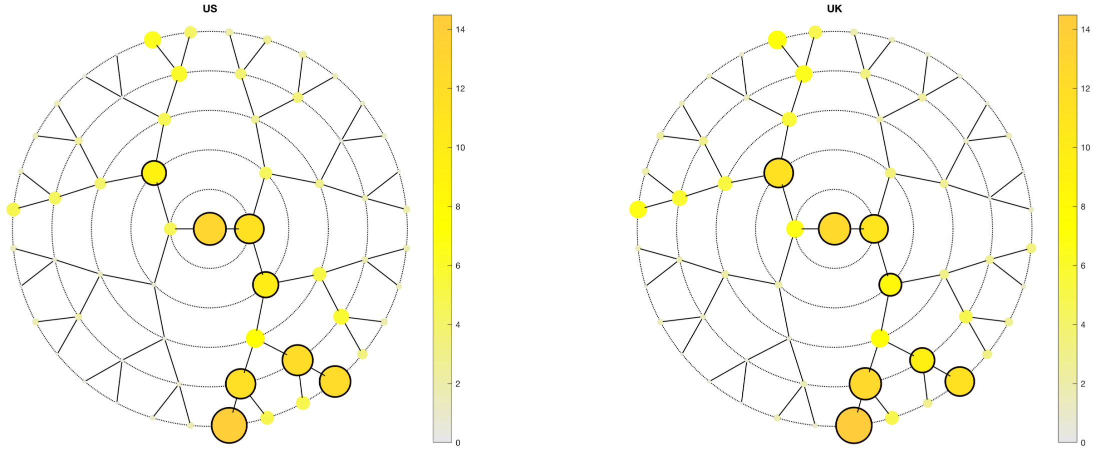
US
UK
14
12
10
8
6
4
2
0

A

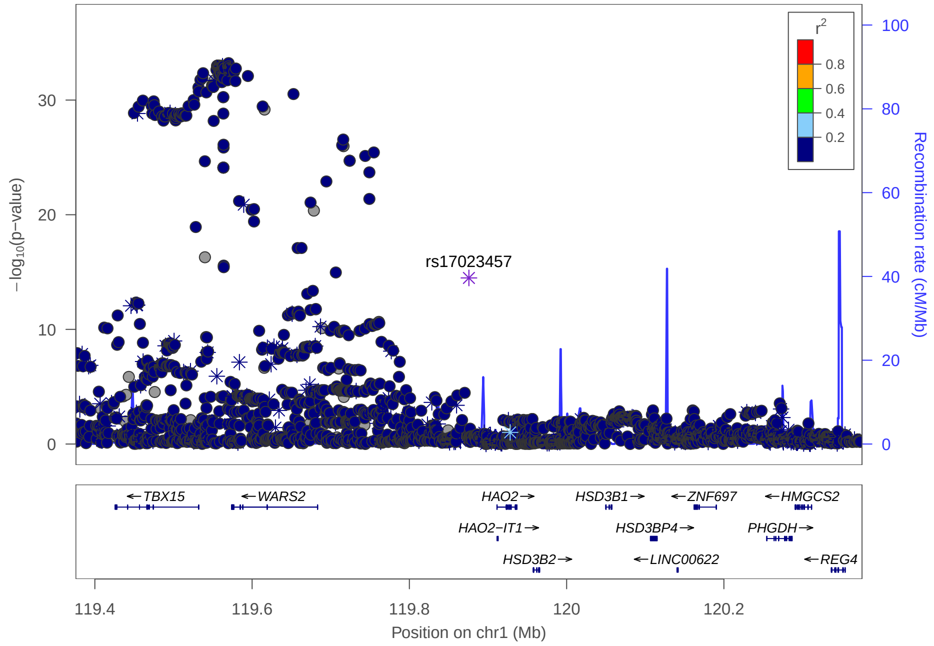
r$^2$
0.8
0.6
0.4
0.2
rs17023457
$-\log_{10}$(p−value)
Recombination rate (cM/Mb)
TBX15
WARS2
HAO2
HSD3B1
ZNF697
HMGCS2
HAO2−IT1
HSD3BP4
PHGDH
HSD3B2
LINC00622
REG4
119.4
119.6
119.8
120
120.2
Position on chr1 (Mb)

C

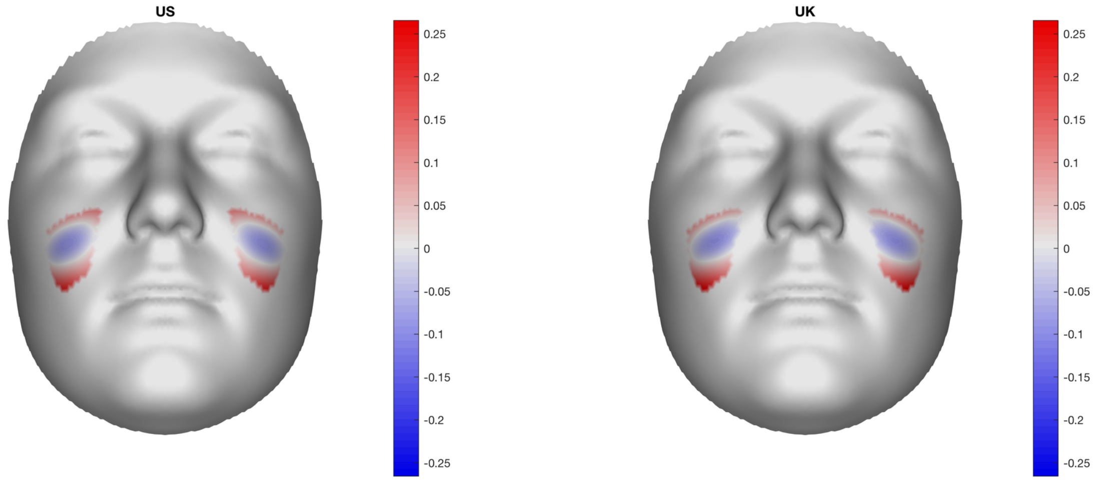
US
UK
0.25
0.2
0.15
0.1
0.05
0
-0.05
-0.1
-0.15
-0.2
-0.25

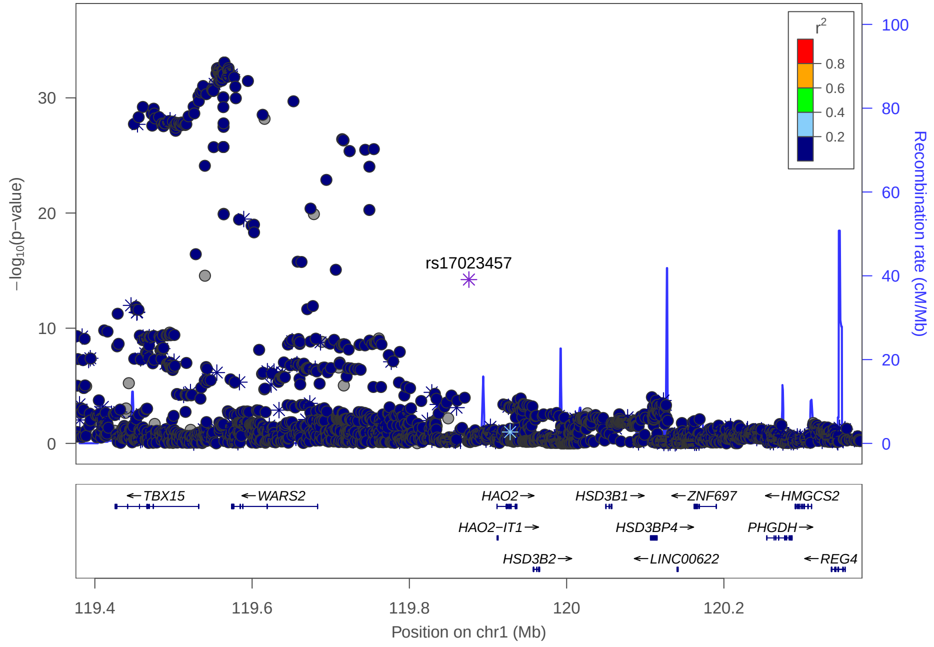
r$^2$
0.8
0.6
0.4
0.2
rs17023457
$-\log_{10}$(p−value)
Recombination rate (cM/Mb)
TBX15
WARS2
HAO2
HSD3B1
ZNF697
HMGCS2
HAO2−IT1
HSD3BP4
PHGDH
HSD3B2
LINC00622
REG4
119.4
119.6
119.8
120
120.2
Position on chr1 (Mb)

B

### Multi-peak locus 3: rs577676

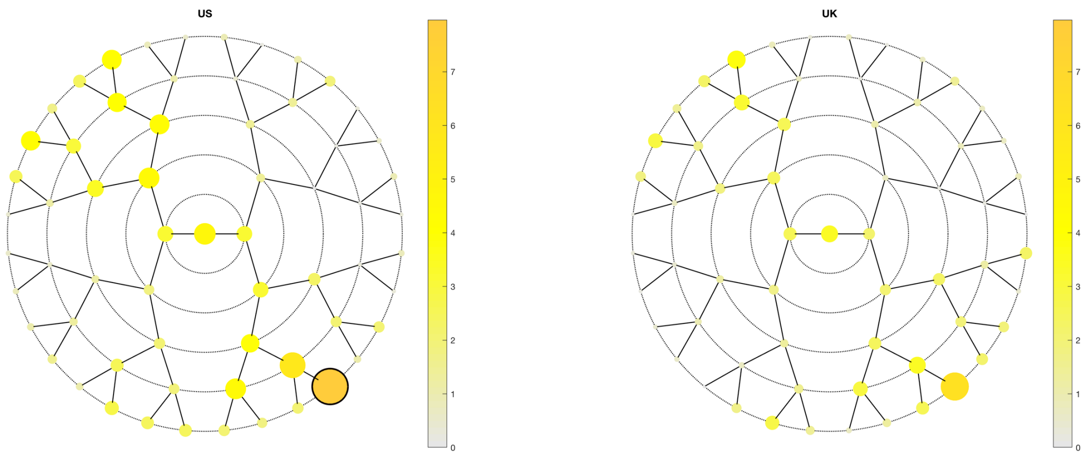

**A**

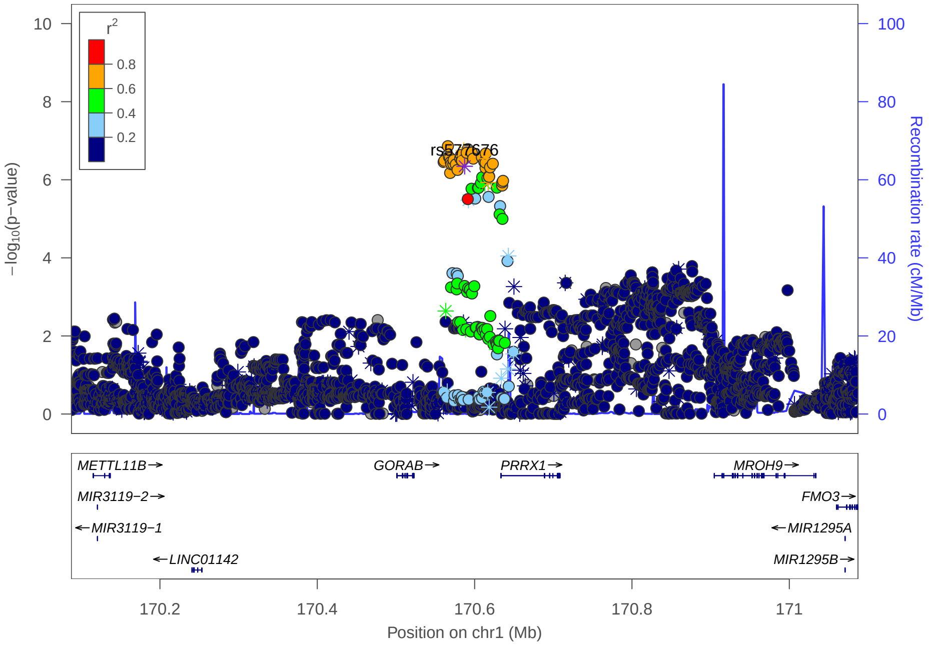

**C**

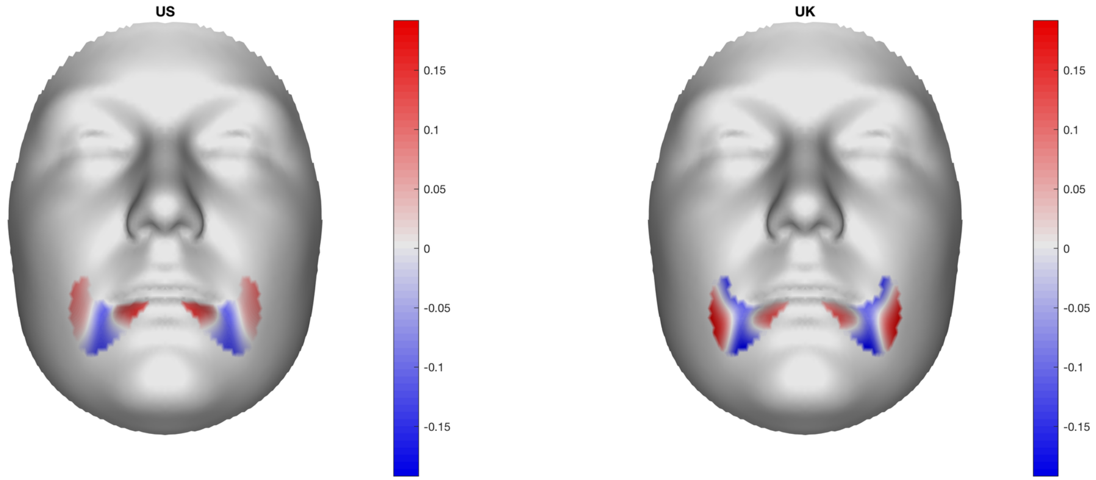

**B**

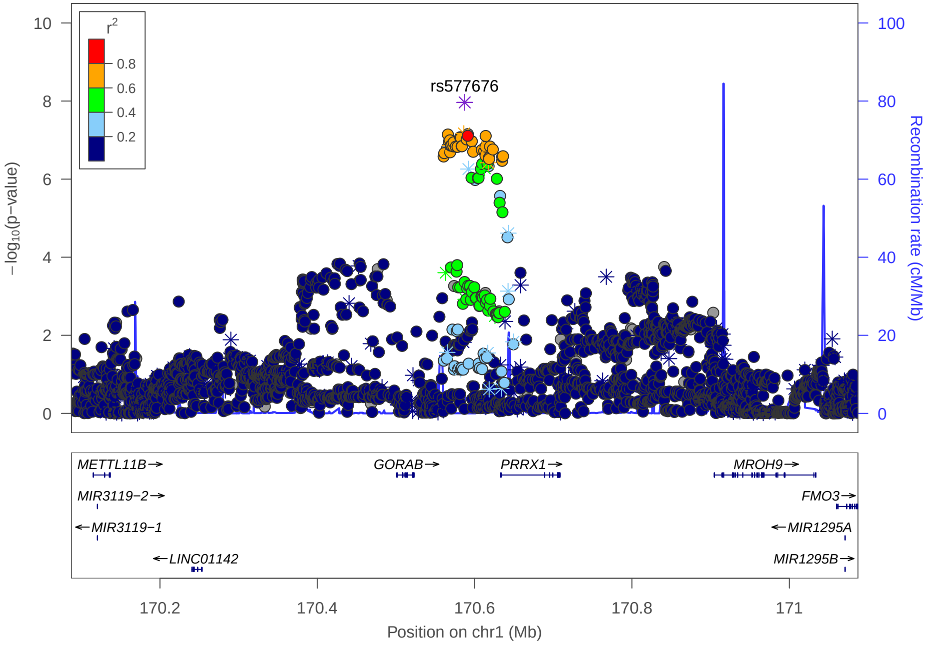

### Multi-peak locus 3: rs10919462

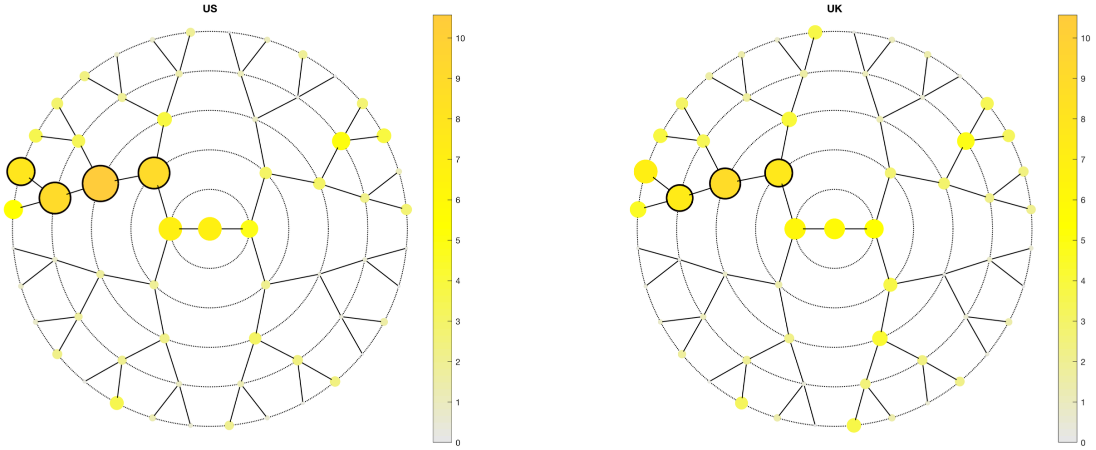

A

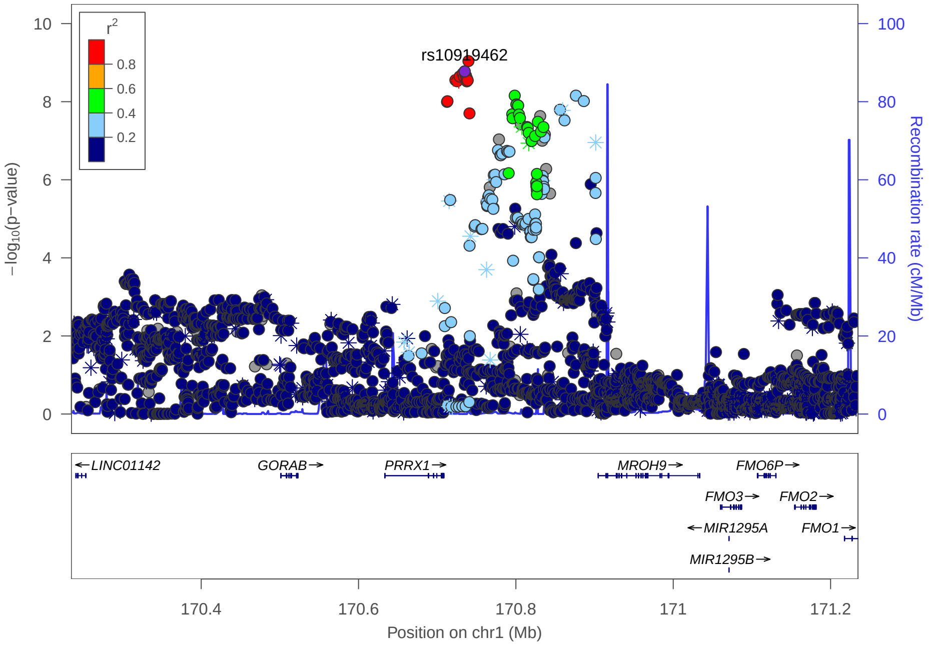

C

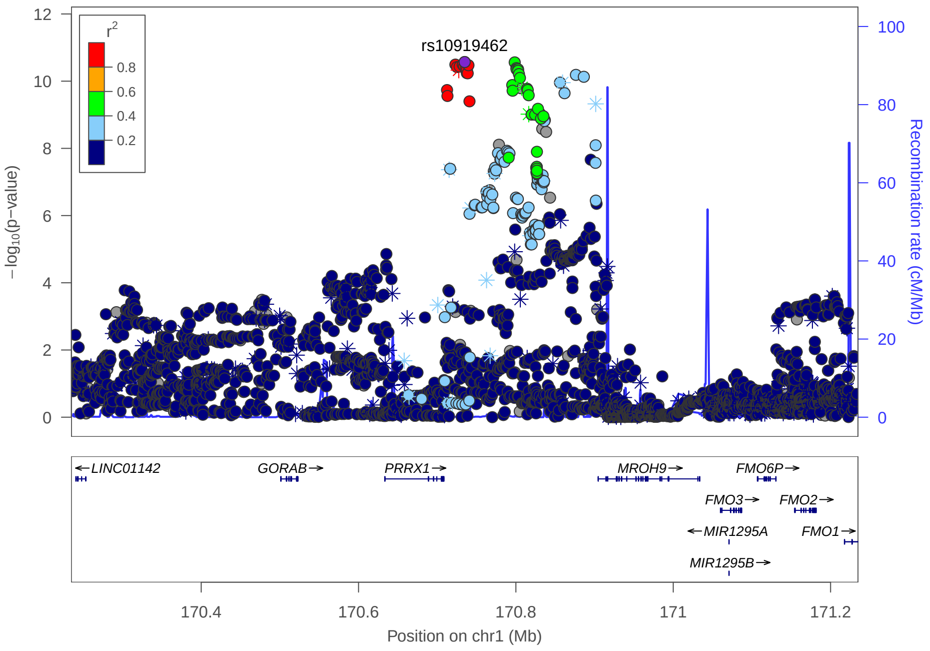

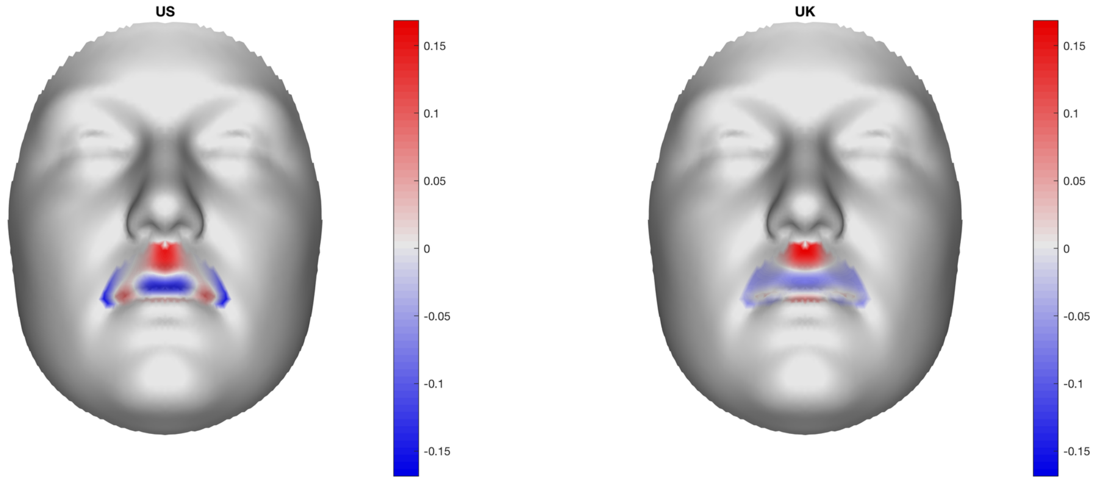

B

### Multi-peak locus 4: rs6715010

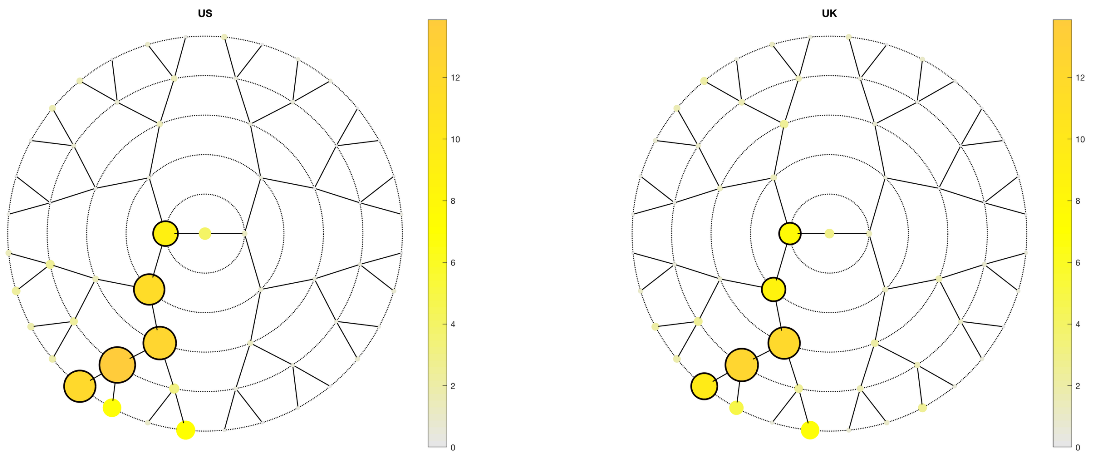

**A**

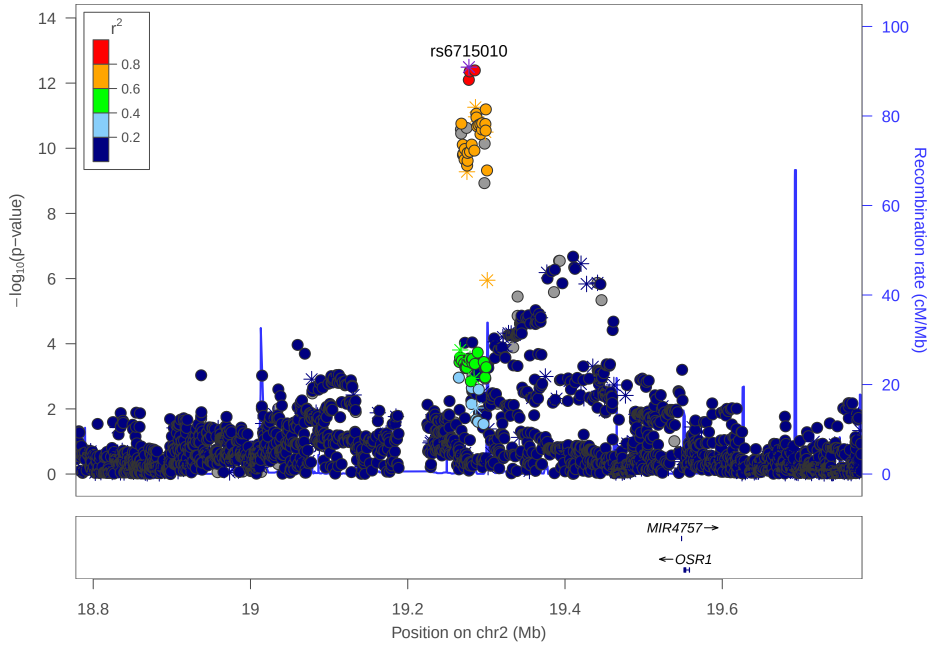

**C**

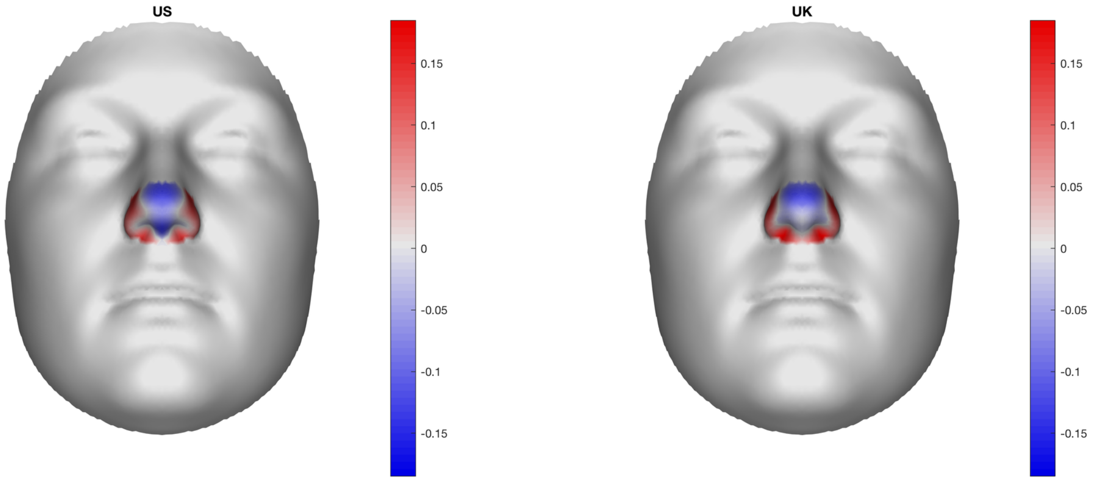

**B**

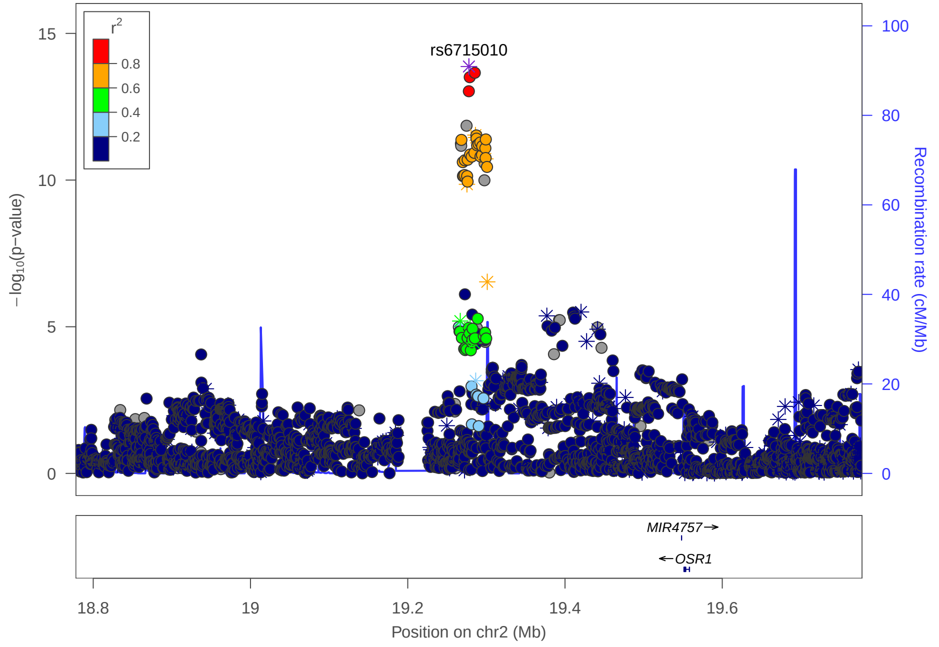

### Multi-peak locus 4: rs79037251

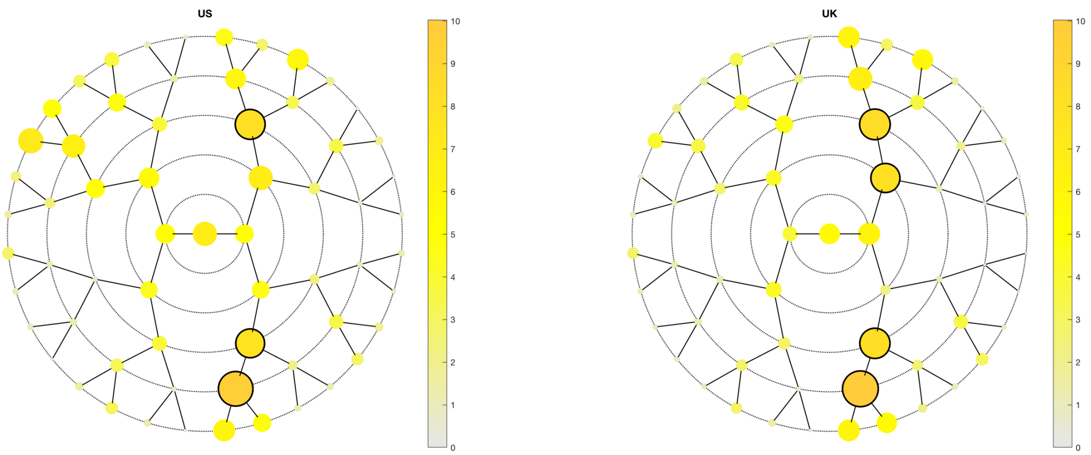

**A**

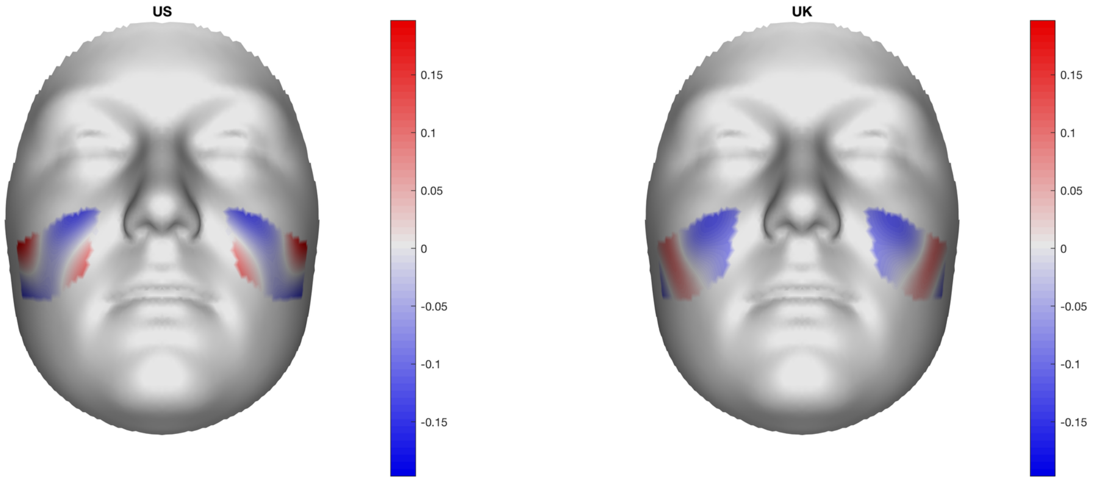

**B**

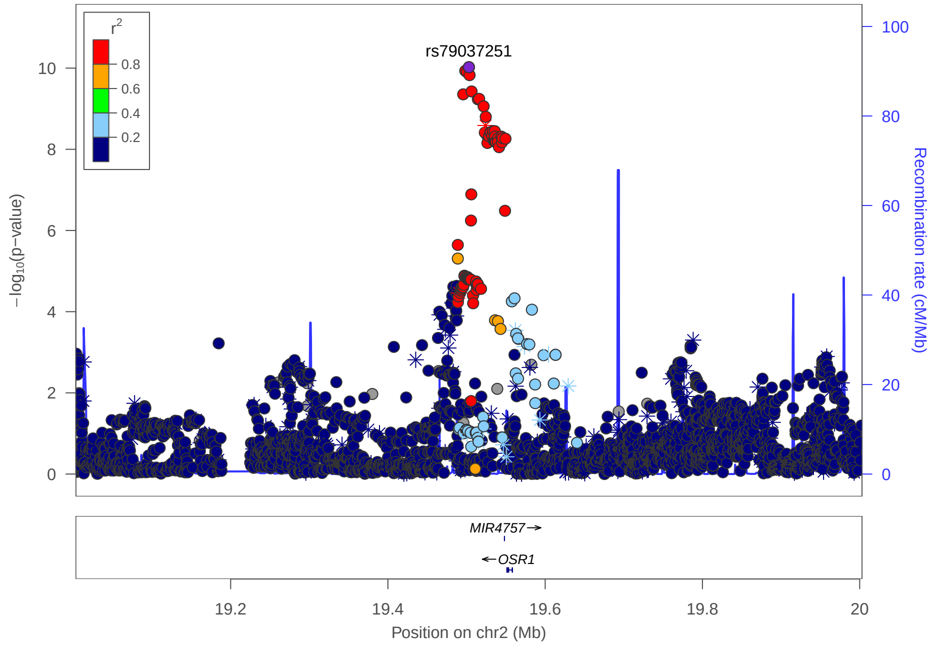

**C**

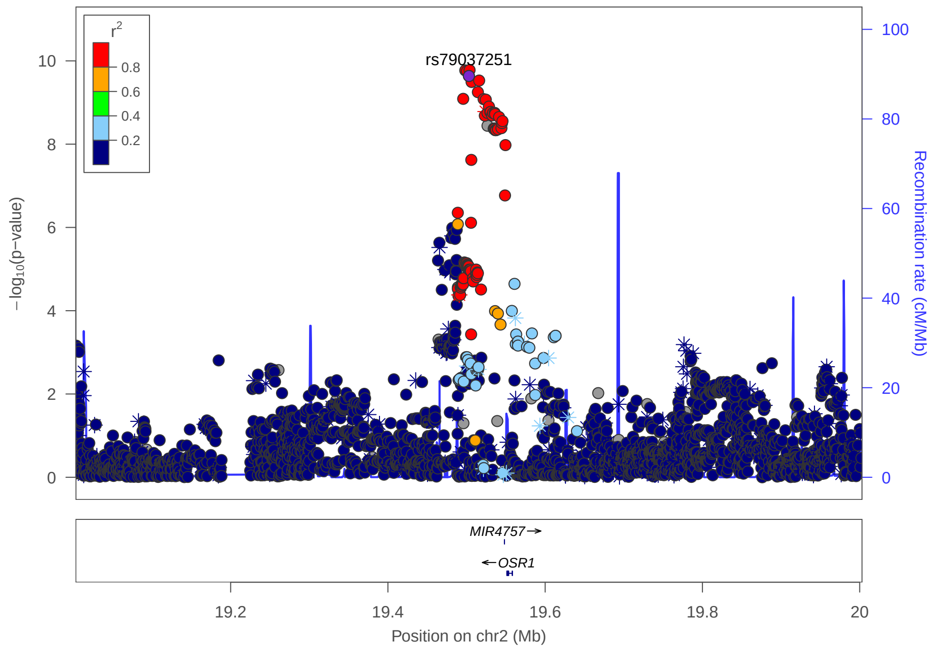

### Multi-peak locus 4: rs1427539

**CHR2** - See final page for captions

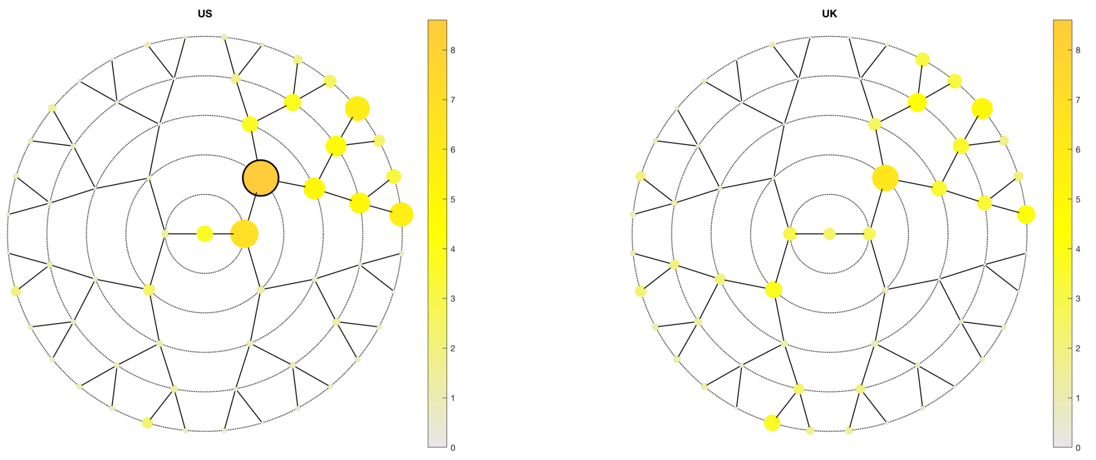

**A**

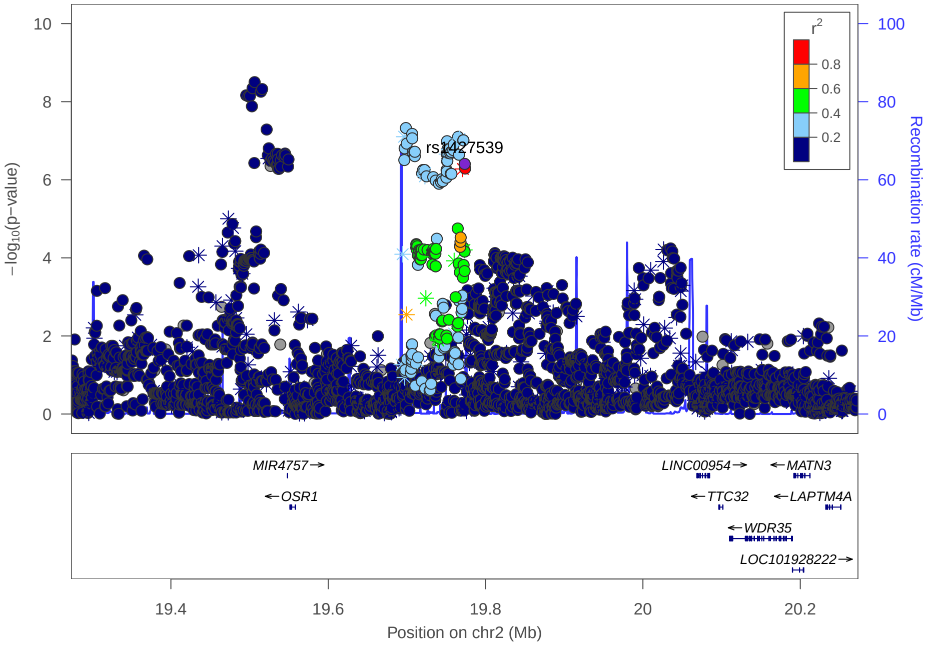

**C**

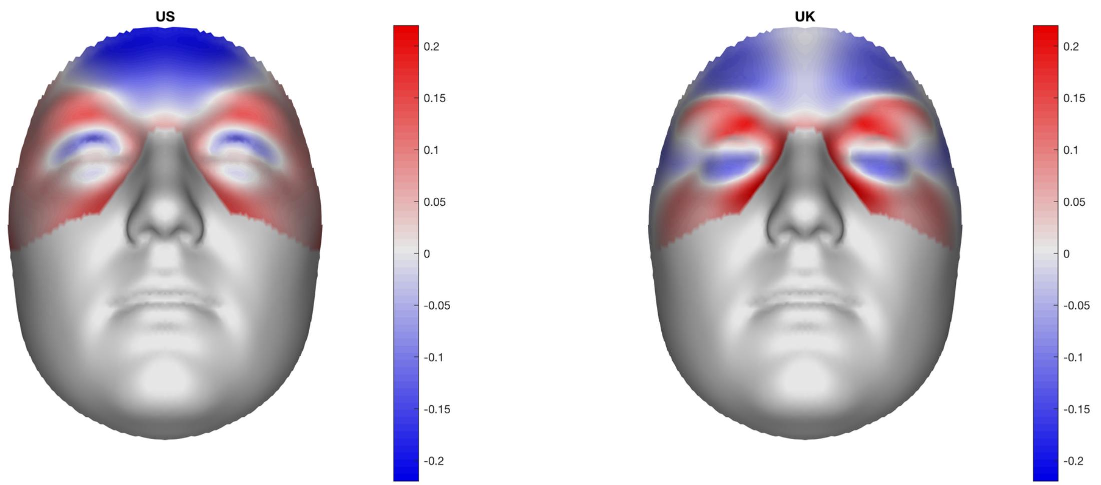

**B**

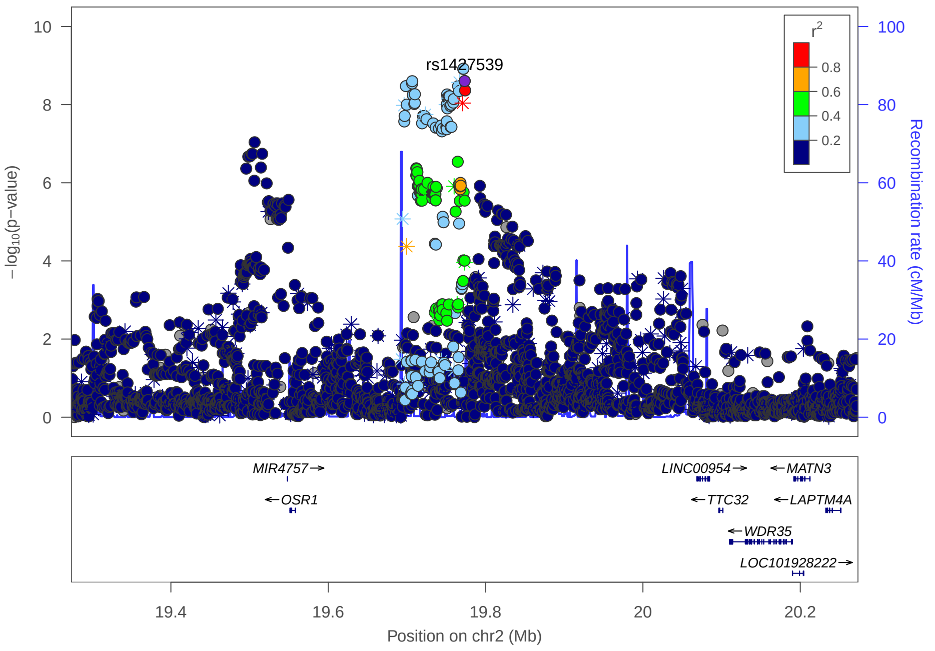

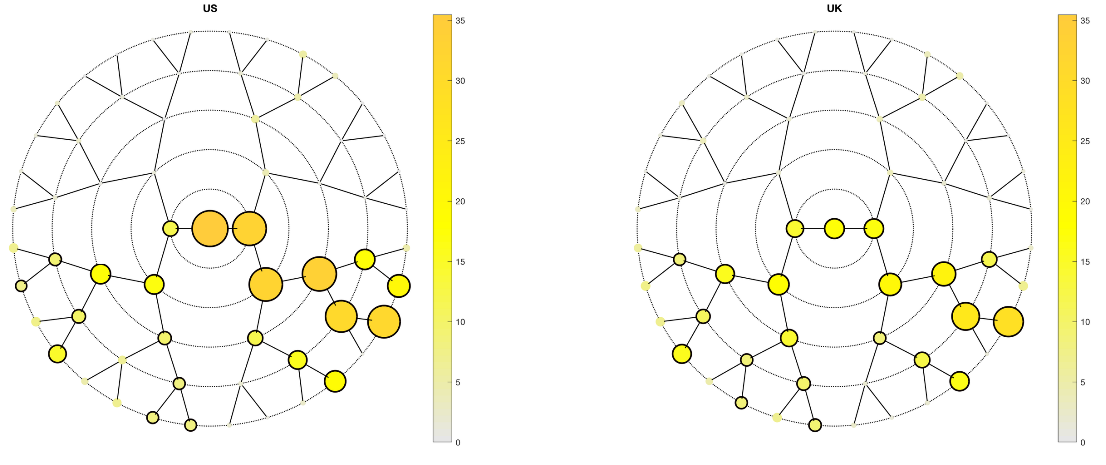
US
UK
35
30
25
20
15
10
5
0

**A**

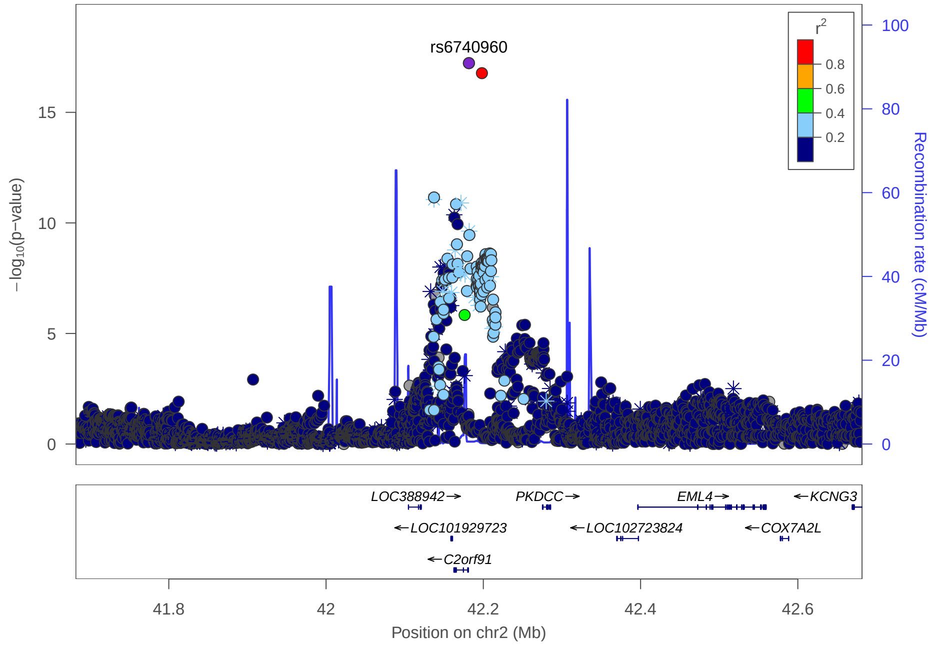
rs6740960
$r^2$
0.8
0.6
0.4
0.2
−log10(p−value)
Recombination rate (cM/Mb)
LOC388942→
PKDCC→
EML4→
←KCNG3
←LOC101929723
←LOC102723824
←COX7A2L
←C2orf91
41.8
42
42.2
42.4
42.6
Position on chr2 (Mb)

**C**

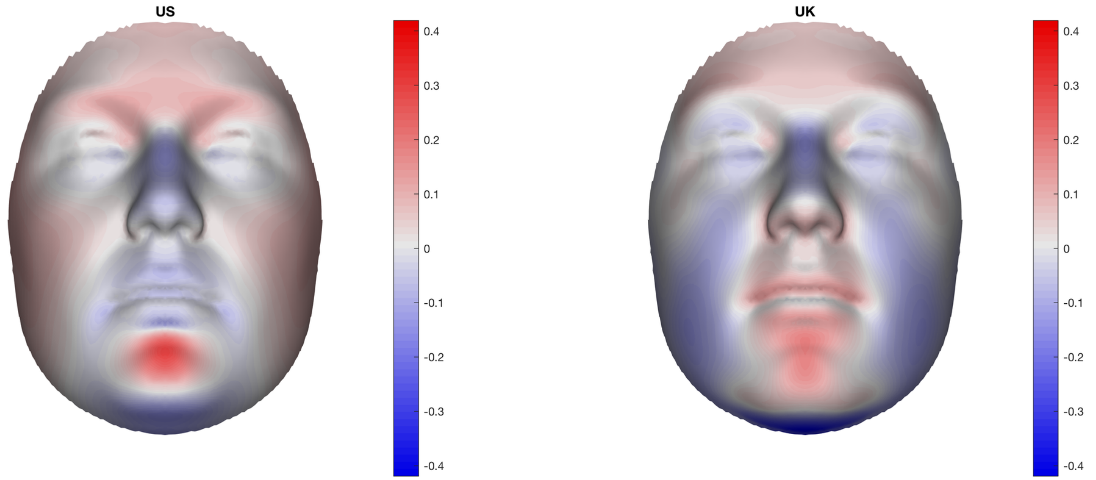
US
UK
0.4
0.3
0.2
0.1
0
-0.1
-0.2
-0.3
-0.4

**B**

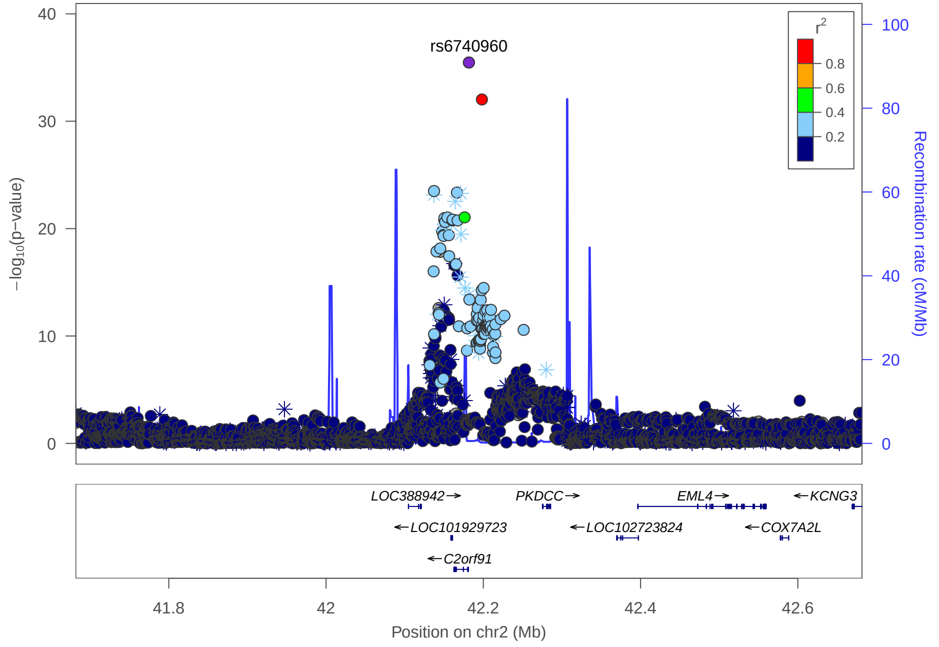
rs6740960
$r^2$
0.8
0.6
0.4
0.2
−log10(p−value)
Recombination rate (cM/Mb)
LOC388942→
PKDCC→
EML4→
←KCNG3
←LOC101929723
←LOC102723824
←COX7A2L
←C2orf91
41.8
42
42.2
42.4
42.6
Position on chr2 (Mb)

### Multi-peak locus 5: rs10189338

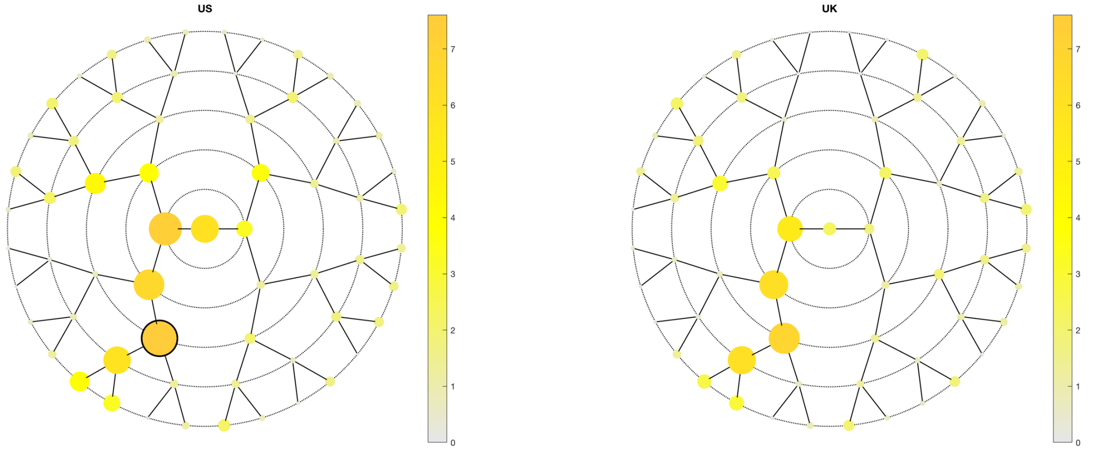

A

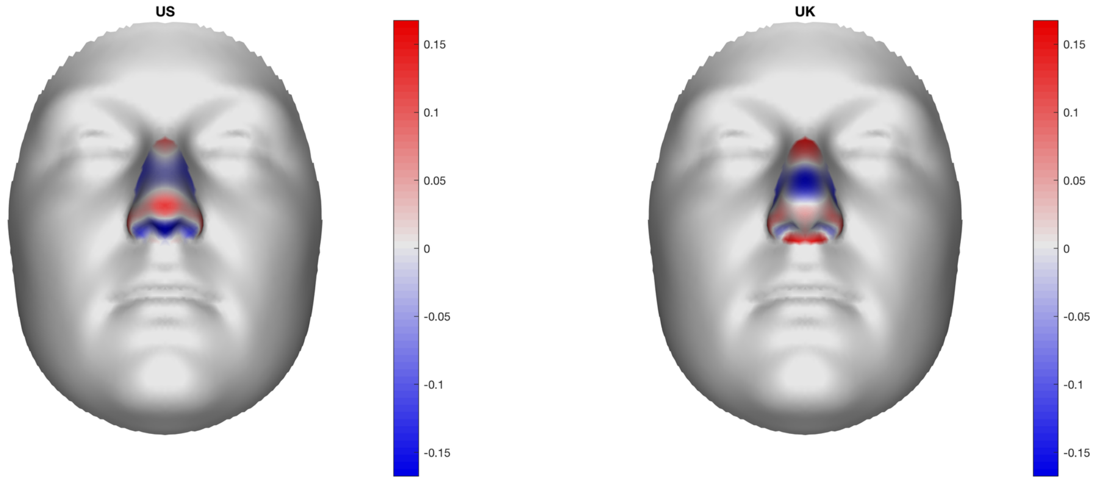

B

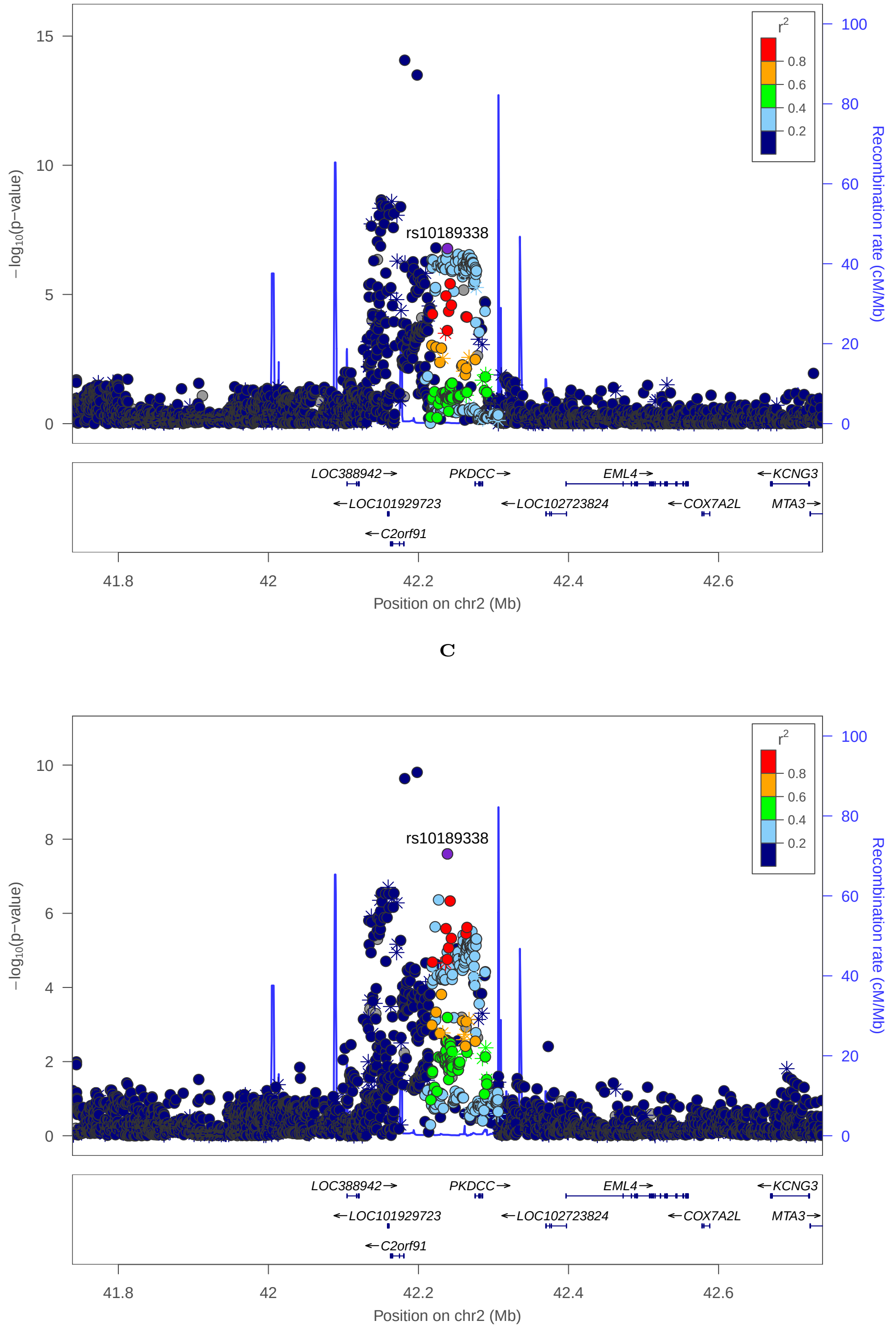

C

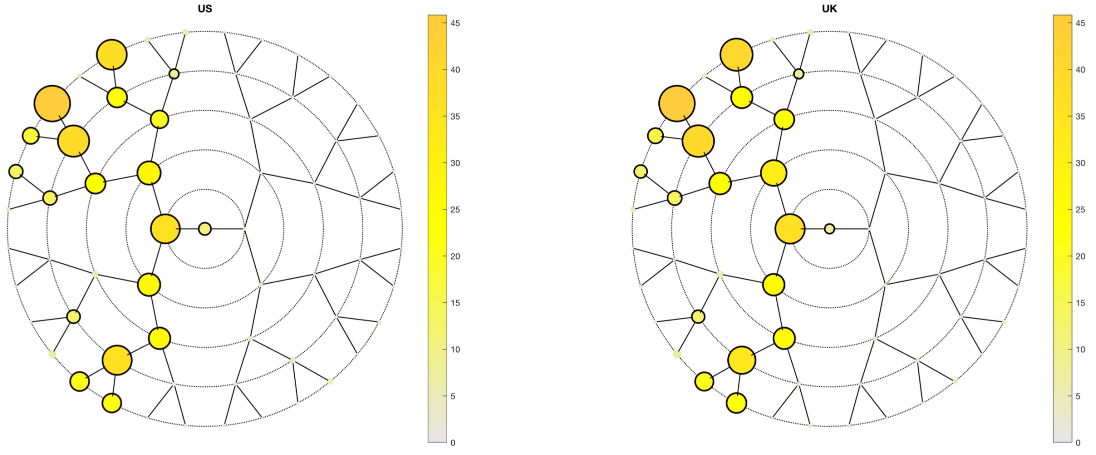
US
UK

**A**

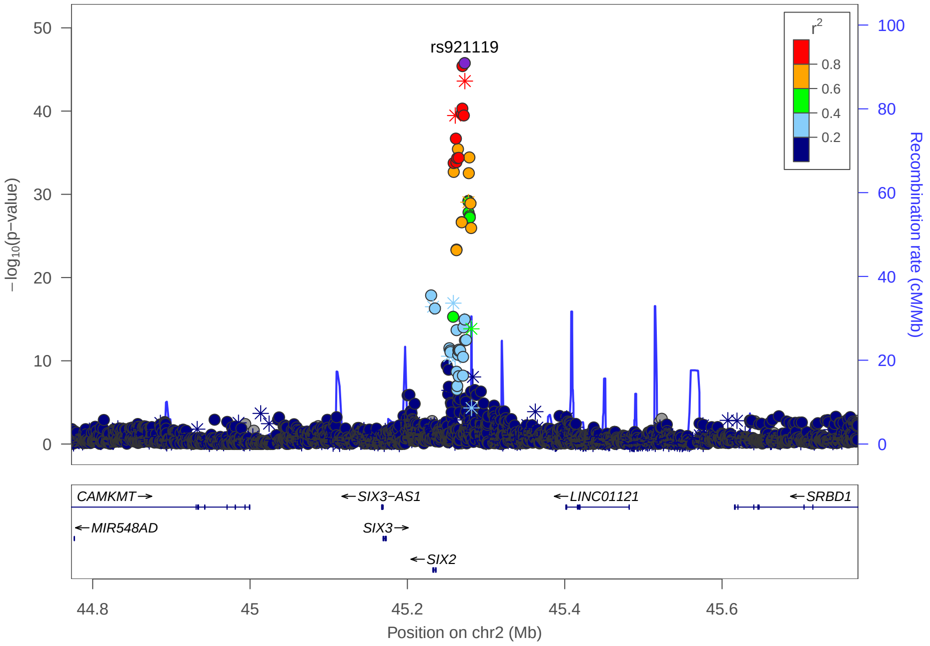
rs921119
−log10(p−value)
Recombination rate (cM/Mb)
r2
CAMKMT→
←SIX3−AS1
←LINC01121
←SRBD1
←MIR548AD
SIX3→
←SIX2
Position on chr2 (Mb)

**C**

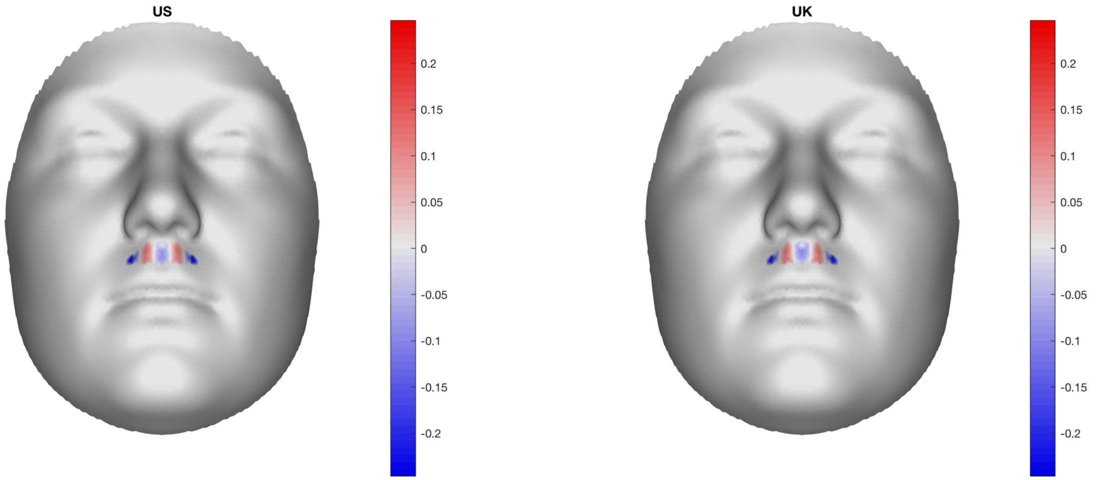
US
UK

**B**

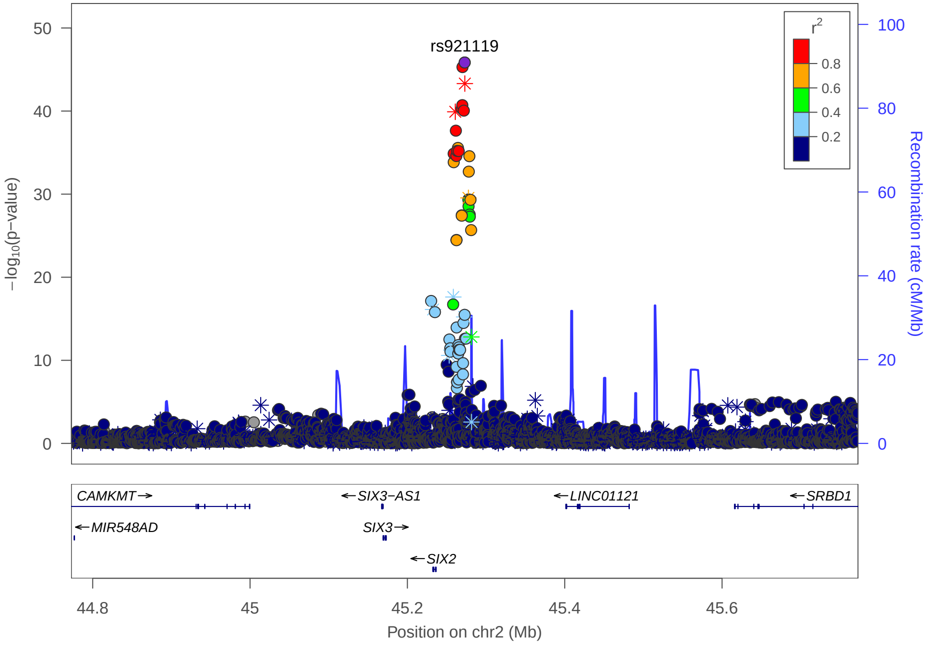
rs921119
−log10(p−value)
Recombination rate (cM/Mb)
r2
CAMKMT→
←SIX3−AS1
←LINC01121
←SRBD1
←MIR548AD
SIX3→
←SIX2
Position on chr2 (Mb)

### Multi-peak locus 6: rs35395759

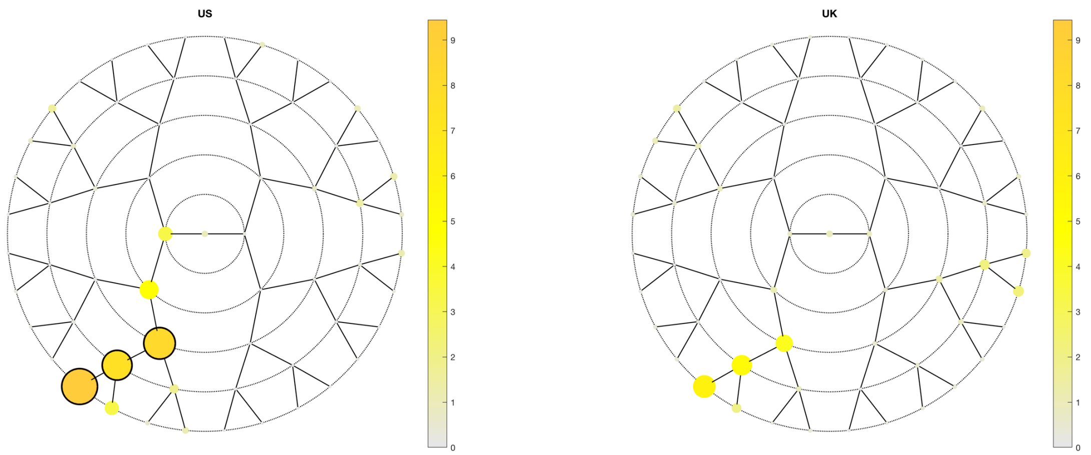

**A**

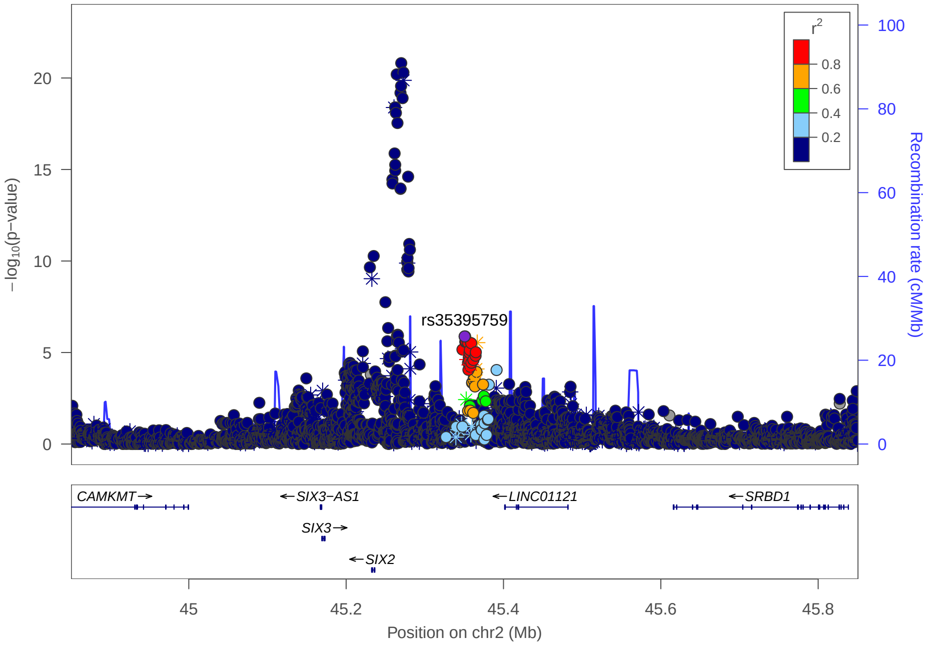

**C**

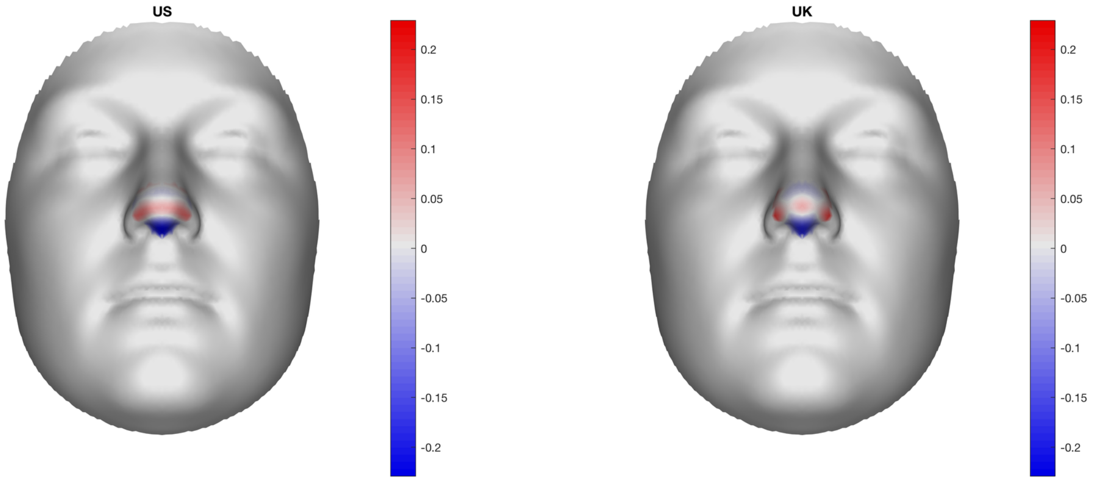

**B**

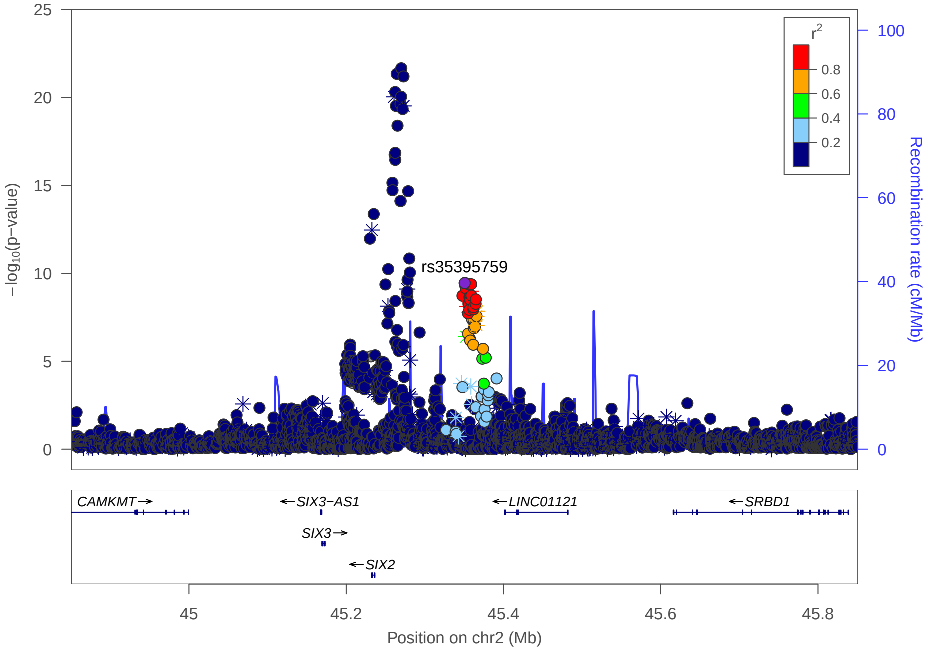

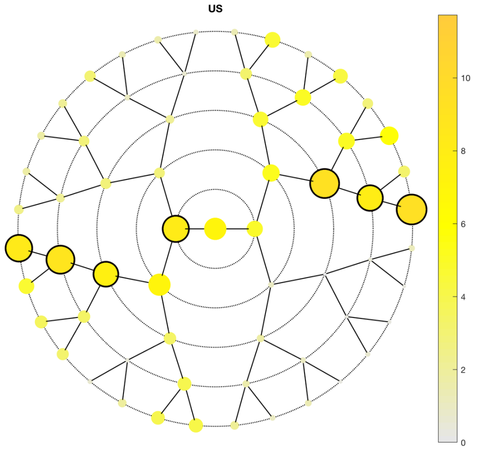
US
UK

**A**

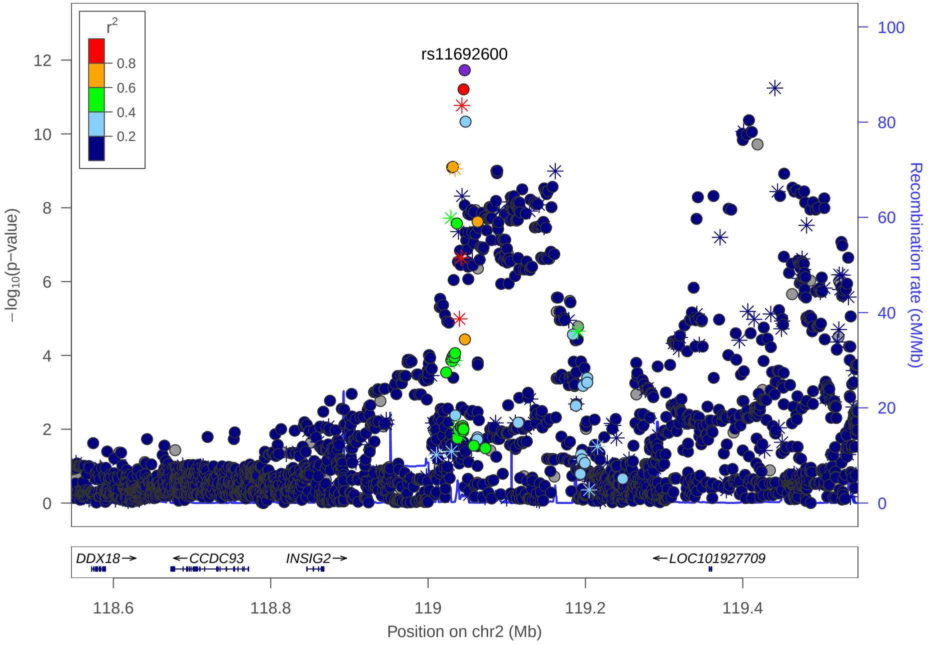
rs11692600
r2
−log10(p−value)
Recombination rate (cM/Mb)
DDX18
CCDC93
INSIG2
LOC101927709
Position on chr2 (Mb)

**C**

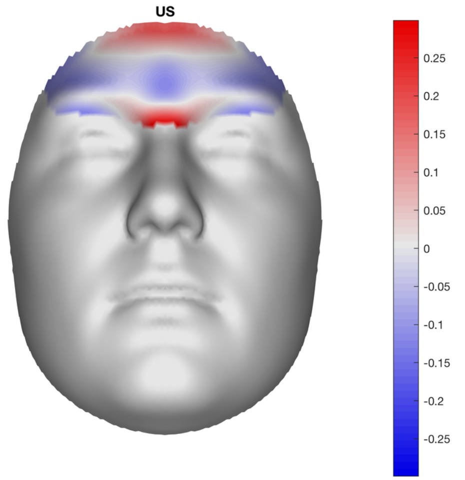
US
UK

**B**

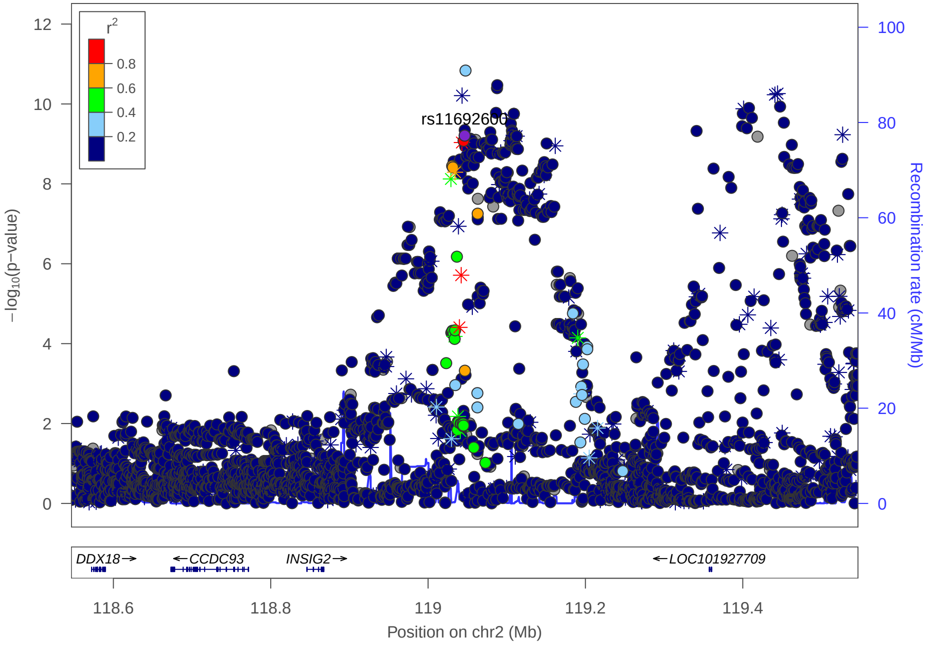
rs11692600
r2
−log10(p−value)
Recombination rate (cM/Mb)
DDX18
CCDC93
INSIG2
LOC101927709
Position on chr2 (Mb)

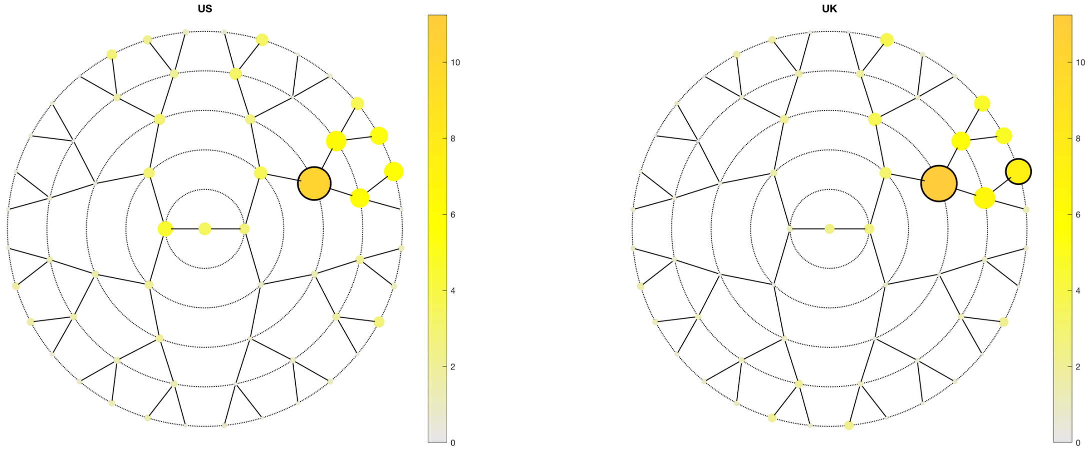
US
UK
0
2
4
6
8
10

**A**

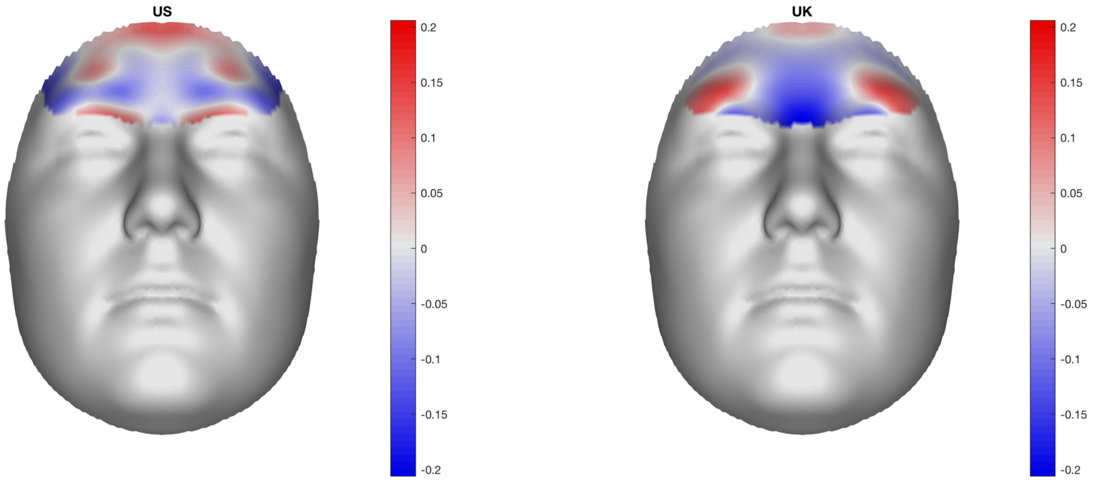
US
UK
0.2
0.15
0.1
0.05
0
-0.05
-0.1
-0.15
-0.2

**B**

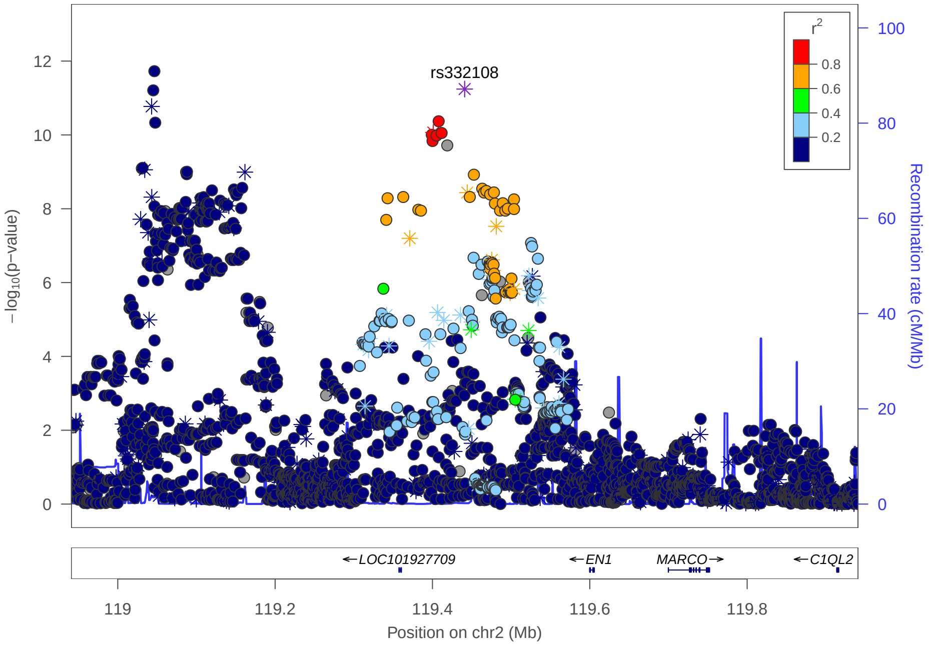
rs332108
r²
0.8
0.6
0.4
0.2
−log10(p−value)
Recombination rate (cM/Mb)
←LOC101927709
←EN1
MARCO→
←C1QL2
119
119.2
119.4
119.6
119.8
Position on chr2 (Mb)

**C**

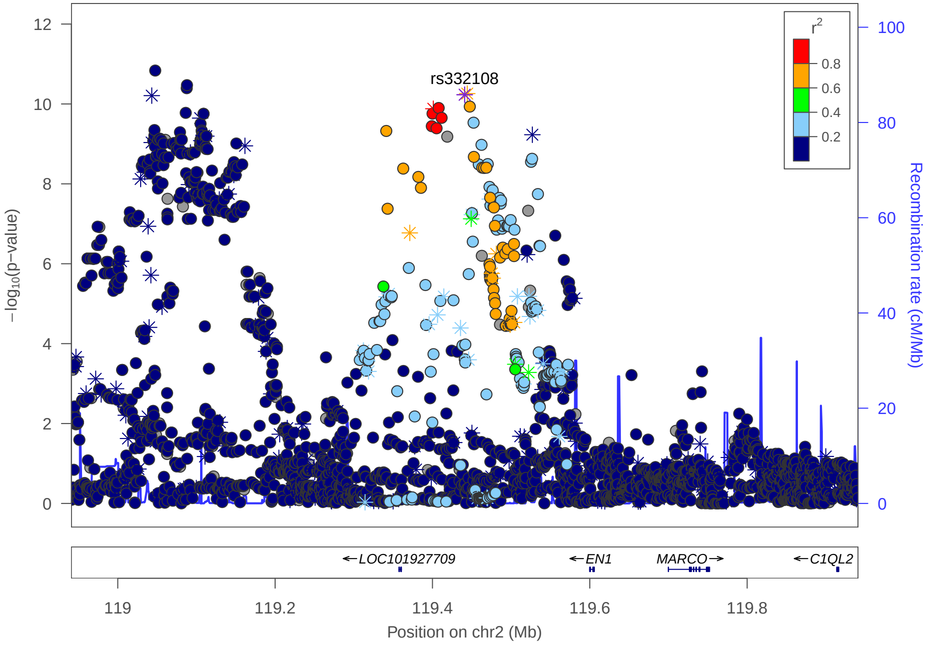
rs332108
r²
0.8
0.6
0.4
0.2
−log10(p−value)
Recombination rate (cM/Mb)
←LOC101927709
←EN1
MARCO→
←C1QL2
119
119.2
119.4
119.6
119.8
Position on chr2 (Mb)

### Multi-peak locus 8: rs970797

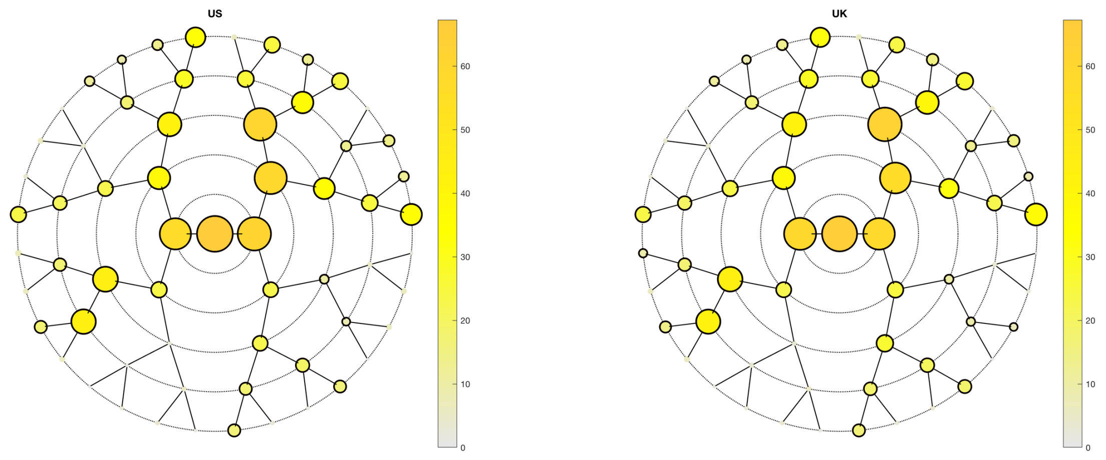

**A**

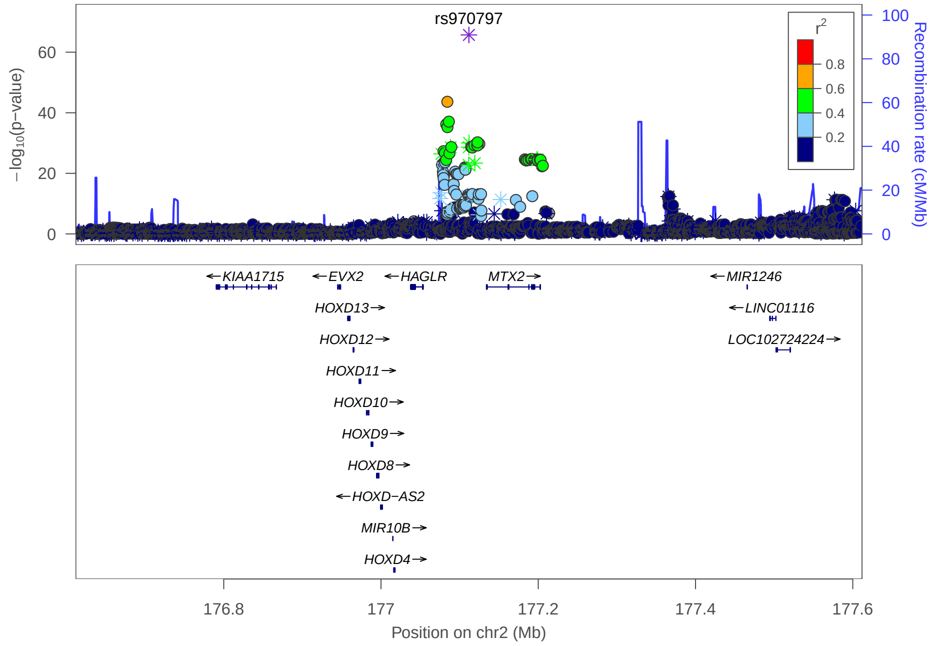

**C**

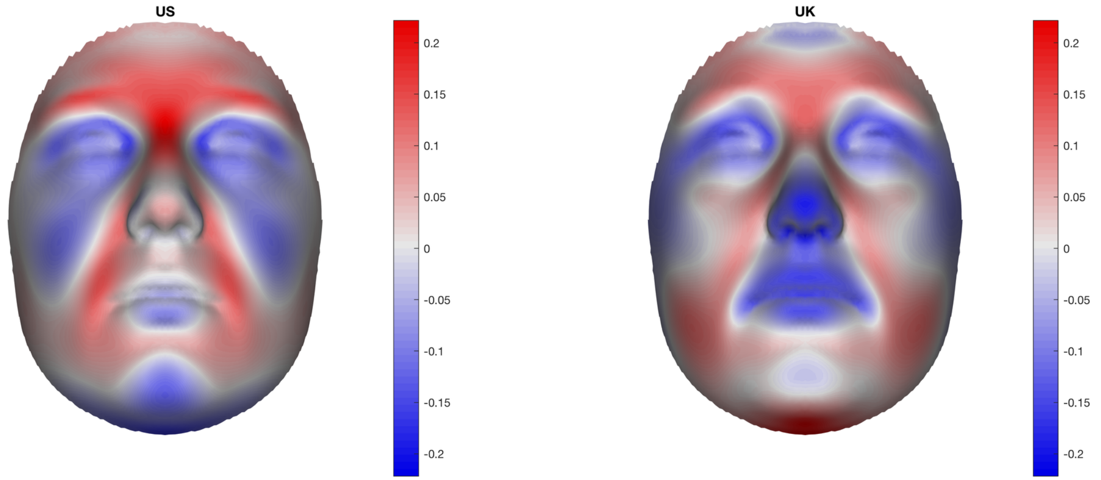

**B**

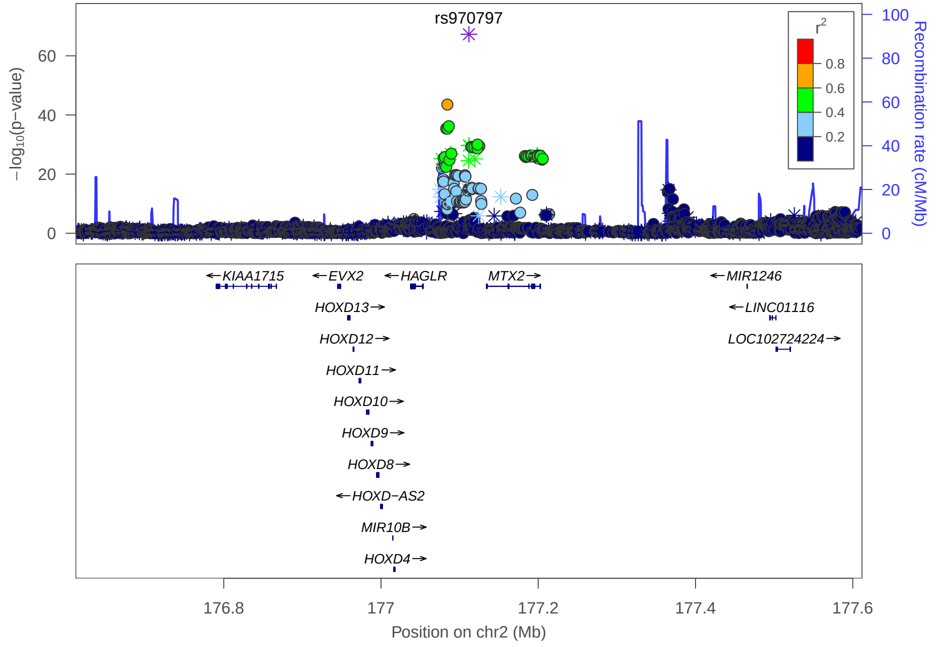

Multi-peak locus 8: rs10178696

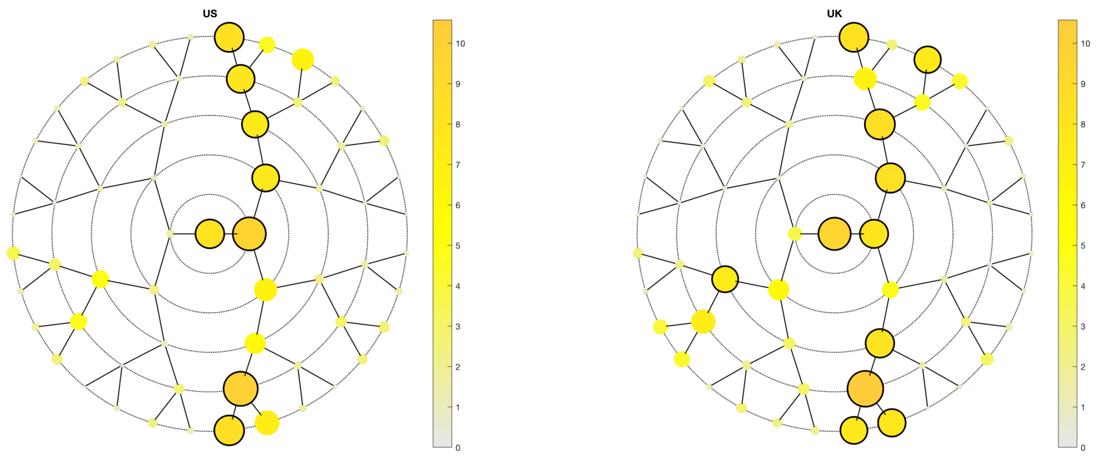

A

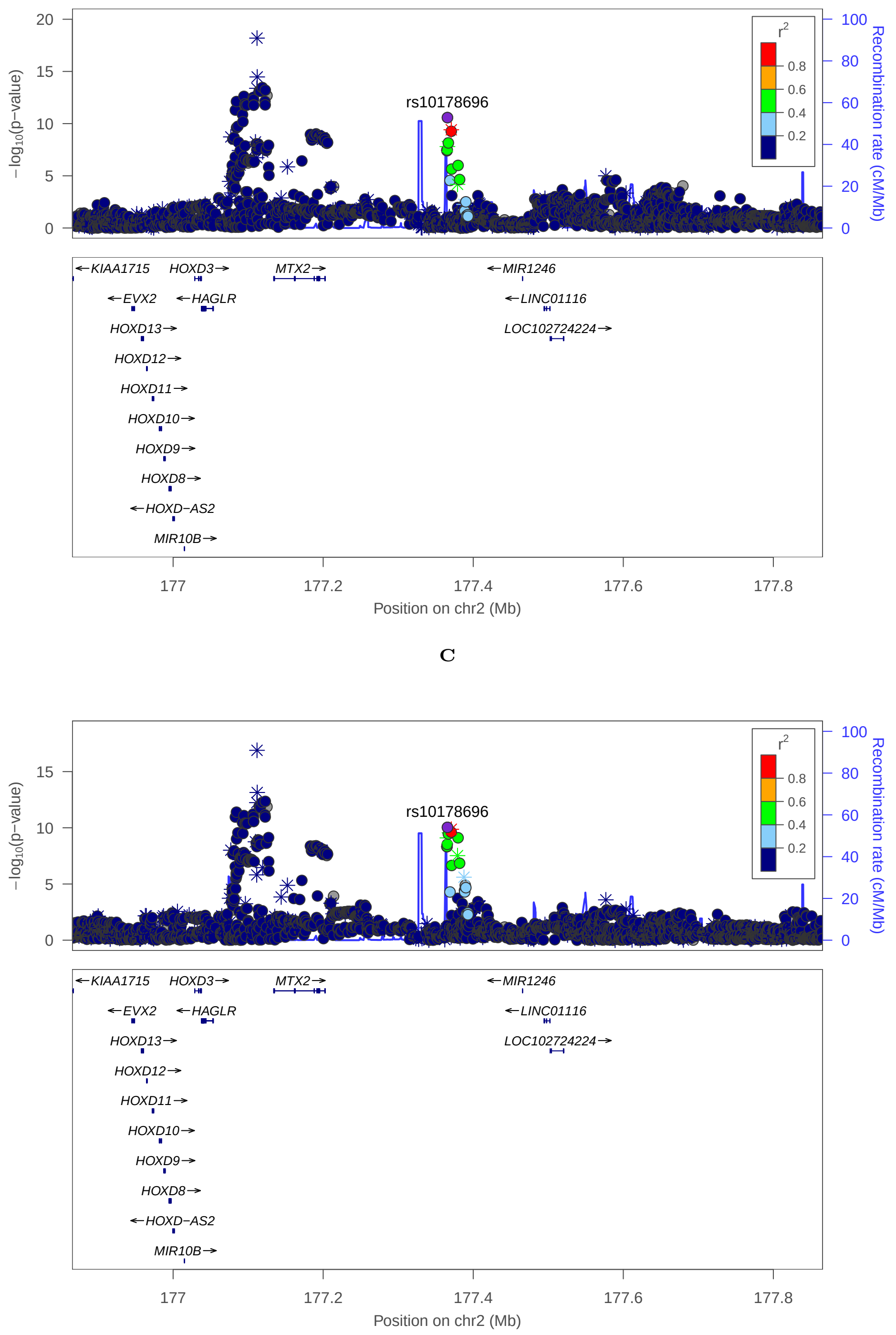

C

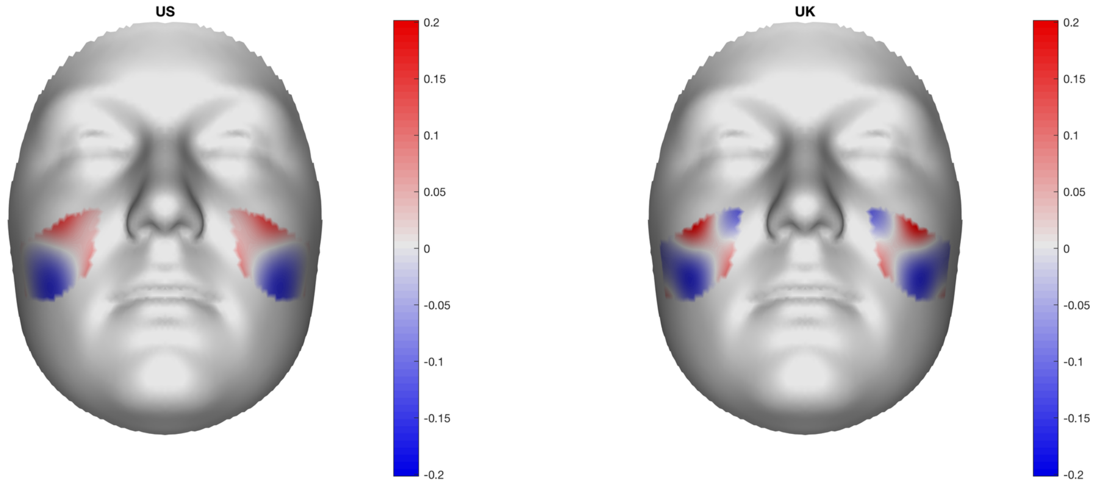

B

### Multi-peak locus 9: rs58022575

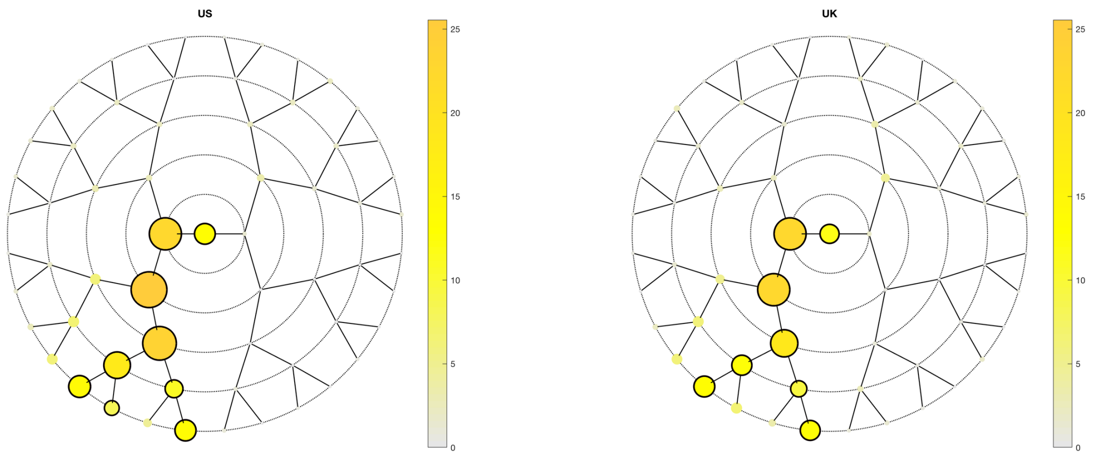

**A**

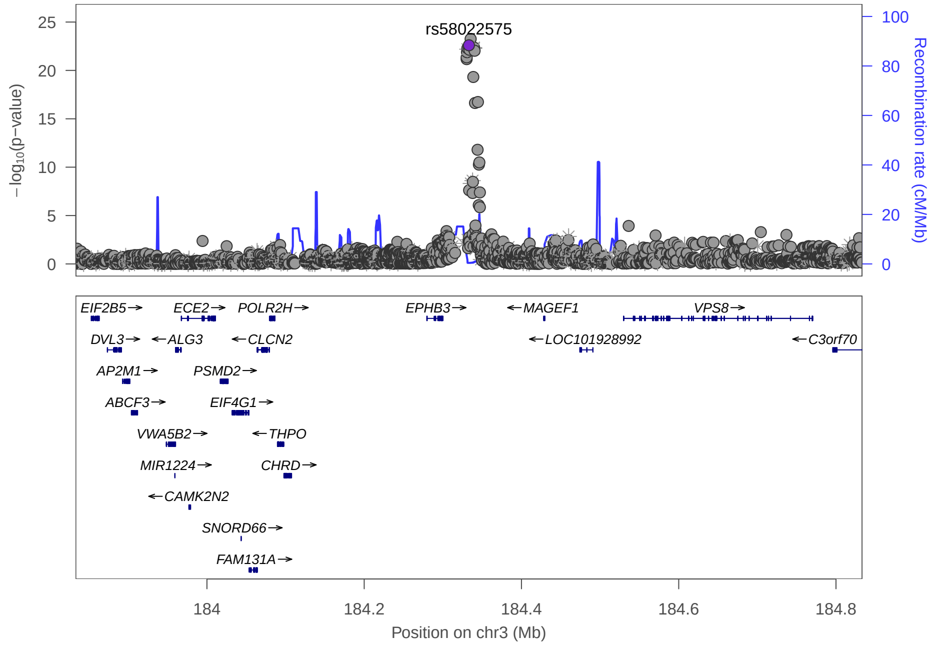

**C**

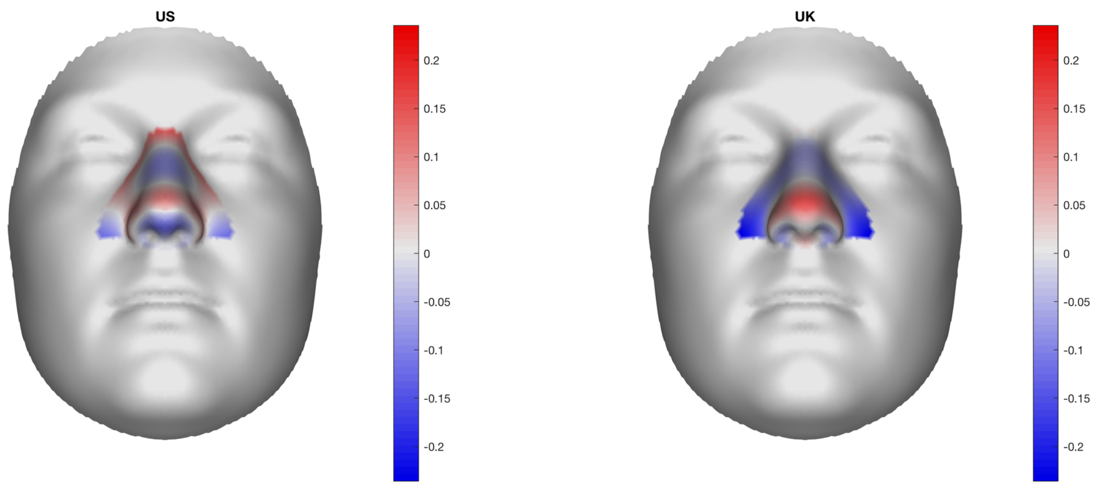

**B**

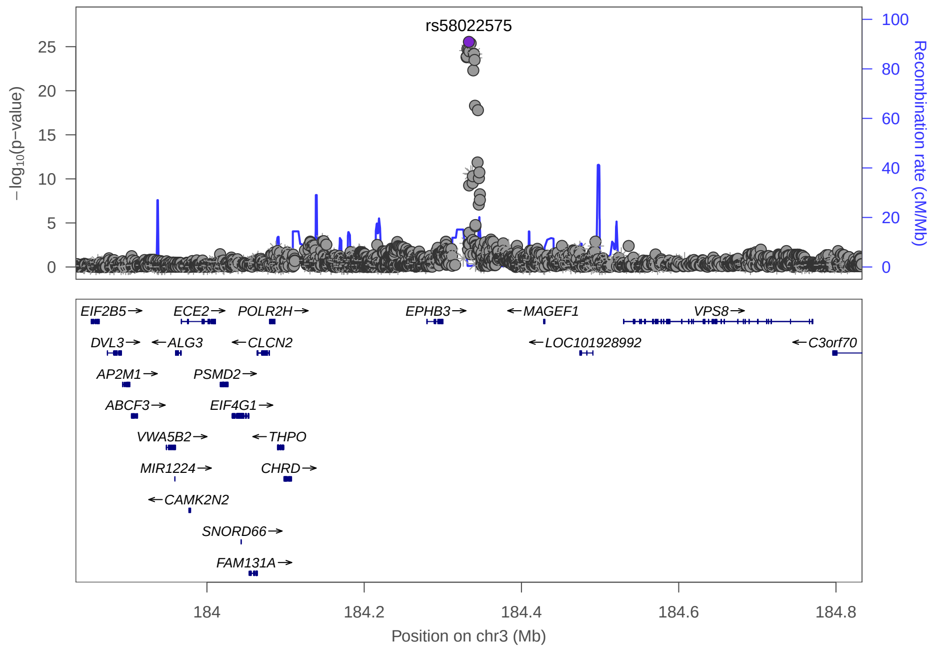

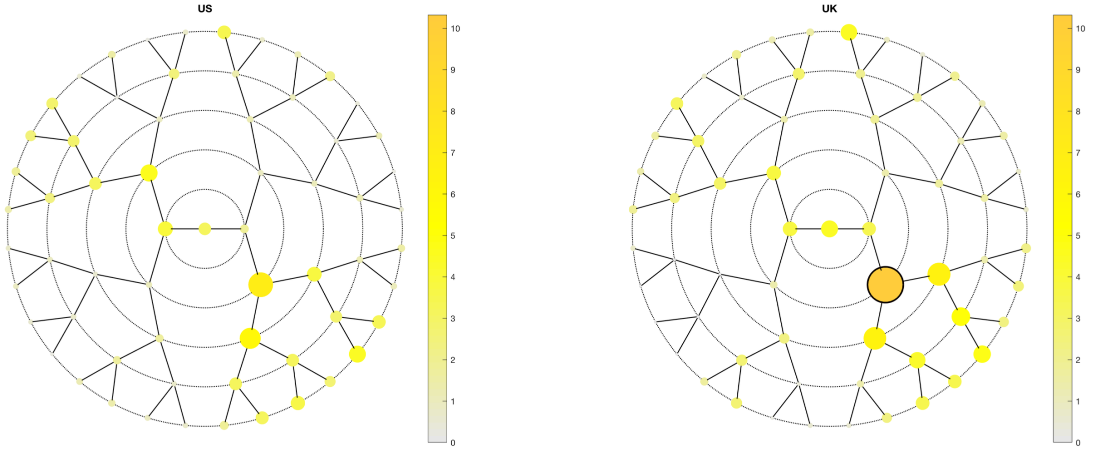
US
UK
10
9
8
7
6
5
4
3
2
1
0

A

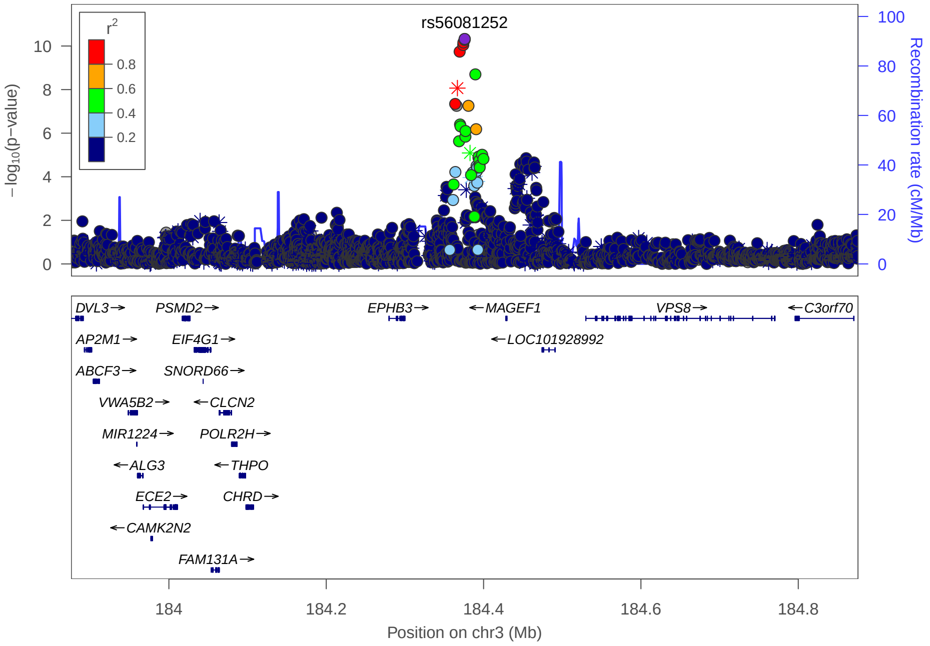
rs56081252
r²
0.8
0.6
0.4
0.2
−log₁₀(p−value)
Recombination rate (cM/Mb)
DVL3→
PSMD2→
EPHB3→
←MAGEF1
VPS8→
←C3orf70
AP2M1→
EIF4G1→
←LOC101928992
ABCF3→
SNORD66→
VWA5B2→
←CLCN2
MIR1224→
POLR2H→
←ALG3
←THPO
ECE2→
CHRD→
←CAMK2N2
FAM131A→
184
184.2
184.4
184.6
184.8
Position on chr3 (Mb)

C

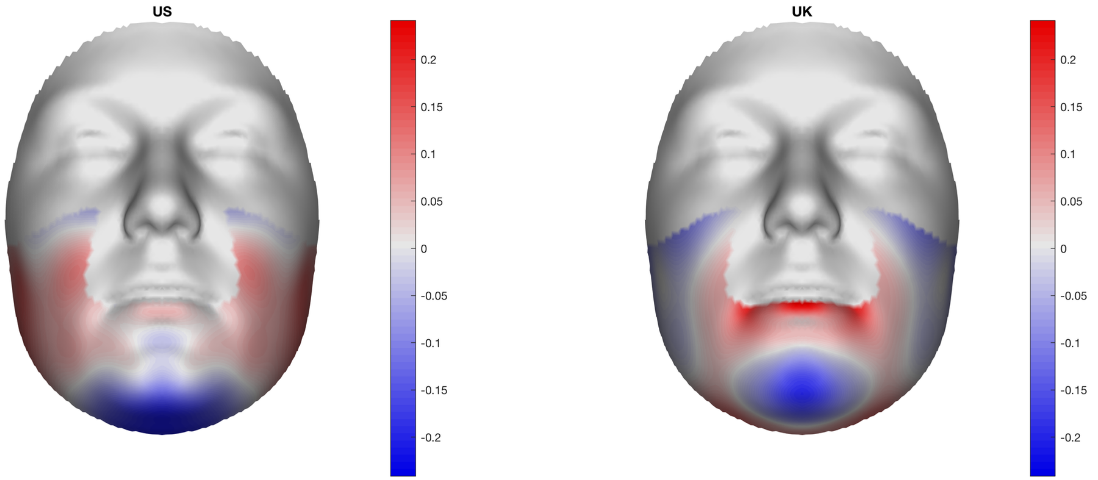
US
UK
0.2
0.15
0.1
0.05
0
-0.05
-0.1
-0.15
-0.2

B

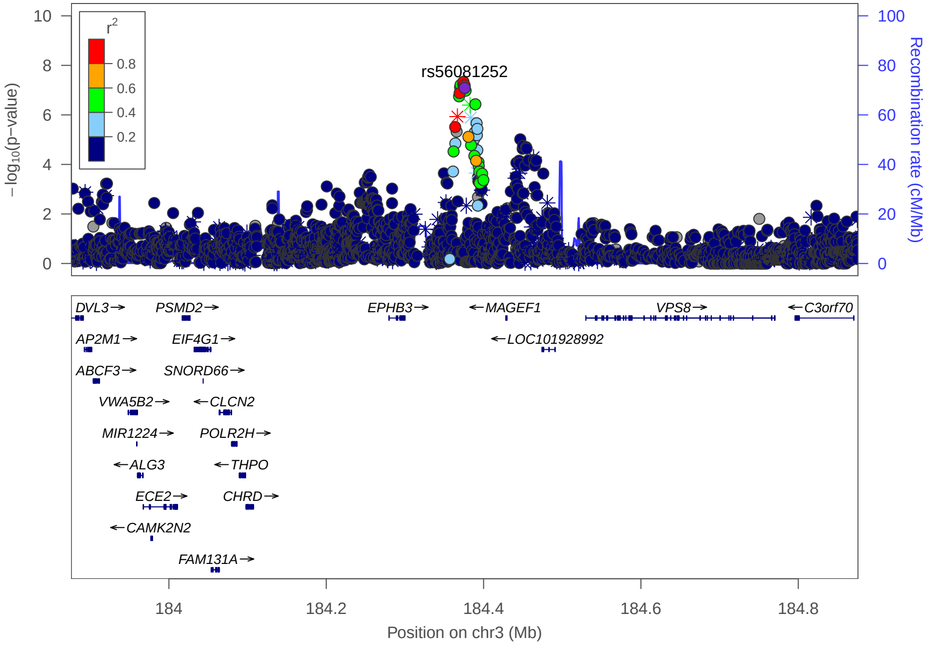
rs56081252
r²
0.8
0.6
0.4
0.2
−log₁₀(p−value)
Recombination rate (cM/Mb)
DVL3→
PSMD2→
EPHB3→
←MAGEF1
VPS8→
←C3orf70
AP2M1→
EIF4G1→
←LOC101928992
ABCF3→
SNORD66→
VWA5B2→
←CLCN2
MIR1224→
POLR2H→
←ALG3
←THPO
ECE2→
CHRD→
←CAMK2N2
FAM131A→
184
184.2
184.4
184.6
184.8
Position on chr3 (Mb)

### Multi-peak locus 10: rs3910659

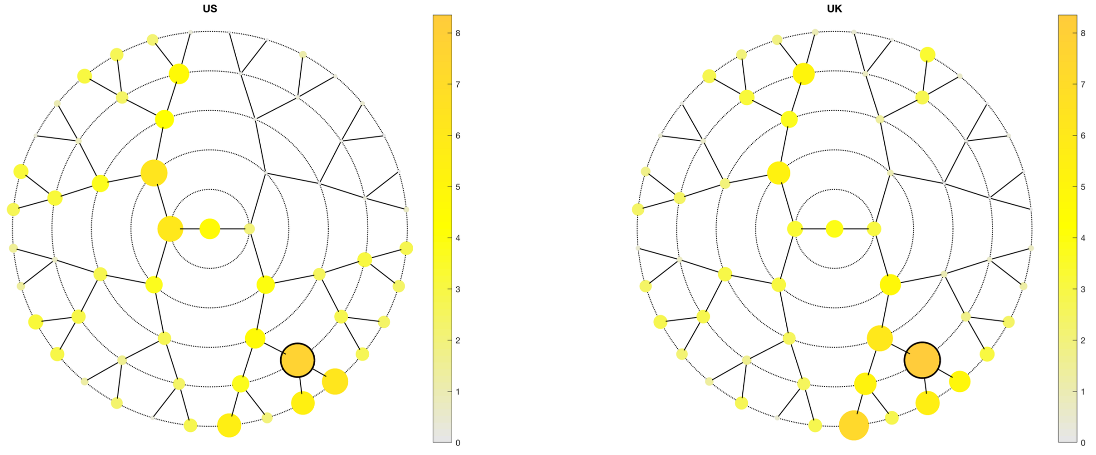

**A**

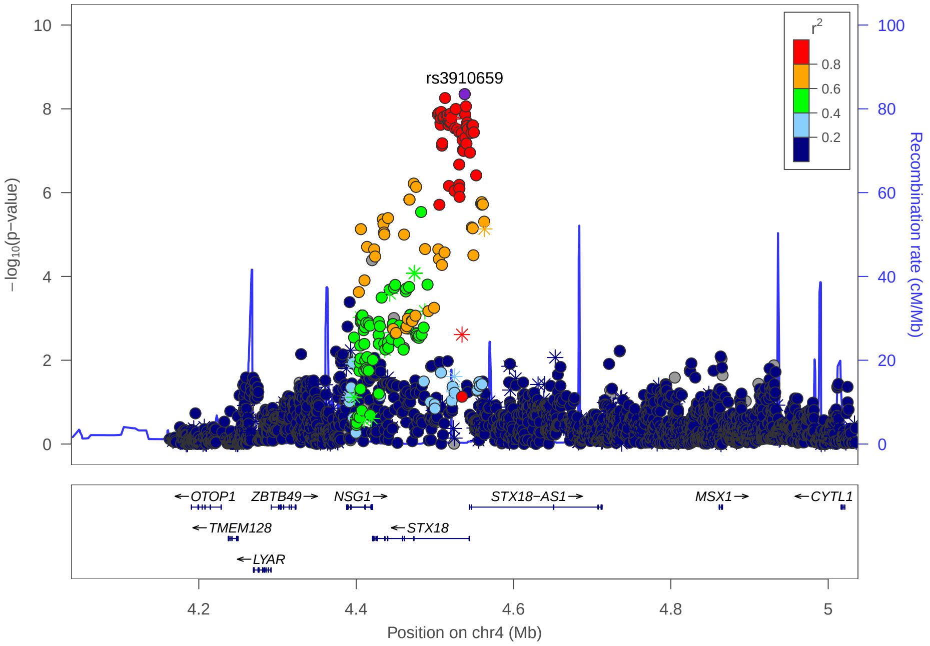

**C**

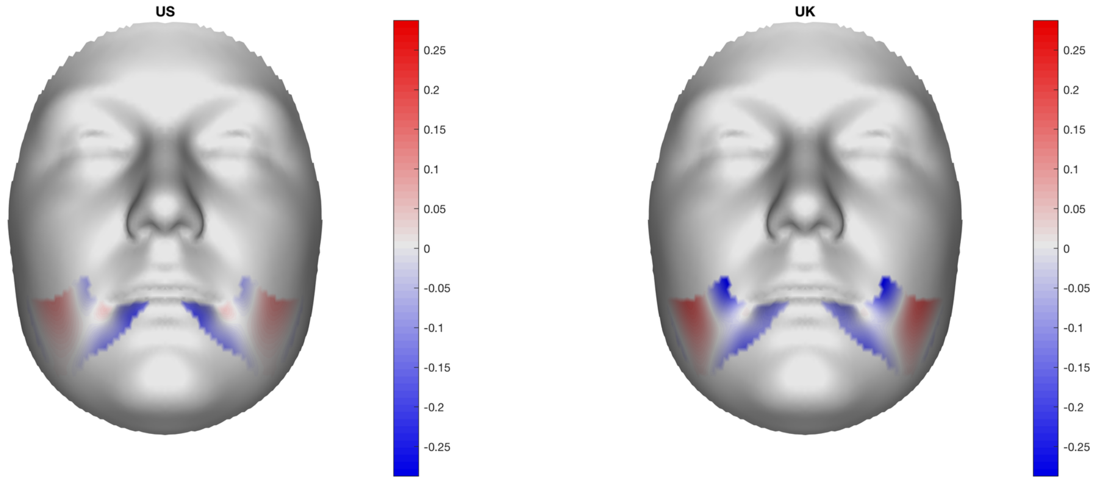

**B**

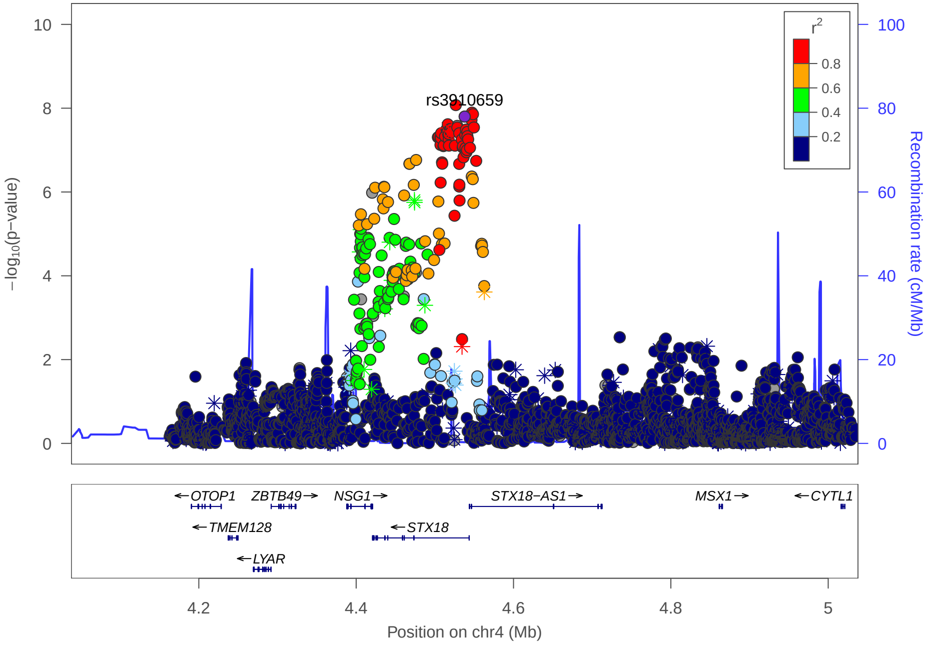

### Multi-peak locus 10: rs13117653

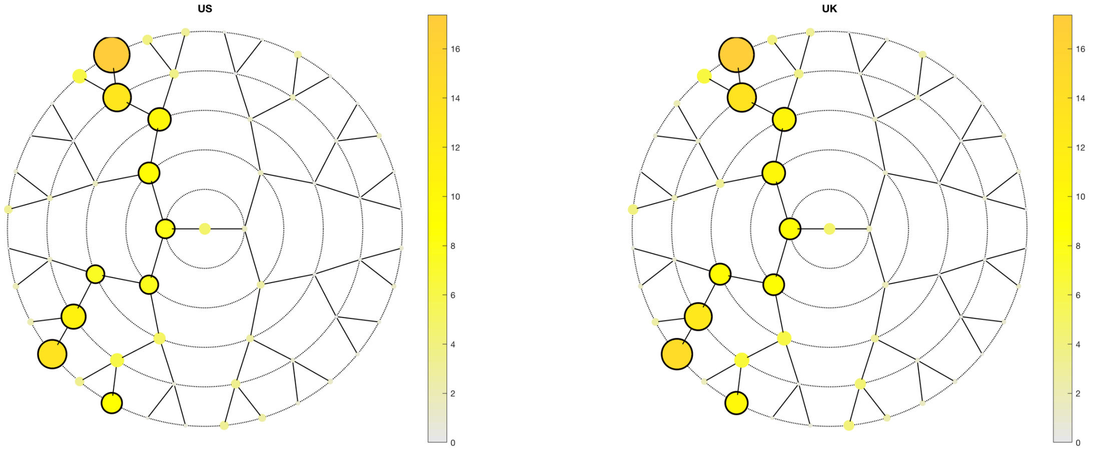

**A**

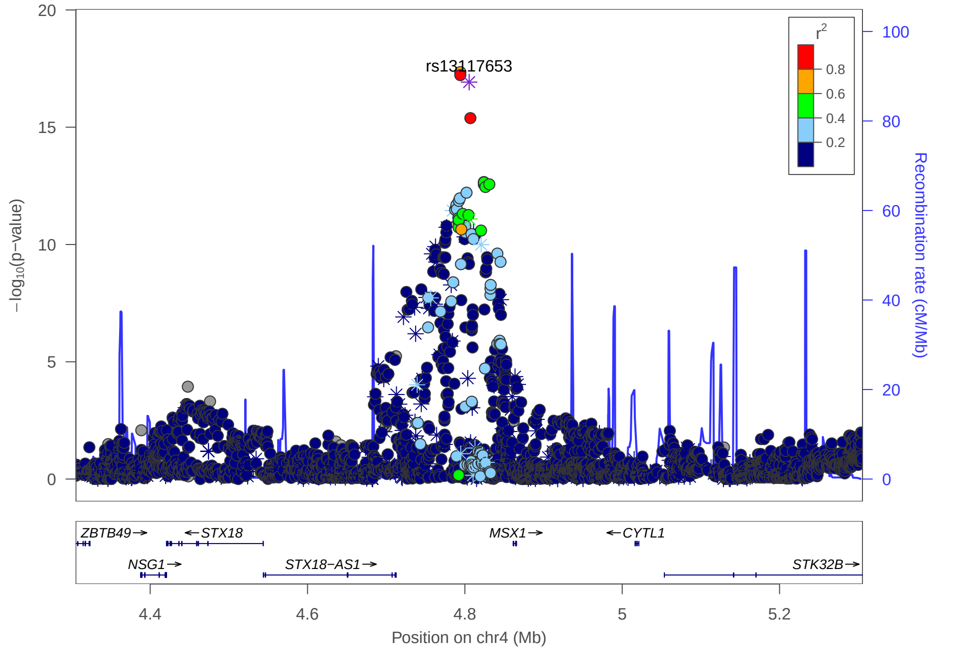

**C**

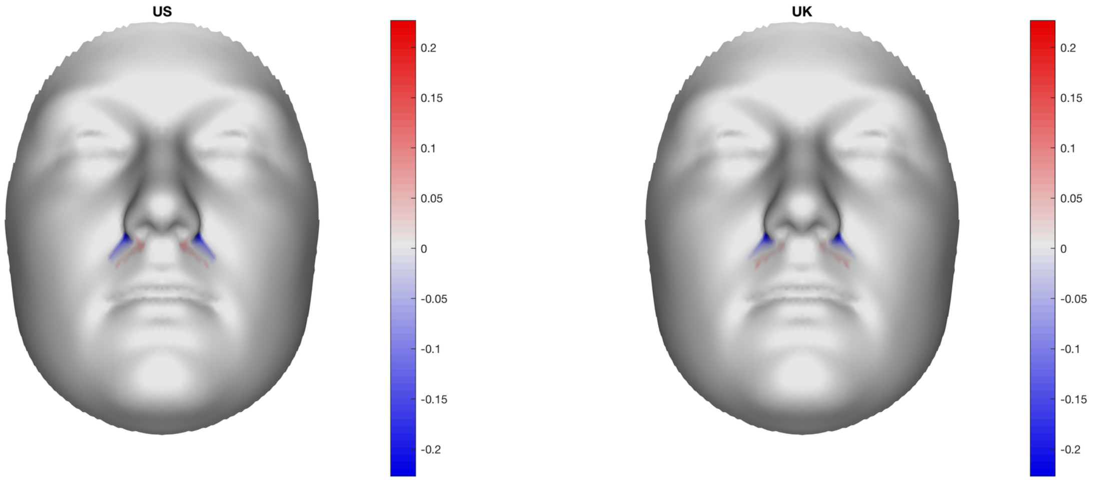

**B**

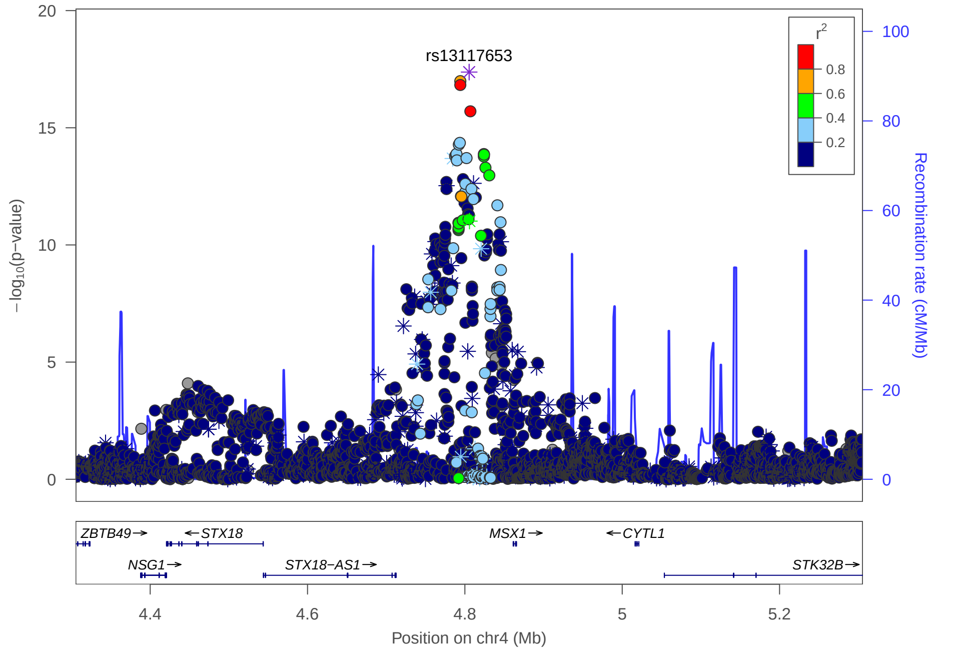

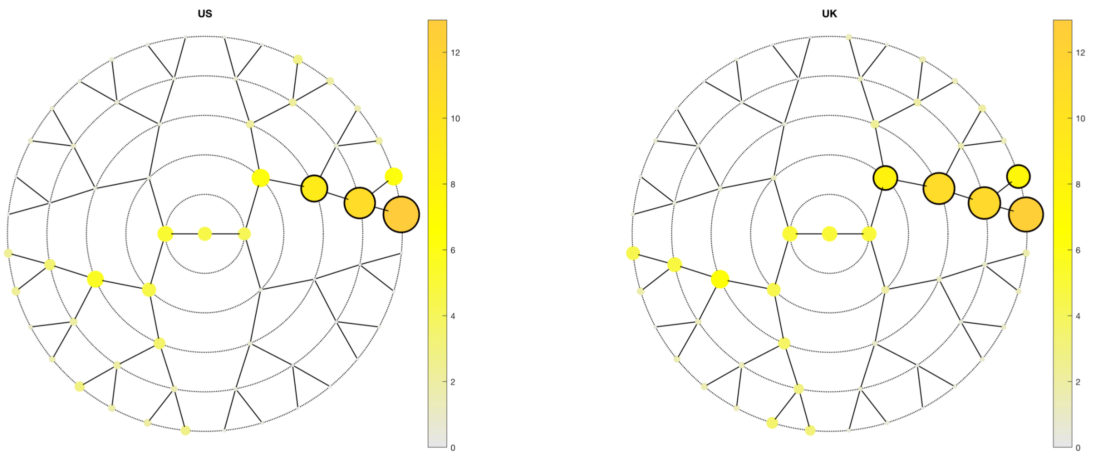
US
UK

A

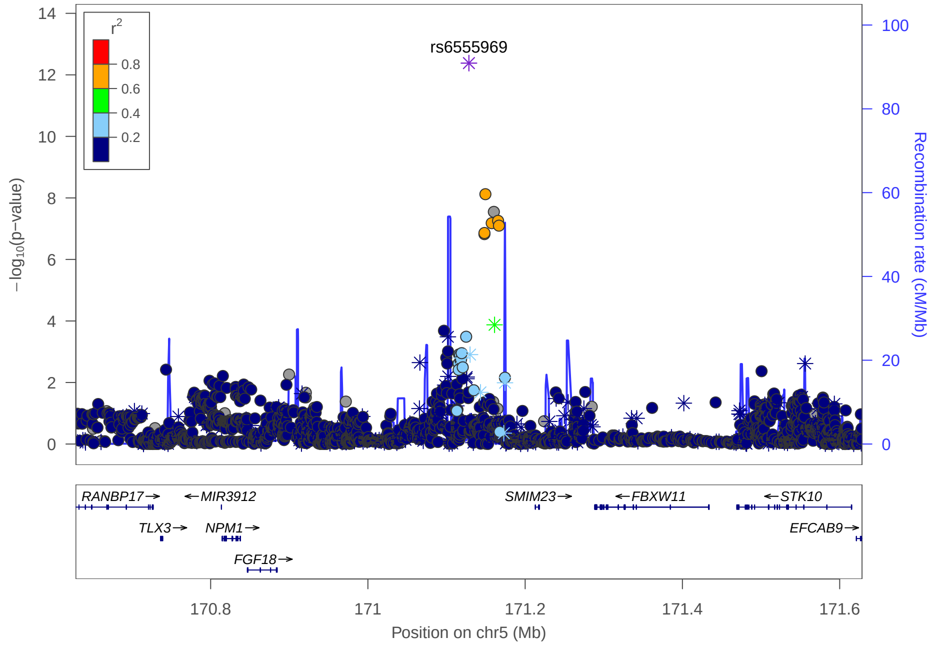
rs6555969
Position on chr5 (Mb)
Recombination rate (cM/Mb)
RANBP17
MIR3912
SMIM23
FBXW11
STK10
TLX3
NPM1
EFCAB9
FGF18

C

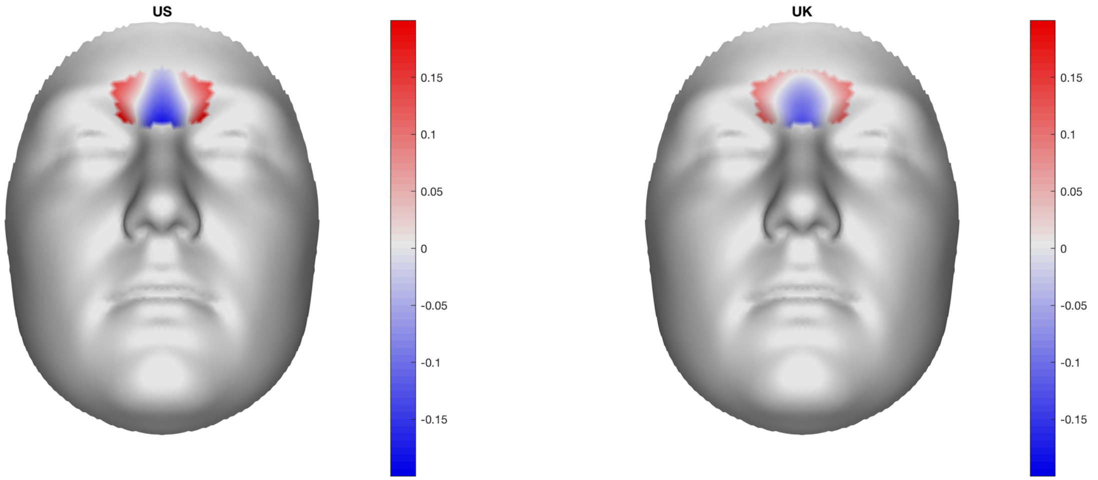
US
UK

B

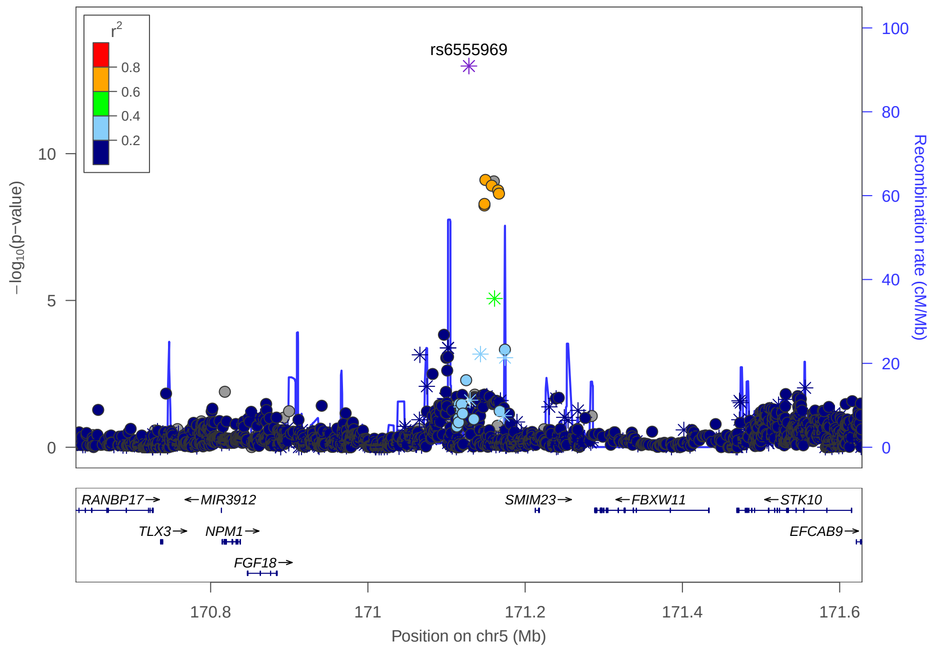
rs6555969
Position on chr5 (Mb)
Recombination rate (cM/Mb)
RANBP17
MIR3912
SMIM23
FBXW11
STK10
TLX3
NPM1
EFCAB9
FGF18

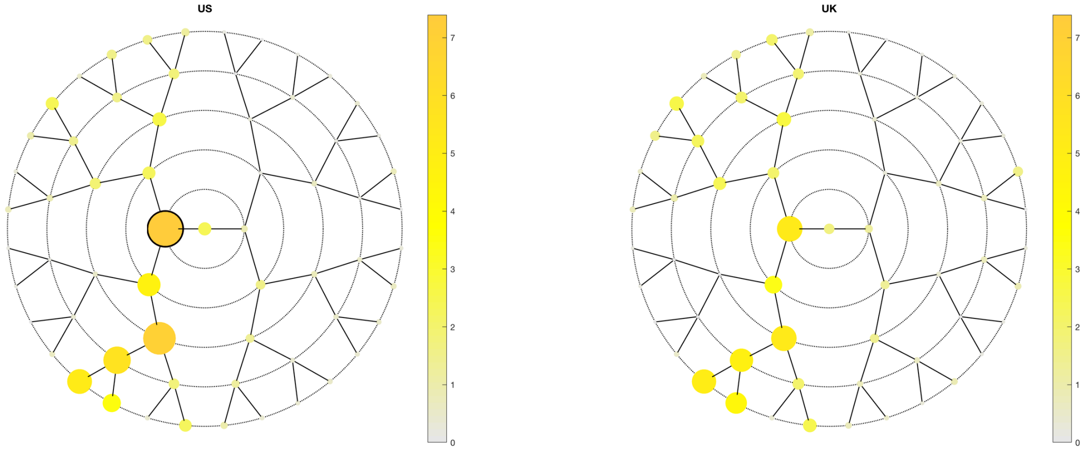
US
UK
0
1
2
3
4
5
6
7

A

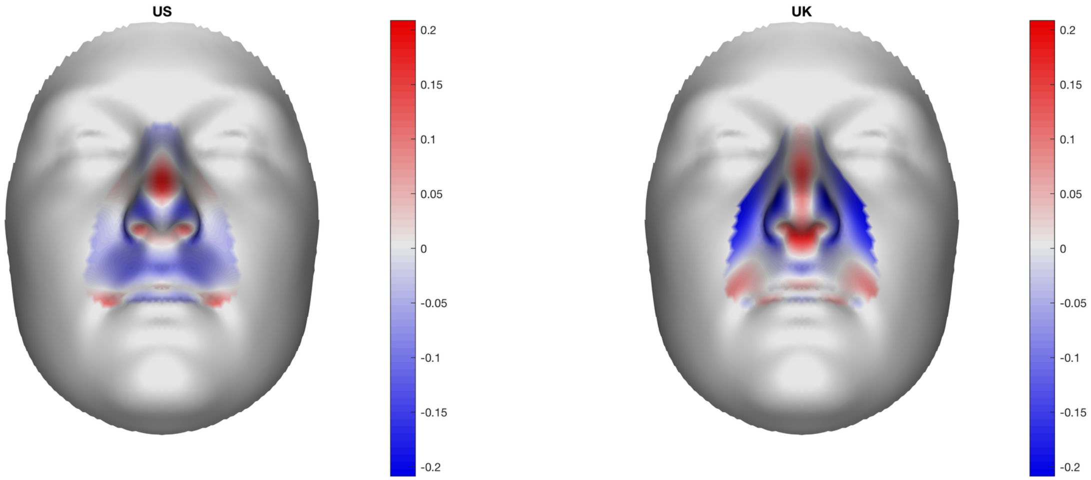
US
UK
0.2
0.15
0.1
0.05
0
-0.05
-0.1
-0.15
-0.2

B

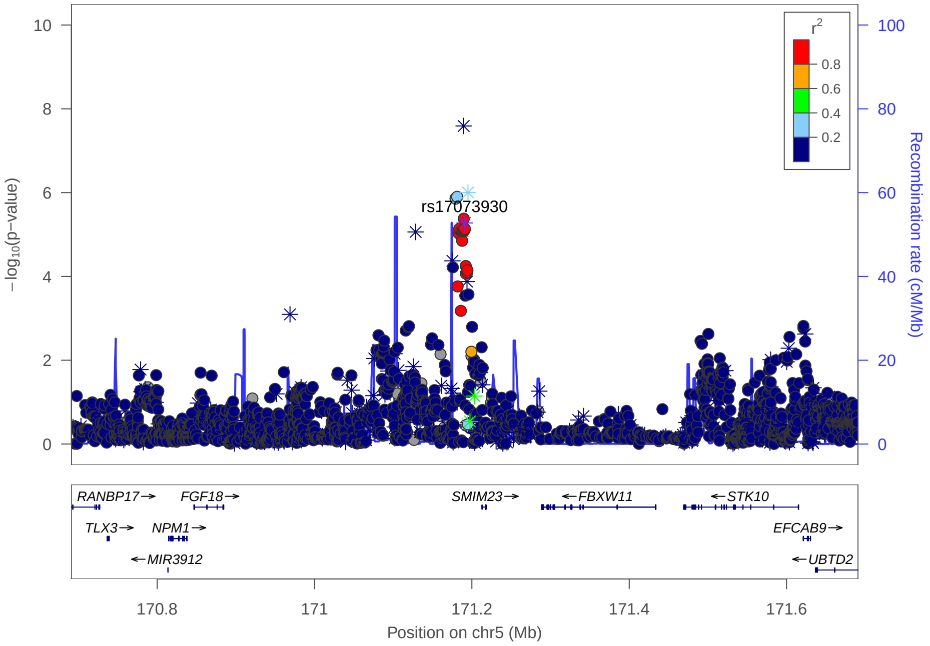
rs17073930
r2
0.8
0.6
0.4
0.2
−log10(p−value)
Recombination rate (cM/Mb)
RANBP17
FGF18
SMIM23
FBXW11
STK10
TLX3
NPM1
EFCAB9
MIR3912
UBTD2
170.8
171
171.2
171.4
171.6
Position on chr5 (Mb)

C

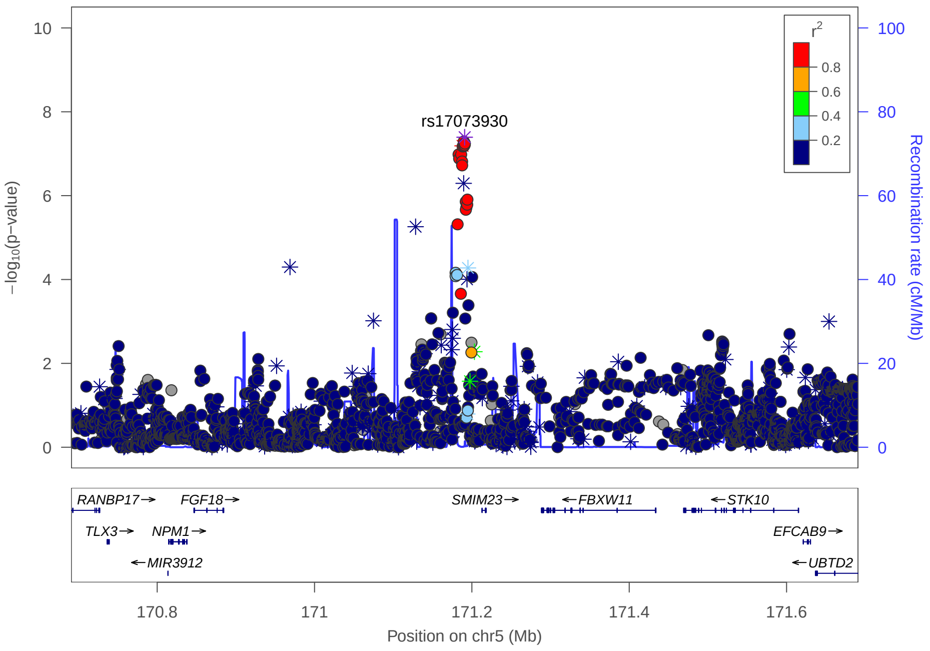
rs17073930
r2
0.8
0.6
0.4
0.2
−log10(p−value)
Recombination rate (cM/Mb)
RANBP17
FGF18
SMIM23
FBXW11
STK10
TLX3
NPM1
EFCAB9
MIR3912
UBTD2
170.8
171
171.2
171.4
171.6
Position on chr5 (Mb)

### Multi-peak locus 12: rs2245221

**CHR8** - See final page for captions

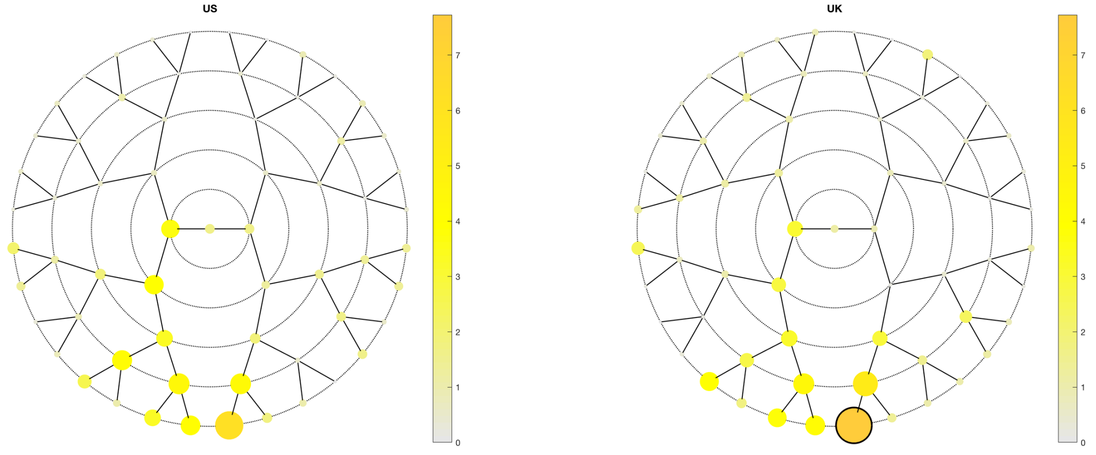

A

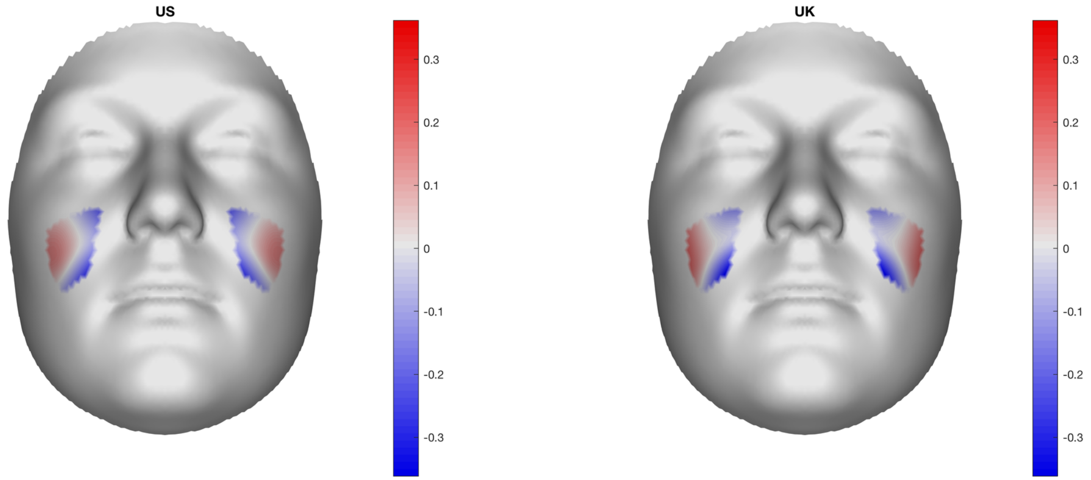

B

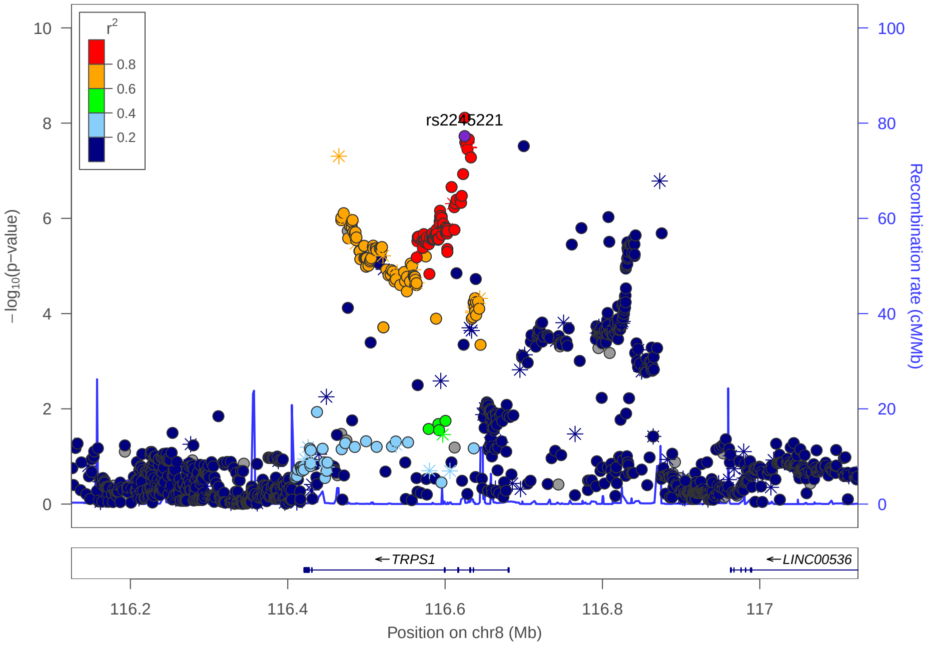

C

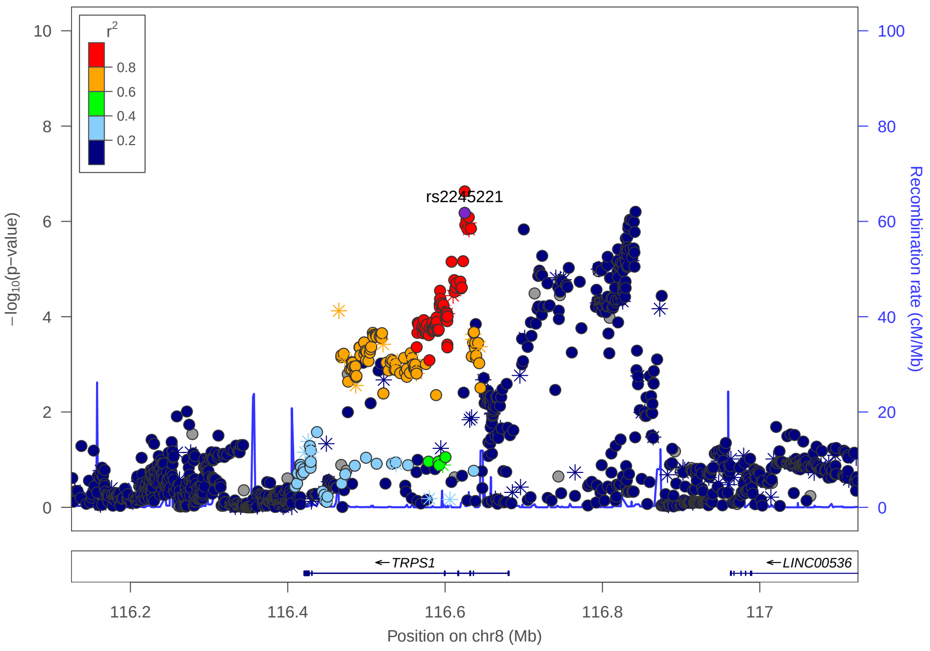

### Multi-peak locus 12: rs11337200

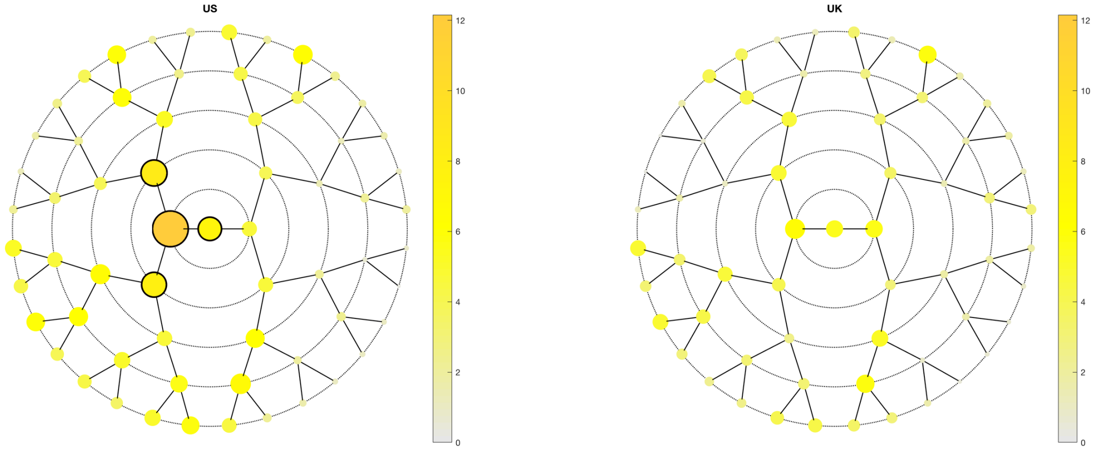

**A**

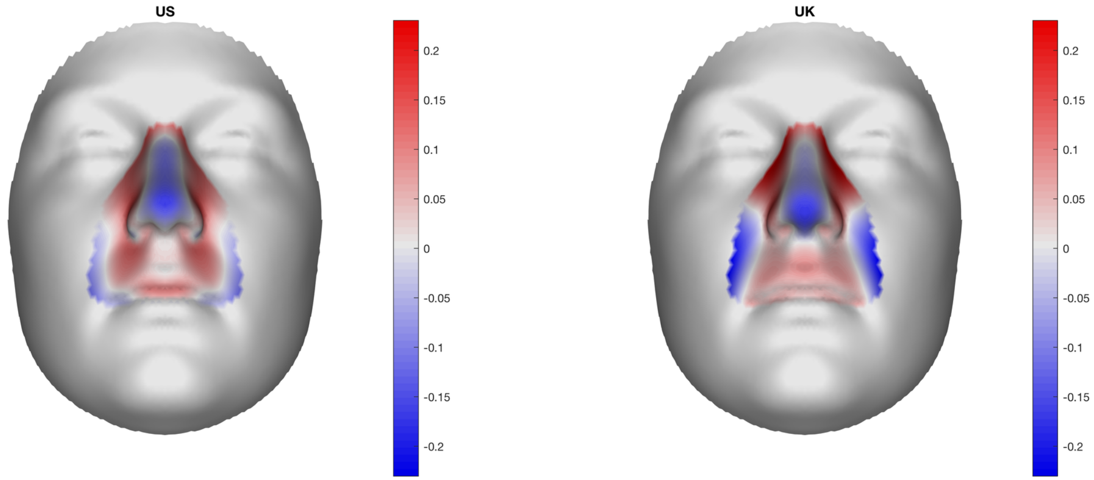

**B**

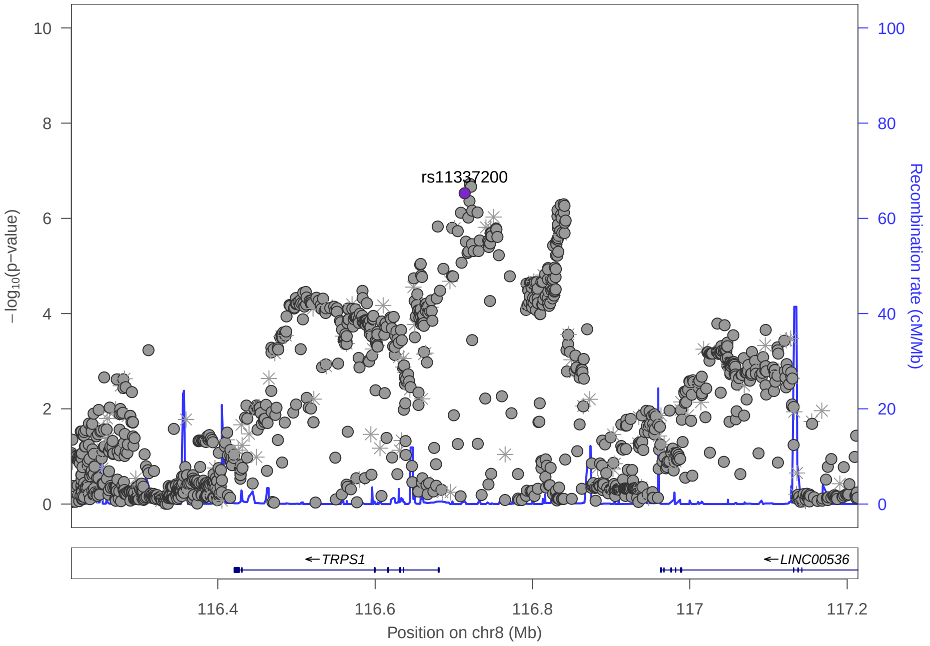

**C**

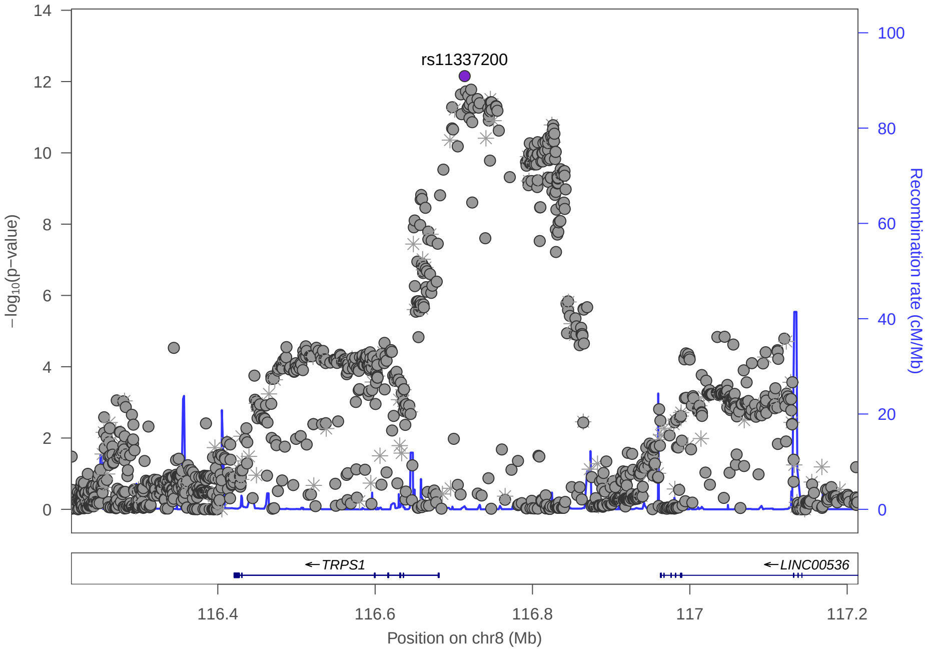

### Multi-peak locus 13: rs7843236

**CHR8** - See final page for captions

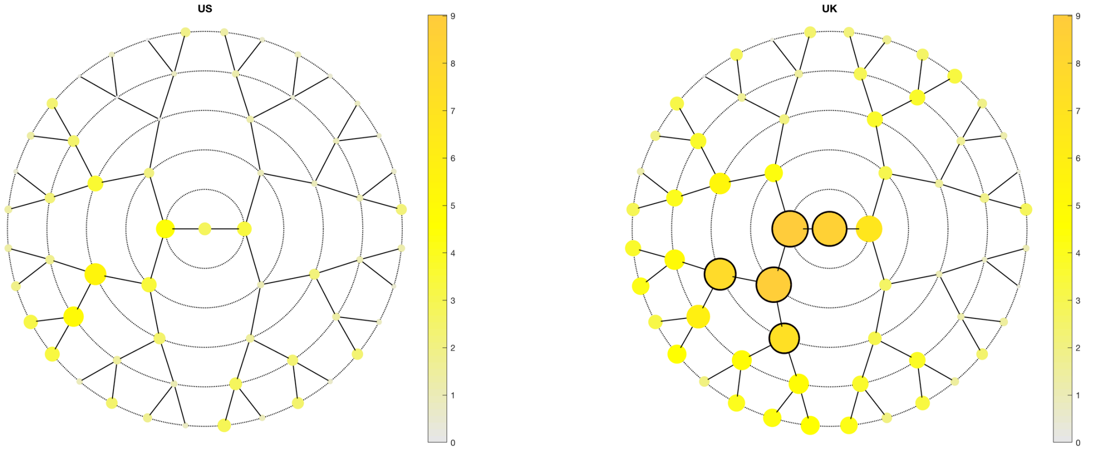

**A**

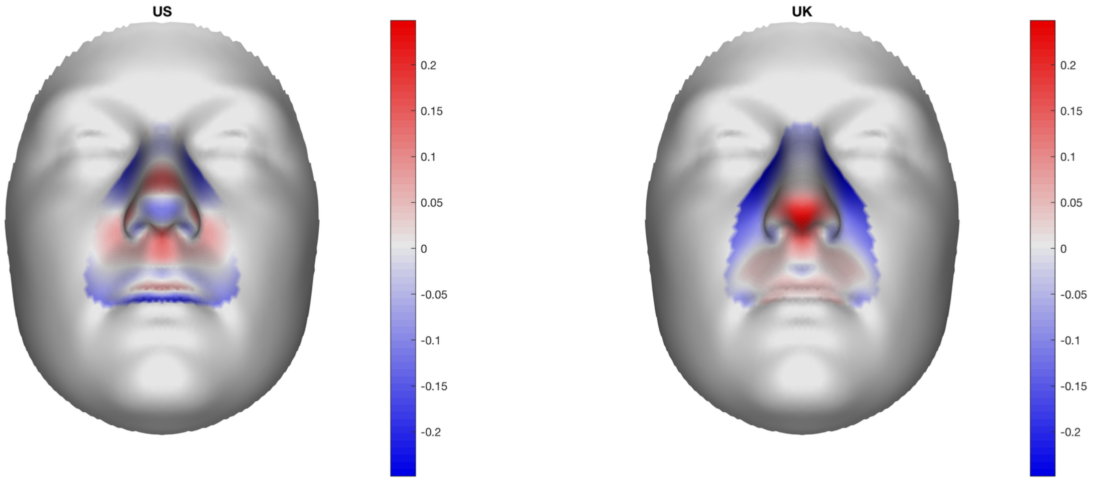

**B**

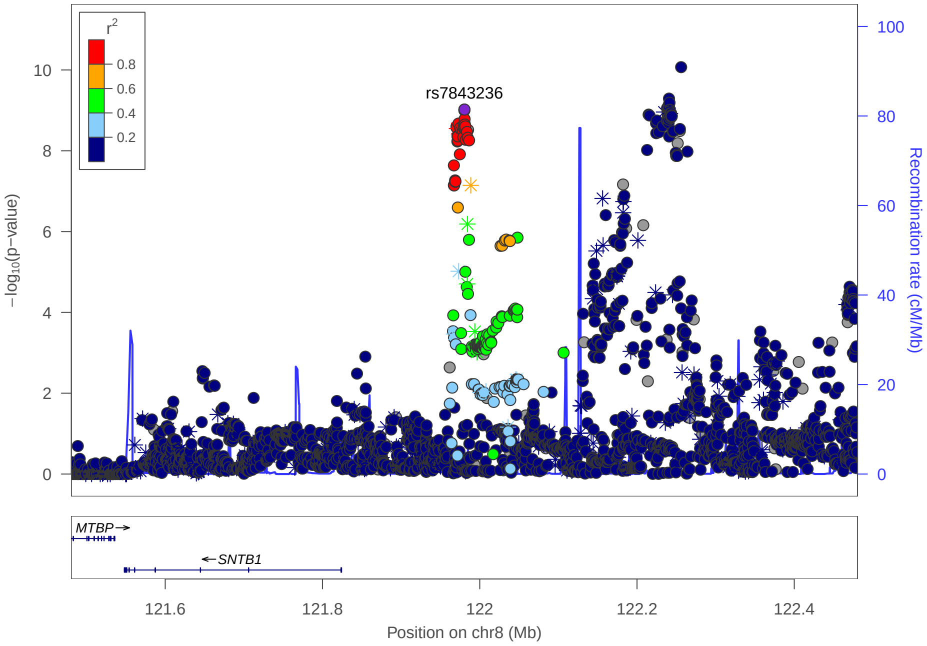

**C**

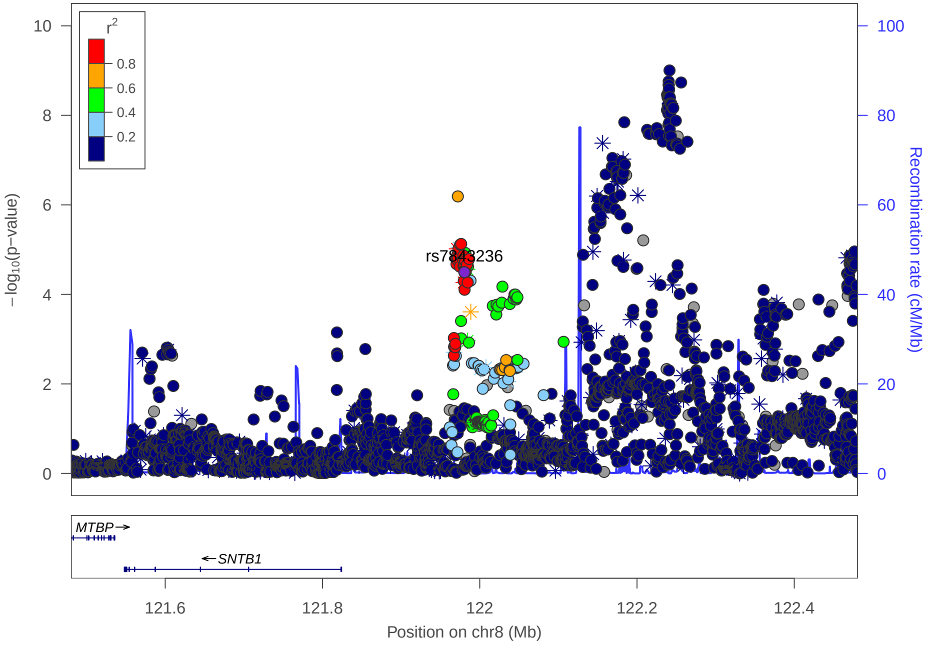

### Multi-peak locus 13: rs2581548

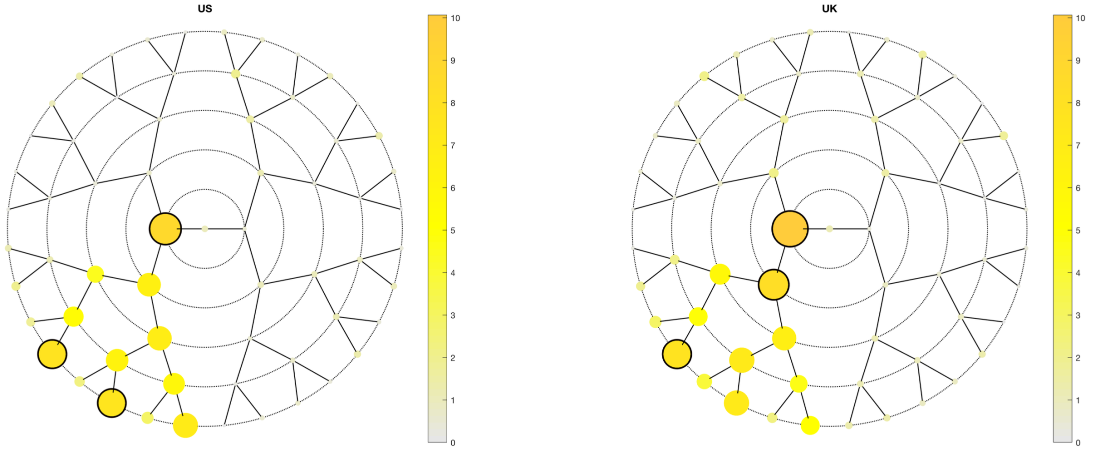

**A**

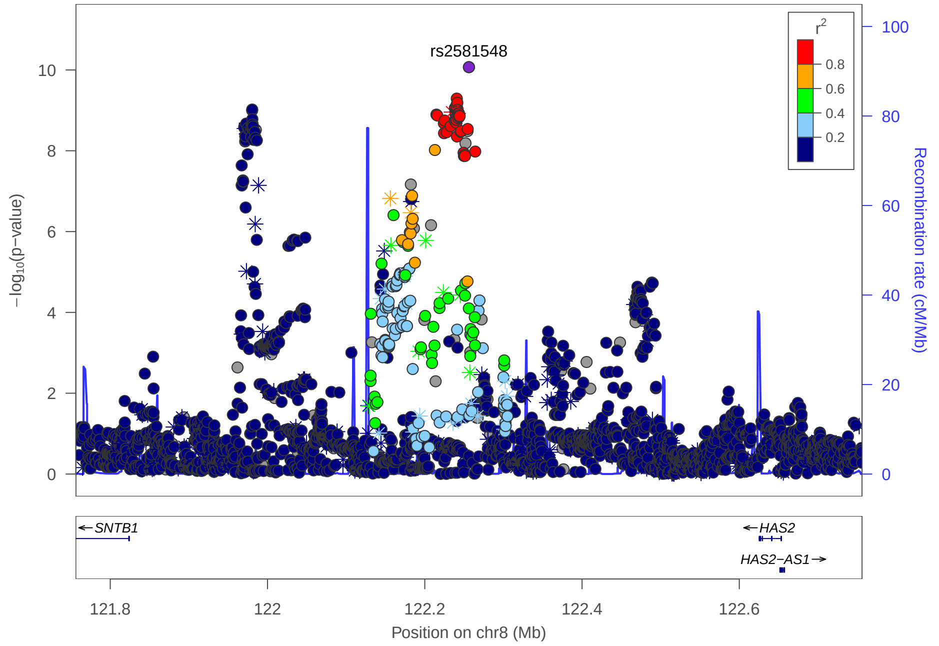

**C**

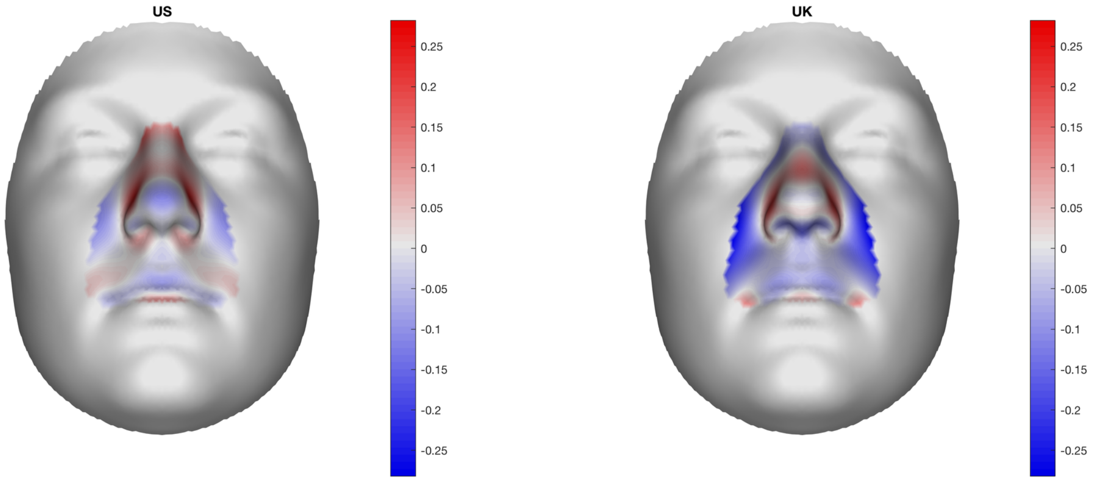

**B**

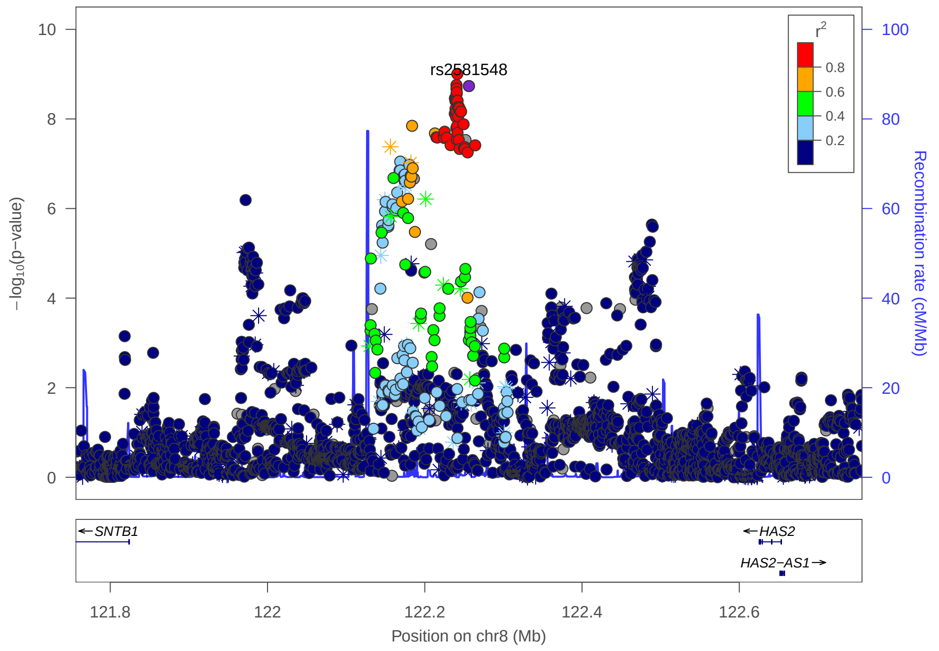

### Multi-peak locus 14: rs242983

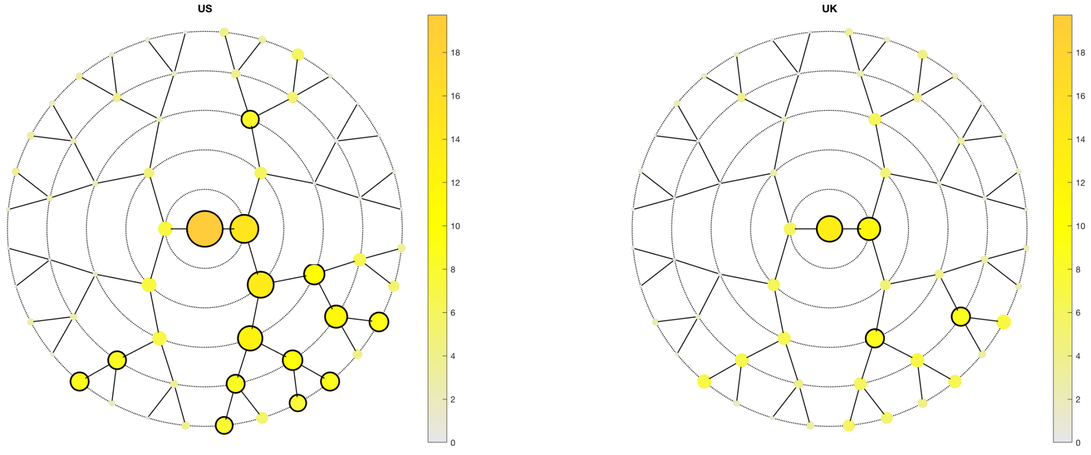

**A**

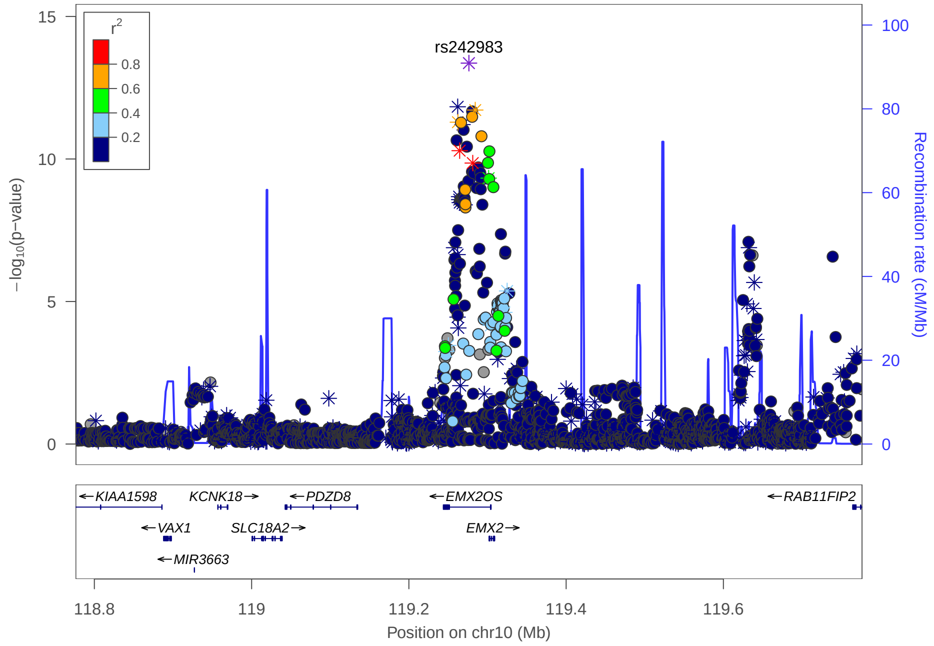

**C**

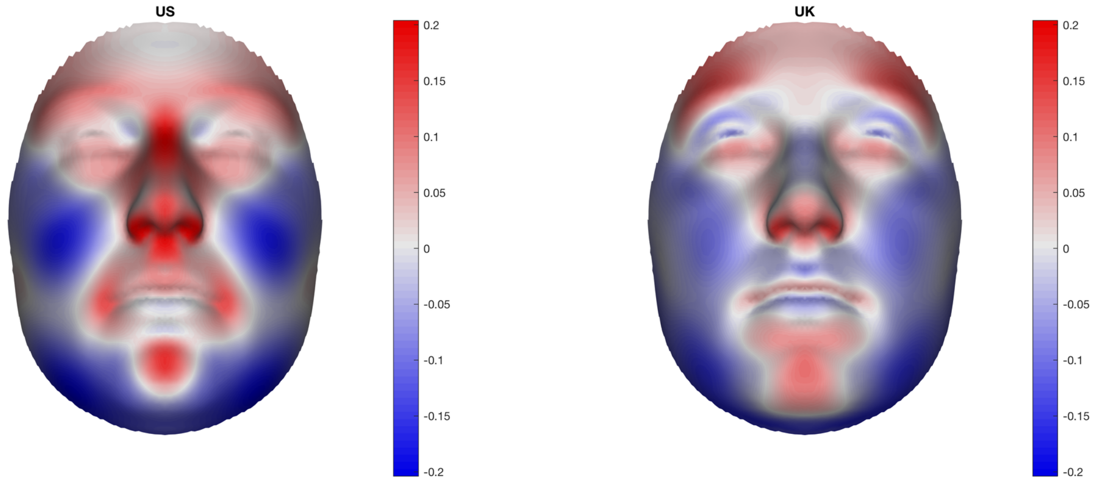

**B**

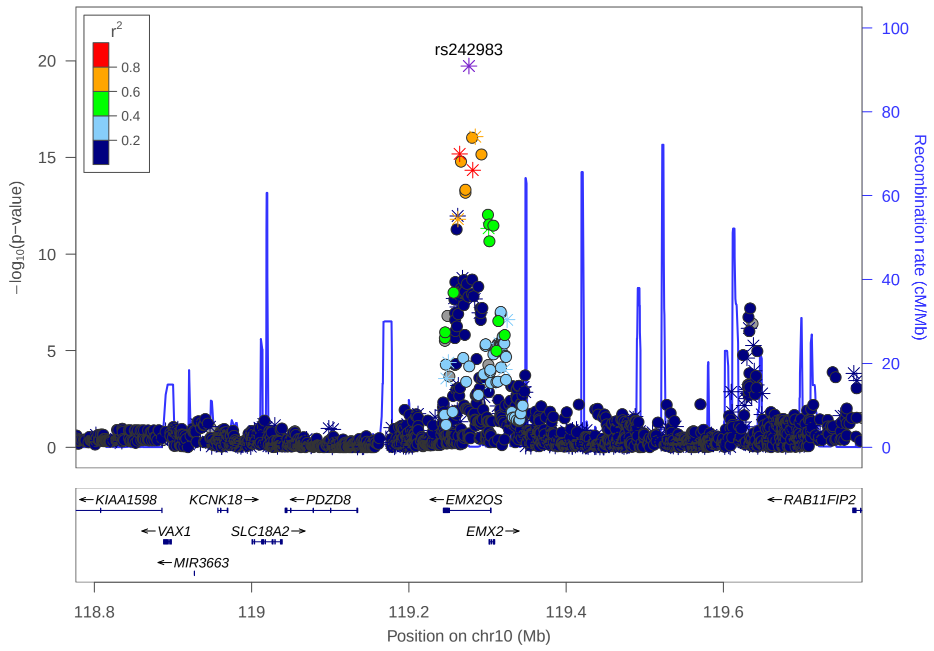

Multi-peak locus 14: rs34988394

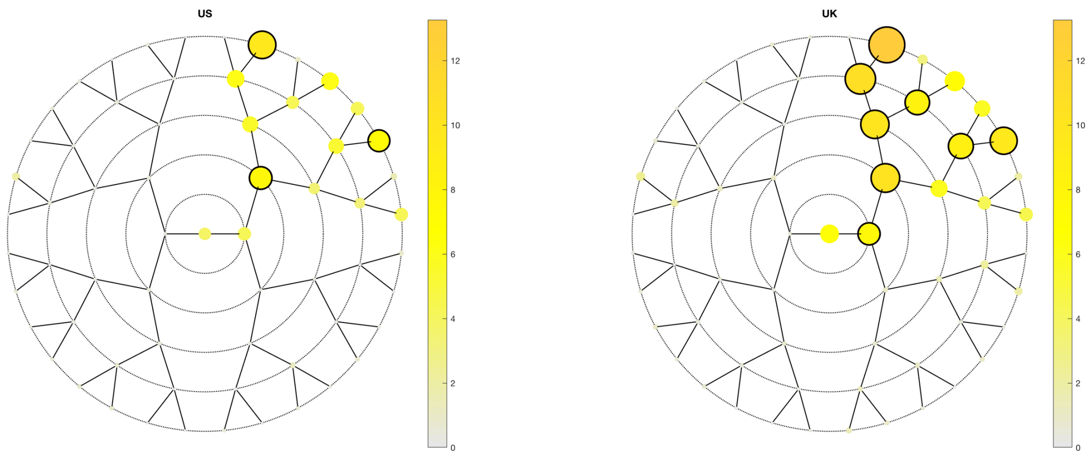

A

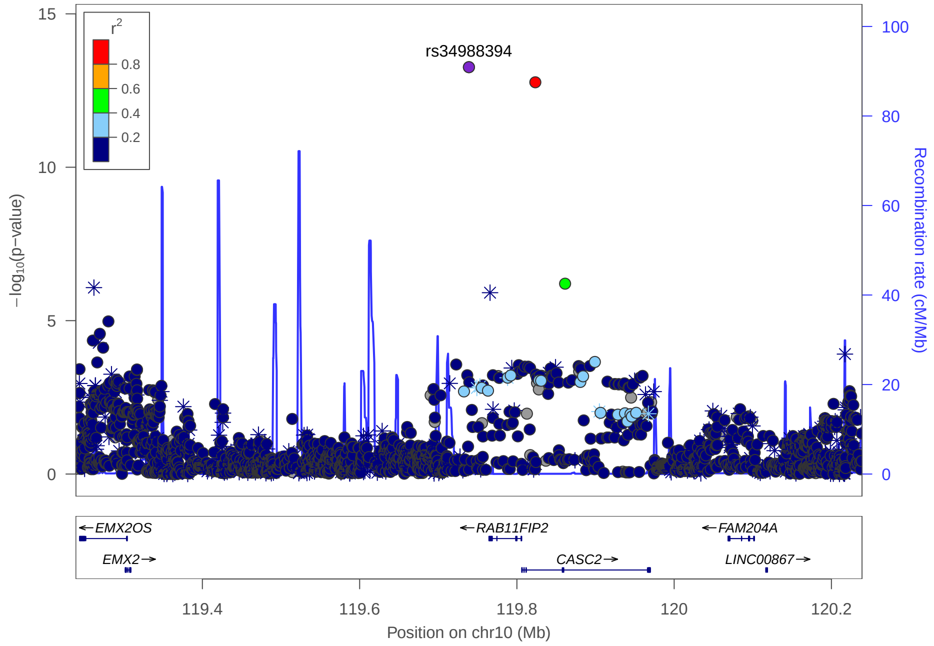

C

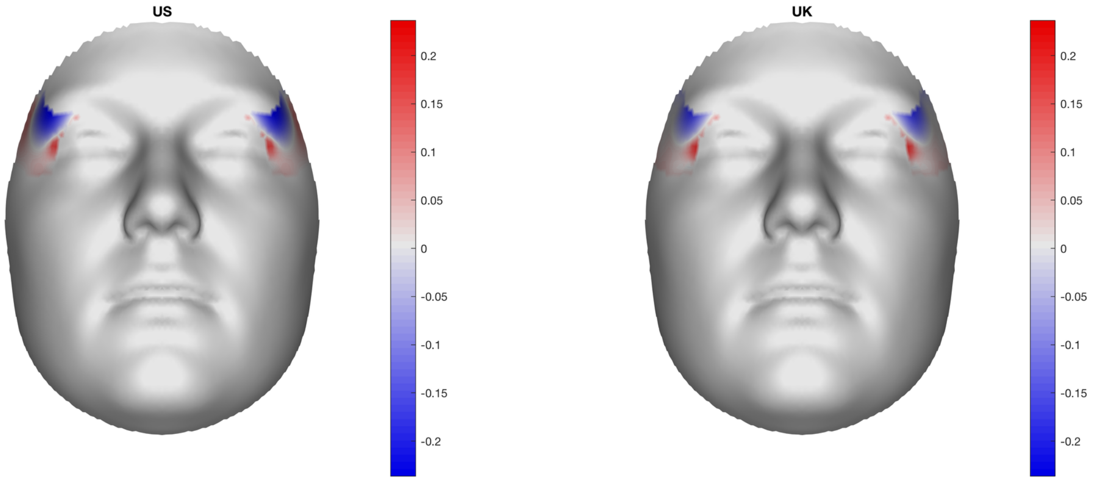

B

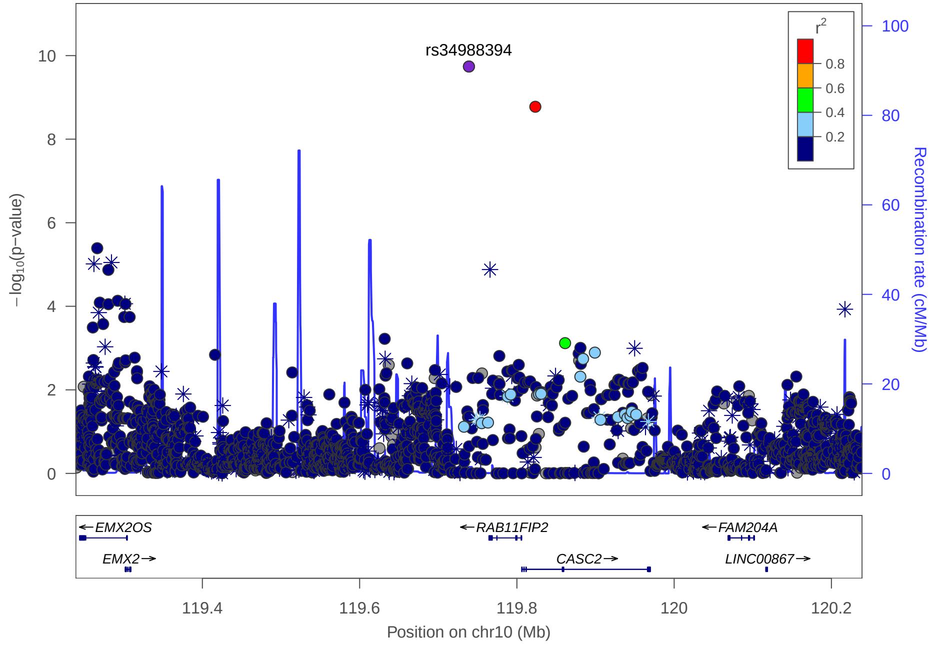

#### **Multi-peak locus 15: rs7121535**

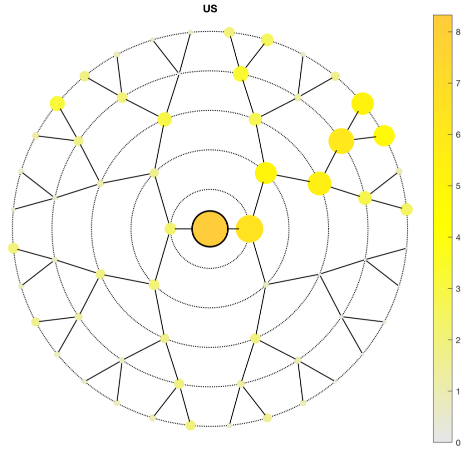

**A**

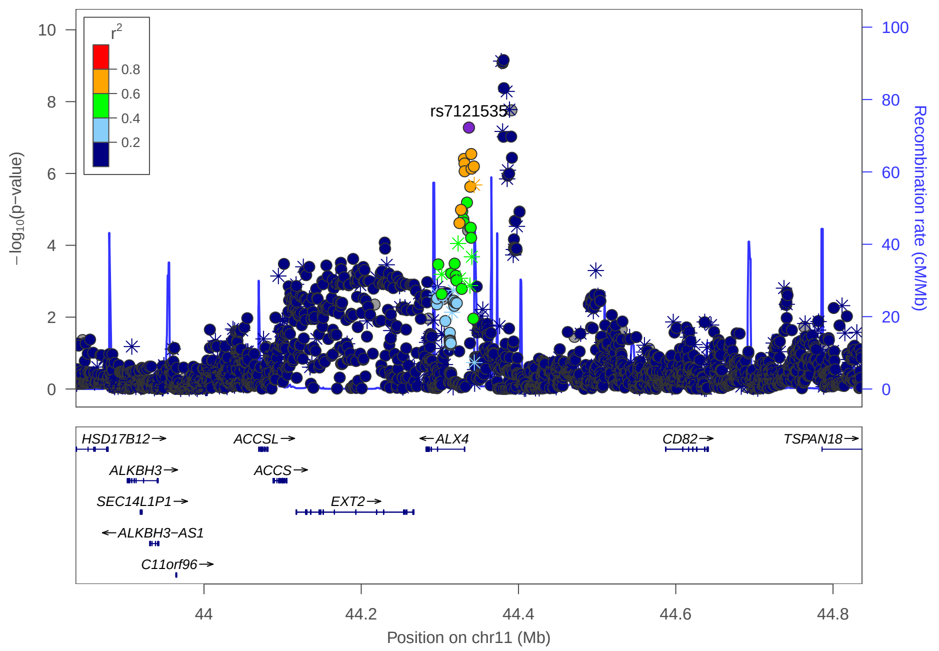

**C**

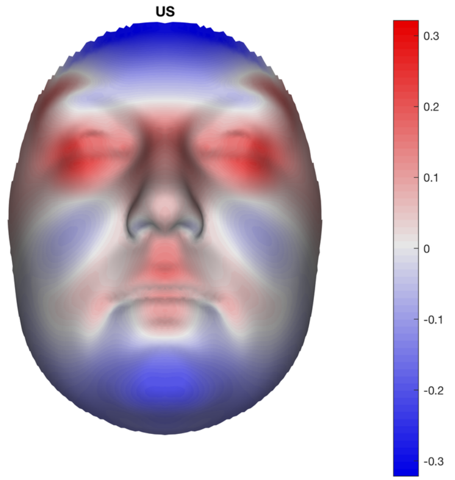

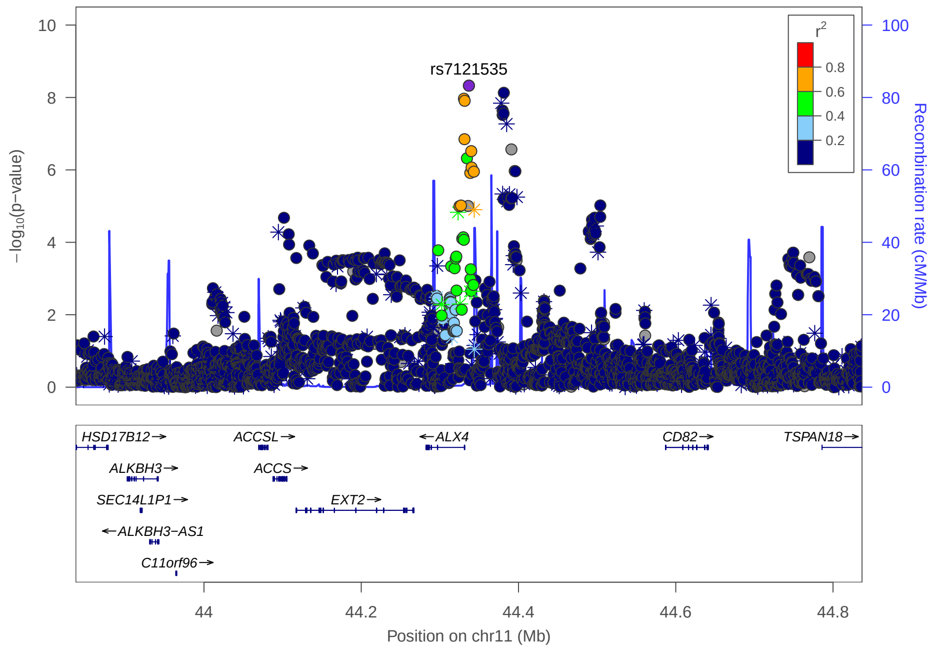

**B**

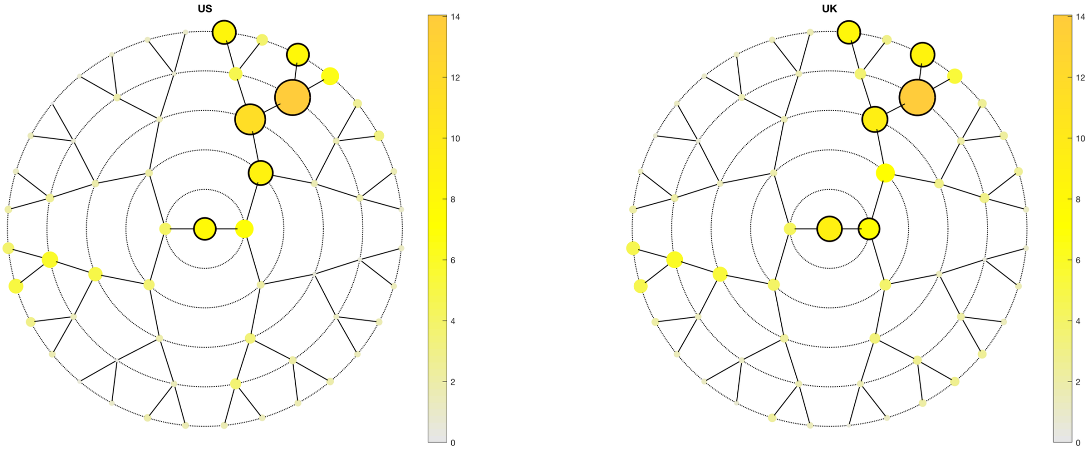
US
UK
0
2
4
6
8
10
12
14

**A**

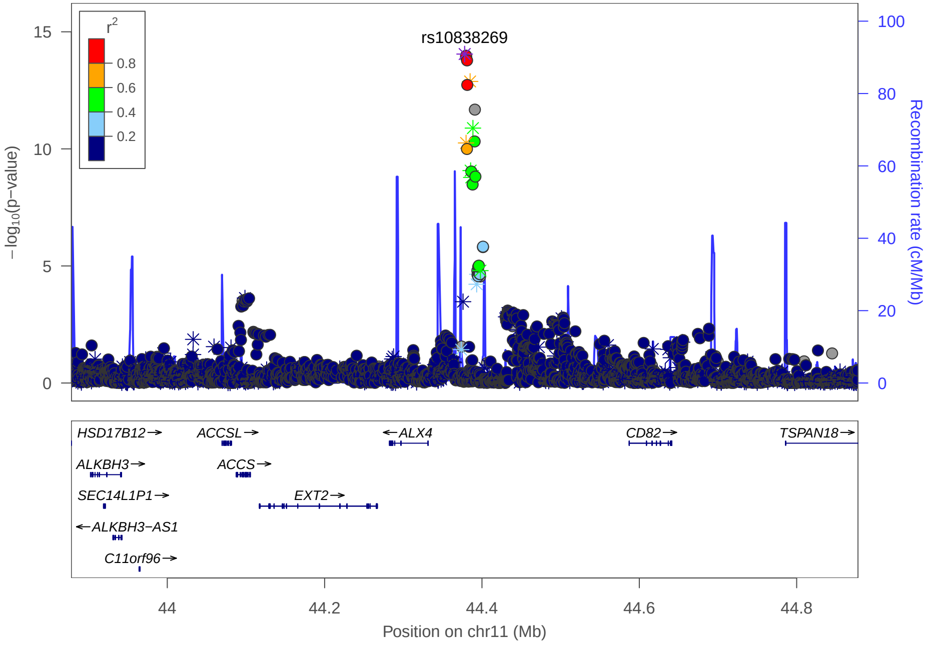
rs10838269
r²
0.8
0.6
0.4
0.2
−log10(p−value)
Recombination rate (cM/Mb)
HSD17B12→
ACCSL→
←ALX4
CD82→
TSPAN18→
ALKBH3→
ACCS→
SEC14L1P1→
EXT2→
←ALKBH3−AS1
C11orf96→
44
44.2
44.4
44.6
44.8
Position on chr11 (Mb)

**C**

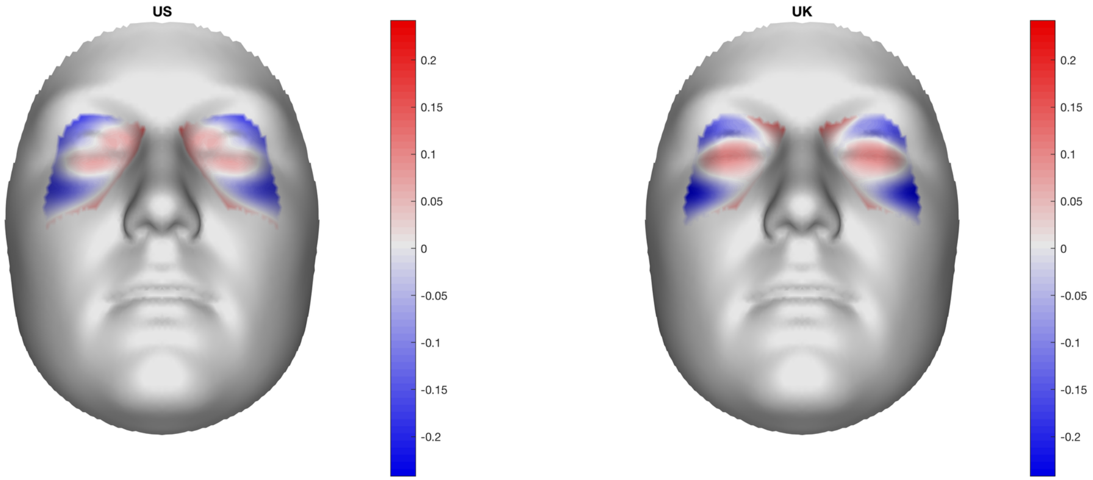
US
UK
0.2
0.15
0.1
0.05
0
-0.05
-0.1
-0.15
-0.2

**B**

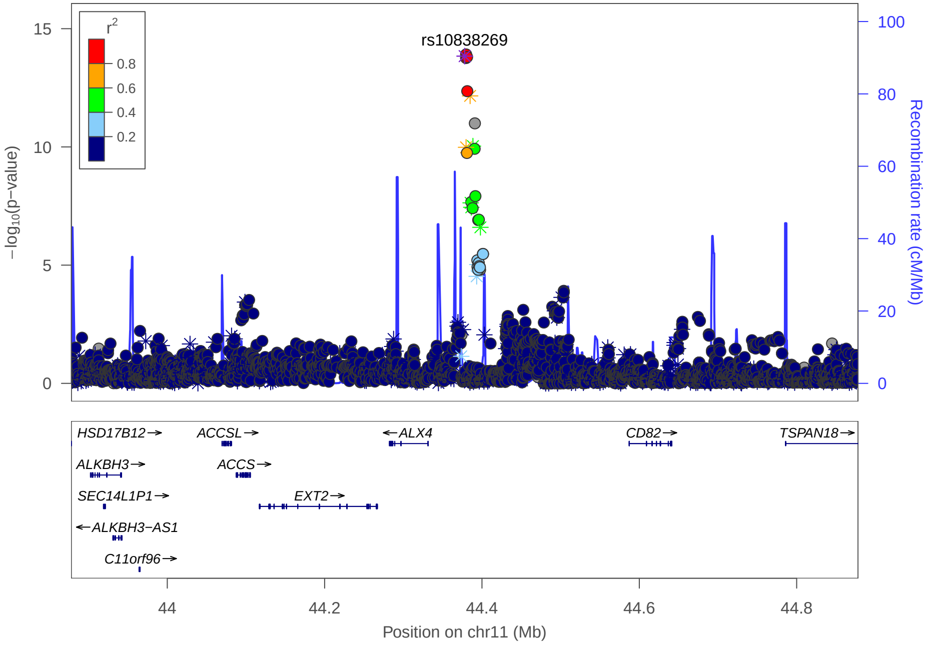
rs10838269
r²
0.8
0.6
0.4
0.2
−log10(p−value)
Recombination rate (cM/Mb)
HSD17B12→
ACCSL→
←ALX4
CD82→
TSPAN18→
ALKBH3→
ACCS→
SEC14L1P1→
EXT2→
←ALKBH3−AS1
C11orf96→
44
44.2
44.4
44.6
44.8
Position on chr11 (Mb)

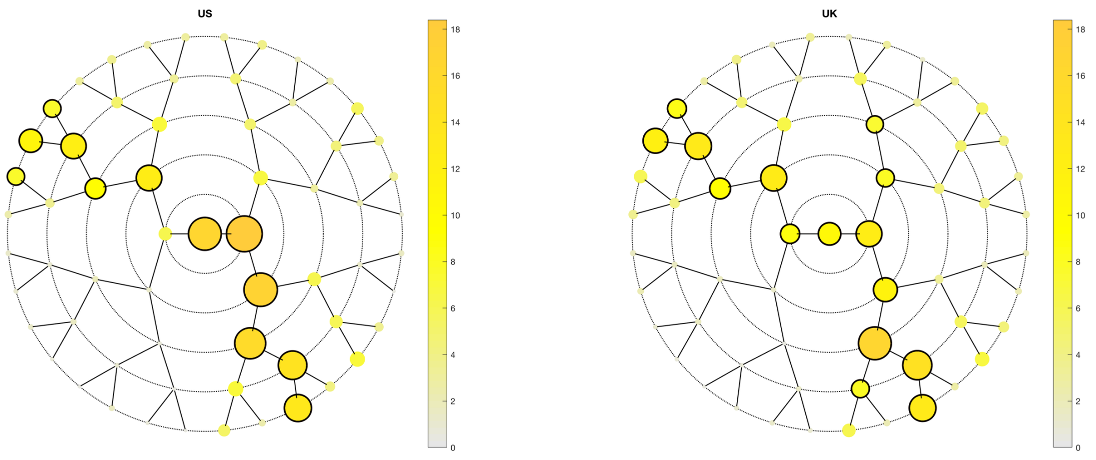
US
UK

A

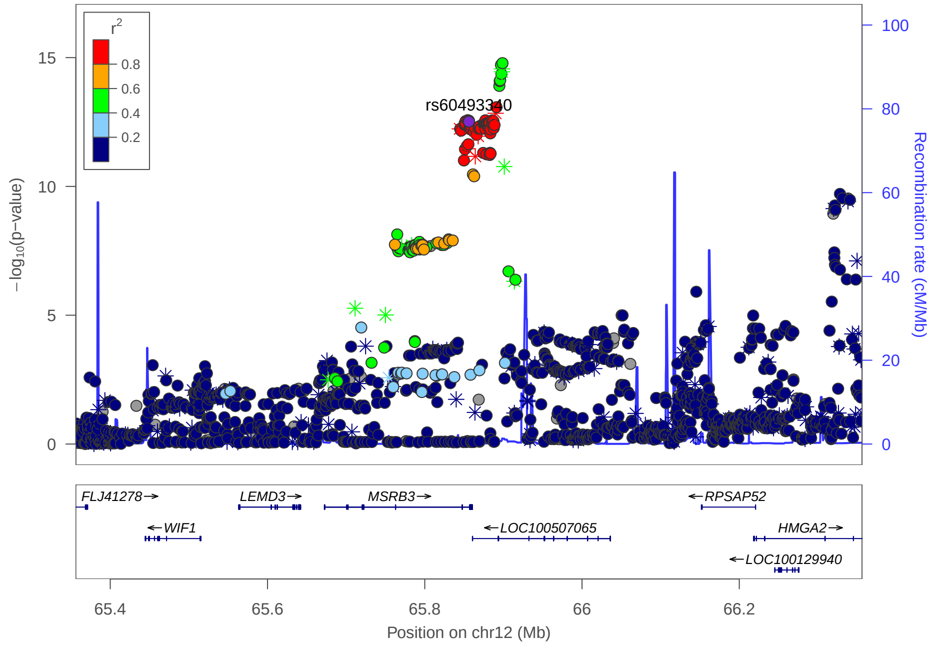
rs60493340
r2
−log10(p−value)
Recombination rate (cM/Mb)
FLJ41278
WIF1
LEMD3
MSRB3
LOC100507065
RPSAP52
HMGA2
LOC100129940
Position on chr12 (Mb)

C

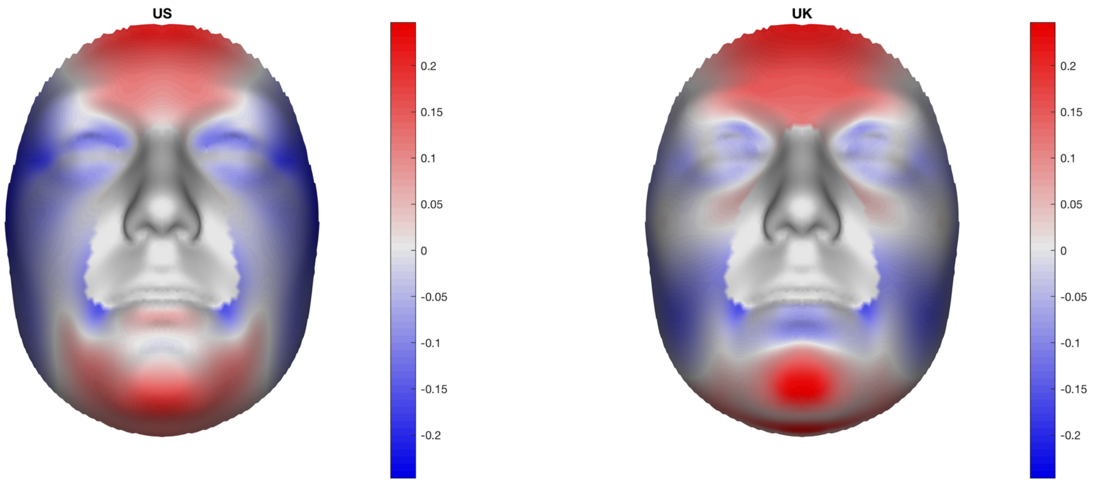
US
UK

B

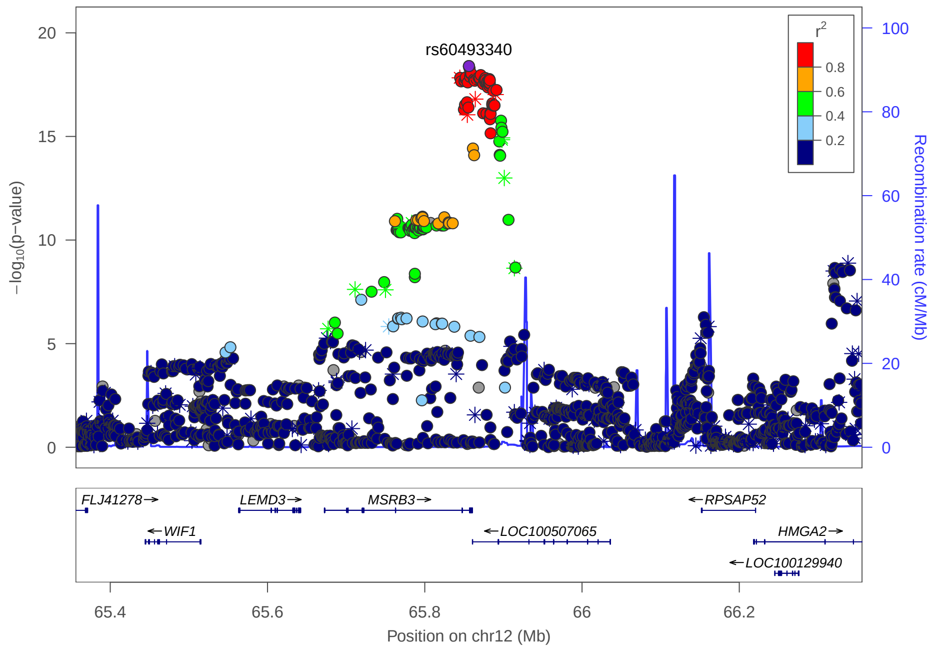
rs60493340
r2
−log10(p−value)
Recombination rate (cM/Mb)
FLJ41278
WIF1
LEMD3
MSRB3
LOC100507065
RPSAP52
HMGA2
LOC100129940
Position on chr12 (Mb)

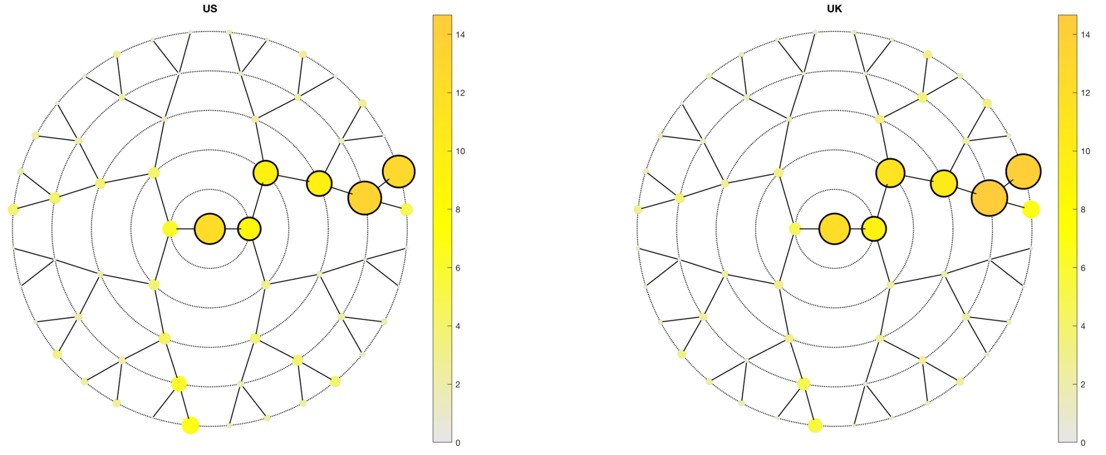

US
UK

A

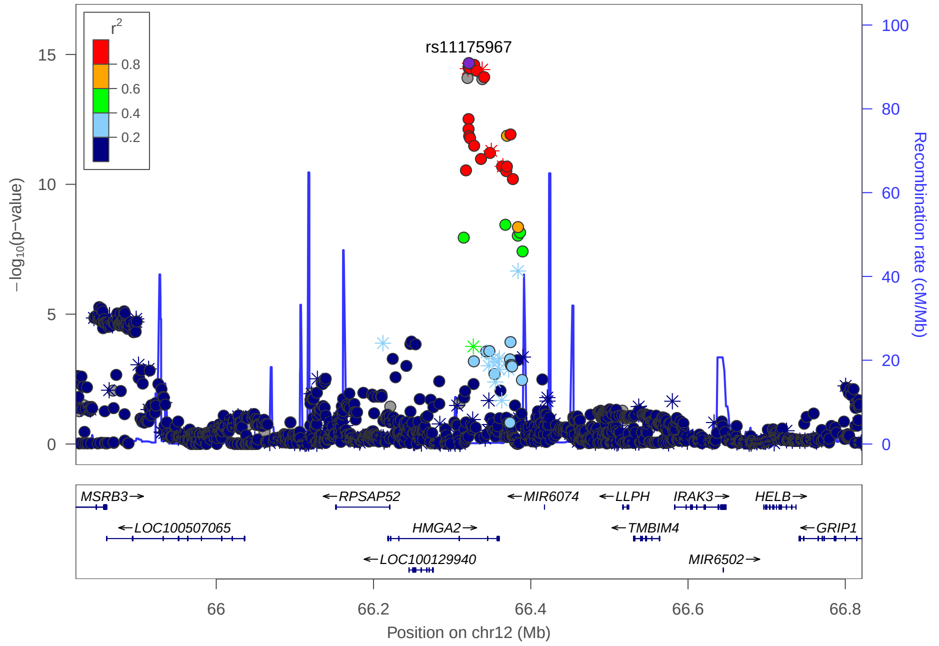

rs11175967
r2
−log10(p−value)
Recombination rate (cM/Mb)
MSRB3
RPSAP52
MIR6074
LLPH
IRAK3
HELB
LOC100507065
HMGA2
TMBIM4
GRIP1
LOC100129940
MIR6502
Position on chr12 (Mb)

C

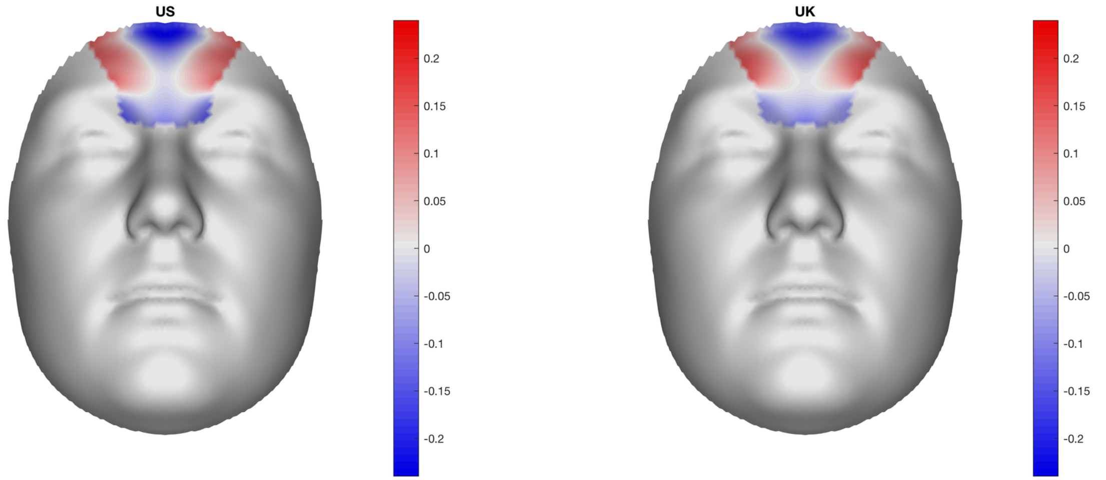

US
UK

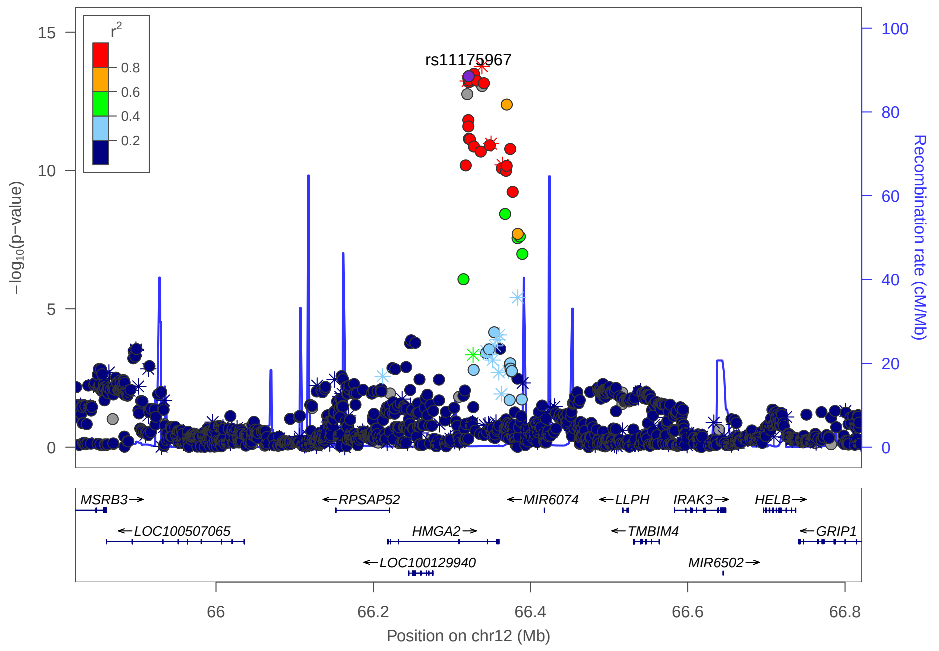

rs11175967
r2
−log10(p−value)
Recombination rate (cM/Mb)
MSRB3
RPSAP52
MIR6074
LLPH
IRAK3
HELB
LOC100507065
HMGA2
TMBIM4
GRIP1
LOC100129940
MIR6502
Position on chr12 (Mb)

B

### Multi-peak locus 17: rs2695152

**CHR12 -** See final page for captions

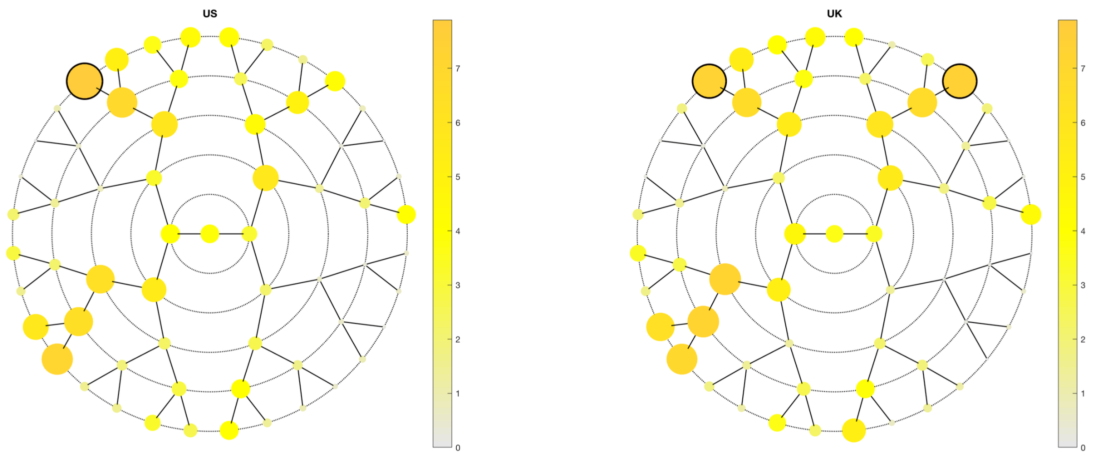

**A**

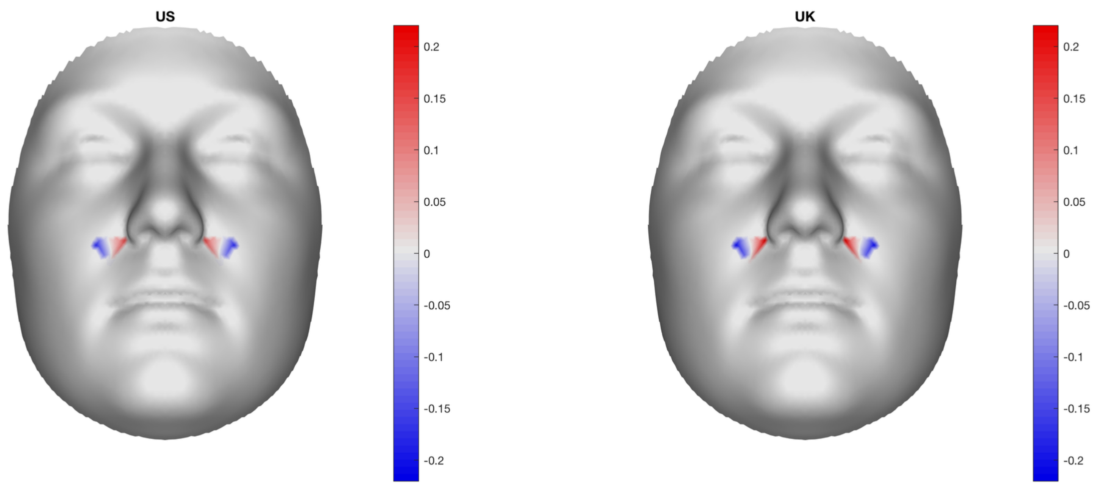

**B**

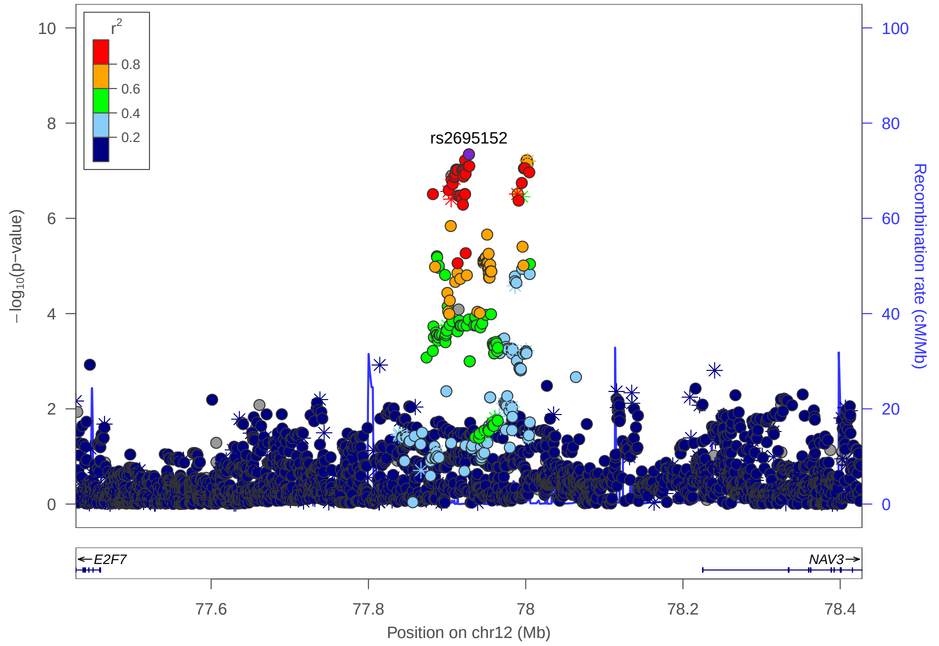

**C**

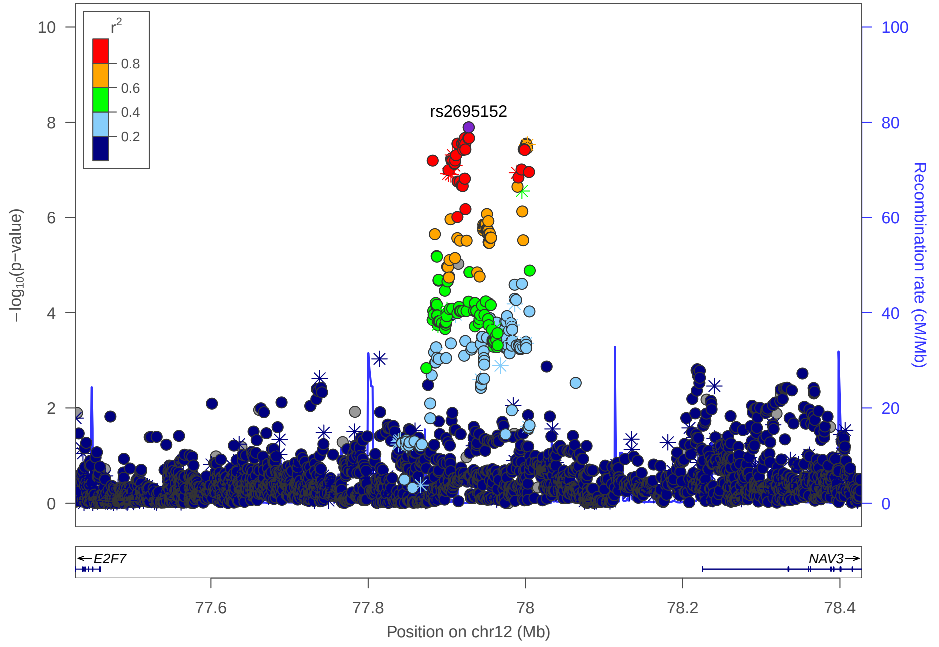

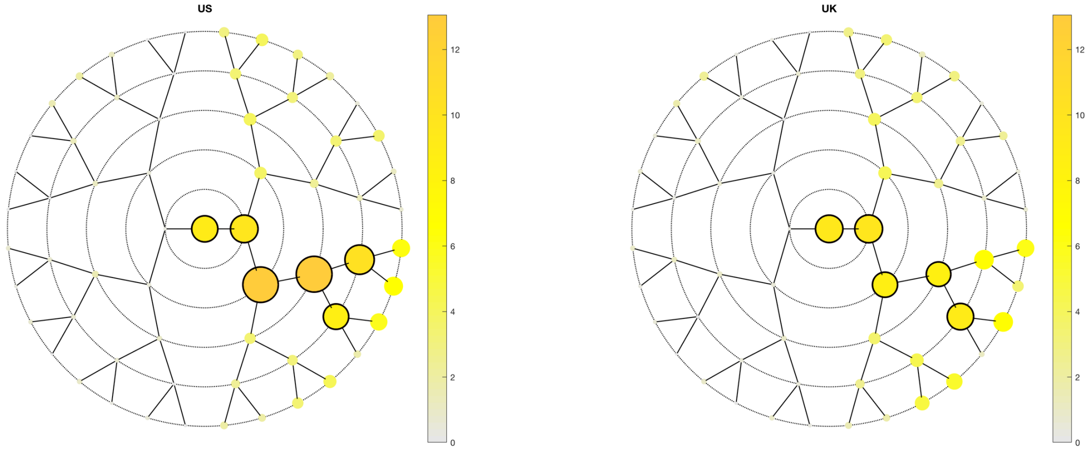

US
UK
0
2
4
6
8
10
12

**A**

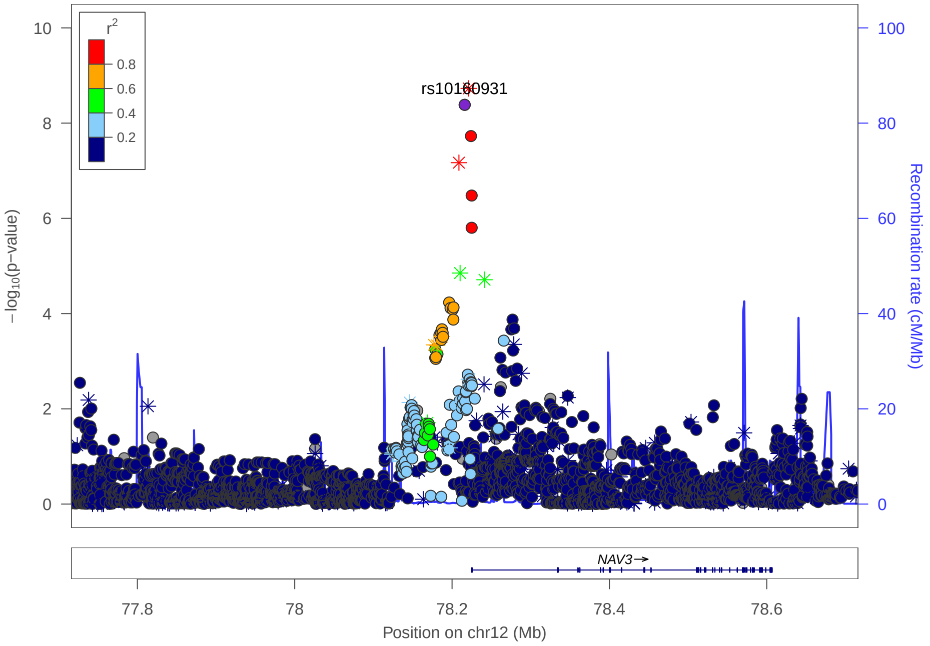

rs10160931
−log10(p−value)
Recombination rate (cM/Mb)
r2
0.8
0.6
0.4
0.2
NAV3→
77.8
78
78.2
78.4
78.6
Position on chr12 (Mb)

**C**

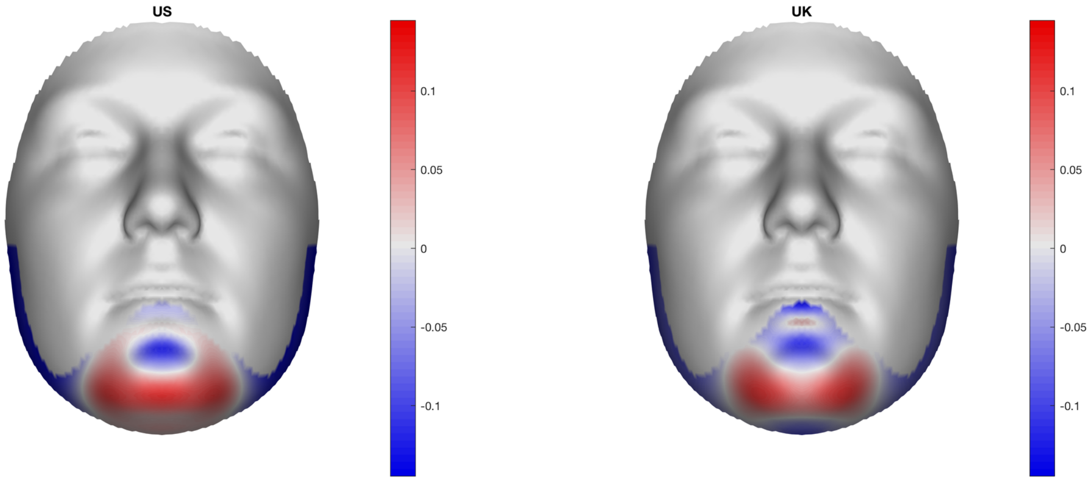

US
UK
0.1
0.05
0
-0.05
-0.1

**B**

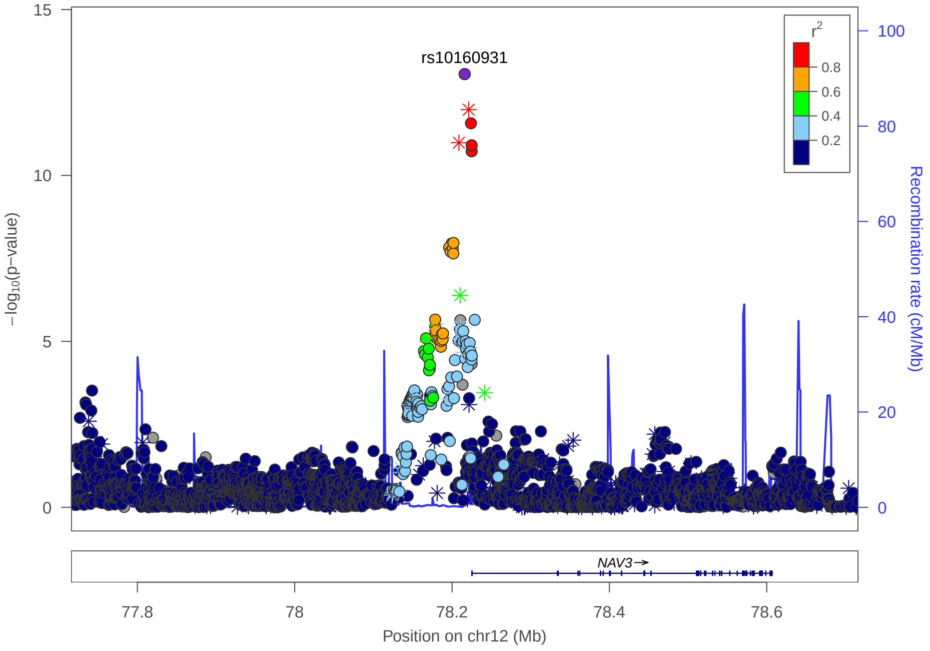

rs10160931
−log10(p−value)
Recombination rate (cM/Mb)
r2
0.8
0.6
0.4
0.2
NAV3→
77.8
78
78.2
78.4
78.6
Position on chr12 (Mb)

### Multi-peak locus 18: rs58687115

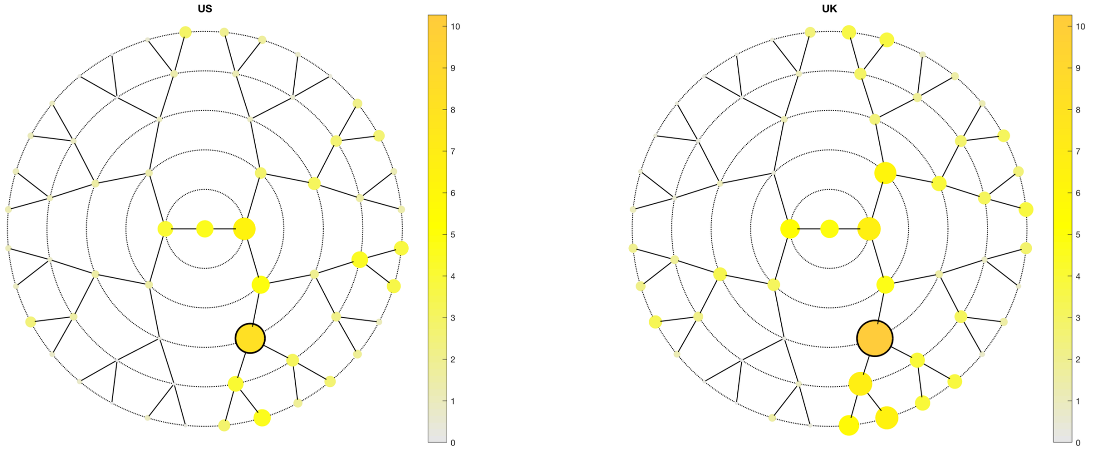

**A**

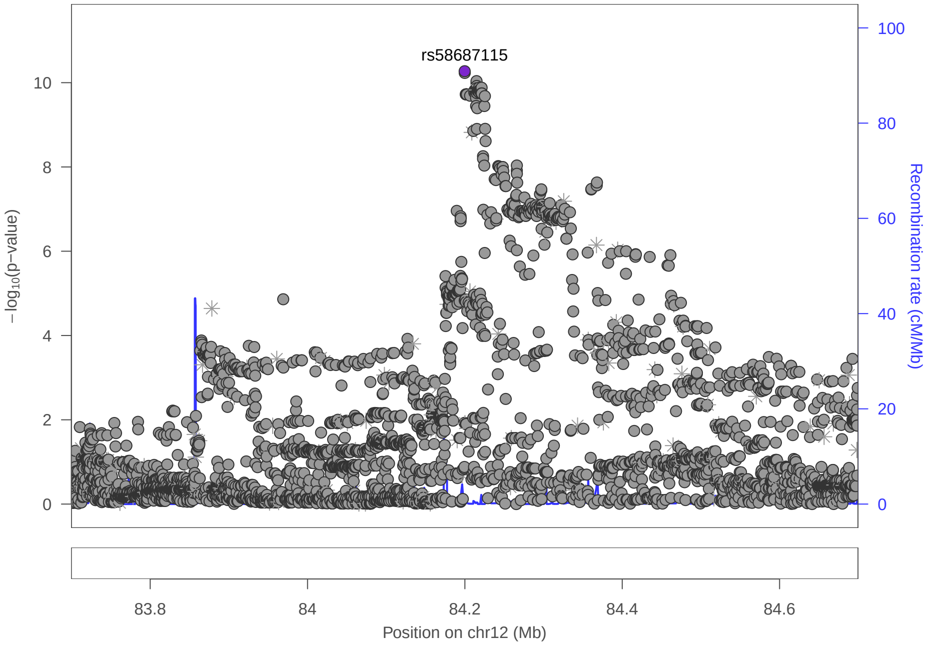

**C**

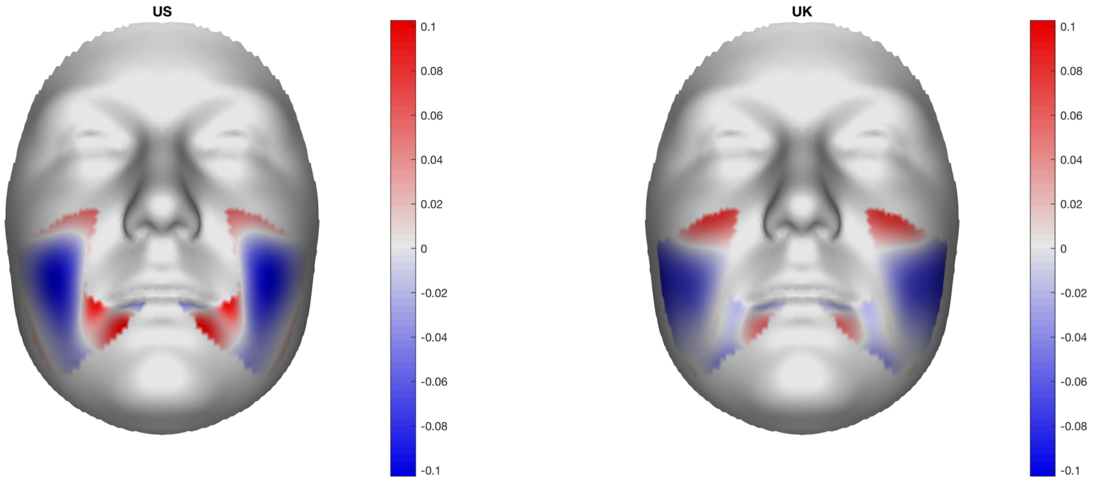

**B**

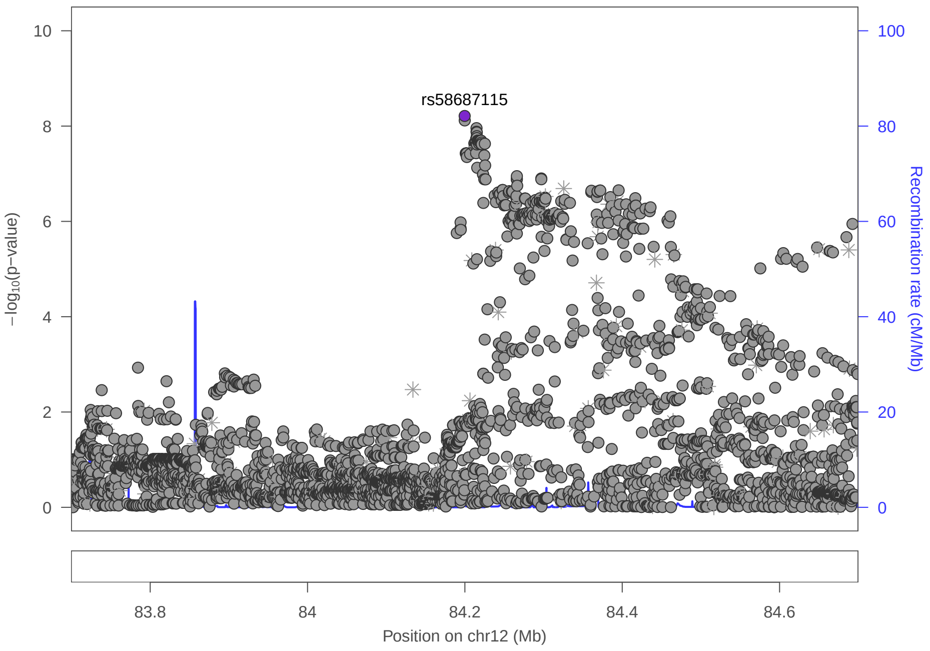

### Multi-peak locus 18: rs4385969

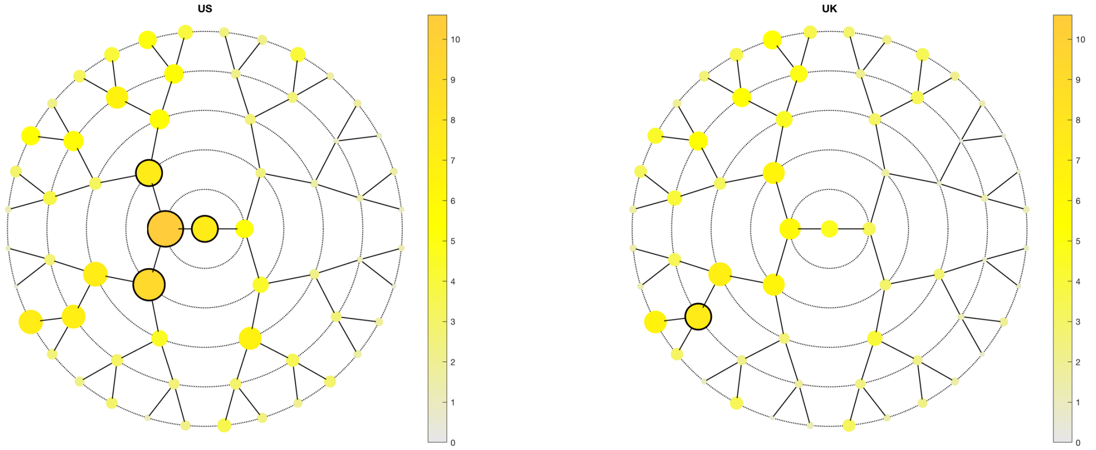

**A**

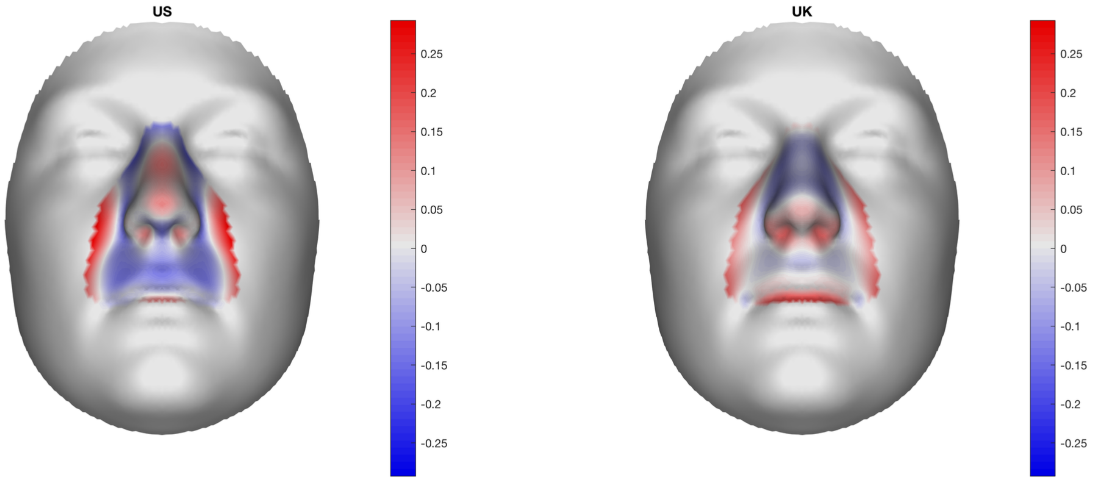

**B**

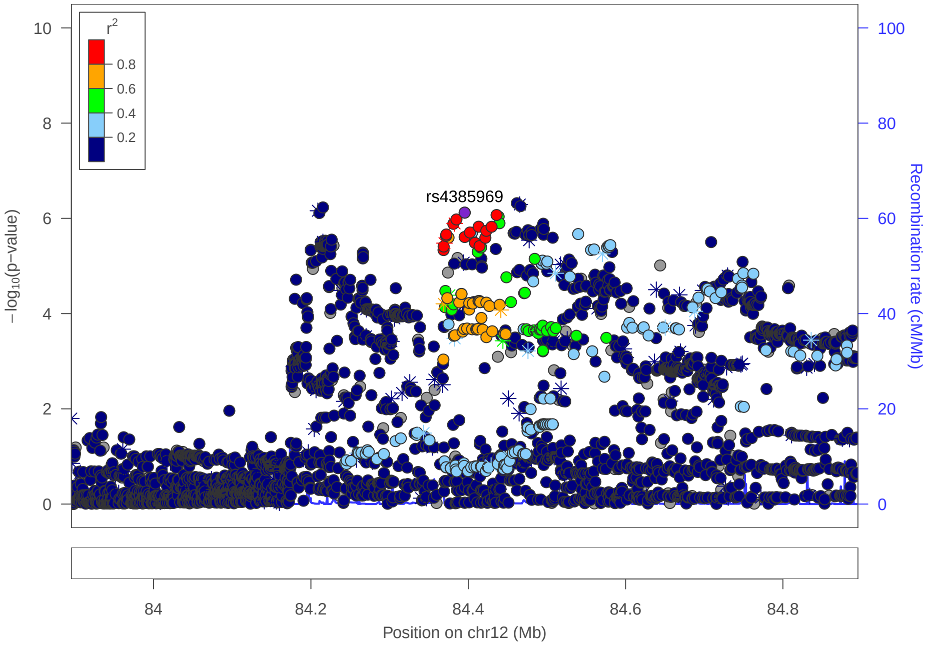

**C**

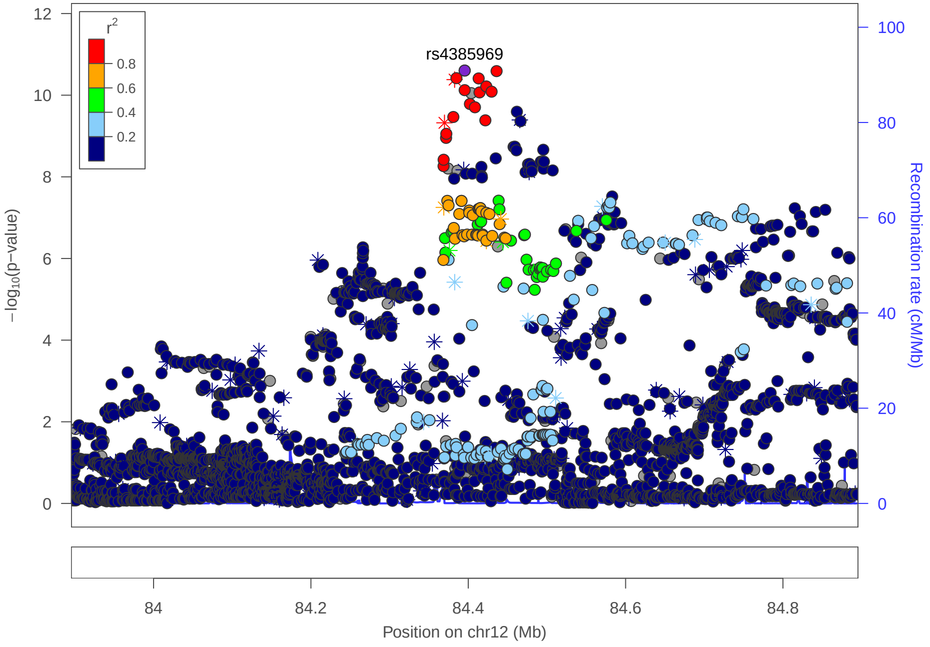

### Multi-peak locus 19: rs11609649

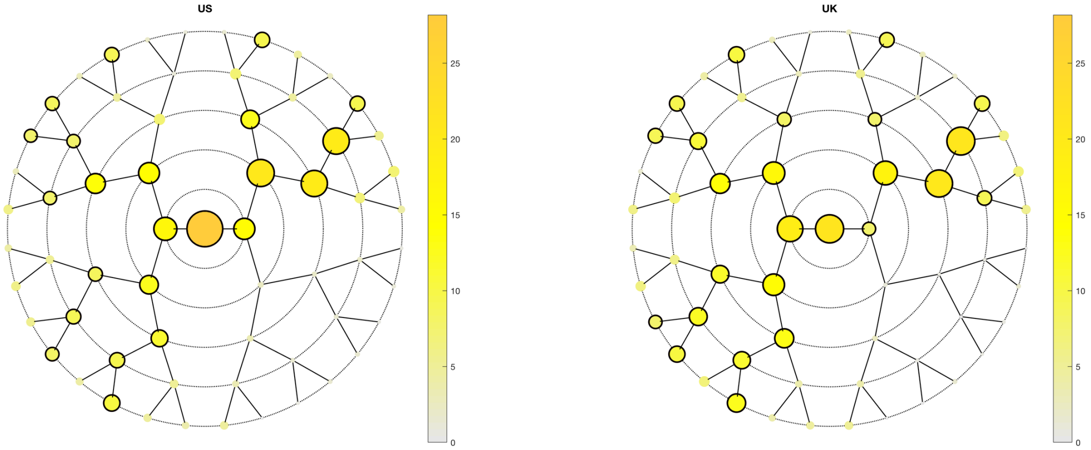

**A**

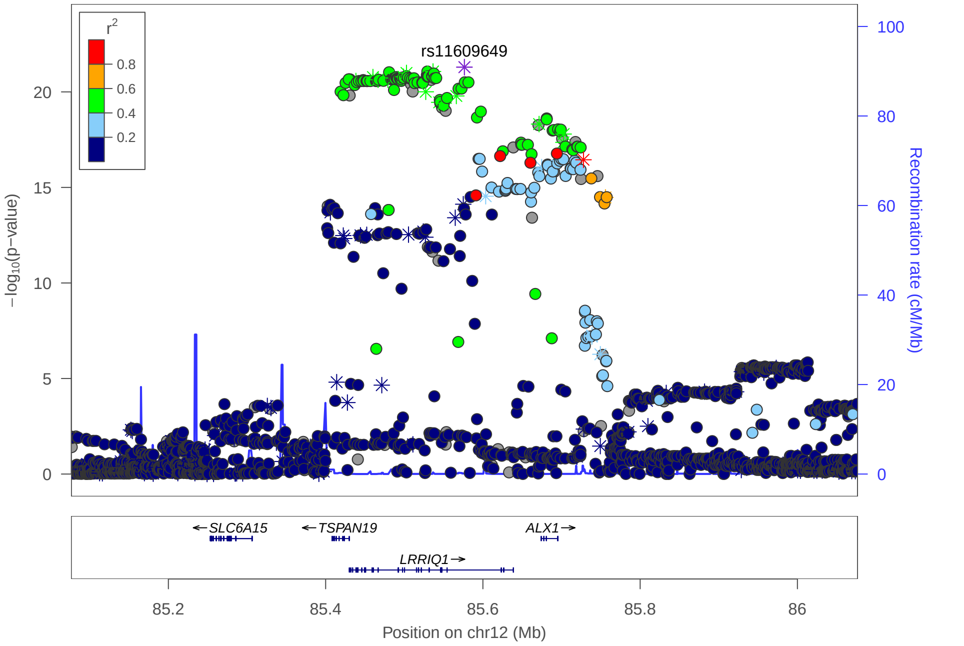

**C**

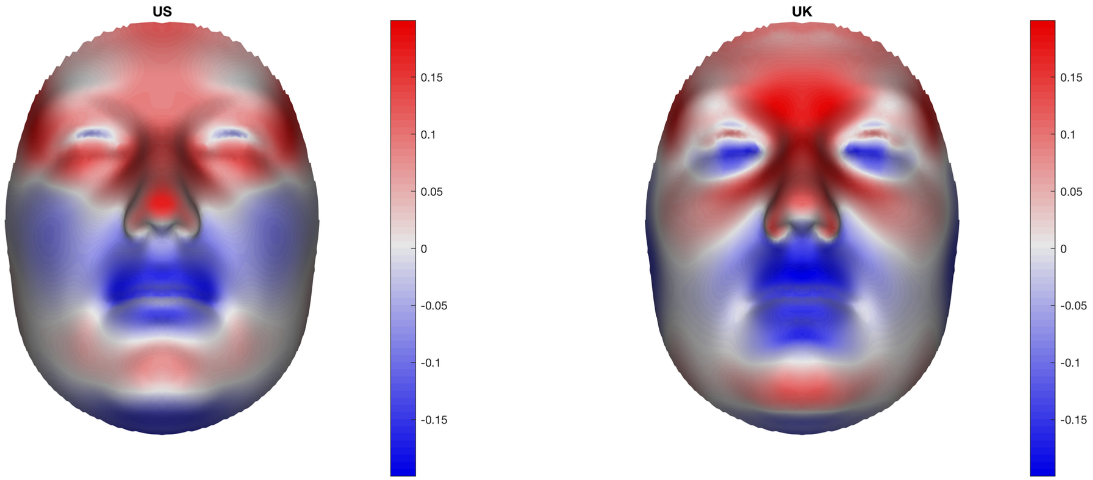

**B**

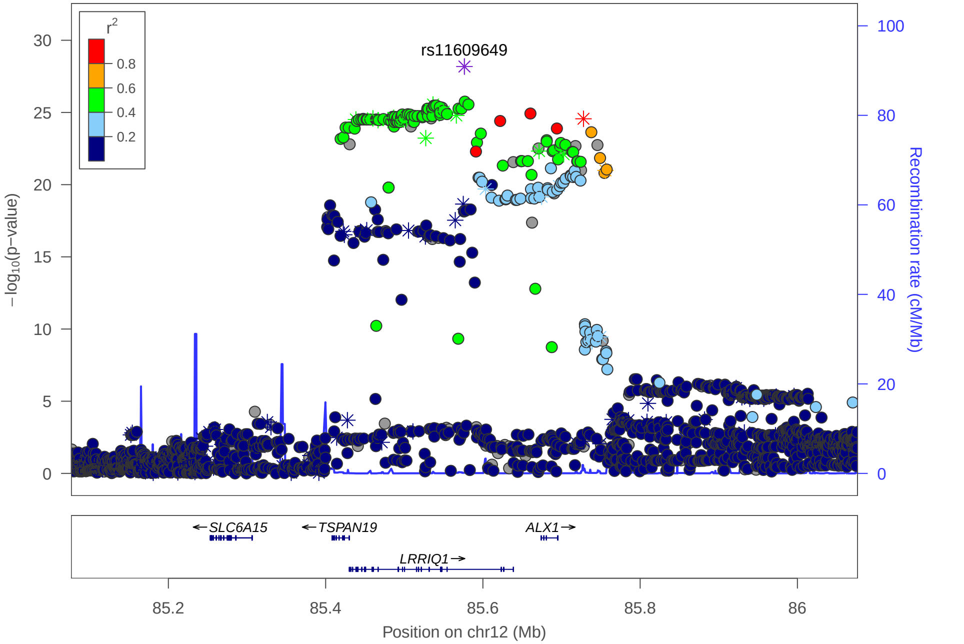

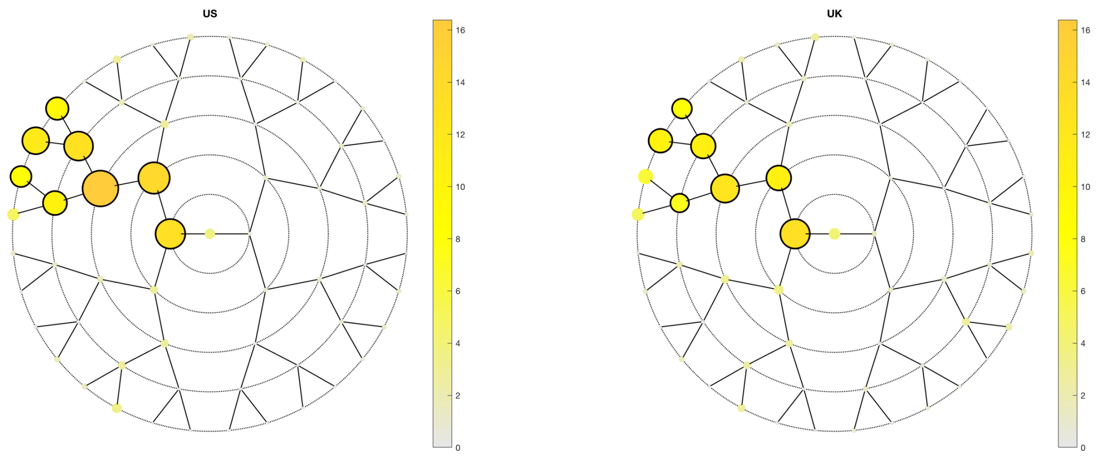

US
UK

**A**

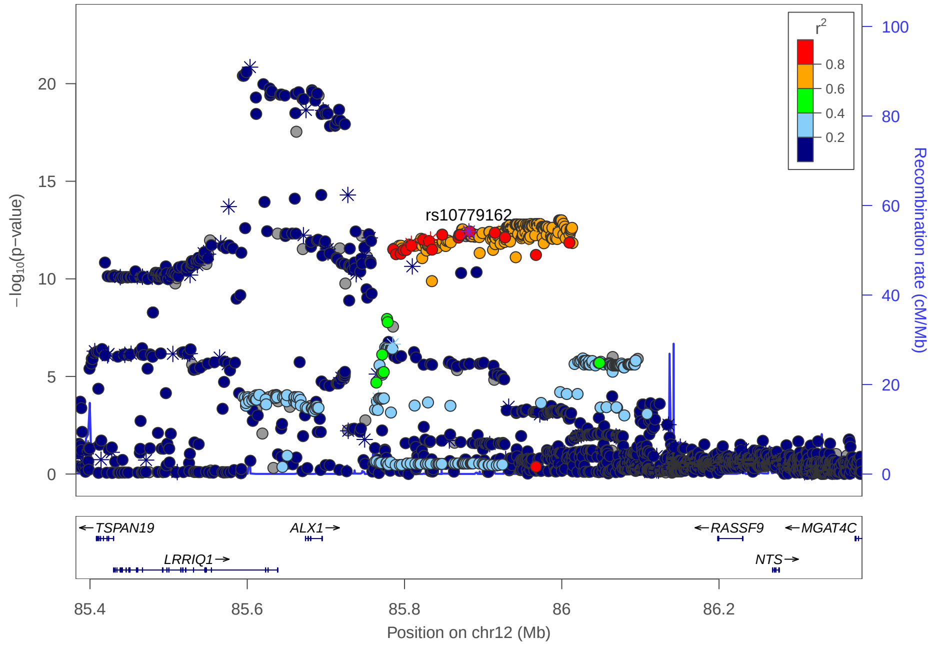

rs10779162
$r^2$
0.8
0.6
0.4
0.2
−log10(p−value)
Recombination rate (cM/Mb)
TSPAN19
ALX1
RASSF9
MGAT4C
LRRIQ1
NTS
85.4
85.6
85.8
86
86.2
Position on chr12 (Mb)

**C**

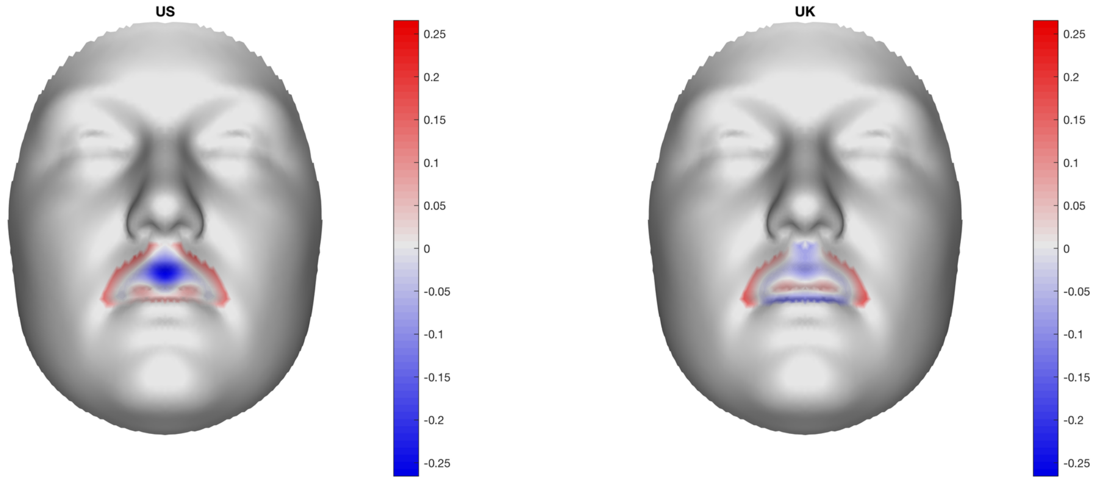

US
UK

**B**

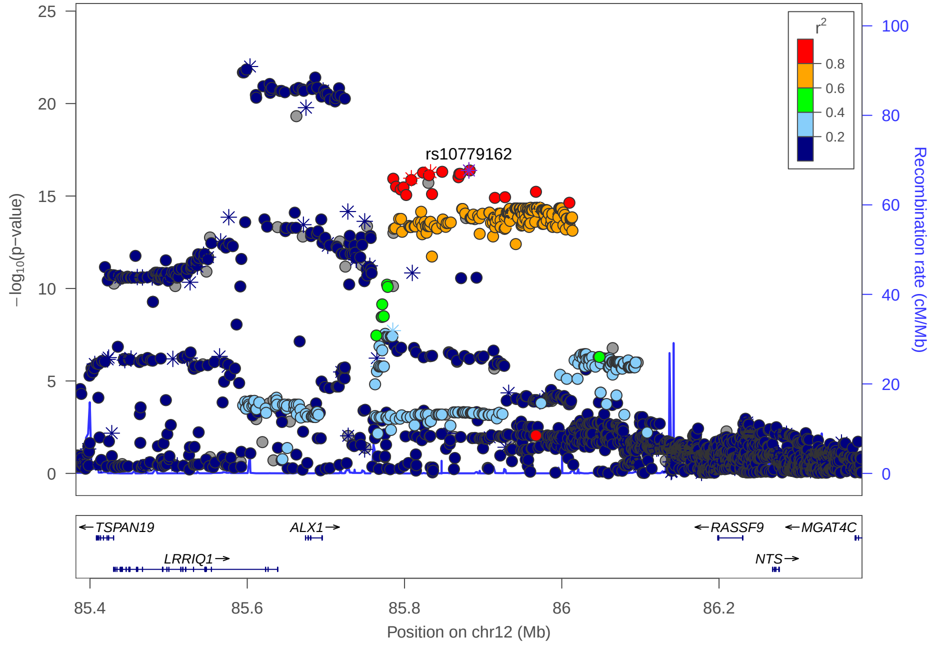

rs10779162
$r^2$
0.8
0.6
0.4
0.2
−log10(p−value)
Recombination rate (cM/Mb)
TSPAN19
ALX1
RASSF9
MGAT4C
LRRIQ1
NTS
85.4
85.6
85.8
86
86.2
Position on chr12 (Mb)

### Multi-peak locus 20: rs11842203

**CHR13** - See final page for captions

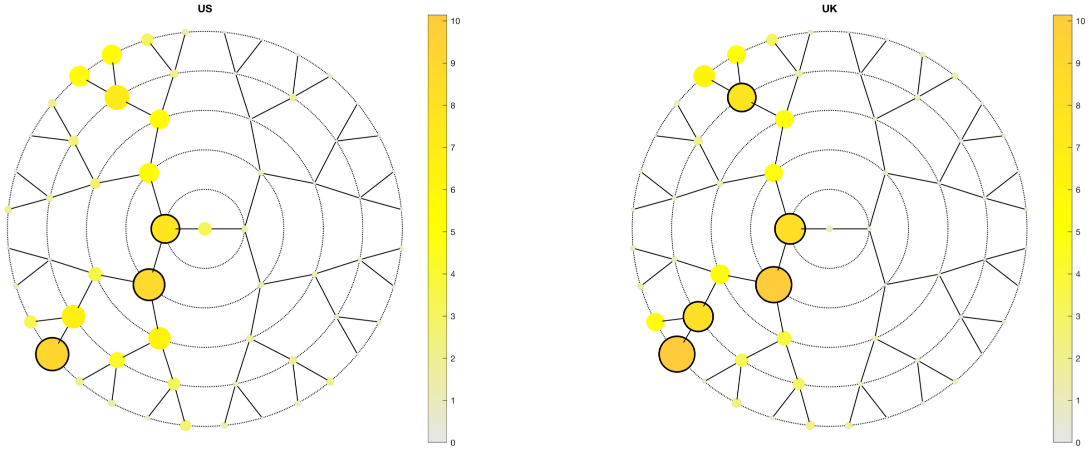

A

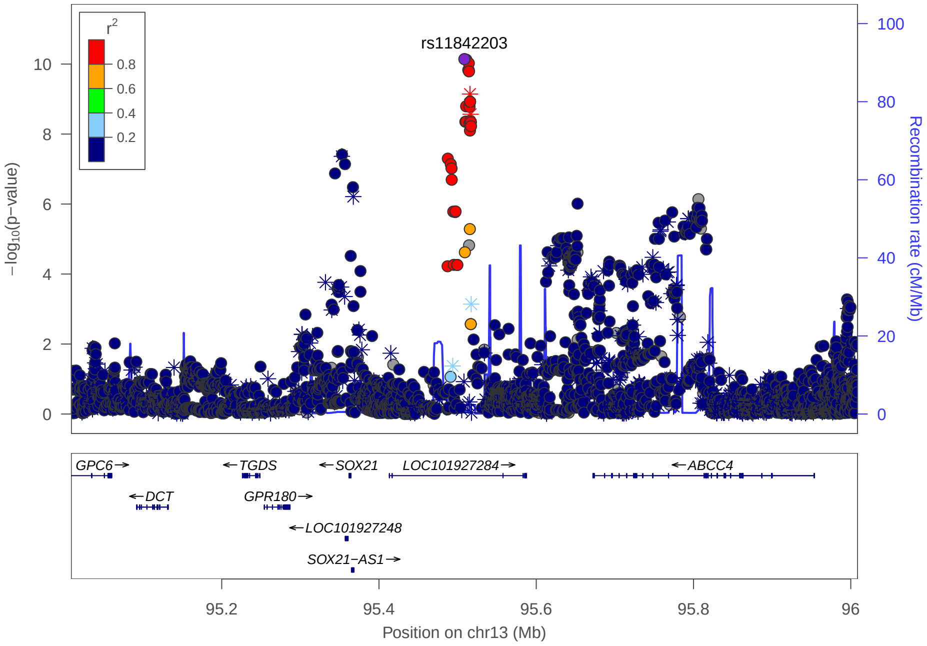

C

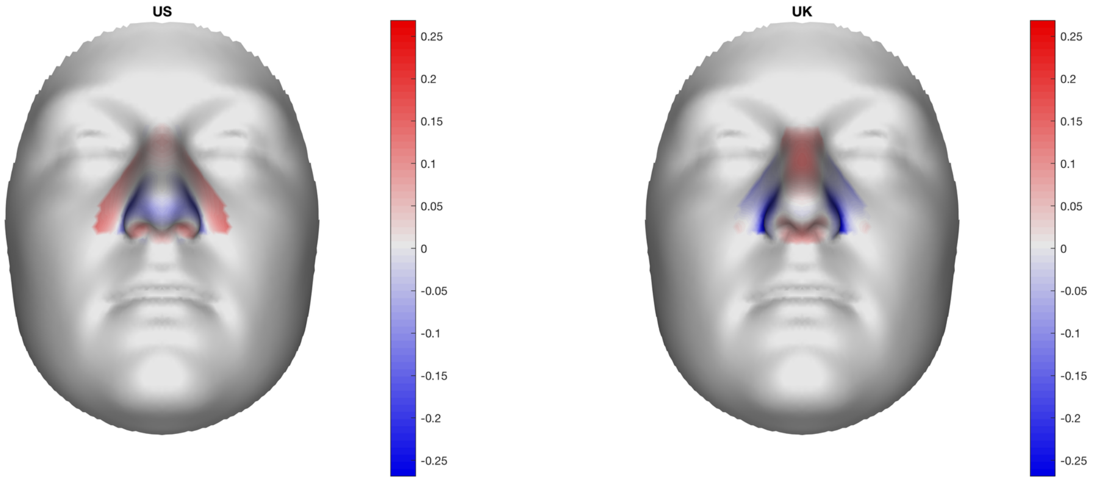

B

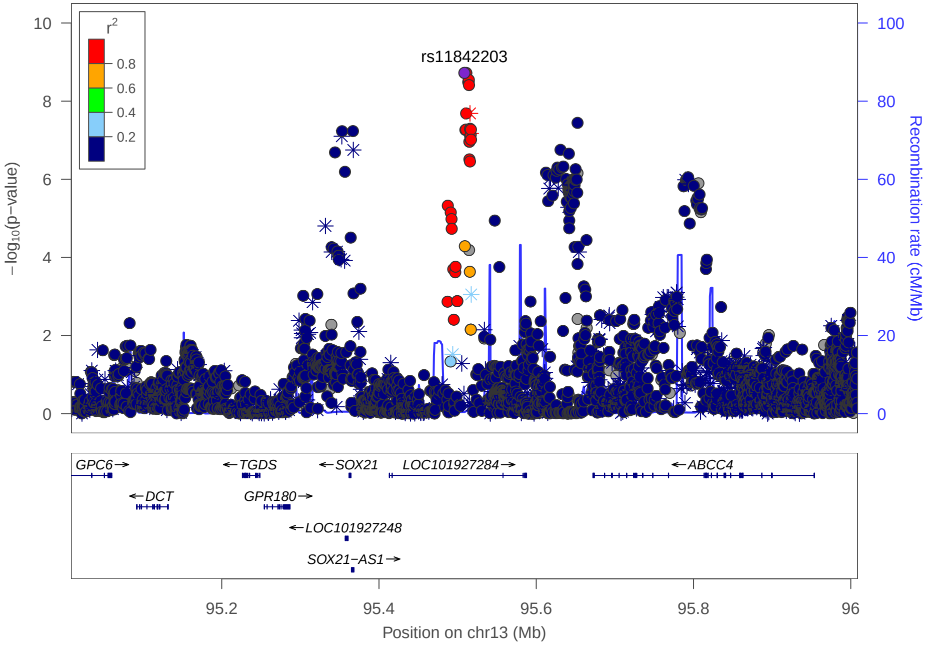

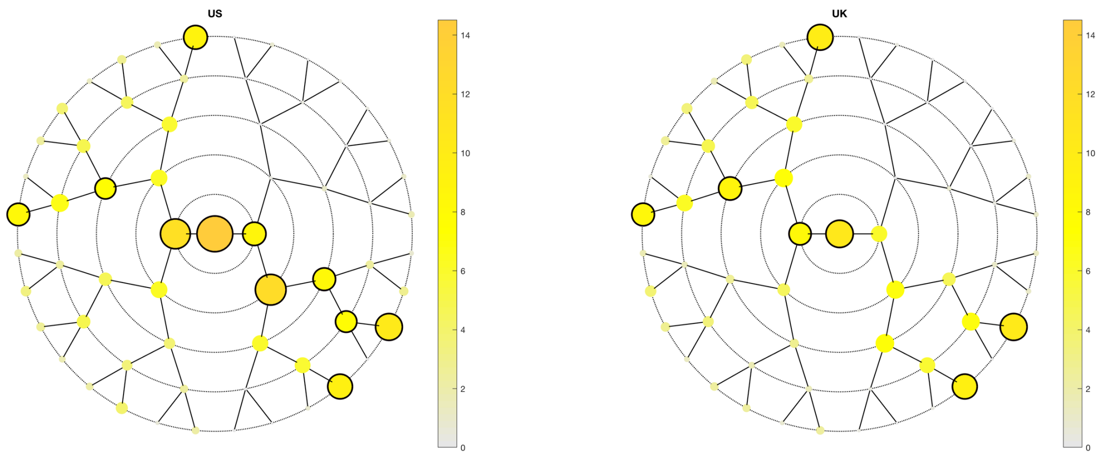
US
UK

A

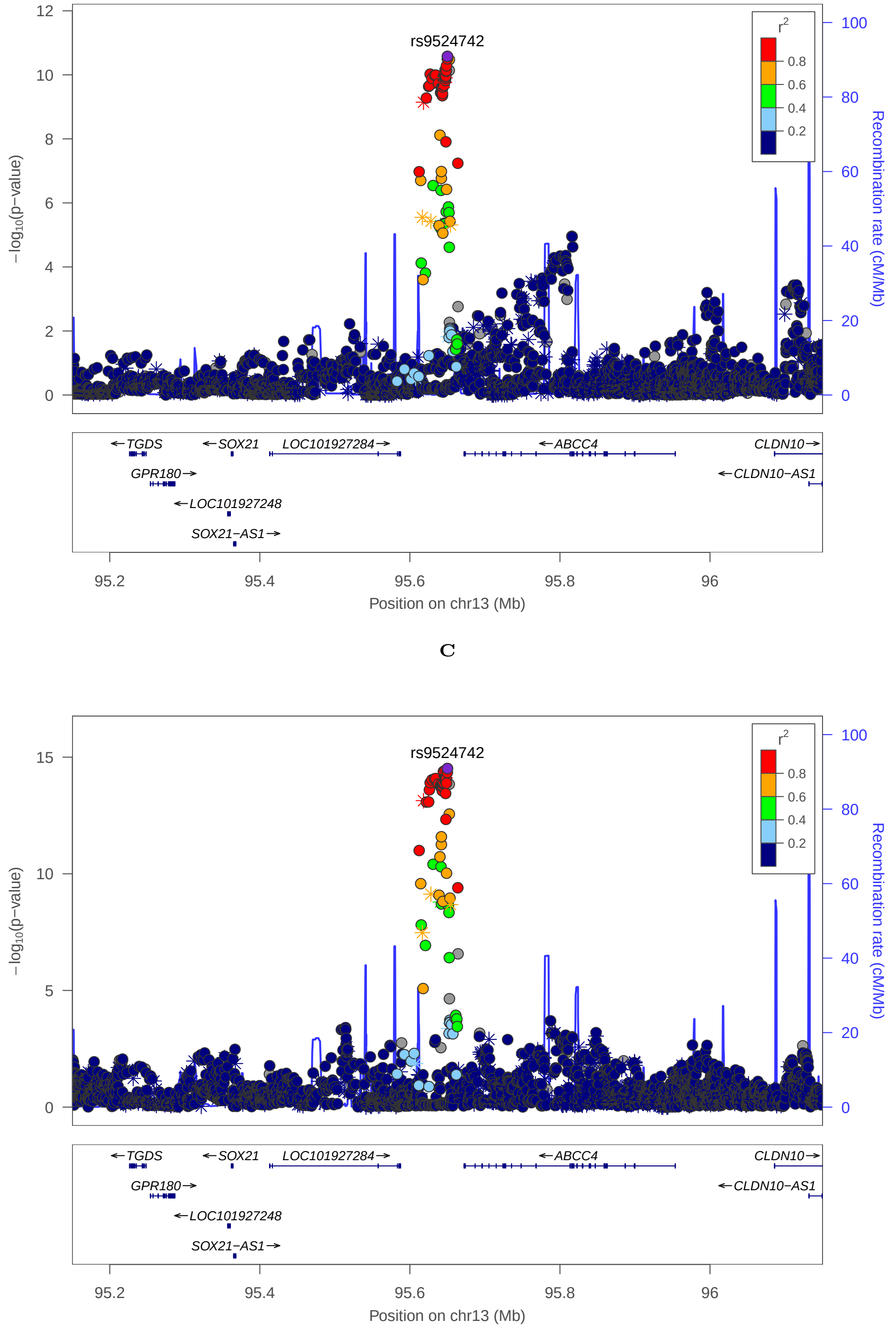
rs9524742
−log10(p−value)
Recombination rate (cM/Mb)
Position on chr13 (Mb)
TGDS
SOX21
LOC101927284
ABCC4
CLDN10
GPR180
CLDN10−AS1
LOC101927248
SOX21−AS1

C

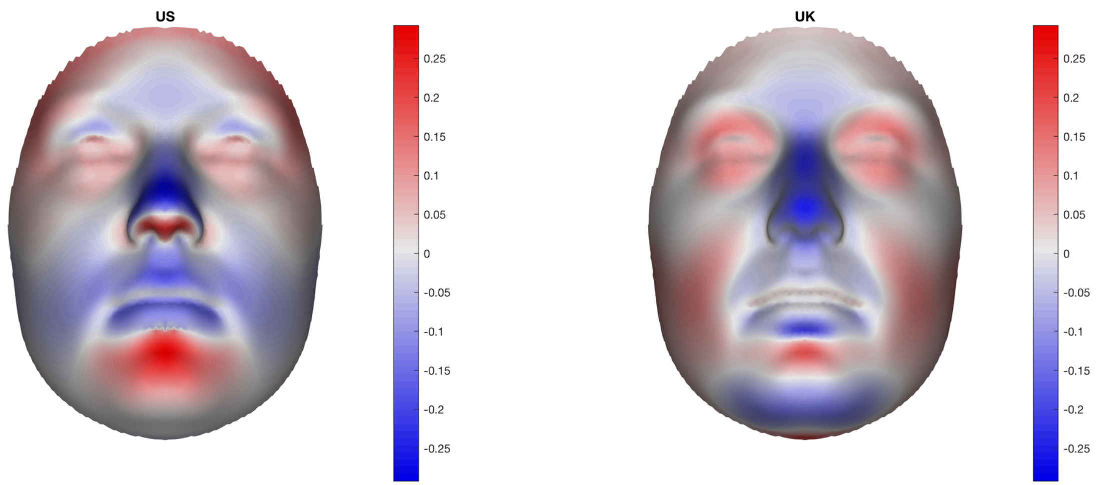
US
UK

B

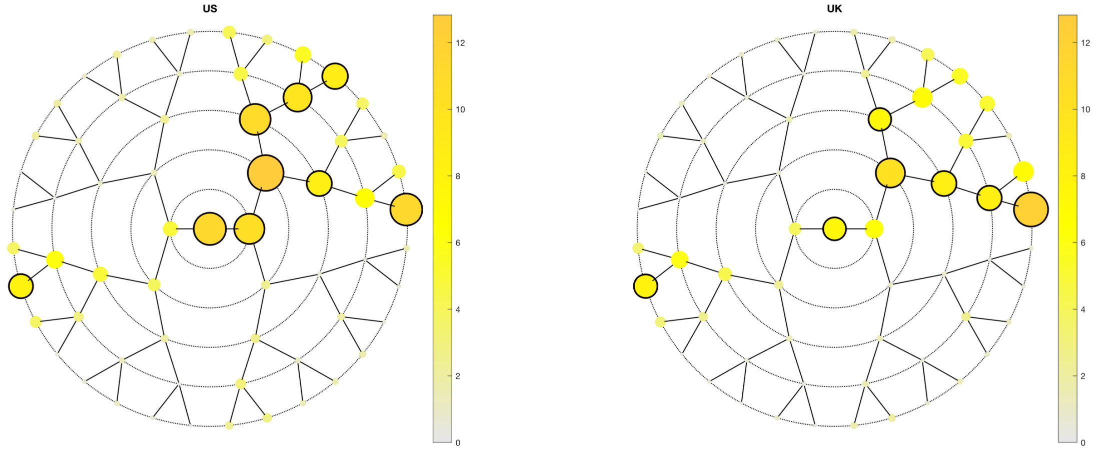
US
UK

**A**

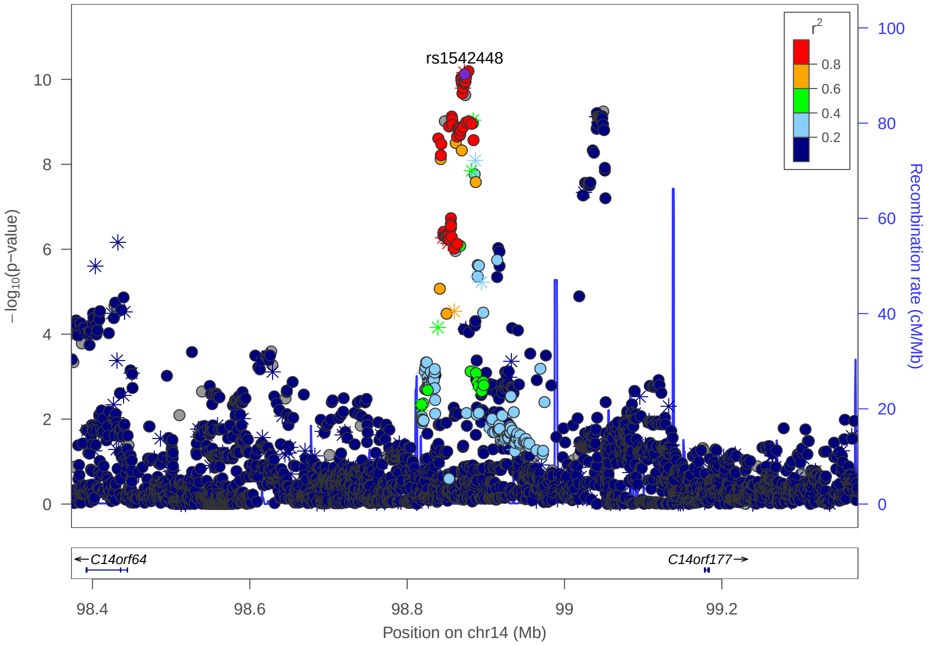
rs1542448
−log10(p−value)
Recombination rate (cM/Mb)
r2
C14orf64
C14orf177
Position on chr14 (Mb)

**C**

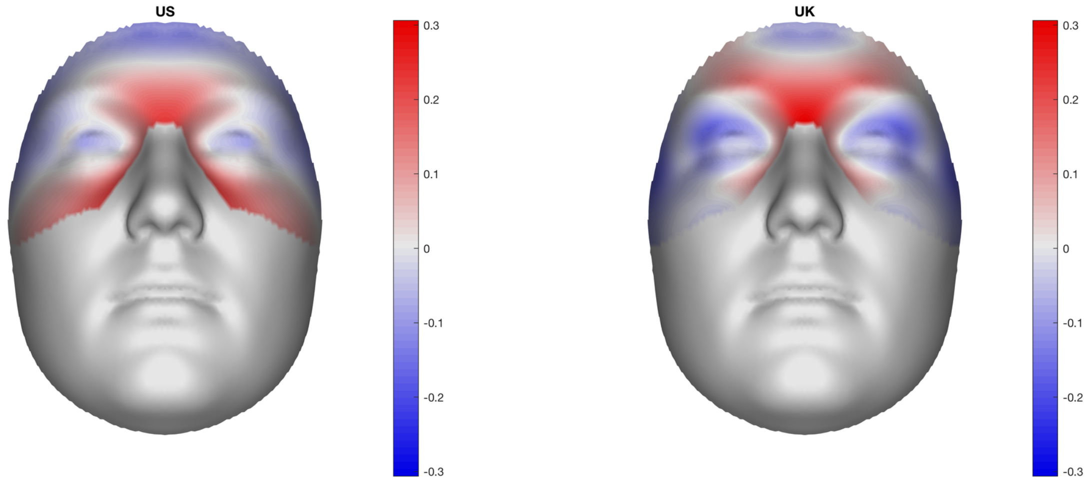
US
UK

**B**

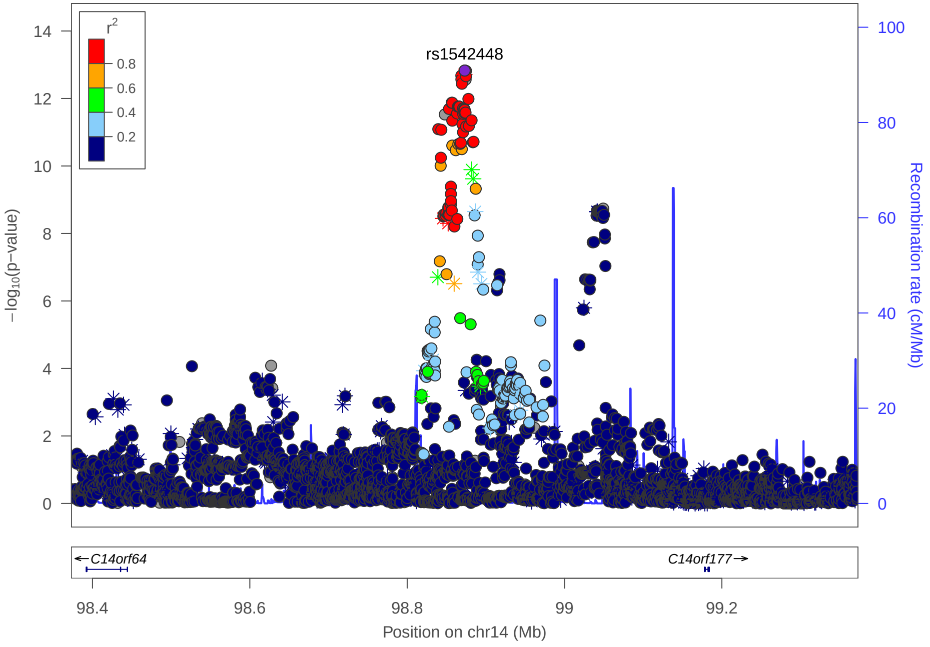
rs1542448
−log10(p−value)
Recombination rate (cM/Mb)
r2
C14orf64
C14orf177
Position on chr14 (Mb)

### Multi-peak locus 21: rs76810699

**CHR14** - See final page for captions

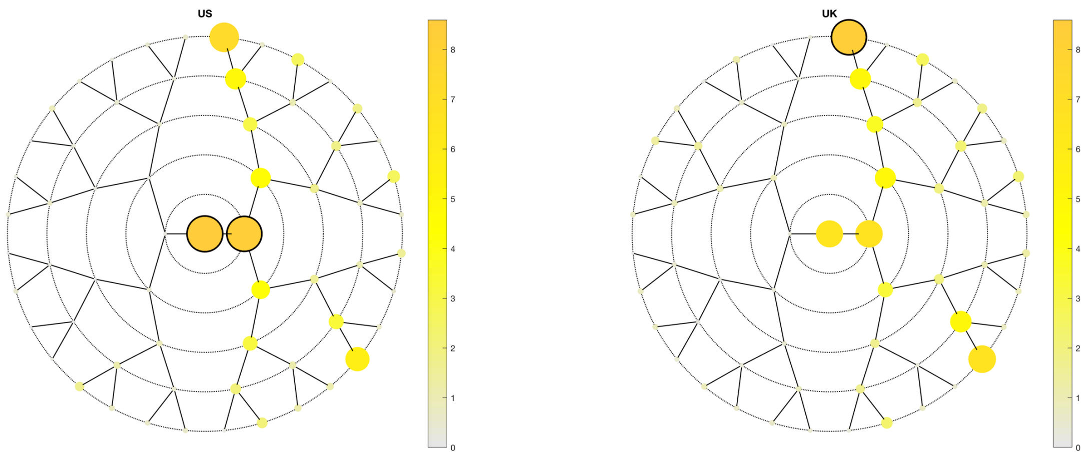

A

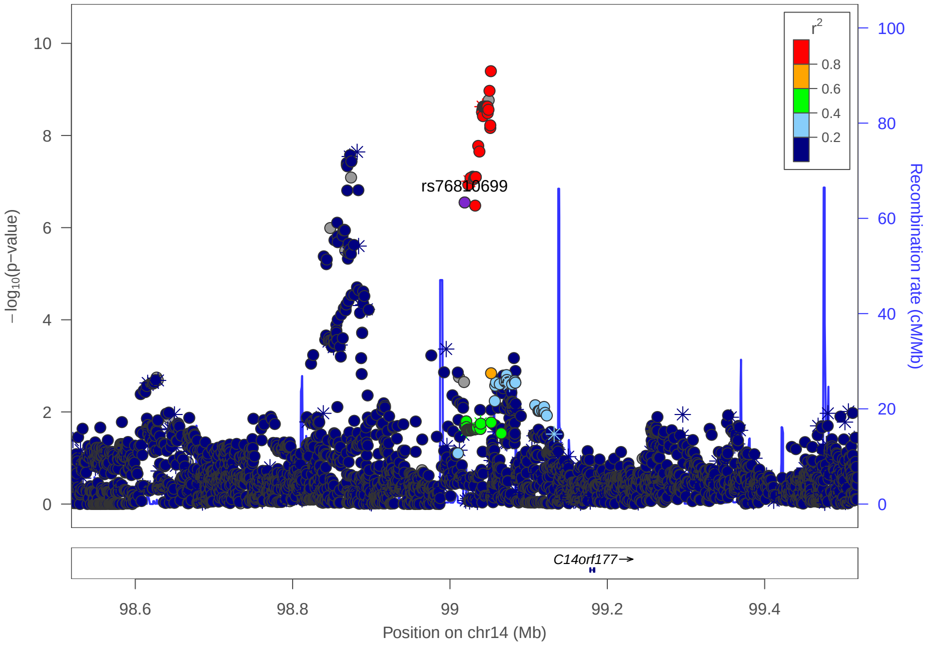

C

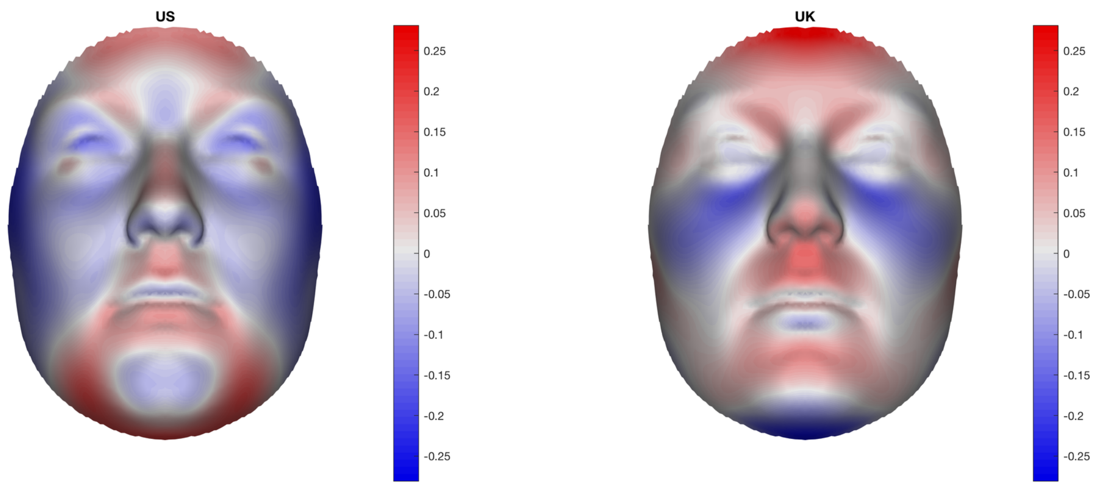

B

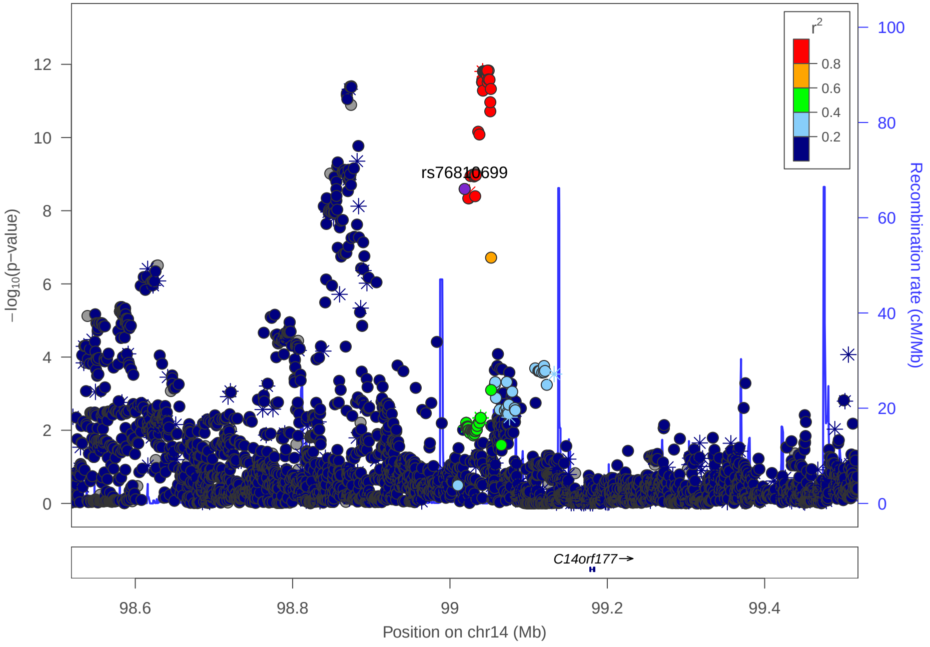

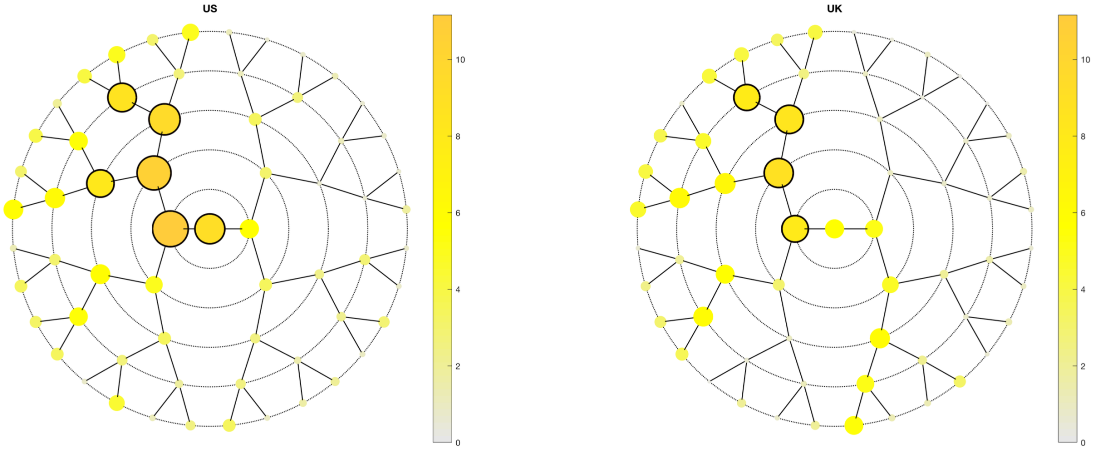

US
UK

**A**

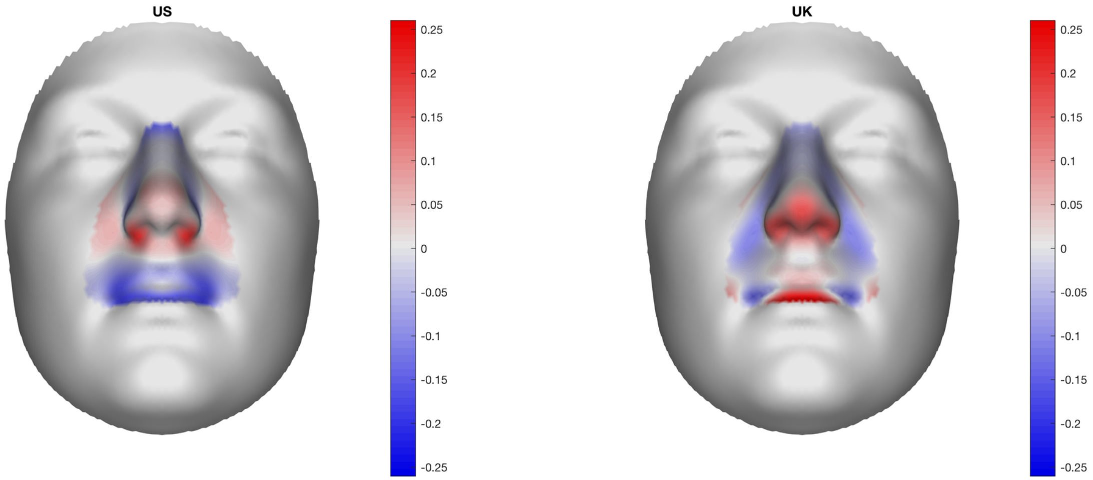

US
UK

**B**

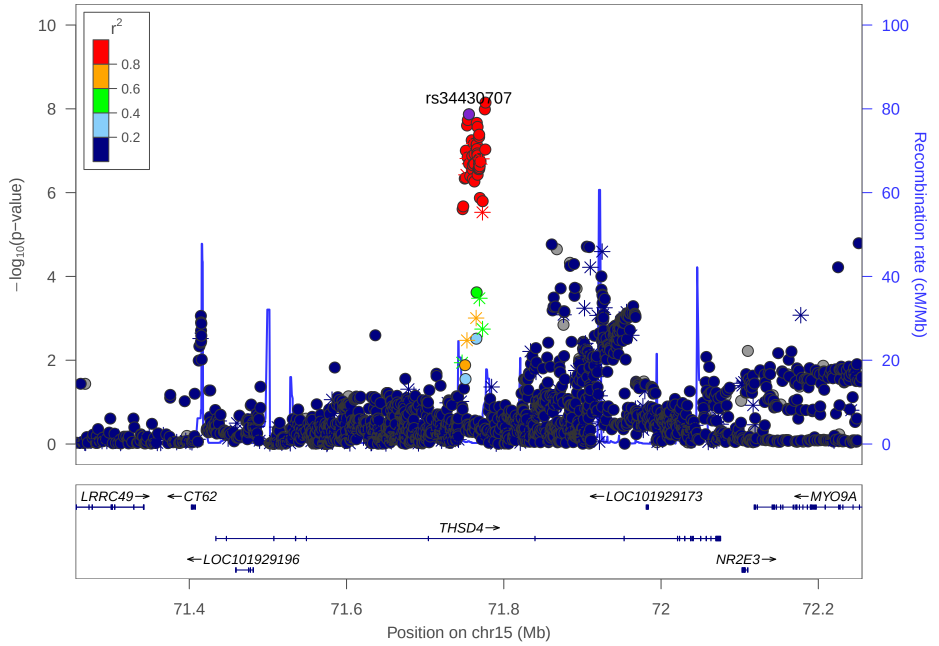

r2
0.8
0.6
0.4
0.2
rs34430707
−log10(p−value)
Recombination rate (cM/Mb)
LRRC49
CT62
LOC101929173
MYO9A
THSD4
LOC101929196
NR2E3
Position on chr15 (Mb)

**C**

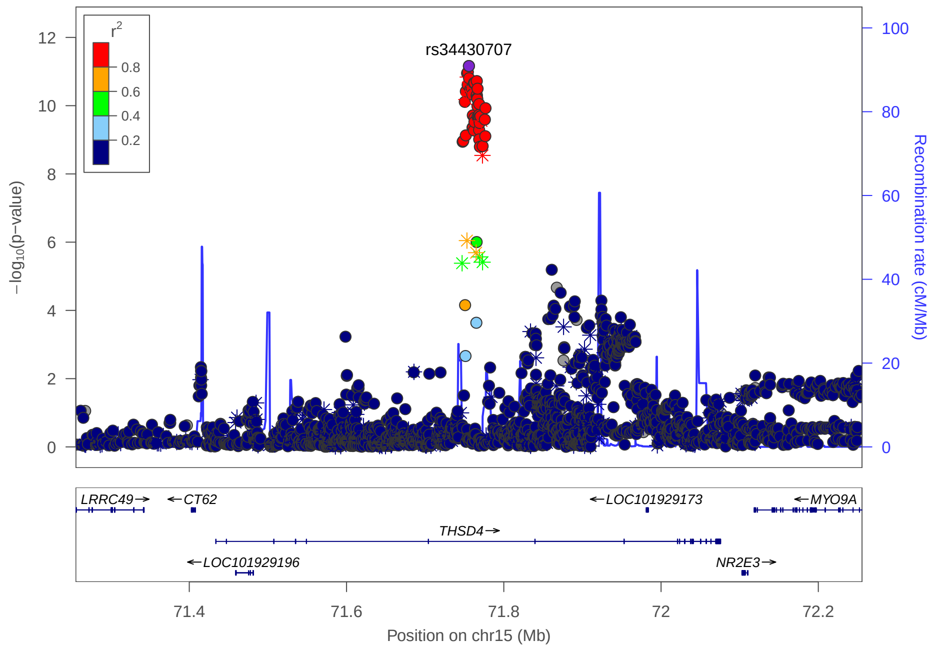

r2
0.8
0.6
0.4
0.2
rs34430707
−log10(p−value)
Recombination rate (cM/Mb)
LRRC49
CT62
LOC101929173
MYO9A
THSD4
LOC101929196
NR2E3
Position on chr15 (Mb)

### Multi-peak locus 22: rs112087864

**CHR15** - See final page for captions

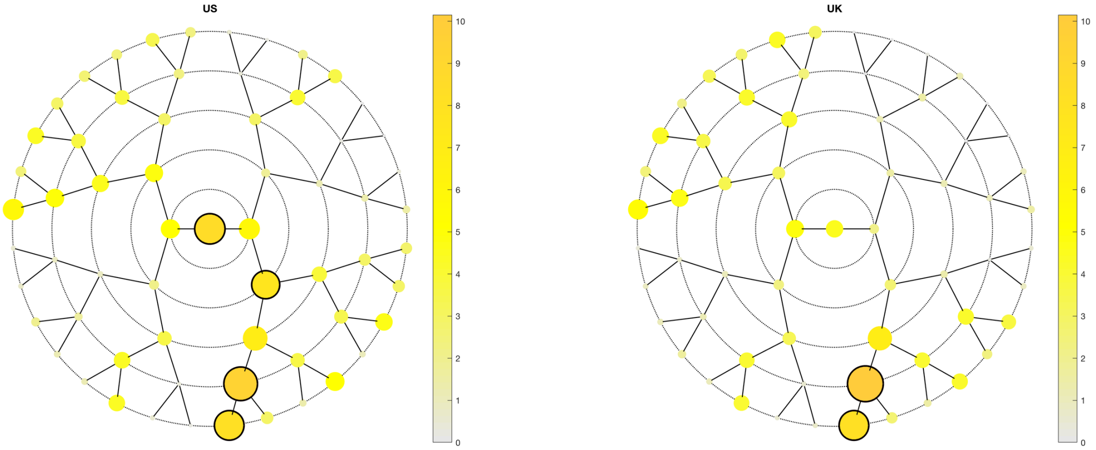

A

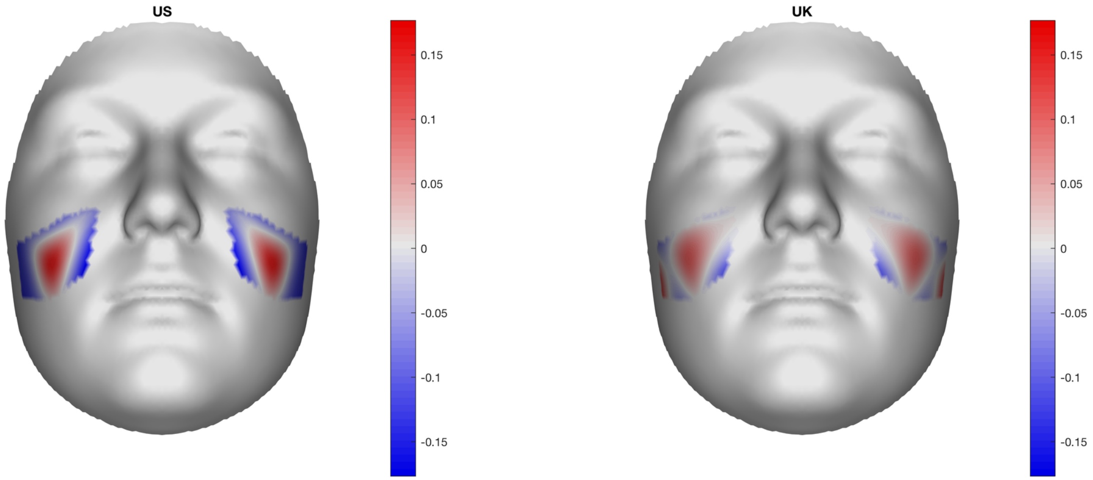

B

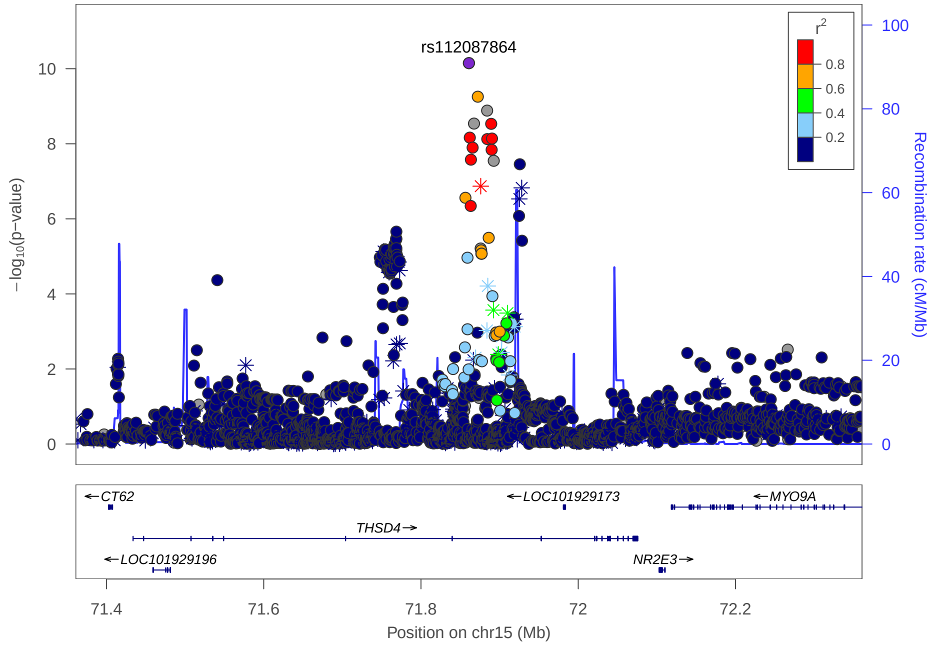

C

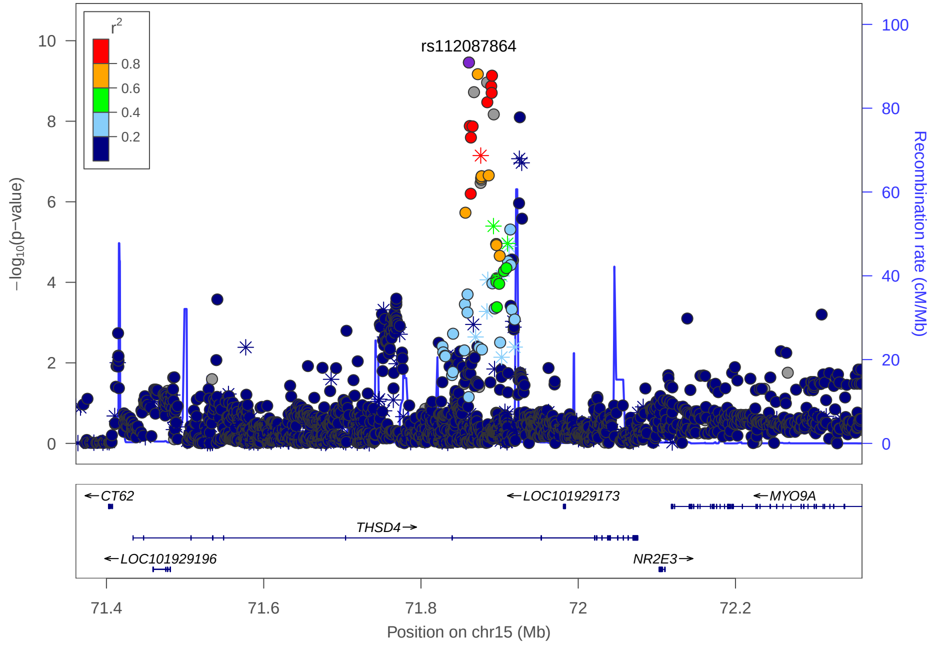

B

### Multi-peak locus 23: rs7218433

**CHR17** - See final page for captions

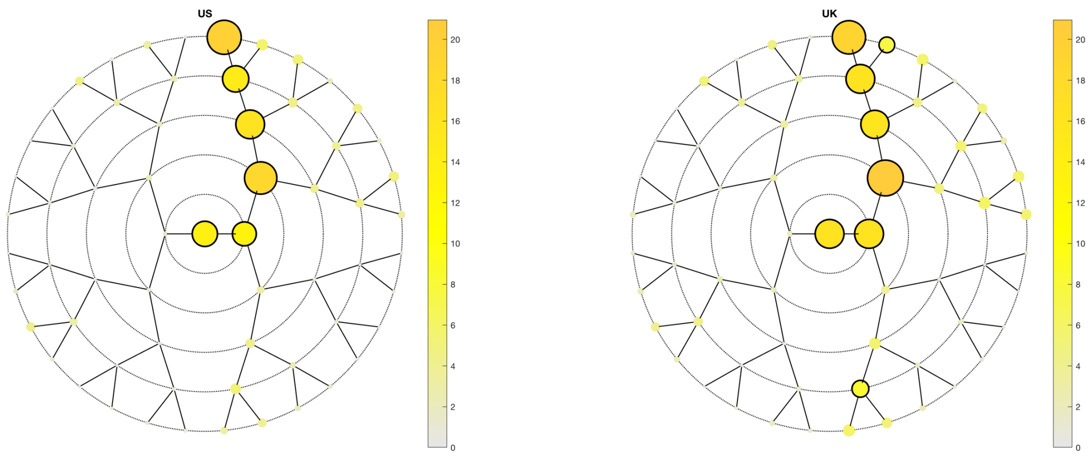

A

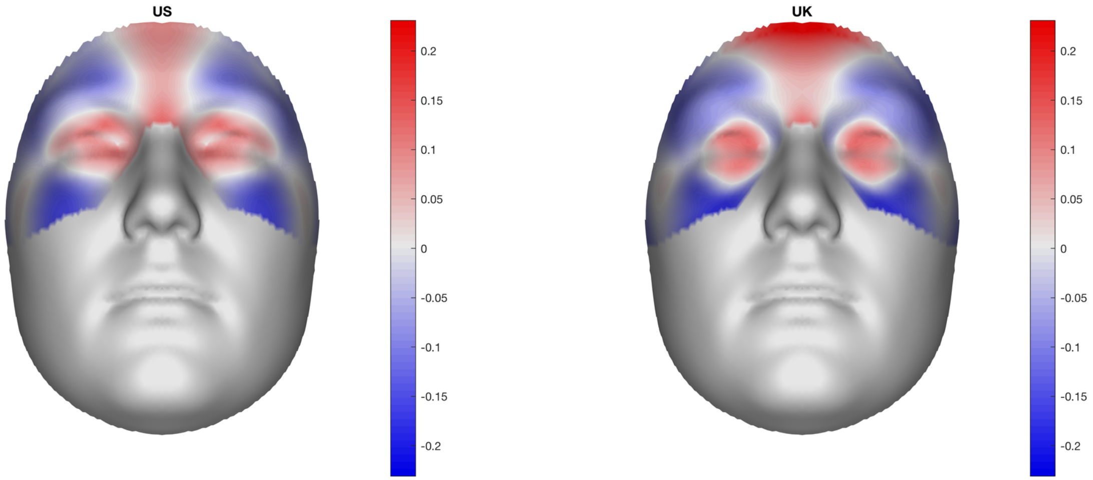

B

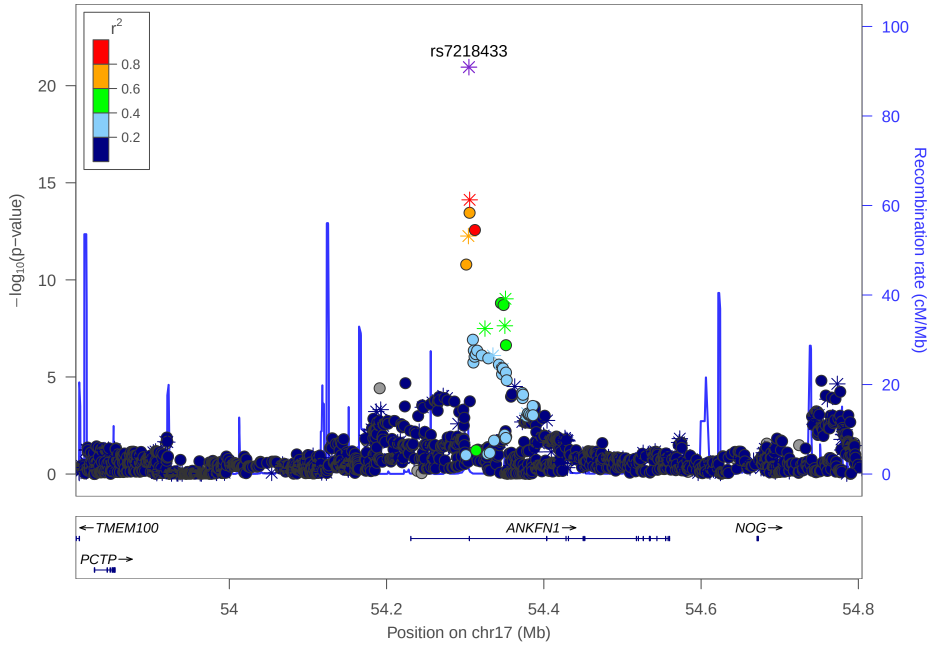

C

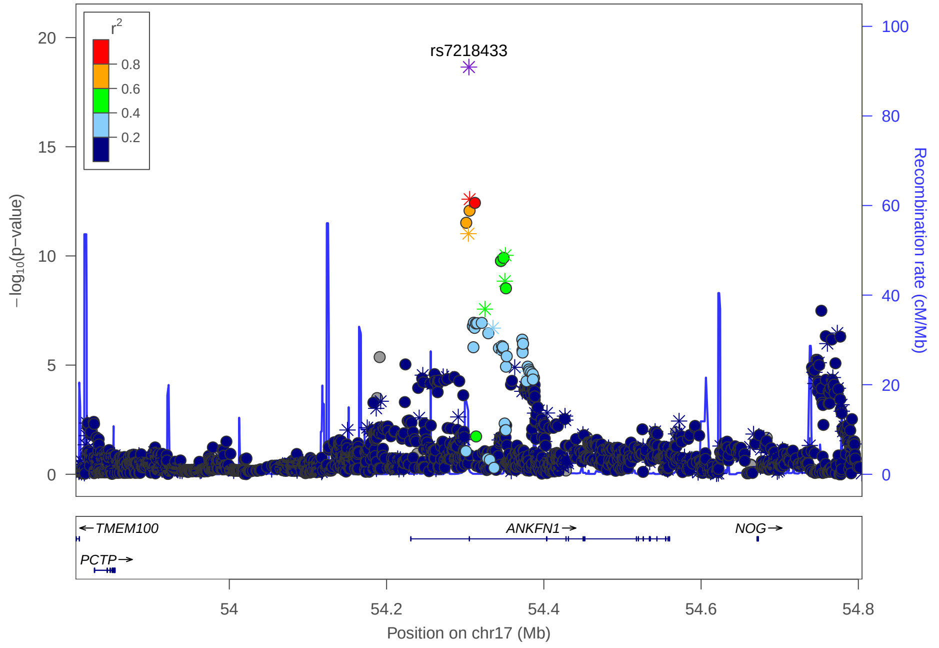

### Multi-peak locus 23: rs227727

**CHR17** - See final page for captions

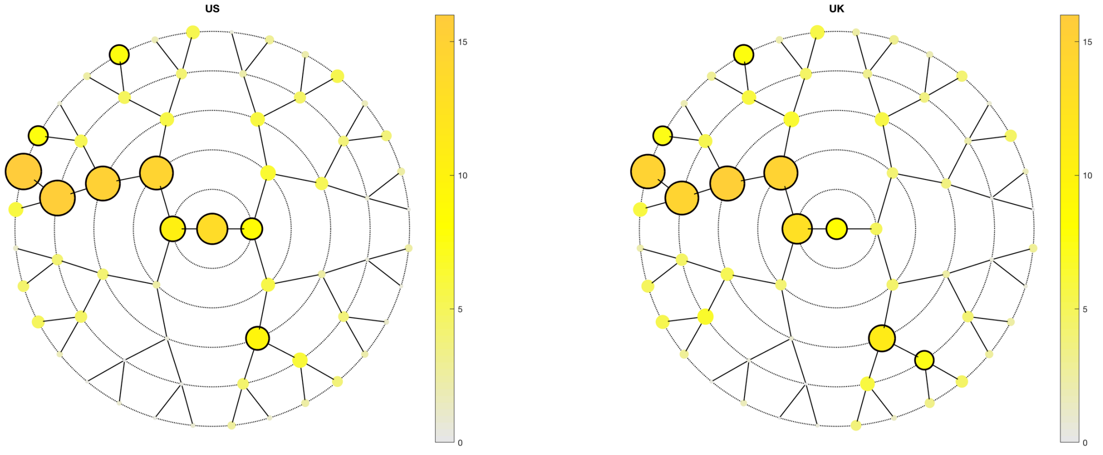

**A**

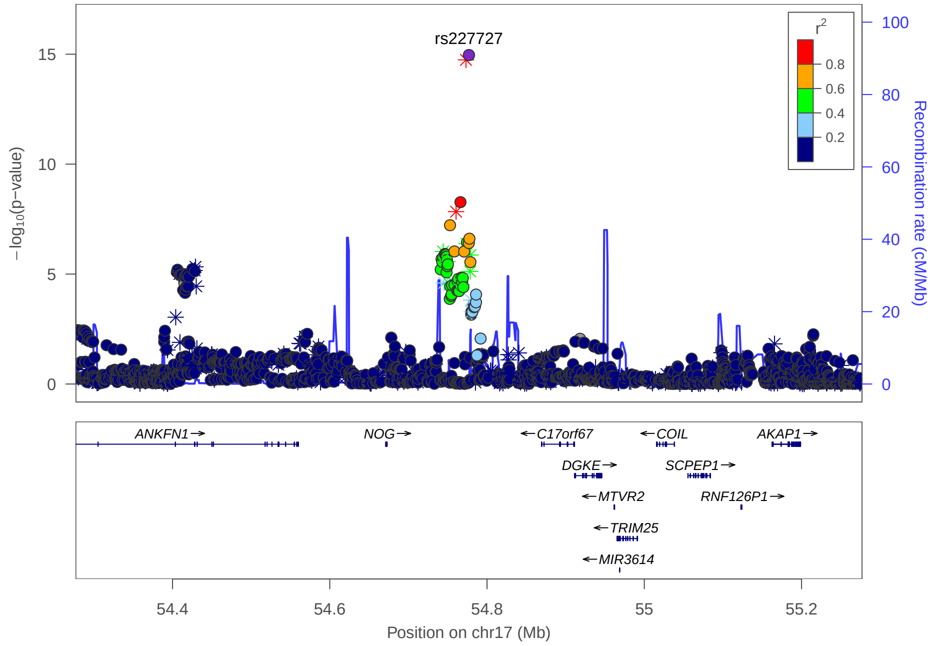

**C**

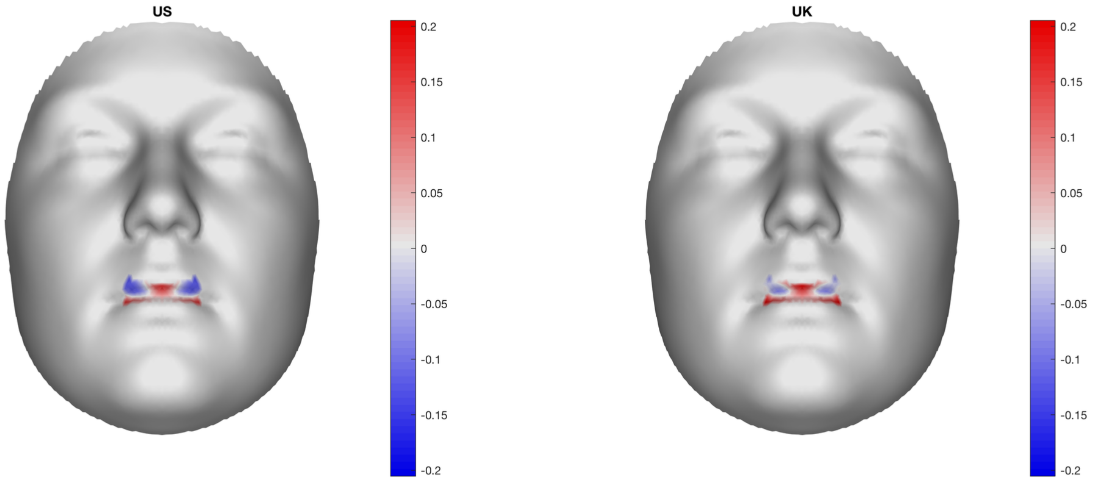

**B**

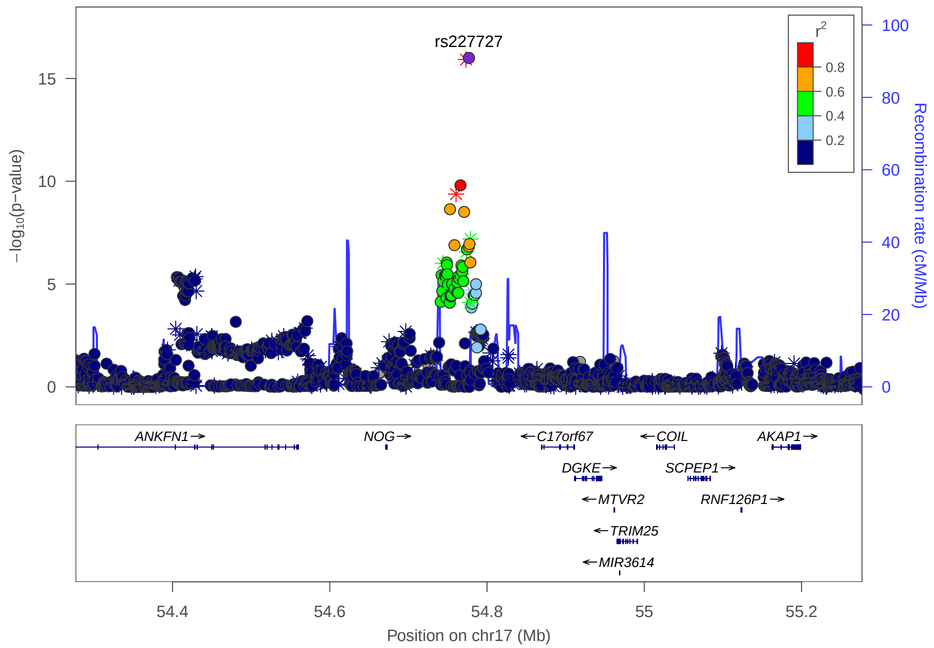

### Multi-peak locus 24: rs6047635

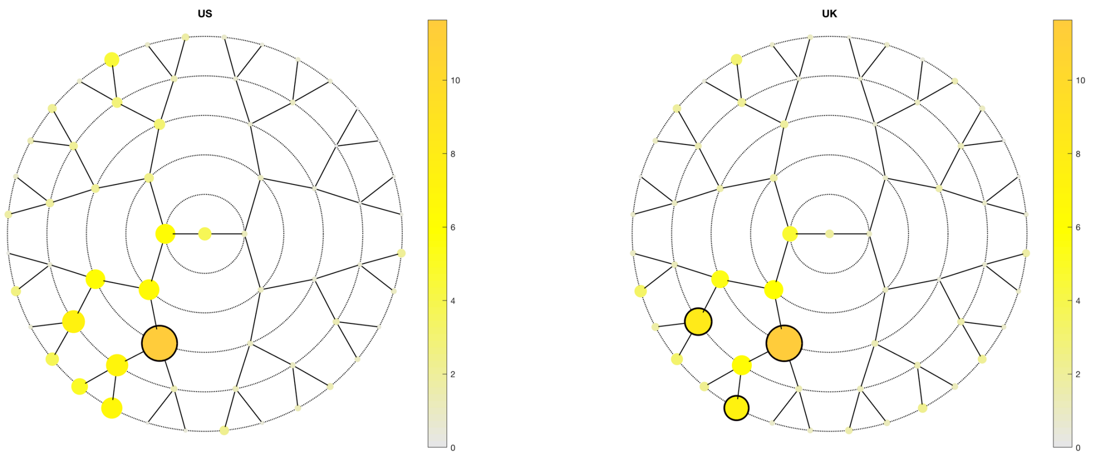

**A**

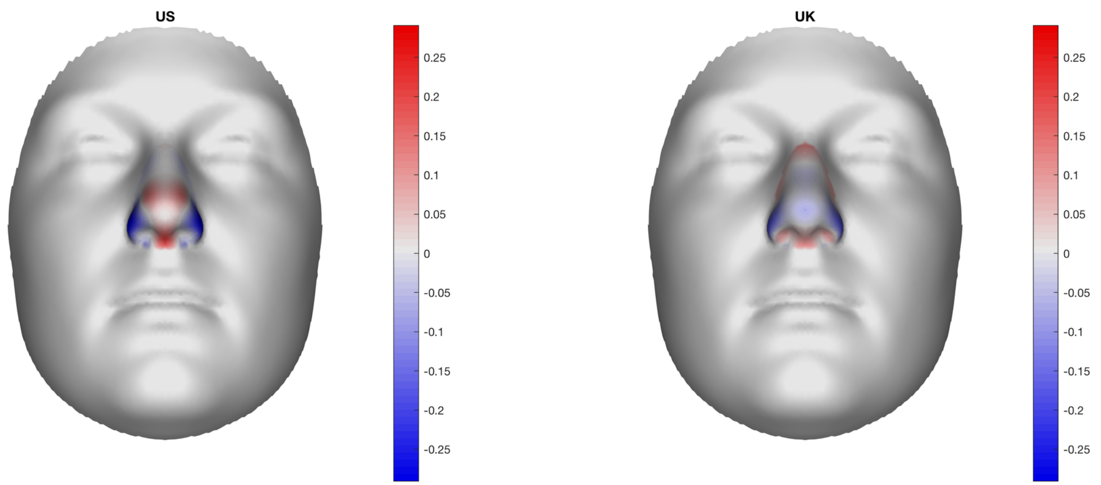

**B**

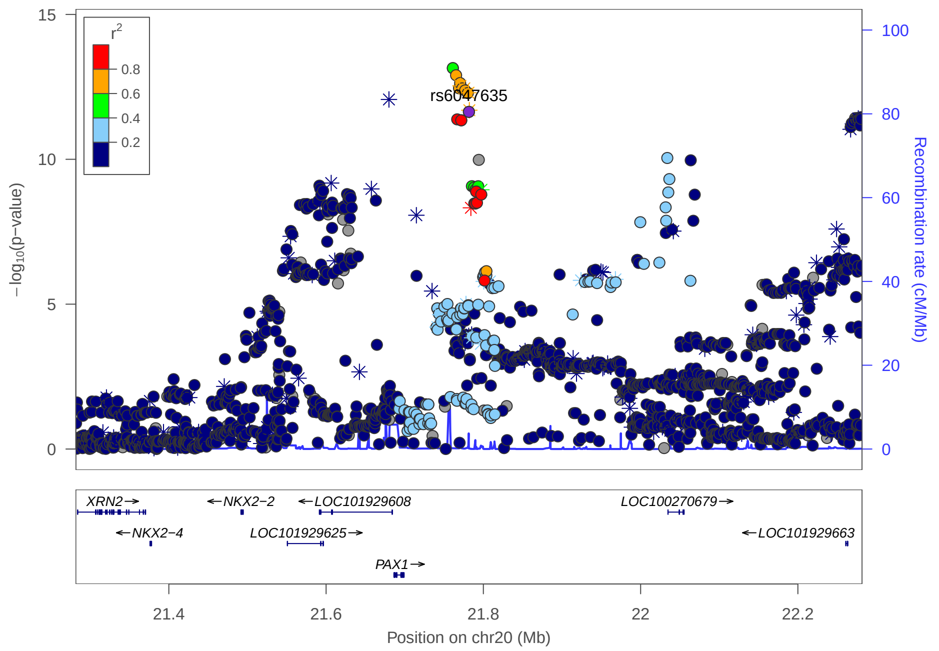

**C**

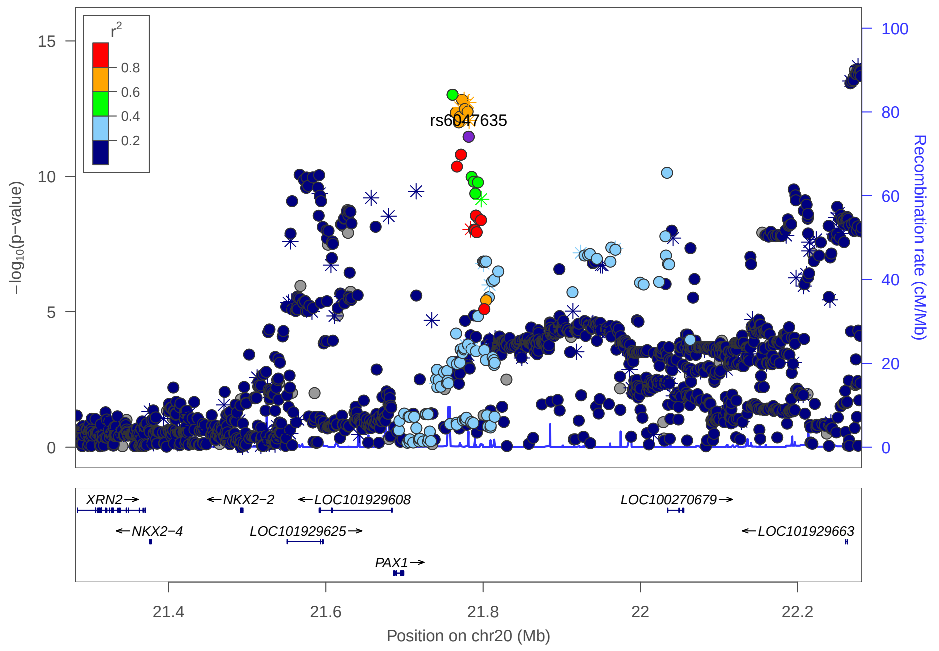

### Multi-peak locus 24: rs6047637

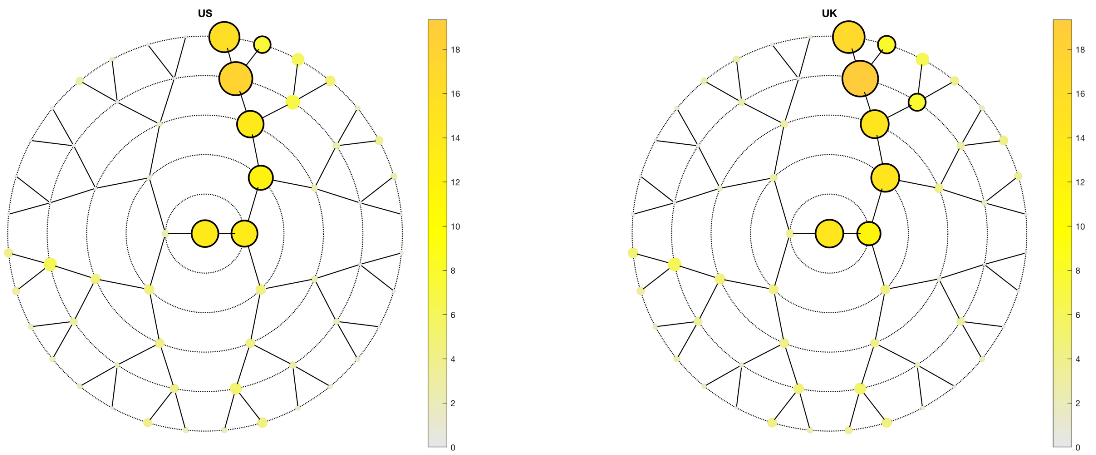

**A**

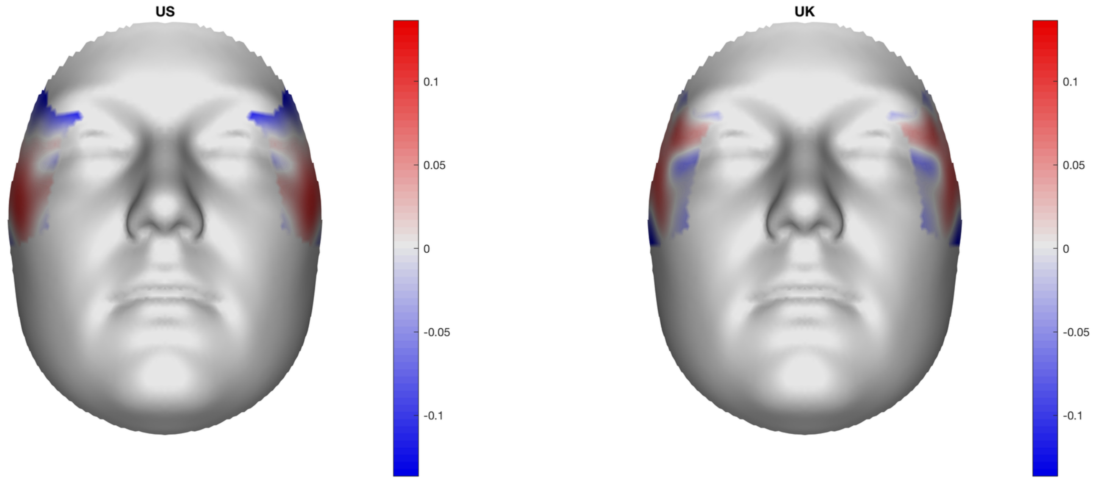

**B**

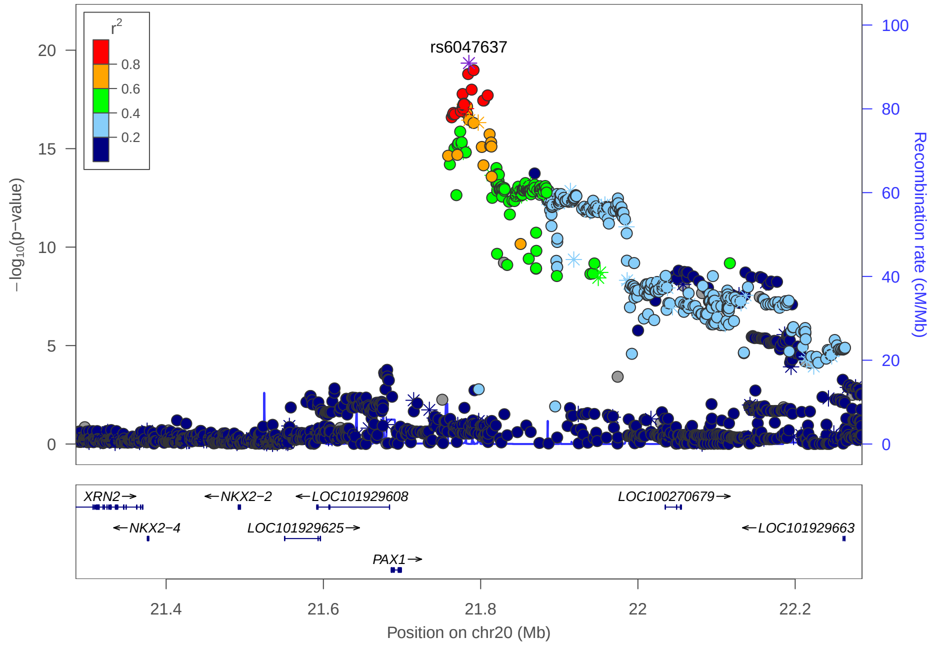

**C**

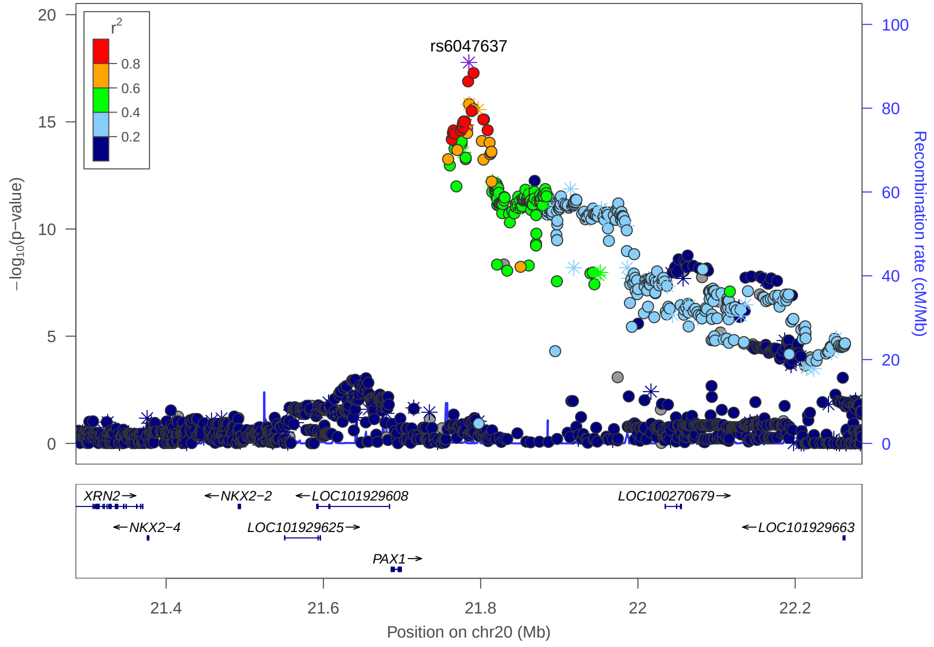

Overview multi-peak loci. **A.** $-\log_{10}$(p-value) of the meta-analysis p-values per facial segment in $\text{Meta}_{\text{US}}$ and $\text{Meta}_{\text{UK}}$. Black-encircled facial segments have reached a genome-wide p-value ($p = 5.10^{-8}$). **B.** The normal displacement (displacement in the direction locally normal to the facial surface) in each quasi-landmark of the facial segment reaching the lowest p-value in $\text{Meta}_{\text{US}}$ and $\text{Meta}_{\text{UK}}$ going from the minor to the major allele SNP variant. Blue, inward depression; red, outward protrusion. **C.** LocusZoom plots in in $\text{Meta}_{\text{US}}$ (top) and $\text{Meta}_{\text{UK}}$ (bottom). Points are color coded based on linkage disequilibrium ($r^2$) in Europeans. The asterisks represent genotyped SNPs, the circles represent imputed SNPs.
